## Supplementary file 7 for "High-Throughput Library Transgenesis in *Caenorhabditis elegans* via Transgenic Arrays Resulting in Diversity of Integrated Sequences (TARDIS)"

PX786 fxEx23[5’ΔHygR::Intron5'::Read1::ACACTATTTCATCTC::Read2::Intron3'::3’ΔHygR + 5’ΔHygR::Intron5'::Read1::CTCCCTTATCATTGC::Read2::Intron3'::3’ΔHygR + 5’ΔHygR::Intron5'::Read1::CACCTATTTCACAAC::Read2::Intron3'::3’ΔHygR + 5’ΔHygR::Intron5'::Read1::GCTCATTCTGACGTA::Read2::Intron3'::3’ΔHygR + 5’ΔHygR::Intron5'::Read1::TATCGGTTTCAGTCA::Read2::Intron3'::3’ΔHygR + 5’ΔHygR::Intron5'::Read1::TCCCCATGTCACTCA::Read2::Intron3'::3’ΔHygR + 5’ΔHygR::Intron5'::Read1::ATACATTCTTACACT::Read2::Intron3'::3’ΔHygR + 5’ΔHygR::Intron5'::Read1::TTTCCTTGTCAGATT::Read2::Intron3'::3’ΔHygR + 5’ΔHygR::Intron5'::Read1::CCTCTGTATTATCCC::Read2::Intron3'::3’ΔHygR + 5’ΔHygR::Intron5'::Read1::TACCCCTTTTAAACT::Read2::Intron3'::3’ΔHygR + 5’ΔHygR::Intron5'::Read1::TTGCTCTCTTATCTG::Read2::Intron3'::3’ΔHygR + 5’ΔHygR::Intron5'::Read1::CGCCATTCTTAATTT::Read2::Intron3'::3’ΔHygR + 5’ΔHygR::Intron5'::Read1::GAGCAATTTAATCAT::Read2::Intron3'::3’ΔHygR + 5’ΔHygR::Intron5'::Read1::TCGCGTTCTCATTCA::Read2::Intron3'::3’ΔHygR + 5’ΔHygR::Intron5'::Read1::AACCTTTATCATGGA::Read2::Intron3'::3’ΔHygR + 5’ΔHygR::Intron5'::Read1::ACCCGTTTTAACTCT::Read2::Intron3'::3’ΔHygR + 5’ΔHygR::Intron5'::Read1::ACACTCTATCACCCC::Read2::Intron3'::3’ΔHygR + 5’ΔHygR::Intron5'::Read1::TAACGGTATGAATAA::Read2::Intron3'::3’ΔHygR + 5’ΔHygR::Intron5'::Read1::TATCTTTGTAACATA::Read2::Intron3'::3’ΔHygR + 5’ΔHygR::Intron5'::Read1::TACCAATTTGACCAT::Read2::Intron3'::3’ΔHygR + 5’ΔHygR::Intron5'::Read1::CGCCAATTTAACTAC::Read2::Intron3'::3’ΔHygR + 5’ΔHygR::Intron5'::Read1::AGGCTTTGTCAAACG::Read2::Intron3'::3’ΔHygR + 5’ΔHygR::Intron5'::Read1::GCACCATCTGATGCC::Read2::Intron3'::3’ΔHygR + 5’ΔHygR::Intron5'::Read1::TCACATTTTAACTCA::Read2::Intron3'::3’ΔHygR + 5’ΔHygR::Intron5'::Read1::CCACCATTTTATTTG::Read2::Intron3'::3’ΔHygR + 5’ΔHygR::Intron5'::Read1::ACTCTATGTCATCTA::Read2::Intron3'::3’ΔHygR + 5’ΔHygR::Intron5'::Read1::ACCCGGTATAACTTC::Read2::Intron3'::3’ΔHygR + 5’ΔHygR::Intron5'::Read1::TGCCCTTTTCAGTAC::Read2::Intron3'::3’ΔHygR + 5’ΔHygR::Intron5'::Read1::ACCCGCTTTTAGTTA::Read2::Intron3'::3’ΔHygR + 5’ΔHygR::Intron5'::Read1::CCGCTGTCTTAAATG::Read2::Intron3'::3’ΔHygR + 5’ΔHygR::Intron5'::Read1::TTCCTTTTTAACCCT::Read2::Intron3'::3’ΔHygR + 5’ΔHygR::Intron5'::Read1::TAGCAATGTTACGAA::Read2::Intron3'::3’ΔHygR + 5’ΔHygR::Intron5'::Read1::CACCCCTGTGACTAC::Read2::Intron3'::3’ΔHygR + 5’ΔHygR::Intron5'::Read1::ACTCGGTTTCAACCA::Read2::Intron3'::3’ΔHygR + 5’ΔHygR::Intron5'::Read1::GTACCGTTTAACGTC::Read2::Intron3'::3’ΔHygR + 5’ΔHygR::Intron5'::Read1::CCTCTTTGTAAATCA::Read2::Intron3'::3’ΔHygR + 5’ΔHygR::Intron5'::Read1::TTAAATTATCACATG::Read2::Intron3'::3’ΔHygR + 5’ΔHygR::Intron5'::Read1::AAACCATGTTAAACC::Read2::Intron3'::3’ΔHygR + 5’ΔHygR::Intron5'::Read1::CTACCCTATCATCAC::Read2::Intron3'::3’ΔHygR + 5’ΔHygR::Intron5'::Read1::GTCCTGTTTCATCCC::Read2::Intron3'::3’ΔHygR + 5’ΔHygR::Intron5'::Read1::ATTCAATTTAACAAT::Read2::Intron3'::3’ΔHygR + 5’ΔHygR::Intron5'::Read1::TTACCGTTTTACTAT::Read2::Intron3'::3’ΔHygR + 5’ΔHygR::Intron5'::Read1::TGTCGGTTTCAATCC::Read2::Intron3'::3’ΔHygR + 5’ΔHygR::Intron5'::Read1::AGGGTTAAAAAGGAA::Read2::Intron3'::3’ΔHygR + 5’ΔHygR::Intron5'::Read1::GGGATAATACAGAGG::Read2::Intron3'::3’ΔHygR + 5’ΔHygR::Intron5'::Read1::AGTGTAAGAATGTAT::Read2::Intron3'::3’ΔHygR + 5’ΔHygR::Intron5'::Read1::TTATTCATACCGTTA::Read2::Intron3'::3’ΔHygR + 5’ΔHygR::Intron5'::Read1::GGATTGAAACCGACA::Read2::Intron3'::3’ΔHygR + 5’ΔHygR::Intron5'::Read1::TACGTCAGAATGAGC::Read2::Intron3'::3’ΔHygR + 5’ΔHygR::Intron5'::Read1::AATCTCTATTATCAC::Read2::Intron3'::3’ΔHygR + 5’ΔHygR::Intron5'::Read1::AGAGTTAAAACGGGT::Read2::Intron3'::3’ΔHygR + 5’ΔHygR::Intron5'::Read1::GGGGTGATAGAGTGT::Read2::Intron3'::3’ΔHygR + 5’ΔHygR::Intron5'::Read1::TATGTTACAAAGATA::Read2::Intron3'::3’ΔHygR + 5’ΔHygR::Intron5'::Read1::ATGATTAAATTGCTC::Read2::Intron3'::3’ΔHygR + 5’ΔHygR::Intron5'::Read1::CAGATAAGAGAGCAA::Read2::Intron3'::3’ΔHygR + 5’ΔHygR::Intron5'::Read1::GGTTTAACATGGTTT::Read2::Intron3'::3’ΔHygR + 5’ΔHygR::Intron5'::Read1::GTTGTGAAATAGGTG::Read2::Intron3'::3’ΔHygR + 5’ΔHygR::Intron5'::Read1::TGAGTGACATGGGGA::Read2::Intron3'::3’ΔHygR + 5’ΔHygR::Intron5'::Read1::TGAATGAGAACGCGA::Read2::Intron3'::3’ΔHygR + 5’ΔHygR::Intron5'::Read1::TGACTGAAACCGATA::Read2::Intron3'::3’ΔHygR + 5’ΔHygR::Intron5'::Read1::AACTCTATGGCACAG::Read2::Intron3'::3’ΔHygR + 5’ΔHygR::Intron5'::Read1::CATTTAAGACAGCGG::Read2::Intron3'::3’ΔHygR + 5’ΔHygR::Intron5'::Read1::TTCGTAACATTGCTA::Read2::Intron3'::3’ΔHygR + 5’ΔHygR::Intron5'::Read1::AAATTAAGAATGGCG::Read2::Intron3'::3’ΔHygR + 5’ΔHygR::Intron5'::Read1::CATCTTGTTCCCATC::Read2::Intron3'::3’ΔHygR + 5’ΔHygR::Intron5'::Read1::GACGTTAAACGGTAC::Read2::Intron3'::3’ΔHygR + 5’ΔHygR::Intron5'::Read1::GTACTGAAAAGGGCA::Read2::Intron3'::3’ΔHygR + 5’ΔHygR::Intron5'::Read1::ATCCAATCTTAAATT::Read2::Intron3'::3’ΔHygR + 5’ΔHygR::Intron5'::Read1::GAAGTTATACCGGGT::Read2::Intron3'::3’ΔHygR +

5’ΔHygR::Intron5'::Read1::GAGATGAAATAGTGT::Read2::Intron3'::3’ΔHygR + 5’ΔHygR::Intron5'::Read1::GGCATCAGATGGTGC::Read2::Intron3'::3’ΔHygR + 5’ΔHygR::Intron5'::Read1::TAACTAAAAGCGGGT::Read2::Intron3'::3’ΔHygR + 5’ΔHygR::Intron5'::Read1::TGATTTACAAAGAGG::Read2::Intron3'::3’ΔHygR + 5’ΔHygR::Intron5'::Read1::TGGTTGAAACCGAGT::Read2::Intron3'::3’ΔHygR + 5’ΔHygR::Intron5'::Read1::AGTTTAAAAGGGGTA::Read2::Intron3'::3’ΔHygR + 5’ΔHygR::Intron5'::Read1::CAAATAAAATGGTGG::Read2::Intron3'::3’ΔHygR + 5’ΔHygR::Intron5'::Read1::GTAGTCACAGGGGTG::Read2::Intron3'::3’ΔHygR + 5’ΔHygR::Intron5'::Read1::AAACTCTATAAACAC::Read2::Intron3'::3’ΔHygR + 5’ΔHygR::Intron5'::Read1::AATAGTGTCACATGA::Read2::Intron3'::3’ΔHygR + 5’ΔHygR::Intron5'::Read1::AATCTGACAAGGAAA::Read2::Intron3'::3’ΔHygR + 5’ΔHygR::Intron5'::Read1::ACACAATGTCAAAAA::Read2::Intron3'::3’ΔHygR + 5’ΔHygR::Intron5'::Read1::ACCCATTCTAAATAA::Read2::Intron3'::3’ΔHygR + 5’ΔHygR::Intron5'::Read1::ACTCTTTATAATACA::Read2::Intron3'::3’ΔHygR + 5’ΔHygR::Intron5'::Read1::ATCCAATATAATATC::Read2::Intron3'::3’ΔHygR + 5’ΔHygR::Intron5'::Read1::ATGGTCAAATTGGTA::Read2::Intron3'::3’ΔHygR + 5’ΔHygR::Intron5'::Read1::CATAGTTTTCCTTAT::Read2::Intron3'::3’ΔHygR + 5’ΔHygR::Intron5'::Read1::CATGTGATAATTTAA::Read2::Intron3'::3’ΔHygR + 5’ΔHygR::Intron5'::Read1::GATAATATACCTAGT::Read2::Intron3'::3’ΔHygR + 5’ΔHygR::Intron5'::Read1::GTAGTTAAATTGGCG::Read2::Intron3'::3’ΔHygR + 5’ΔHygR::Intron5'::Read1::GTGATGATAGGGTAG::Read2::Intron3'::3’ΔHygR + 5’ΔHygR::Intron5'::Read1::TGAGTTAAAATGTGA::Read2::Intron3'::3’ΔHygR + 5’ΔHygR::Intron5'::Read1::AAACCATGTGACAAA::Read2::Intron3'::3’ΔHygR + 5’ΔHygR::Intron5'::Read1::AAACCTTTTTATGTT::Read2::Intron3'::3’ΔHygR + 5’ΔHygR::Intron5'::Read1::AAACGCTTTAAGTAA::Read2::Intron3'::3’ΔHygR + 5’ΔHygR::Intron5'::Read1::AACCACTTTAAGCTT::Read2::Intron3'::3’ΔHygR + 5’ΔHygR::Intron5'::Read1::AACCCTTATAAATTC::Read2::Intron3'::3’ΔHygR + 5’ΔHygR::Intron5'::Read1::AATCAATATTAAGAA::Read2::Intron3'::3’ΔHygR + 5’ΔHygR::Intron5'::Read1::AATCCATCTCATACA::Read2::Intron3'::3’ΔHygR + 5’ΔHygR::Intron5'::Read1::ACACAATATAAAACA::Read2::Intron3'::3’ΔHygR + 5’ΔHygR::Intron5'::Read1::ACACAATATAAAACC::Read2::Intron3'::3’ΔHygR + 5’ΔHygR::Intron5'::Read1::ACACCATATAATTAT::Read2::Intron3'::3’ΔHygR + 5’ΔHygR::Intron5'::Read1::ACCCAATCTAAATAA::Read2::Intron3'::3’ΔHygR + 5’ΔHygR::Intron5'::Read1::ACCCGATATAATTAA::Read2::Intron3'::3’ΔHygR + 5’ΔHygR::Intron5'::Read1::ATACCTTATCATATC::Read2::Intron3'::3’ΔHygR + 5’ΔHygR::Intron5'::Read1::ATCCAATATAAAAAC::Read2::Intron3'::3’ΔHygR + 5’ΔHygR::Intron5'::Read1::ATTGTTAAATTGAAT::Read2::Intron3'::3’ΔHygR + 5’ΔHygR::Intron5'::Read1::CAACATTATAATACC::Read2::Intron3'::3’ΔHygR + 5’ΔHygR::Intron5'::Read1::CAACCATATAATTTC::Read2::Intron3'::3’ΔHygR + 5’ΔHygR::Intron5'::Read1::CCTCATTTTTACATC::Read2::Intron3'::3’ΔHygR + 5’ΔHygR::Intron5'::Read1::GCAATGATAAGGGAG::Read2::Intron3'::3’ΔHygR + 5’ΔHygR::Intron5'::Read1::TAAGTTATATCGTAC::Read2::Intron3'::3’ΔHygR + 5’ΔHygR::Intron5'::Read1::TACCTTCTGTCTGTT::Read2::Intron3'::3’ΔHygR + 5’ΔHygR::Intron5'::Read1::TAGATGACATAGAGT::Read2::Intron3'::3’ΔHygR + 5’ΔHygR::Intron5'::Read1::TCACTCTTTTACCTG::Read2::Intron3'::3’ΔHygR + 5’ΔHygR::Intron5'::Read1::TTACCTTGTTATCAC::Read2::Intron3'::3’ΔHygR + hsp-16.41p::Cas9(dpiRNA)::tbb-2 3’ UTR + prsp-27::NeoR::unc- 54 3' UTR + U6p:: GCGAAGTGACGGTAGACCGT]; fxSi47[ prsp-0p::5’ΔHygR:: GCGAAGTGACGGTAGACCGT ::3’ΔHygR::unc-54 3’::LoxP]

PX816 fxEx25 [5’ΔHygR::Intron5'::Read1::TAACCGTATTAGCTC::Read2::Intron3'::3’ΔHygR + 5’ΔHygR::Intron5'::Read1::AGTCCTTTTGACGTA::Read2::Intron3'::3’ΔHygR + 5’ΔHygR::Intron5'::Read1::GAACTATGTGACGTT::Read2::Intron3'::3’ΔHygR + 5’ΔHygR::Intron5'::Read1::GCATTCAGAATGCTT::Read2::Intron3'::3’ΔHygR + 5’ΔHygR::Intron5'::Read1::TGCCTATTTCACCTC::Read2::Intron3'::3’ΔHygR + 5’ΔHygR::Intron5'::Read1::ACGCTCTATAAGTTC::Read2::Intron3'::3’ΔHygR + 5’ΔHygR::Intron5'::Read1::CAGCGATATAACCTA::Read2::Intron3'::3’ΔHygR + 5’ΔHygR::Intron5'::Read1::TTGCCATTTTATCTT::Read2::Intron3'::3’ΔHygR + 5’ΔHygR::Intron5'::Read1::GCCCTCTTTCATCGG::Read2::Intron3'::3’ΔHygR + 5’ΔHygR::Intron5'::Read1::TGCCAGTCTCAATAA::Read2::Intron3'::3’ΔHygR + 5’ΔHygR::Intron5'::Read1::ATTCATTTTCAACGT::Read2::Intron3'::3’ΔHygR + 5’ΔHygR::Intron5'::Read1::CCCCCATCTAACGAT::Read2::Intron3'::3’ΔHygR + 5’ΔHygR::Intron5'::Read1::CCCCCGTTTTAATCT::Read2::Intron3'::3’ΔHygR + 5’ΔHygR::Intron5'::Read1::GCACATTCTAATACA::Read2::Intron3'::3’ΔHygR + 5’ΔHygR::Intron5'::Read1::CATCCGTATGAAGTT::Read2::Intron3'::3’ΔHygR + 5’ΔHygR::Intron5'::Read1::CTTCTGTGTGATCAC::Read2::Intron3'::3’ΔHygR + 5’ΔHygR::Intron5'::Read1::CAACGGTTTCAGCAT::Read2::Intron3'::3’ΔHygR + 5’ΔHygR::Intron5'::Read1::CCACCCTATCATGCC::Read2::Intron3'::3’ΔHygR + 5’ΔHygR::Intron5'::Read1::GAGCATTATAATCGT::Read2::Intron3'::3’ΔHygR +

5’ΔHygR::Intron5'::Read1::GACCAGTCTCACTTG::Read2::Intron3'::3’ΔHygR + 5’ΔHygR::Intron5'::Read1::AGCCATTTTAACTCC::Read2::Intron3'::3’ΔHygR + 5’ΔHygR::Intron5'::Read1::GGGCACTCTGACTCA::Read2::Intron3'::3’ΔHygR + 5’ΔHygR::Intron5'::Read1::TGCCTGTGTAATCTC::Read2::Intron3'::3’ΔHygR + 5’ΔHygR::Intron5'::Read1::CCTCGTTCTGATCCA::Read2::Intron3'::3’ΔHygR + 5’ΔHygR::Intron5'::Read1::TTCCTTTCTCAGGCG::Read2::Intron3'::3’ΔHygR + 5’ΔHygR::Intron5'::Read1::CCTCAGTCTTACCTC::Read2::Intron3'::3’ΔHygR + 5’ΔHygR::Intron5'::Read1::GATCCCTTTTAGGTA::Read2::Intron3'::3’ΔHygR + 5’ΔHygR::Intron5'::Read1::CTTCCATCTCATCAC::Read2::Intron3'::3’ΔHygR + 5’ΔHygR::Intron5'::Read1::TCACGCTATAACGCC::Read2::Intron3'::3’ΔHygR + 5’ΔHygR::Intron5'::Read1::CATCGATGTAATACT::Read2::Intron3'::3’ΔHygR + 5’ΔHygR::Intron5'::Read1::GAGCGCTATAATACA::Read2::Intron3'::3’ΔHygR + 5’ΔHygR::Intron5'::Read1::ATCCAGTTTCATCCG::Read2::Intron3'::3’ΔHygR + 5’ΔHygR::Intron5'::Read1::CGCCCATATTACCTA::Read2::Intron3'::3’ΔHygR + 5’ΔHygR::Intron5'::Read1::TCGCTGTATGAGGAT::Read2::Intron3'::3’ΔHygR + 5’ΔHygR::Intron5'::Read1::CTTCGCTGTCATATT::Read2::Intron3'::3’ΔHygR + 5’ΔHygR::Intron5'::Read1::CAACTTTTTCAATCG::Read2::Intron3'::3’ΔHygR + 5’ΔHygR::Intron5'::Read1::CTTCAGTGTCAATGT::Read2::Intron3'::3’ΔHygR + 5’ΔHygR::Intron5'::Read1::GTGCATTTTGATATG::Read2::Intron3'::3’ΔHygR + 5’ΔHygR::Intron5'::Read1::AGACGATTTCAAGGA::Read2::Intron3'::3’ΔHygR + 5’ΔHygR::Intron5'::Read1::CCACCTTTTCACTCG::Read2::Intron3'::3’ΔHygR + 5’ΔHygR::Intron5'::Read1::ACTCACTATGACTGG::Read2::Intron3'::3’ΔHygR + 5’ΔHygR::Intron5'::Read1::TAGCATTATTAGCCG::Read2::Intron3'::3’ΔHygR + 5’ΔHygR::Intron5'::Read1::CAACTTTTTAAACGA::Read2::Intron3'::3’ΔHygR + 5’ΔHygR::Intron5'::Read1::GGCCTATTTAACTTA::Read2::Intron3'::3’ΔHygR + 5’ΔHygR::Intron5'::Read1::CGCCTCTCTCAGTTT::Read2::Intron3'::3’ΔHygR + 5’ΔHygR::Intron5'::Read1::GTGCCGTATCAACTC::Read2::Intron3'::3’ΔHygR + 5’ΔHygR::Intron5'::Read1::AAGCCTTGTCATCAT::Read2::Intron3'::3’ΔHygR + 5’ΔHygR::Intron5'::Read1::TTTCGTTCTGAATCC::Read2::Intron3'::3’ΔHygR + 5’ΔHygR::Intron5'::Read1::TAACTGTCTGAATAC::Read2::Intron3'::3’ΔHygR + 5’ΔHygR::Intron5'::Read1::CATCTTTCTCAACTC::Read2::Intron3'::3’ΔHygR + 5’ΔHygR::Intron5'::Read1::AGTCTATTTCAAATT::Read2::Intron3'::3’ΔHygR + 5’ΔHygR::Intron5'::Read1::CAACTCTTTTACCGT::Read2::Intron3'::3’ΔHygR + 5’ΔHygR::Intron5'::Read1::AGCCCGTTTTACGAG::Read2::Intron3'::3’ΔHygR + 5’ΔHygR::Intron5'::Read1::AACCGCTGTAACATC::Read2::Intron3'::3’ΔHygR + 5’ΔHygR::Intron5'::Read1::CGTCACTTTAACATC::Read2::Intron3'::3’ΔHygR + 5’ΔHygR::Intron5'::Read1::CAGCCATCTCAATTC::Read2::Intron3'::3’ΔHygR + 5’ΔHygR::Intron5'::Read1::CTCCGCTATTACAAA::Read2::Intron3'::3’ΔHygR + 5’ΔHygR::Intron5'::Read1::GGGCCATTTCATCTT::Read2::Intron3'::3’ΔHygR + 5’ΔHygR::Intron5'::Read1::CTGCCGTGTTACATC::Read2::Intron3'::3’ΔHygR + 5’ΔHygR::Intron5'::Read1::GCCCGCTCTGAGTAT::Read2::Intron3'::3’ΔHygR + 5’ΔHygR::Intron5'::Read1::GTTCCATATCAGTTT::Read2::Intron3'::3’ΔHygR + 5’ΔHygR::Intron5'::Read1::GTTCTTTTTGAACCA::Read2::Intron3'::3’ΔHygR + 5’ΔHygR::Intron5'::Read1::CGTCACTTTCACCGC::Read2::Intron3'::3’ΔHygR + 5’ΔHygR::Intron5'::Read1::TTGCCATTTAAACCG::Read2::Intron3'::3’ΔHygR + 5’ΔHygR::Intron5'::Read1::TACCGTTATGATTCC::Read2::Intron3'::3’ΔHygR + 5’ΔHygR::Intron5'::Read1::AAACAGTCTCAACAA::Read2::Intron3'::3’ΔHygR + 5’ΔHygR::Intron5'::Read1::GTCCATTCTAACTCT::Read2::Intron3'::3’ΔHygR + 5’ΔHygR::Intron5'::Read1::GTCCACTCTAATTCA::Read2::Intron3'::3’ΔHygR + 5’ΔHygR::Intron5'::Read1::CACCCTTTTCAGTCT::Read2::Intron3'::3’ΔHygR + 5’ΔHygR::Intron5'::Read1::AATCGTTTTTAGCTA::Read2::Intron3'::3’ΔHygR + 5’ΔHygR::Intron5'::Read1::CAACGTTTTCATATC::Read2::Intron3'::3’ΔHygR + 5’ΔHygR::Intron5'::Read1::GCACACTCTGAGACC::Read2::Intron3'::3’ΔHygR + 5’ΔHygR::Intron5'::Read1::TCCCTGTCTCAATAA::Read2::Intron3'::3’ΔHygR + 5’ΔHygR::Intron5'::Read1::ACTCACTCTGAATCA::Read2::Intron3'::3’ΔHygR + 5’ΔHygR::Intron5'::Read1::GGACATTTTGAATTT::Read2::Intron3'::3’ΔHygR + 5’ΔHygR::Intron5'::Read1::TATCAATCTGAGCCC::Read2::Intron3'::3’ΔHygR + 5’ΔHygR::Intron5'::Read1::GACCCATTTTACTAC::Read2::Intron3'::3’ΔHygR + 5’ΔHygR::Intron5'::Read1::CAGCCTTTTTATACA::Read2::Intron3'::3’ΔHygR + 5’ΔHygR::Intron5'::Read1::CAACTGTCTCAAGAA::Read2::Intron3'::3’ΔHygR + 5’ΔHygR::Intron5'::Read1::GAACACTGTTAAAAA::Read2::Intron3'::3’ΔHygR + 5’ΔHygR::Intron5'::Read1::GTACTGTGTCACAAC::Read2::Intron3'::3’ΔHygR + 5’ΔHygR::Intron5'::Read1::TGACCTTTTGACCTC::Read2::Intron3'::3’ΔHygR + 5’ΔHygR::Intron5'::Read1::CGTCGTTCTGAGACC::Read2::Intron3'::3’ΔHygR + 5’ΔHygR::Intron5'::Read1::CCGCAGTTTCAATCA::Read2::Intron3'::3’ΔHygR + 5’ΔHygR::Intron5'::Read1::TGGCACTGTCATACA::Read2::Intron3'::3’ΔHygR + 5’ΔHygR::Intron5'::Read1::CTACAATCTAACCGT::Read2::Intron3'::3’ΔHygR + 5’ΔHygR::Intron5'::Read1::ACACCTTATGATAGA::Read2::Intron3'::3’ΔHygR + 5’ΔHygR::Intron5'::Read1::TGTCGCTCTTACTAA::Read2::Intron3'::3’ΔHygR + 5’ΔHygR::Intron5'::Read1::TAACAGTATTAAACC::Read2::Intron3'::3’ΔHygR +

5’ΔHygR::Intron5'::Read1::CGACCTTCTGAAACC::Read2::Intron3'::3’ΔHygR + 5’ΔHygR::Intron5'::Read1::GGCCTATTTGATTGA::Read2::Intron3'::3’ΔHygR + 5’ΔHygR::Intron5'::Read1::CCCCACTATCACTCG::Read2::Intron3'::3’ΔHygR + 5’ΔHygR::Intron5'::Read1::TGCCCATCTGAAAAG::Read2::Intron3'::3’ΔHygR + 5’ΔHygR::Intron5'::Read1::GCACACTATAACATT::Read2::Intron3'::3’ΔHygR + 5’ΔHygR::Intron5'::Read1::ATGCCCTCTCATGTC::Read2::Intron3'::3’ΔHygR + 5’ΔHygR::Intron5'::Read1::TACCTCTATTATTGG::Read2::Intron3'::3’ΔHygR + 5’ΔHygR::Intron5'::Read1::TTACGTTGTAACACG::Read2::Intron3'::3’ΔHygR + 5’ΔHygR::Intron5'::Read1::ATGCTTTCTTACGCT::Read2::Intron3'::3’ΔHygR + 5’ΔHygR::Intron5'::Read1::TCACGGTATAAATAG::Read2::Intron3'::3’ΔHygR + 5’ΔHygR::Intron5'::Read1::TCCCCCTATCAACAT::Read2::Intron3'::3’ΔHygR + 5’ΔHygR::Intron5'::Read1::GTACCATATTACGGC::Read2::Intron3'::3’ΔHygR + 5’ΔHygR::Intron5'::Read1::CTCCCCTCTAAATTT::Read2::Intron3'::3’ΔHygR + 5’ΔHygR::Intron5'::Read1::CCACGCTCTAAGCTC::Read2::Intron3'::3’ΔHygR + 5’ΔHygR::Intron5'::Read1::ATGCTATCTCAACCT::Read2::Intron3'::3’ΔHygR + 5’ΔHygR::Intron5'::Read1::AATCAATCTTAGCCA::Read2::Intron3'::3’ΔHygR + 5’ΔHygR::Intron5'::Read1::CAGCTATCTCAGTAC::Read2::Intron3'::3’ΔHygR + 5’ΔHygR::Intron5'::Read1::CGACCATTTAACATA::Read2::Intron3'::3’ΔHygR + 5’ΔHygR::Intron5'::Read1::AGTCCGTGTTAGCGA::Read2::Intron3'::3’ΔHygR + 5’ΔHygR::Intron5'::Read1::GGCCTGTGTTAGCCT::Read2::Intron3'::3’ΔHygR + 5’ΔHygR::Intron5'::Read1::AGCCTGTCTCAACAG::Read2::Intron3'::3’ΔHygR + 5’ΔHygR::Intron5'::Read1::GCACCATGTCAAAAA::Read2::Intron3'::3’ΔHygR + 5’ΔHygR::Intron5'::Read1::GCCCATTTTTATCGT::Read2::Intron3'::3’ΔHygR + 5’ΔHygR::Intron5'::Read1::TCTCCTTATAAAGGC::Read2::Intron3'::3’ΔHygR + 5’ΔHygR::Intron5'::Read1::TCTCAATATCATTTC::Read2::Intron3'::3’ΔHygR + 5’ΔHygR::Intron5'::Read1::TTGCATTCTTAGTAT::Read2::Intron3'::3’ΔHygR + 5’ΔHygR::Intron5'::Read1::CATCGTTGTAAAGCA::Read2::Intron3'::3’ΔHygR + 5’ΔHygR::Intron5'::Read1::CGTCTATCTCAACTC::Read2::Intron3'::3’ΔHygR + 5’ΔHygR::Intron5'::Read1::CGGCCATTTCAATTC::Read2::Intron3'::3’ΔHygR + 5’ΔHygR::Intron5'::Read1::GTACACTTTTATGAT::Read2::Intron3'::3’ΔHygR + 5’ΔHygR::Intron5'::Read1::CAGCGCTGTCACAGT::Read2::Intron3'::3’ΔHygR + 5’ΔHygR::Intron5'::Read1::ACTCTTTCTCACGCA::Read2::Intron3'::3’ΔHygR + 5’ΔHygR::Intron5'::Read1::TCGCACTTTAACACT::Read2::Intron3'::3’ΔHygR + 5’ΔHygR::Intron5'::Read1::ATTCGTTCTGATTTG::Read2::Intron3'::3’ΔHygR + 5’ΔHygR::Intron5'::Read1::TAGCTATGTAATTCA::Read2::Intron3'::3’ΔHygR + 5’ΔHygR::Intron5'::Read1::ACACCATTTGATCAT::Read2::Intron3'::3’ΔHygR + 5’ΔHygR::Intron5'::Read1::CTACAGTTTTAACGA::Read2::Intron3'::3’ΔHygR + 5’ΔHygR::Intron5'::Read1::AGTCTATCTCAGCCT::Read2::Intron3'::3’ΔHygR + 5’ΔHygR::Intron5'::Read1::TGTCATTCTTAATTC::Read2::Intron3'::3’ΔHygR + 5’ΔHygR::Intron5'::Read1::TATCAGTCTAACTTC::Read2::Intron3'::3’ΔHygR + 5’ΔHygR::Intron5'::Read1::CATCAATCTTAACGG::Read2::Intron3'::3’ΔHygR + 5’ΔHygR::Intron5'::Read1::GTACACTATAACGAA::Read2::Intron3'::3’ΔHygR + 5’ΔHygR::Intron5'::Read1::AACCTCTCTCAGATT::Read2::Intron3'::3’ΔHygR + 5’ΔHygR::Intron5'::Read1::ATGCGCTTTAACACG::Read2::Intron3'::3’ΔHygR + 5’ΔHygR::Intron5'::Read1::CGGCGATTTGATAAC::Read2::Intron3'::3’ΔHygR + 5’ΔHygR::Intron5'::Read1::GCTCATTGTCATACT::Read2::Intron3'::3’ΔHygR + 5’ΔHygR::Intron5'::Read1::GCTCTCTCTCAGAAC::Read2::Intron3'::3’ΔHygR + 5’ΔHygR::Intron5'::Read1::TTCCCTTATGAATTG::Read2::Intron3'::3’ΔHygR + 5’ΔHygR::Intron5'::Read1::CAACTCTCTCATACA::Read2::Intron3'::3’ΔHygR + 5’ΔHygR::Intron5'::Read1::ACCCATTCTCATATT::Read2::Intron3'::3’ΔHygR + 5’ΔHygR::Intron5'::Read1::TACCGATTTTATGTA::Read2::Intron3'::3’ΔHygR + 5’ΔHygR::Intron5'::Read1::CCCCCGTATAAACGT::Read2::Intron3'::3’ΔHygR + 5’ΔHygR::Intron5'::Read1::AACCTGTCTAACACT::Read2::Intron3'::3’ΔHygR + 5’ΔHygR::Intron5'::Read1::AGGCTCTGTGATTCC::Read2::Intron3'::3’ΔHygR + 5’ΔHygR::Intron5'::Read1::TCCCCCTATCAAATA::Read2::Intron3'::3’ΔHygR + 5’ΔHygR::Intron5'::Read1::CCCCATTATTATGTA::Read2::Intron3'::3’ΔHygR + 5’ΔHygR::Intron5'::Read1::TATCGCTATCATTAC::Read2::Intron3'::3’ΔHygR + 5’ΔHygR::Intron5'::Read1::TCTCGTTATGATGCC::Read2::Intron3'::3’ΔHygR + 5’ΔHygR::Intron5'::Read1::CTACACTCTCATGTT::Read2::Intron3'::3’ΔHygR + 5’ΔHygR::Intron5'::Read1::GGCCTATCTGACATG::Read2::Intron3'::3’ΔHygR + 5’ΔHygR::Intron5'::Read1::CGTCCATCTAACCAA::Read2::Intron3'::3’ΔHygR + 5’ΔHygR::Intron5'::Read1::GCCCTCTTTAACCAG::Read2::Intron3'::3’ΔHygR + 5’ΔHygR::Intron5'::Read1::TATCTGTCTGAGCAT::Read2::Intron3'::3’ΔHygR + 5’ΔHygR::Intron5'::Read1::AAGCTGTCTTACTGT::Read2::Intron3'::3’ΔHygR + 5’ΔHygR::Intron5'::Read1::GGCCTCTCTCAACCA::Read2::Intron3'::3’ΔHygR + 5’ΔHygR::Intron5'::Read1::AGGCCTTTTGATCCC::Read2::Intron3'::3’ΔHygR + 5’ΔHygR::Intron5'::Read1::GAACATTTTTATCAC::Read2::Intron3'::3’ΔHygR + 5’ΔHygR::Intron5'::Read1::TCGCATTTTTAGAAT::Read2::Intron3'::3’ΔHygR + 5’ΔHygR::Intron5'::Read1::CAACTTTCTCAGACT::Read2::Intron3'::3’ΔHygR + 5’ΔHygR::Intron5'::Read1::CTACAGTTTGACAAC::Read2::Intron3'::3’ΔHygR +

5’ΔHygR::Intron5'::Read1::TAACAGTCTTATTGG::Read2::Intron3'::3’ΔHygR + 5’ΔHygR::Intron5'::Read1::TAGCAGTGTCATCTC::Read2::Intron3'::3’ΔHygR + 5’ΔHygR::Intron5'::Read1::GGTCCATTTTAAAAC::Read2::Intron3'::3’ΔHygR + 5’ΔHygR::Intron5'::Read1::TTTCGCTCTTAACAT::Read2::Intron3'::3’ΔHygR + 5’ΔHygR::Intron5'::Read1::TAACCATATAATACC::Read2::Intron3'::3’ΔHygR + 5’ΔHygR::Intron5'::Read1::AAACGATGTCAATTA::Read2::Intron3'::3’ΔHygR + 5’ΔHygR::Intron5'::Read1::TTGCCTTATTATCTC::Read2::Intron3'::3’ΔHygR + 5’ΔHygR::Intron5'::Read1::AGTCCATATGATGAT::Read2::Intron3'::3’ΔHygR + 5’ΔHygR::Intron5'::Read1::TGACTTTCTGACATC::Read2::Intron3'::3’ΔHygR + 5’ΔHygR::Intron5'::Read1::TGCCTTTTTAATGTA::Read2::Intron3'::3’ΔHygR + 5’ΔHygR::Intron5'::Read1::TTGCAATGTCACTTT::Read2::Intron3'::3’ΔHygR + 5’ΔHygR::Intron5'::Read1::ATACGTTTTGATAAT::Read2::Intron3'::3’ΔHygR + 5’ΔHygR::Intron5'::Read1::GTCCGATGTTATTAA::Read2::Intron3'::3’ΔHygR + 5’ΔHygR::Intron5'::Read1::GTCCGTTTTGAGGCT::Read2::Intron3'::3’ΔHygR + 5’ΔHygR::Intron5'::Read1::ACACCATTTTACTTC::Read2::Intron3'::3’ΔHygR + 5’ΔHygR::Intron5'::Read1::TCCCCATGTCATACT::Read2::Intron3'::3’ΔHygR + 5’ΔHygR::Intron5'::Read1::AGTCGGTCTCATCTA::Read2::Intron3'::3’ΔHygR + 5’ΔHygR::Intron5'::Read1::TTCCCTTGTGAACCG::Read2::Intron3'::3’ΔHygR + 5’ΔHygR::Intron5'::Read1::CCACTCTTTGATTAA::Read2::Intron3'::3’ΔHygR + 5’ΔHygR::Intron5'::Read1::ACTCTCTCTTACCTC::Read2::Intron3'::3’ΔHygR + 5’ΔHygR::Intron5'::Read1::TGCCAGTATCAATGA::Read2::Intron3'::3’ΔHygR + 5’ΔHygR::Intron5'::Read1::GAACATTATCATAAA::Read2::Intron3'::3’ΔHygR + 5’ΔHygR::Intron5'::Read1::TGGCCTTTTGACTGG::Read2::Intron3'::3’ΔHygR + 5’ΔHygR::Intron5'::Read1::AGACTATATCACGCC::Read2::Intron3'::3’ΔHygR + 5’ΔHygR::Intron5'::Read1::CGCCCATTTGAAAAA::Read2::Intron3'::3’ΔHygR + 5’ΔHygR::Intron5'::Read1::TGGCACTGTTACTAC::Read2::Intron3'::3’ΔHygR + 5’ΔHygR::Intron5'::Read1::AACCGTTTTAATCCT::Read2::Intron3'::3’ΔHygR + 5’ΔHygR::Intron5'::Read1::ATCCTGTTTCATATT::Read2::Intron3'::3’ΔHygR + 5’ΔHygR::Intron5'::Read1::TTGCATTTTAAATTC::Read2::Intron3'::3’ΔHygR + 5’ΔHygR::Intron5'::Read1::AGTCTCTTTCAGGCC::Read2::Intron3'::3’ΔHygR + 5’ΔHygR::Intron5'::Read1::AAACTCTGTAAAACA::Read2::Intron3'::3’ΔHygR + 5’ΔHygR::Intron5'::Read1::CACCATTCTCAACCC::Read2::Intron3'::3’ΔHygR + 5’ΔHygR::Intron5'::Read1::TAGCTATTTCACCAA::Read2::Intron3'::3’ΔHygR + 5’ΔHygR::Intron5'::Read1::AGTCTGTTTCACCGG::Read2::Intron3'::3’ΔHygR + 5’ΔHygR::Intron5'::Read1::ATTCTGTCTAAATAA::Read2::Intron3'::3’ΔHygR + 5’ΔHygR::Intron5'::Read1::CGTCTGTCTCACAAA::Read2::Intron3'::3’ΔHygR + 5’ΔHygR::Intron5'::Read1::GTACATTGTAATATT::Read2::Intron3'::3’ΔHygR + 5’ΔHygR::Intron5'::Read1::CACCAGTTTAAGCCC::Read2::Intron3'::3’ΔHygR + 5’ΔHygR::Intron5'::Read1::GGACTATTTGACCTC::Read2::Intron3'::3’ΔHygR + 5’ΔHygR::Intron5'::Read1::GTTCCATTTCAAATT::Read2::Intron3'::3’ΔHygR + 5’ΔHygR::Intron5'::Read1::AATCAGTGTTAGGTT::Read2::Intron3'::3’ΔHygR + 5’ΔHygR::Intron5'::Read1::ATGCCATATTACCCC::Read2::Intron3'::3’ΔHygR + 5’ΔHygR::Intron5'::Read1::TCCCCATCTAACCTT::Read2::Intron3'::3’ΔHygR + 5’ΔHygR::Intron5'::Read1::ATCCTATGTAACAAG::Read2::Intron3'::3’ΔHygR + 5’ΔHygR::Intron5'::Read1::GGTCCATCTAAGCGA::Read2::Intron3'::3’ΔHygR + 5’ΔHygR::Intron5'::Read1::CGCCGGTTTAACCTC::Read2::Intron3'::3’ΔHygR + 5’ΔHygR::Intron5'::Read1::TGGCCCTTTAAAAAC::Read2::Intron3'::3’ΔHygR + 5’ΔHygR::Intron5'::Read1::TGACTATGTCAGACC::Read2::Intron3'::3’ΔHygR + 5’ΔHygR::Intron5'::Read1::CCCCGTTGTCATCAA::Read2::Intron3'::3’ΔHygR + 5’ΔHygR::Intron5'::Read1::TTTCTGTATCATCTT::Read2::Intron3'::3’ΔHygR + 5’ΔHygR::Intron5'::Read1::AATCGCTCTCACTGT::Read2::Intron3'::3’ΔHygR + 5’ΔHygR::Intron5'::Read1::GACCTTTTTCACCTA::Read2::Intron3'::3’ΔHygR + 5’ΔHygR::Intron5'::Read1::TGTCTGTTTCACGGC::Read2::Intron3'::3’ΔHygR + 5’ΔHygR::Intron5'::Read1::CTTCTCTTTCAGTGT::Read2::Intron3'::3’ΔHygR + 5’ΔHygR::Intron5'::Read1::TACCTCTGTAACTAT::Read2::Intron3'::3’ΔHygR + 5’ΔHygR::Intron5'::Read1::TCACACTTTGATAAG::Read2::Intron3'::3’ΔHygR + 5’ΔHygR::Intron5'::Read1::ACGCACTTTCATGTG::Read2::Intron3'::3’ΔHygR + 5’ΔHygR::Intron5'::Read1::TGTCGTTATGAACTA::Read2::Intron3'::3’ΔHygR + 5’ΔHygR::Intron5'::Read1::CGTCCTTATGATACT::Read2::Intron3'::3’ΔHygR + 5’ΔHygR::Intron5'::Read1::TCGCAGTGTCAAAAA::Read2::Intron3'::3’ΔHygR + 5’ΔHygR::Intron5'::Read1::TGTCGTTGTTATCGC::Read2::Intron3'::3’ΔHygR + 5’ΔHygR::Intron5'::Read1::GTGCTTTTTAATTTT::Read2::Intron3'::3’ΔHygR + 5’ΔHygR::Intron5'::Read1::TAACTGTTTTACAAC::Read2::Intron3'::3’ΔHygR + 5’ΔHygR::Intron5'::Read1::GTTCCTTTTTAAGAG::Read2::Intron3'::3’ΔHygR + 5’ΔHygR::Intron5'::Read1::ACACTATCTTACTAT::Read2::Intron3'::3’ΔHygR + 5’ΔHygR::Intron5'::Read1::AGTCCGTCTTACCGC::Read2::Intron3'::3’ΔHygR + 5’ΔHygR::Intron5'::Read1::TCTCACTGTTACTAA::Read2::Intron3'::3’ΔHygR + 5’ΔHygR::Intron5'::Read1::TATCCCTGTTACCAC::Read2::Intron3'::3’ΔHygR + 5’ΔHygR::Intron5'::Read1::CAGCCCTGTTATGCC::Read2::Intron3'::3’ΔHygR + 5’ΔHygR::Intron5'::Read1::TGCCCTTCTGACATG::Read2::Intron3'::3’ΔHygR +

5’ΔHygR::Intron5'::Read1::GTCCAATTTCACCAA::Read2::Intron3'::3’ΔHygR + 5’ΔHygR::Intron5'::Read1::TTTCTTTTTGATATC::Read2::Intron3'::3’ΔHygR + 5’ΔHygR::Intron5'::Read1::CGTCACTCTTAACGA::Read2::Intron3'::3’ΔHygR + 5’ΔHygR::Intron5'::Read1::GCTCAGTATCAATAG::Read2::Intron3'::3’ΔHygR + 5’ΔHygR::Intron5'::Read1::TAACTCTGTAATGAA::Read2::Intron3'::3’ΔHygR + 5’ΔHygR::Intron5'::Read1::CATCATTTTTACCAA::Read2::Intron3'::3’ΔHygR + 5’ΔHygR::Intron5'::Read1::CGGCATTCTAATAGA::Read2::Intron3'::3’ΔHygR + 5’ΔHygR::Intron5'::Read1::CAGCTATGTAAGGTC::Read2::Intron3'::3’ΔHygR + 5’ΔHygR::Intron5'::Read1::ACGCTTTTTGAAGTT::Read2::Intron3'::3’ΔHygR + 5’ΔHygR::Intron5'::Read1::GAACAGTGTCATCCG::Read2::Intron3'::3’ΔHygR + 5’ΔHygR::Intron5'::Read1::TGGCTTTCTCACACT::Read2::Intron3'::3’ΔHygR + 5’ΔHygR::Intron5'::Read1::TATCCGTGTAAATAC::Read2::Intron3'::3’ΔHygR + 5’ΔHygR::Intron5'::Read1::TAACTGTATTATGCC::Read2::Intron3'::3’ΔHygR + 5’ΔHygR::Intron5'::Read1::TGACAATCTCAAGTT::Read2::Intron3'::3’ΔHygR + 5’ΔHygR::Intron5'::Read1::TGTCCTTTTTACGGT::Read2::Intron3'::3’ΔHygR + 5’ΔHygR::Intron5'::Read1::GATCTGTATGAACAA::Read2::Intron3'::3’ΔHygR + 5’ΔHygR::Intron5'::Read1::TCGCATTATAATTCT::Read2::Intron3'::3’ΔHygR + 5’ΔHygR::Intron5'::Read1::CCCCCTTTTAAATCA::Read2::Intron3'::3’ΔHygR + 5’ΔHygR::Intron5'::Read1::TCCCCATGTCACTGC::Read2::Intron3'::3’ΔHygR + 5’ΔHygR::Intron5'::Read1::TGTCCTTTTTAGTTT::Read2::Intron3'::3’ΔHygR + 5’ΔHygR::Intron5'::Read1::TCCCGATTTCAATTT::Read2::Intron3'::3’ΔHygR + 5’ΔHygR::Intron5'::Read1::CCTCGATCTAATTTG::Read2::Intron3'::3’ΔHygR + 5’ΔHygR::Intron5'::Read1::AAGCACTTTCACTCA::Read2::Intron3'::3’ΔHygR + 5’ΔHygR::Intron5'::Read1::CAACATTATCACAAA::Read2::Intron3'::3’ΔHygR + 5’ΔHygR::Intron5'::Read1::CTGCGATTTCAAACT::Read2::Intron3'::3’ΔHygR + 5’ΔHygR::Intron5'::Read1::CTGCGATATCACCAT::Read2::Intron3'::3’ΔHygR + 5’ΔHygR::Intron5'::Read1::TATCTGTATAACACG::Read2::Intron3'::3’ΔHygR + 5’ΔHygR::Intron5'::Read1::ATCCGGTTTTACACA::Read2::Intron3'::3’ΔHygR + 5’ΔHygR::Intron5'::Read1::TAACAATCTTAGTAC::Read2::Intron3'::3’ΔHygR + 5’ΔHygR::Intron5'::Read1::ATGCCATGTCACTTA::Read2::Intron3'::3’ΔHygR + 5’ΔHygR::Intron5'::Read1::TAACACTTTTACCAC::Read2::Intron3'::3’ΔHygR + 5’ΔHygR::Intron5'::Read1::TTACAGTATTACCCC::Read2::Intron3'::3’ΔHygR + 5’ΔHygR::Intron5'::Read1::CCTCTTTTTCAGCCC::Read2::Intron3'::3’ΔHygR + 5’ΔHygR::Intron5'::Read1::CTACACTTTGAGAAA::Read2::Intron3'::3’ΔHygR + 5’ΔHygR::Intron5'::Read1::TCTCTTTGTTATCAC::Read2::Intron3'::3’ΔHygR + 5’ΔHygR::Intron5'::Read1::TCCCCATATTACACG::Read2::Intron3'::3’ΔHygR + 5’ΔHygR::Intron5'::Read1::TGGCACTATTACAAG::Read2::Intron3'::3’ΔHygR + 5’ΔHygR::Intron5'::Read1::CTTCGATATTACGTT::Read2::Intron3'::3’ΔHygR + 5’ΔHygR::Intron5'::Read1::TCACGCTTTGATAGT::Read2::Intron3'::3’ΔHygR + 5’ΔHygR::Intron5'::Read1::GTTCCTTCTAAGAAT::Read2::Intron3'::3’ΔHygR + 5’ΔHygR::Intron5'::Read1::GGGCCGTATTATTCG::Read2::Intron3'::3’ΔHygR + 5’ΔHygR::Intron5'::Read1::CTTCGTTATAAGAAC::Read2::Intron3'::3’ΔHygR + 5’ΔHygR::Intron5'::Read1::ACACGGTCTTACGGA::Read2::Intron3'::3’ΔHygR + 5’ΔHygR::Intron5'::Read1::CAGCGATGTAACAGC::Read2::Intron3'::3’ΔHygR + 5’ΔHygR::Intron5'::Read1::ACTCACTGTCAACCT::Read2::Intron3'::3’ΔHygR + 5’ΔHygR::Intron5'::Read1::TCTCCTTGTAACTGA::Read2::Intron3'::3’ΔHygR + 5’ΔHygR::Intron5'::Read1::TGTCGGTTTTATGTC::Read2::Intron3'::3’ΔHygR + 5’ΔHygR::Intron5'::Read1::TAACCGTCTTACTTC::Read2::Intron3'::3’ΔHygR + 5’ΔHygR::Intron5'::Read1::CGCCTGTCTGAGCAA::Read2::Intron3'::3’ΔHygR + 5’ΔHygR::Intron5'::Read1::TACCGATCTTAAAGT::Read2::Intron3'::3’ΔHygR + 5’ΔHygR::Intron5'::Read1::CGCCCGTCTCACGCC::Read2::Intron3'::3’ΔHygR + 5’ΔHygR::Intron5'::Read1::CACCCTTTTCACTCA::Read2::Intron3'::3’ΔHygR + 5’ΔHygR::Intron5'::Read1::AATCTGTTTTATTCT::Read2::Intron3'::3’ΔHygR + 5’ΔHygR::Intron5'::Read1::TCCCGTTATGAATAT::Read2::Intron3'::3’ΔHygR + 5’ΔHygR::Intron5'::Read1::CCTCTTTCTCATTGA::Read2::Intron3'::3’ΔHygR + 5’ΔHygR::Intron5'::Read1::GCTCAATCTGACTCA::Read2::Intron3'::3’ΔHygR + 5’ΔHygR::Intron5'::Read1::GGTCGGTCTCAAGAT::Read2::Intron3'::3’ΔHygR + 5’ΔHygR::Intron5'::Read1::AGTCGATGTCATGCC::Read2::Intron3'::3’ΔHygR + 5’ΔHygR::Intron5'::Read1::TTACCGTCTGACTCT::Read2::Intron3'::3’ΔHygR + 5’ΔHygR::Intron5'::Read1::CTCCTCTCTTATCCT::Read2::Intron3'::3’ΔHygR + 5’ΔHygR::Intron5'::Read1::TTCCCCTTTTACTCC::Read2::Intron3'::3’ΔHygR + 5’ΔHygR::Intron5'::Read1::AATCATTGTCAGGCC::Read2::Intron3'::3’ΔHygR + 5’ΔHygR::Intron5'::Read1::AAGCTGTTTTATTAT::Read2::Intron3'::3’ΔHygR + 5’ΔHygR::Intron5'::Read1::TTTCCCTTTTAAGTA::Read2::Intron3'::3’ΔHygR + 5’ΔHygR::Intron5'::Read1::CATCTCTGTAATGTC::Read2::Intron3'::3’ΔHygR + 5’ΔHygR::Intron5'::Read1::GTCCCATTTCACCCA::Read2::Intron3'::3’ΔHygR + 5’ΔHygR::Intron5'::Read1::TACCACTCTTATTCT::Read2::Intron3'::3’ΔHygR + 5’ΔHygR::Intron5'::Read1::CTTCACTTTTACACA::Read2::Intron3'::3’ΔHygR + 5’ΔHygR::Intron5'::Read1::CACCCGTGTCACAAC::Read2::Intron3'::3’ΔHygR + 5’ΔHygR::Intron5'::Read1::CTACAATCTTATTCT::Read2::Intron3'::3’ΔHygR +

5’ΔHygR::Intron5'::Read1::TCACCCTCTGATTTT::Read2::Intron3'::3’ΔHygR + 5’ΔHygR::Intron5'::Read1::GGTCCATTTTAAAGC::Read2::Intron3'::3’ΔHygR + 5’ΔHygR::Intron5'::Read1::TGTCATTATTATGGC::Read2::Intron3'::3’ΔHygR + 5’ΔHygR::Intron5'::Read1::CTCCCTTTTTAAGCA::Read2::Intron3'::3’ΔHygR + 5’ΔHygR::Intron5'::Read1::ACTCCATATAACGTC::Read2::Intron3'::3’ΔHygR + 5’ΔHygR::Intron5'::Read1::CAGCTTTTTTAACGA::Read2::Intron3'::3’ΔHygR + 5’ΔHygR::Intron5'::Read1::TAACGCTTTTAAGCG::Read2::Intron3'::3’ΔHygR + 5’ΔHygR::Intron5'::Read1::TGTCACTATGAATTC::Read2::Intron3'::3’ΔHygR + 5’ΔHygR::Intron5'::Read1::ACACGCTGTAACCAT::Read2::Intron3'::3’ΔHygR + 5’ΔHygR::Intron5'::Read1::CACCACTGTGATCAC::Read2::Intron3'::3’ΔHygR + 5’ΔHygR::Intron5'::Read1::GAACTCTATAAAAGG::Read2::Intron3'::3’ΔHygR + 5’ΔHygR::Intron5'::Read1::CTACGCTTTGACAAG::Read2::Intron3'::3’ΔHygR + 5’ΔHygR::Intron5'::Read1::TTACTGTTTTATAAT::Read2::Intron3'::3’ΔHygR + 5’ΔHygR::Intron5'::Read1::AGCCTCTCTAACTCG::Read2::Intron3'::3’ΔHygR + 5’ΔHygR::Intron5'::Read1::AAACTTTGTTAGCCA::Read2::Intron3'::3’ΔHygR + 5’ΔHygR::Intron5'::Read1::TTTCAATCTTATGTC::Read2::Intron3'::3’ΔHygR + 5’ΔHygR::Intron5'::Read1::CTGCATTATCAAAAG::Read2::Intron3'::3’ΔHygR + 5’ΔHygR::Intron5'::Read1::GCGCTTTATAAATTT::Read2::Intron3'::3’ΔHygR + 5’ΔHygR::Intron5'::Read1::GATCAATATTAAATA::Read2::Intron3'::3’ΔHygR + 5’ΔHygR::Intron5'::Read1::TCACTTTCTTACCGT::Read2::Intron3'::3’ΔHygR + 5’ΔHygR::Intron5'::Read1::ACCCCCTCTCATGCG::Read2::Intron3'::3’ΔHygR + 5’ΔHygR::Intron5'::Read1::GGGCTCTTTCAATTT::Read2::Intron3'::3’ΔHygR + 5’ΔHygR::Intron5'::Read1::TCTCCGTTTGAAGCC::Read2::Intron3'::3’ΔHygR + 5’ΔHygR::Intron5'::Read1::TTACCCTATAATACA::Read2::Intron3'::3’ΔHygR + 5’ΔHygR::Intron5'::Read1::TTTCGCTCTCACACA::Read2::Intron3'::3’ΔHygR + 5’ΔHygR::Intron5'::Read1::TCACATTTTGAACGA::Read2::Intron3'::3’ΔHygR + 5’ΔHygR::Intron5'::Read1::CAGCACTATTACTCC::Read2::Intron3'::3’ΔHygR + 5’ΔHygR::Intron5'::Read1::TCGCTATTTAACATC::Read2::Intron3'::3’ΔHygR + 5’ΔHygR::Intron5'::Read1::TTCCCGTCTCATGTC::Read2::Intron3'::3’ΔHygR + 5’ΔHygR::Intron5'::Read1::ACCCGTTTTAACAAG::Read2::Intron3'::3’ΔHygR + 5’ΔHygR::Intron5'::Read1::ACCCGCTGTCACCCA::Read2::Intron3'::3’ΔHygR + 5’ΔHygR::Intron5'::Read1::GTTCAATATTAACAT::Read2::Intron3'::3’ΔHygR + 5’ΔHygR::Intron5'::Read1::CTACTATATGACGAC::Read2::Intron3'::3’ΔHygR + 5’ΔHygR::Intron5'::Read1::CATCAATGTAAGACC::Read2::Intron3'::3’ΔHygR + 5’ΔHygR::Intron5'::Read1::CTACCCTTTTAACCT::Read2::Intron3'::3’ΔHygR + 5’ΔHygR::Intron5'::Read1::AATCTCTCTAATCAC::Read2::Intron3'::3’ΔHygR + 5’ΔHygR::Intron5'::Read1::AGCCAATCTCATCAC::Read2::Intron3'::3’ΔHygR + 5’ΔHygR::Intron5'::Read1::TCGCCGTTTTATCGG::Read2::Intron3'::3’ΔHygR + 5’ΔHygR::Intron5'::Read1::ACGCCGTTTCATATT::Read2::Intron3'::3’ΔHygR + 5’ΔHygR::Intron5'::Read1::TCACGGTATGACATT::Read2::Intron3'::3’ΔHygR + 5’ΔHygR::Intron5'::Read1::CCTCAGTTTAACCTA::Read2::Intron3'::3’ΔHygR + 5’ΔHygR::Intron5'::Read1::CGCCTGTTTCAAGTT::Read2::Intron3'::3’ΔHygR + 5’ΔHygR::Intron5'::Read1::TAGCATTCTCAGTAC::Read2::Intron3'::3’ΔHygR + 5’ΔHygR::Intron5'::Read1::TATCGGTCTCAATTT::Read2::Intron3'::3’ΔHygR + 5’ΔHygR::Intron5'::Read1::CAACCATCTCAAGGT::Read2::Intron3'::3’ΔHygR + 5’ΔHygR::Intron5'::Read1::TATCTATTTAAAACA::Read2::Intron3'::3’ΔHygR + 5’ΔHygR::Intron5'::Read1::CCGCGATGTGACCGT::Read2::Intron3'::3’ΔHygR + 5’ΔHygR::Intron5'::Read1::CCTCTGTCTGAGTAG::Read2::Intron3'::3’ΔHygR + 5’ΔHygR::Intron5'::Read1::CGCCTCTCTAACAGC::Read2::Intron3'::3’ΔHygR + 5’ΔHygR::Intron5'::Read1::GTCCCCTATTACCAT::Read2::Intron3'::3’ΔHygR + 5’ΔHygR::Intron5'::Read1::CCCCTCTATCAATGT::Read2::Intron3'::3’ΔHygR + 5’ΔHygR::Intron5'::Read1::CGCCCCTTTTATCCA::Read2::Intron3'::3’ΔHygR + 5’ΔHygR::Intron5'::Read1::GAACGCTTTTAGTAA::Read2::Intron3'::3’ΔHygR + 5’ΔHygR::Intron5'::Read1::ATTCCGTATCATTCC::Read2::Intron3'::3’ΔHygR + 5’ΔHygR::Intron5'::Read1::CCGCAATCTTAACTG::Read2::Intron3'::3’ΔHygR + 5’ΔHygR::Intron5'::Read1::GCTCGCTATGACCGA::Read2::Intron3'::3’ΔHygR + 5’ΔHygR::Intron5'::Read1::AACCGCTGTAACGCC::Read2::Intron3'::3’ΔHygR + 5’ΔHygR::Intron5'::Read1::CTACGCTGTAACAAT::Read2::Intron3'::3’ΔHygR + 5’ΔHygR::Intron5'::Read1::CTACCATTTCAAACC::Read2::Intron3'::3’ΔHygR + 5’ΔHygR::Intron5'::Read1::GTTCTATTTCAACAT::Read2::Intron3'::3’ΔHygR + 5’ΔHygR::Intron5'::Read1::CCGCATTTTCATCAG::Read2::Intron3'::3’ΔHygR + 5’ΔHygR::Intron5'::Read1::GCACCGTTTCATCTC::Read2::Intron3'::3’ΔHygR + 5’ΔHygR::Intron5'::Read1::TAGCTCTCTCATCCT::Read2::Intron3'::3’ΔHygR + 5’ΔHygR::Intron5'::Read1::TCACTCTATGAGTCC::Read2::Intron3'::3’ΔHygR + 5’ΔHygR::Intron5'::Read1::GAACTATGTAAATCC::Read2::Intron3'::3’ΔHygR + 5’ΔHygR::Intron5'::Read1::TTTCCATCTAATCCA::Read2::Intron3'::3’ΔHygR + 5’ΔHygR::Intron5'::Read1::TCACCTTTTGAATAC::Read2::Intron3'::3’ΔHygR + 5’ΔHygR::Intron5'::Read1::CCCCCCTGTAACCAG::Read2::Intron3'::3’ΔHygR + 5’ΔHygR::Intron5'::Read1::TACCGTTTTAACGTT::Read2::Intron3'::3’ΔHygR + 5’ΔHygR::Intron5'::Read1::CATCATTGTAACGCA::Read2::Intron3'::3’ΔHygR +

5’ΔHygR::Intron5'::Read1::CGCCGATCTCAGCCA::Read2::Intron3'::3’ΔHygR + 5’ΔHygR::Intron5'::Read1::AATCTTTTTTAATGA::Read2::Intron3'::3’ΔHygR + 5’ΔHygR::Intron5'::Read1::ATCCCCTGTAATCGC::Read2::Intron3'::3’ΔHygR + 5’ΔHygR::Intron5'::Read1::CGGCTCTCTAATACT::Read2::Intron3'::3’ΔHygR + 5’ΔHygR::Intron5'::Read1::TCGCGCTTTCACTGT::Read2::Intron3'::3’ΔHygR + 5’ΔHygR::Intron5'::Read1::CTCCATTTTGACTCA::Read2::Intron3'::3’ΔHygR + 5’ΔHygR::Intron5'::Read1::TTTCTCTATCACTTA::Read2::Intron3'::3’ΔHygR + 5’ΔHygR::Intron5'::Read1::GGTCCTTCTTATCCT::Read2::Intron3'::3’ΔHygR + 5’ΔHygR::Intron5'::Read1::TTTCGATCTGATCAA::Read2::Intron3'::3’ΔHygR + 5’ΔHygR::Intron5'::Read1::TAACATTGTAAATAC::Read2::Intron3'::3’ΔHygR + 5’ΔHygR::Intron5'::Read1::CACCCTTTTTAATCA::Read2::Intron3'::3’ΔHygR + 5’ΔHygR::Intron5'::Read1::CGACTCTCTCAGACA::Read2::Intron3'::3’ΔHygR + 5’ΔHygR::Intron5'::Read1::CTCCGCTTTAACCTA::Read2::Intron3'::3’ΔHygR + 5’ΔHygR::Intron5'::Read1::TCTCAATGTTATCCT::Read2::Intron3'::3’ΔHygR + 5’ΔHygR::Intron5'::Read1::CCACCTTGTTAGTTC::Read2::Intron3'::3’ΔHygR + 5’ΔHygR::Intron5'::Read1::CAGCTTTATAAGATC::Read2::Intron3'::3’ΔHygR + 5’ΔHygR::Intron5'::Read1::TCACCTTGTCAACCT::Read2::Intron3'::3’ΔHygR + 5’ΔHygR::Intron5'::Read1::TCACGCTGTGATGAC::Read2::Intron3'::3’ΔHygR + 5’ΔHygR::Intron5'::Read1::ACGCTCTCTTAACTA::Read2::Intron3'::3’ΔHygR + 5’ΔHygR::Intron5'::Read1::CATCCCTCTTAATTG::Read2::Intron3'::3’ΔHygR + 5’ΔHygR::Intron5'::Read1::TAACCCTATCACAAT::Read2::Intron3'::3’ΔHygR + 5’ΔHygR::Intron5'::Read1::TTACATTCTTATCCG::Read2::Intron3'::3’ΔHygR + 5’ΔHygR::Intron5'::Read1::ACGCTATTTAACGAA::Read2::Intron3'::3’ΔHygR + 5’ΔHygR::Intron5'::Read1::TTTCGCTTTTACCCT::Read2::Intron3'::3’ΔHygR + 5’ΔHygR::Intron5'::Read1::ACACATTCTCACAAC::Read2::Intron3'::3’ΔHygR + 5’ΔHygR::Intron5'::Read1::ACCCCTTGTAACGGC::Read2::Intron3'::3’ΔHygR + 5’ΔHygR::Intron5'::Read1::CTCCTATTTAATGTA::Read2::Intron3'::3’ΔHygR + 5’ΔHygR::Intron5'::Read1::TTACGGTTTAATATA::Read2::Intron3'::3’ΔHygR + 5’ΔHygR::Intron5'::Read1::AAGCGGTATCAGTGG::Read2::Intron3'::3’ΔHygR + 5’ΔHygR::Intron5'::Read1::TCACAGTATGAATCC::Read2::Intron3'::3’ΔHygR + 5’ΔHygR::Intron5'::Read1::GTGCTATATCACAAC::Read2::Intron3'::3’ΔHygR + 5’ΔHygR::Intron5'::Read1::TGTCGGTGTAATATA::Read2::Intron3'::3’ΔHygR + 5’ΔHygR::Intron5'::Read1::ACGCATTCTCATATT::Read2::Intron3'::3’ΔHygR + 5’ΔHygR::Intron5'::Read1::ATCCTTTCTAACTAA::Read2::Intron3'::3’ΔHygR + 5’ΔHygR::Intron5'::Read1::ACGCAATGTGAGCAC::Read2::Intron3'::3’ΔHygR + 5’ΔHygR::Intron5'::Read1::ATTCACTTTCAGTGT::Read2::Intron3'::3’ΔHygR + 5’ΔHygR::Intron5'::Read1::ATTCTGTGTTATCCC::Read2::Intron3'::3’ΔHygR + 5’ΔHygR::Intron5'::Read1::CCACTCTATTAGAGA::Read2::Intron3'::3’ΔHygR + 5’ΔHygR::Intron5'::Read1::CAACTTTTTTACTAC::Read2::Intron3'::3’ΔHygR + 5’ΔHygR::Intron5'::Read1::CCCCTTTTTCAGTAT::Read2::Intron3'::3’ΔHygR + 5’ΔHygR::Intron5'::Read1::TTCCTTTCTTATTCC::Read2::Intron3'::3’ΔHygR + 5’ΔHygR::Intron5'::Read1::ATACTCTTTAACCCC::Read2::Intron3'::3’ΔHygR + 5’ΔHygR::Intron5'::Read1::GAACCGTATGACCCC::Read2::Intron3'::3’ΔHygR + 5’ΔHygR::Intron5'::Read1::TTCCGTTGTCATGCT::Read2::Intron3'::3’ΔHygR + 5’ΔHygR::Intron5'::Read1::ACTCCATTTGACTGA::Read2::Intron3'::3’ΔHygR + 5’ΔHygR::Intron5'::Read1::ACTCACTATAACAGC::Read2::Intron3'::3’ΔHygR + 5’ΔHygR::Intron5'::Read1::TTCCTTTATAATCAA::Read2::Intron3'::3’ΔHygR + 5’ΔHygR::Intron5'::Read1::CTCCCATATTAGAGT::Read2::Intron3'::3’ΔHygR + 5’ΔHygR::Intron5'::Read1::TCTCGGTATAATCAC::Read2::Intron3'::3’ΔHygR + 5’ΔHygR::Intron5'::Read1::CCTCGTTATCACGCT::Read2::Intron3'::3’ΔHygR + 5’ΔHygR::Intron5'::Read1::CAACACTATGAGCAC::Read2::Intron3'::3’ΔHygR + 5’ΔHygR::Intron5'::Read1::TTTCTATTTCACCAC::Read2::Intron3'::3’ΔHygR + 5’ΔHygR::Intron5'::Read1::TTCCTATTTTACGTT::Read2::Intron3'::3’ΔHygR + 5’ΔHygR::Intron5'::Read1::CGACACTCTTAGTTC::Read2::Intron3'::3’ΔHygR + 5’ΔHygR::Intron5'::Read1::TACCTCTTTTATAAA::Read2::Intron3'::3’ΔHygR + 5’ΔHygR::Intron5'::Read1::GGACTATTTTAGATT::Read2::Intron3'::3’ΔHygR + 5’ΔHygR::Intron5'::Read1::CACCAATGTCAAAGC::Read2::Intron3'::3’ΔHygR + 5’ΔHygR::Intron5'::Read1::CTCCATTGTTAAGGC::Read2::Intron3'::3’ΔHygR + 5’ΔHygR::Intron5'::Read1::CTCCATTTTTATCAC::Read2::Intron3'::3’ΔHygR + 5’ΔHygR::Intron5'::Read1::GCTCGCTTTAATGCA::Read2::Intron3'::3’ΔHygR + 5’ΔHygR::Intron5'::Read1::TTTCAATTTGATTCT::Read2::Intron3'::3’ΔHygR + 5’ΔHygR::Intron5'::Read1::AGACCCTTTAATCTC::Read2::Intron3'::3’ΔHygR + 5’ΔHygR::Intron5'::Read1::GACCCGTCTTAAACT::Read2::Intron3'::3’ΔHygR + 5’ΔHygR::Intron5'::Read1::TCTCTATCTCACCTC::Read2::Intron3'::3’ΔHygR + 5’ΔHygR::Intron5'::Read1::GTTCATTGTTAATCC::Read2::Intron3'::3’ΔHygR + 5’ΔHygR::Intron5'::Read1::CCTCTATGTTATTAA::Read2::Intron3'::3’ΔHygR + 5’ΔHygR::Intron5'::Read1::GCTCAGTCTCACTGC::Read2::Intron3'::3’ΔHygR + 5’ΔHygR::Intron5'::Read1::CCCCCATCTTAAGTC::Read2::Intron3'::3’ΔHygR + 5’ΔHygR::Intron5'::Read1::CGTCCCTCTTAAGTC::Read2::Intron3'::3’ΔHygR + 5’ΔHygR::Intron5'::Read1::TCACGGTGTCAACCT::Read2::Intron3'::3’ΔHygR +

5’ΔHygR::Intron5'::Read1::CCGCATTTTAACTAC::Read2::Intron3'::3’ΔHygR + 5’ΔHygR::Intron5'::Read1::ACGCGCTTTCACATC::Read2::Intron3'::3’ΔHygR + 5’ΔHygR::Intron5'::Read1::TCACGGTCTCATCCT::Read2::Intron3'::3’ΔHygR + 5’ΔHygR::Intron5'::Read1::CAGCCATCTAAGTCT::Read2::Intron3'::3’ΔHygR + 5’ΔHygR::Intron5'::Read1::TATCATTCTGAATAC::Read2::Intron3'::3’ΔHygR + 5’ΔHygR::Intron5'::Read1::GCCCGGTCTAATGTG::Read2::Intron3'::3’ΔHygR + 5’ΔHygR::Intron5'::Read1::GCCCCGTATTAAAAG::Read2::Intron3'::3’ΔHygR + 5’ΔHygR::Intron5'::Read1::CCTCGATCTAACTCG::Read2::Intron3'::3’ΔHygR + 5’ΔHygR::Intron5'::Read1::ATACTGTCTAATTCC::Read2::Intron3'::3’ΔHygR + 5’ΔHygR::Intron5'::Read1::TAGCTGTATGAGTCC::Read2::Intron3'::3’ΔHygR + 5’ΔHygR::Intron5'::Read1::AACCTTTCTTACGAT::Read2::Intron3'::3’ΔHygR + 5’ΔHygR::Intron5'::Read1::TTTCTATGTAAGTCG::Read2::Intron3'::3’ΔHygR + 5’ΔHygR::Intron5'::Read1::CGTCTCTATCATGAA::Read2::Intron3'::3’ΔHygR + 5’ΔHygR::Intron5'::Read1::TACCTGTATTAACCC::Read2::Intron3'::3’ΔHygR + 5’ΔHygR::Intron5'::Read1::TGCCTCTGTCATAAT::Read2::Intron3'::3’ΔHygR + 5’ΔHygR::Intron5'::Read1::TCACCCTGTCACACA::Read2::Intron3'::3’ΔHygR + 5’ΔHygR::Intron5'::Read1::TCTCTCTTTTAAGCC::Read2::Intron3'::3’ΔHygR + 5’ΔHygR::Intron5'::Read1::CGCCTGTGTAAGCCT::Read2::Intron3'::3’ΔHygR + 5’ΔHygR::Intron5'::Read1::ACTCTTTATCACATT::Read2::Intron3'::3’ΔHygR + 5’ΔHygR::Intron5'::Read1::AAGCCCTATAATACC::Read2::Intron3'::3’ΔHygR + 5’ΔHygR::Intron5'::Read1::CATCGATTTTACGAA::Read2::Intron3'::3’ΔHygR + 5’ΔHygR::Intron5'::Read1::TCCCTTTATCAGCAT::Read2::Intron3'::3’ΔHygR + 5’ΔHygR::Intron5'::Read1::CGTCACTTTGACCAT::Read2::Intron3'::3’ΔHygR + 5’ΔHygR::Intron5'::Read1::GCACTTTCTCATTTA::Read2::Intron3'::3’ΔHygR + 5’ΔHygR::Intron5'::Read1::TATCTCTCTCACCTA::Read2::Intron3'::3’ΔHygR + 5’ΔHygR::Intron5'::Read1::TCACTCTTTTAATTC::Read2::Intron3'::3’ΔHygR + 5’ΔHygR::Intron5'::Read1::ACCCTTTCTCATATA::Read2::Intron3'::3’ΔHygR + 5’ΔHygR::Intron5'::Read1::GACCTTTGTAATGAC::Read2::Intron3'::3’ΔHygR + 5’ΔHygR::Intron5'::Read1::AACCATTCTTATTGT::Read2::Intron3'::3’ΔHygR + 5’ΔHygR::Intron5'::Read1::AACCCTTTTCACTTG::Read2::Intron3'::3’ΔHygR + 5’ΔHygR::Intron5'::Read1::AACCACTCTCACTGA::Read2::Intron3'::3’ΔHygR + 5’ΔHygR::Intron5'::Read1::TACCGTTTTAAGTTC::Read2::Intron3'::3’ΔHygR + 5’ΔHygR::Intron5'::Read1::TACCTCTCTCAATTG::Read2::Intron3'::3’ΔHygR + 5’ΔHygR::Intron5'::Read1::CCACTCTGTAAATAT::Read2::Intron3'::3’ΔHygR + 5’ΔHygR::Intron5'::Read1::GCCCCTTTTAACCAC::Read2::Intron3'::3’ΔHygR + 5’ΔHygR::Intron5'::Read1::CCGCCGTGTGACTTT::Read2::Intron3'::3’ΔHygR + 5’ΔHygR::Intron5'::Read1::CTCCATTTTGACCCA::Read2::Intron3'::3’ΔHygR + 5’ΔHygR::Intron5'::Read1::GTGCAATTTCATCAG::Read2::Intron3'::3’ΔHygR + 5’ΔHygR::Intron5'::Read1::TGACTTTCTAATTAA::Read2::Intron3'::3’ΔHygR + 5’ΔHygR::Intron5'::Read1::AACCACTGTGACGTT::Read2::Intron3'::3’ΔHygR + 5’ΔHygR::Intron5'::Read1::GGACAATATTACCTT::Read2::Intron3'::3’ΔHygR + 5’ΔHygR::Intron5'::Read1::CTCCACTGTCAACAA::Read2::Intron3'::3’ΔHygR + 5’ΔHygR::Intron5'::Read1::GCCCGTTCTAACAAA::Read2::Intron3'::3’ΔHygR + 5’ΔHygR::Intron5'::Read1::CGACATTCTAAGAGA::Read2::Intron3'::3’ΔHygR + 5’ΔHygR::Intron5'::Read1::AGACCTTCTAAGCAT::Read2::Intron3'::3’ΔHygR + 5’ΔHygR::Intron5'::Read1::GGGCGCTGTGATTTC::Read2::Intron3'::3’ΔHygR + 5’ΔHygR::Intron5'::Read1::GGACCGTATCACCCA::Read2::Intron3'::3’ΔHygR + 5’ΔHygR::Intron5'::Read1::GAGCCATTTAACTCT::Read2::Intron3'::3’ΔHygR + 5’ΔHygR::Intron5'::Read1::ACCCGATATAACTCC::Read2::Intron3'::3’ΔHygR + 5’ΔHygR::Intron5'::Read1::TTTCGATTTAACTTT::Read2::Intron3'::3’ΔHygR + 5’ΔHygR::Intron5'::Read1::AATCATTTTAATTCC::Read2::Intron3'::3’ΔHygR + 5’ΔHygR::Intron5'::Read1::CCACCATGTAACCAA::Read2::Intron3'::3’ΔHygR + 5’ΔHygR::Intron5'::Read1::TTGCGCTGTTATTCC::Read2::Intron3'::3’ΔHygR + 5’ΔHygR::Intron5'::Read1::CTCCTTTATTAGCCT::Read2::Intron3'::3’ΔHygR + 5’ΔHygR::Intron5'::Read1::TAGCCCTGTTATCTA::Read2::Intron3'::3’ΔHygR + 5’ΔHygR::Intron5'::Read1::TTGCTCTCTGACCCA::Read2::Intron3'::3’ΔHygR + 5’ΔHygR::Intron5'::Read1::GATCTGTATTATTAG::Read2::Intron3'::3’ΔHygR + 5’ΔHygR::Intron5'::Read1::GCTCGATGTGAGTCA::Read2::Intron3'::3’ΔHygR + 5’ΔHygR::Intron5'::Read1::TGTCCCTGTCATGGA::Read2::Intron3'::3’ΔHygR + 5’ΔHygR::Intron5'::Read1::AAACCCTCTGAGTAT::Read2::Intron3'::3’ΔHygR + 5’ΔHygR::Intron5'::Read1::GCACACTTTTAATGT::Read2::Intron3'::3’ΔHygR + 5’ΔHygR::Intron5'::Read1::TATCCCTCTCATCTC::Read2::Intron3'::3’ΔHygR + 5’ΔHygR::Intron5'::Read1::CGACCTTCTCAGCCC::Read2::Intron3'::3’ΔHygR + 5’ΔHygR::Intron5'::Read1::TCCCAGTTTCAAAGC::Read2::Intron3'::3’ΔHygR + 5’ΔHygR::Intron5'::Read1::TCTCAATCTTACCAA::Read2::Intron3'::3’ΔHygR + 5’ΔHygR::Intron5'::Read1::CCCCACTTTCATTGT::Read2::Intron3'::3’ΔHygR + 5’ΔHygR::Intron5'::Read1::TACCCTTATCACCGT::Read2::Intron3'::3’ΔHygR + 5’ΔHygR::Intron5'::Read1::CCCCTATATGATGCT::Read2::Intron3'::3’ΔHygR + 5’ΔHygR::Intron5'::Read1::GCTCGGTATGACTAT::Read2::Intron3'::3’ΔHygR + 5’ΔHygR::Intron5'::Read1::AGACCGTGTGATTTT::Read2::Intron3'::3’ΔHygR +

5’ΔHygR::Intron5'::Read1::TAACTTTCTTAAACT::Read2::Intron3'::3’ΔHygR + 5’ΔHygR::Intron5'::Read1::AGCCGTTGTTATTGT::Read2::Intron3'::3’ΔHygR + 5’ΔHygR::Intron5'::Read1::AATCGATATCACCTG::Read2::Intron3'::3’ΔHygR + 5’ΔHygR::Intron5'::Read1::TTTCATTTTAATGGA::Read2::Intron3'::3’ΔHygR + 5’ΔHygR::Intron5'::Read1::CAACATTGTGATCCG::Read2::Intron3'::3’ΔHygR + 5’ΔHygR::Intron5'::Read1::GGCCGCTGTTACCTA::Read2::Intron3'::3’ΔHygR + 5’ΔHygR::Intron5'::Read1::CACCCTTTTTACTAC::Read2::Intron3'::3’ΔHygR + 5’ΔHygR::Intron5'::Read1::CAGCACTCTCAAACA::Read2::Intron3'::3’ΔHygR + 5’ΔHygR::Intron5'::Read1::CTCCATTTTCAGCTT::Read2::Intron3'::3’ΔHygR + 5’ΔHygR::Intron5'::Read1::GTGCGATATAACAGC::Read2::Intron3'::3’ΔHygR + 5’ΔHygR::Intron5'::Read1::GAGCCTTTTCACTTC::Read2::Intron3'::3’ΔHygR + 5’ΔHygR::Intron5'::Read1::TACCCTTATCAATTA::Read2::Intron3'::3’ΔHygR + 5’ΔHygR::Intron5'::Read1::TGCCTCTCTCAATCG::Read2::Intron3'::3’ΔHygR + 5’ΔHygR::Intron5'::Read1::ACGCCCTTTCATACC::Read2::Intron3'::3’ΔHygR + 5’ΔHygR::Intron5'::Read1::ATACAATATCACCAG::Read2::Intron3'::3’ΔHygR + 5’ΔHygR::Intron5'::Read1::GGTCCTTCTTACCTT::Read2::Intron3'::3’ΔHygR + 5’ΔHygR::Intron5'::Read1::CGGCCCTCTCAAACC::Read2::Intron3'::3’ΔHygR + 5’ΔHygR::Intron5'::Read1::ACGCCCTTTGAAGTT::Read2::Intron3'::3’ΔHygR + 5’ΔHygR::Intron5'::Read1::ATTCATTATTAGATC::Read2::Intron3'::3’ΔHygR + 5’ΔHygR::Intron5'::Read1::CATCACTCTTATGCC::Read2::Intron3'::3’ΔHygR + 5’ΔHygR::Intron5'::Read1::TAGCGCTGTTATTCT::Read2::Intron3'::3’ΔHygR + 5’ΔHygR::Intron5'::Read1::AAGCTCTGTCATTTA::Read2::Intron3'::3’ΔHygR + 5’ΔHygR::Intron5'::Read1::TAACGATGTGACGGT::Read2::Intron3'::3’ΔHygR + 5’ΔHygR::Intron5'::Read1::TTGCCGTGTCACAAT::Read2::Intron3'::3’ΔHygR + 5’ΔHygR::Intron5'::Read1::GACCAGTTTTACCTT::Read2::Intron3'::3’ΔHygR + 5’ΔHygR::Intron5'::Read1::CCTCATTTTCAAACC::Read2::Intron3'::3’ΔHygR + 5’ΔHygR::Intron5'::Read1::GTACCTTCTTAAAAA::Read2::Intron3'::3’ΔHygR + 5’ΔHygR::Intron5'::Read1::AACCGCTCTCAATCG::Read2::Intron3'::3’ΔHygR + 5’ΔHygR::Intron5'::Read1::AACCGTTGTCAATCA::Read2::Intron3'::3’ΔHygR + 5’ΔHygR::Intron5'::Read1::CAGCGCTTTAATACC::Read2::Intron3'::3’ΔHygR + 5’ΔHygR::Intron5'::Read1::ACGCGATGTTATCTT::Read2::Intron3'::3’ΔHygR + 5’ΔHygR::Intron5'::Read1::CGCCCCTATTACTCG::Read2::Intron3'::3’ΔHygR + 5’ΔHygR::Intron5'::Read1::ATGCTGTCTTAATGT::Read2::Intron3'::3’ΔHygR + 5’ΔHygR::Intron5'::Read1::CCGCCCTGTTATCGT::Read2::Intron3'::3’ΔHygR + 5’ΔHygR::Intron5'::Read1::CAGCCTTTTCAGAGT::Read2::Intron3'::3’ΔHygR + 5’ΔHygR::Intron5'::Read1::GGGCGGTCTAATCCA::Read2::Intron3'::3’ΔHygR + 5’ΔHygR::Intron5'::Read1::TATCTATCTCACAGT::Read2::Intron3'::3’ΔHygR + 5’ΔHygR::Intron5'::Read1::GTTCATTCTTACATC::Read2::Intron3'::3’ΔHygR + 5’ΔHygR::Intron5'::Read1::AAACTGTCTTATTCC::Read2::Intron3'::3’ΔHygR + 5’ΔHygR::Intron5'::Read1::ACGCCTTATGAAATC::Read2::Intron3'::3’ΔHygR + 5’ΔHygR::Intron5'::Read1::GTACATTTTTACGCA::Read2::Intron3'::3’ΔHygR + 5’ΔHygR::Intron5'::Read1::CCTCACTCTAAGCAG::Read2::Intron3'::3’ΔHygR + 5’ΔHygR::Intron5'::Read1::TGTCTATCTTATCCC::Read2::Intron3'::3’ΔHygR + 5’ΔHygR::Intron5'::Read1::TTGCCCTCTTAAGCT::Read2::Intron3'::3’ΔHygR + 5’ΔHygR::Intron5'::Read1::CGGCTGTTTGATTTT::Read2::Intron3'::3’ΔHygR + 5’ΔHygR::Intron5'::Read1::TCACTTTCTTAACAG::Read2::Intron3'::3’ΔHygR + 5’ΔHygR::Intron5'::Read1::TCCCCGTCTTAACCA::Read2::Intron3'::3’ΔHygR + 5’ΔHygR::Intron5'::Read1::CCACACTTTTATTTC::Read2::Intron3'::3’ΔHygR + 5’ΔHygR::Intron5'::Read1::GCACCCTATCAACGA::Read2::Intron3'::3’ΔHygR + 5’ΔHygR::Intron5'::Read1::AAACGCTCTCAATTT::Read2::Intron3'::3’ΔHygR + 5’ΔHygR::Intron5'::Read1::AGTCGATATGACGAA::Read2::Intron3'::3’ΔHygR + 5’ΔHygR::Intron5'::Read1::AGACTGTCTTATGCA::Read2::Intron3'::3’ΔHygR + 5’ΔHygR::Intron5'::Read1::CTCCTGTATCAGCCG::Read2::Intron3'::3’ΔHygR + 5’ΔHygR::Intron5'::Read1::TAACAATTTTAGCAA::Read2::Intron3'::3’ΔHygR + 5’ΔHygR::Intron5'::Read1::ATCCTATGTCATCCG::Read2::Intron3'::3’ΔHygR + 5’ΔHygR::Intron5'::Read1::GAGCAGTTTTAGGTT::Read2::Intron3'::3’ΔHygR + 5’ΔHygR::Intron5'::Read1::TTTCCATCTAATATG::Read2::Intron3'::3’ΔHygR + 5’ΔHygR::Intron5'::Read1::CCTCACTCTAAAAGC::Read2::Intron3'::3’ΔHygR + 5’ΔHygR::Intron5'::Read1::AGGCCCTCTCAACAC::Read2::Intron3'::3’ΔHygR + 5’ΔHygR::Intron5'::Read1::CAACCATATTAATCA::Read2::Intron3'::3’ΔHygR + 5’ΔHygR::Intron5'::Read1::CTCCTGTATTACTAC::Read2::Intron3'::3’ΔHygR + 5’ΔHygR::Intron5'::Read1::TACCATTTTCAACGT::Read2::Intron3'::3’ΔHygR + 5’ΔHygR::Intron5'::Read1::CAACGCTATAAATAC::Read2::Intron3'::3’ΔHygR + 5’ΔHygR::Intron5'::Read1::TACCCGTTTAAACCC::Read2::Intron3'::3’ΔHygR + 5’ΔHygR::Intron5'::Read1::TATCTTTGTAACATA::Read2::Intron3'::3’ΔHygR + 5’ΔHygR::Intron5'::Read1::ATTCTATCTGAACAC::Read2::Intron3'::3’ΔHygR + 5’ΔHygR::Intron5'::Read1::GTCCTATTTGAGCTT::Read2::Intron3'::3’ΔHygR + 5’ΔHygR::Intron5'::Read1::TACCATTTTCATCTT::Read2::Intron3'::3’ΔHygR + 5’ΔHygR::Intron5'::Read1::ACCCTATCTCACAGT::Read2::Intron3'::3’ΔHygR + 5’ΔHygR::Intron5'::Read1::AGTCGTTCTCACTCC::Read2::Intron3'::3’ΔHygR +

5’ΔHygR::Intron5'::Read1::ACGCACTATCATGCT::Read2::Intron3'::3’ΔHygR + 5’ΔHygR::Intron5'::Read1::TGCCACTATGAACGT::Read2::Intron3'::3’ΔHygR + 5’ΔHygR::Intron5'::Read1::AGTCTCTTTCACGTA::Read2::Intron3'::3’ΔHygR + 5’ΔHygR::Intron5'::Read1::ACTCAATCTGACATA::Read2::Intron3'::3’ΔHygR + 5’ΔHygR::Intron5'::Read1::TCACAGTTTAAGACA::Read2::Intron3'::3’ΔHygR + 5’ΔHygR::Intron5'::Read1::CTCCTTTCTCACAAC::Read2::Intron3'::3’ΔHygR + 5’ΔHygR::Intron5'::Read1::GCACCATCTCACCAA::Read2::Intron3'::3’ΔHygR + 5’ΔHygR::Intron5'::Read1::TAACCGTATAATGCT::Read2::Intron3'::3’ΔHygR + 5’ΔHygR::Intron5'::Read1::TCCCTCTTTGACAAT::Read2::Intron3'::3’ΔHygR + 5’ΔHygR::Intron5'::Read1::TGGCAATGTCAAATC::Read2::Intron3'::3’ΔHygR + 5’ΔHygR::Intron5'::Read1::TCGCAATGTTACGAA::Read2::Intron3'::3’ΔHygR + 5’ΔHygR::Intron5'::Read1::GTGCCTTATAACCCA::Read2::Intron3'::3’ΔHygR + 5’ΔHygR::Intron5'::Read1::ATCCTTTTTTAAGAG::Read2::Intron3'::3’ΔHygR + 5’ΔHygR::Intron5'::Read1::CAGCTATATTAAGTC::Read2::Intron3'::3’ΔHygR + 5’ΔHygR::Intron5'::Read1::CGTCGGTGTTACTAC::Read2::Intron3'::3’ΔHygR + 5’ΔHygR::Intron5'::Read1::CGGCCTTCTGACCTA::Read2::Intron3'::3’ΔHygR + 5’ΔHygR::Intron5'::Read1::AACCTCTATAAATCA::Read2::Intron3'::3’ΔHygR + 5’ΔHygR::Intron5'::Read1::AAACCCTCTTATTCC::Read2::Intron3'::3’ΔHygR + 5’ΔHygR::Intron5'::Read1::TTACATTGTAACAGT::Read2::Intron3'::3’ΔHygR + 5’ΔHygR::Intron5'::Read1::GGGCGTTCTCACATG::Read2::Intron3'::3’ΔHygR + 5’ΔHygR::Intron5'::Read1::GGTCAATGTTAAACA::Read2::Intron3'::3’ΔHygR + 5’ΔHygR::Intron5'::Read1::ATTCGCTATCACTAG::Read2::Intron3'::3’ΔHygR + 5’ΔHygR::Intron5'::Read1::TGCCACTCTTACATC::Read2::Intron3'::3’ΔHygR + 5’ΔHygR::Intron5'::Read1::TTCCACTATCACCGC::Read2::Intron3'::3’ΔHygR + 5’ΔHygR::Intron5'::Read1::CATCCGTCTGAAGGC::Read2::Intron3'::3’ΔHygR + 5’ΔHygR::Intron5'::Read1::CCCCACTGTAACCGC::Read2::Intron3'::3’ΔHygR + 5’ΔHygR::Intron5'::Read1::CCTCTTTGTAATCTT::Read2::Intron3'::3’ΔHygR + 5’ΔHygR::Intron5'::Read1::TGACCGTTTTAAGTC::Read2::Intron3'::3’ΔHygR + 5’ΔHygR::Intron5'::Read1::CCACTTTGTCAAGCT::Read2::Intron3'::3’ΔHygR + 5’ΔHygR::Intron5'::Read1::TCGCTTTCTCAGTCA::Read2::Intron3'::3’ΔHygR + 5’ΔHygR::Intron5'::Read1::GTCCGGTATGACTAC::Read2::Intron3'::3’ΔHygR + 5’ΔHygR::Intron5'::Read1::CAGCACTGTTACATT::Read2::Intron3'::3’ΔHygR + 5’ΔHygR::Intron5'::Read1::TATCTGTATTACGTC::Read2::Intron3'::3’ΔHygR + 5’ΔHygR::Intron5'::Read1::TTACACTATAATGTG::Read2::Intron3'::3’ΔHygR + 5’ΔHygR::Intron5'::Read1::AGACCATGTCAGCCC::Read2::Intron3'::3’ΔHygR + 5’ΔHygR::Intron5'::Read1::CATCTTTATCACCGC::Read2::Intron3'::3’ΔHygR + 5’ΔHygR::Intron5'::Read1::CTACGTTGTGATTCC::Read2::Intron3'::3’ΔHygR + 5’ΔHygR::Intron5'::Read1::TCCCATTGTTACGAT::Read2::Intron3'::3’ΔHygR + 5’ΔHygR::Intron5'::Read1::CAACGTTATGAATCT::Read2::Intron3'::3’ΔHygR + 5’ΔHygR::Intron5'::Read1::TCTCTTTTTCAAGGT::Read2::Intron3'::3’ΔHygR + 5’ΔHygR::Intron5'::Read1::TTACTCTCTAAAAAA::Read2::Intron3'::3’ΔHygR + 5’ΔHygR::Intron5'::Read1::GTGCATTATGAGTAC::Read2::Intron3'::3’ΔHygR + 5’ΔHygR::Intron5'::Read1::TCCCCATGTCAGTAC::Read2::Intron3'::3’ΔHygR + 5’ΔHygR::Intron5'::Read1::CTGCTGTATTATCCC::Read2::Intron3'::3’ΔHygR + 5’ΔHygR::Intron5'::Read1::TCTCAATTTTAACCT::Read2::Intron3'::3’ΔHygR + 5’ΔHygR::Intron5'::Read1::TCTCAATTTTAACTT::Read2::Intron3'::3’ΔHygR + 5’ΔHygR::Intron5'::Read1::TTCCCATCTCATCTC::Read2::Intron3'::3’ΔHygR + 5’ΔHygR::Intron5'::Read1::TACCCCTTTGAAGTG::Read2::Intron3'::3’ΔHygR + 5’ΔHygR::Intron5'::Read1::TCCCTGTATGACTGC::Read2::Intron3'::3’ΔHygR + 5’ΔHygR::Intron5'::Read1::AATCCATTTGACTGT::Read2::Intron3'::3’ΔHygR + 5’ΔHygR::Intron5'::Read1::TCTCTTTATGATTCA::Read2::Intron3'::3’ΔHygR + 5’ΔHygR::Intron5'::Read1::CATCTTTCTCACCCA::Read2::Intron3'::3’ΔHygR + 5’ΔHygR::Intron5'::Read1::CCTCCCTTTGATAAC::Read2::Intron3'::3’ΔHygR + 5’ΔHygR::Intron5'::Read1::CGACTCTCTTAGAAT::Read2::Intron3'::3’ΔHygR + 5’ΔHygR::Intron5'::Read1::GAGCCCTATAATCAC::Read2::Intron3'::3’ΔHygR + 5’ΔHygR::Intron5'::Read1::GTTCGATTTTATTAG::Read2::Intron3'::3’ΔHygR + 5’ΔHygR::Intron5'::Read1::GAGCCTTATTAACGT::Read2::Intron3'::3’ΔHygR + 5’ΔHygR::Intron5'::Read1::TACCCTTCTAATCAC::Read2::Intron3'::3’ΔHygR + 5’ΔHygR::Intron5'::Read1::ATTCGGTTTCATTTT::Read2::Intron3'::3’ΔHygR + 5’ΔHygR::Intron5'::Read1::TCACCTTGTTATTTC::Read2::Intron3'::3’ΔHygR + 5’ΔHygR::Intron5'::Read1::TTACATTATCAATCA::Read2::Intron3'::3’ΔHygR + 5’ΔHygR::Intron5'::Read1::AGACGTTCTGATATG::Read2::Intron3'::3’ΔHygR + 5’ΔHygR::Intron5'::Read1::TTCCTATTTCAAAAC::Read2::Intron3'::3’ΔHygR + 5’ΔHygR::Intron5'::Read1::ATGCCATGTTACCCT::Read2::Intron3'::3’ΔHygR + 5’ΔHygR::Intron5'::Read1::GCCCGCTATTAGCAC::Read2::Intron3'::3’ΔHygR + 5’ΔHygR::Intron5'::Read1::TCACGGTCTGACTAA::Read2::Intron3'::3’ΔHygR + 5’ΔHygR::Intron5'::Read1::ATACTCTTTCAGAAT::Read2::Intron3'::3’ΔHygR + 5’ΔHygR::Intron5'::Read1::TCACCTTGTCACATT::Read2::Intron3'::3’ΔHygR + 5’ΔHygR::Intron5'::Read1::TCACGCTTTTAAATC::Read2::Intron3'::3’ΔHygR + 5’ΔHygR::Intron5'::Read1::ACTCATTTTAAGATT::Read2::Intron3'::3’ΔHygR +

5’ΔHygR::Intron5'::Read1::CCACAATTTAAGCAA::Read2::Intron3'::3’ΔHygR + 5’ΔHygR::Intron5'::Read1::TCCCCTTCTAACCAA::Read2::Intron3'::3’ΔHygR + 5’ΔHygR::Intron5'::Read1::TCACTATTTAAAAAT::Read2::Intron3'::3’ΔHygR + 5’ΔHygR::Intron5'::Read1::TATCTCTGTTACACA::Read2::Intron3'::3’ΔHygR + 5’ΔHygR::Intron5'::Read1::TCTCACTATAATGGC::Read2::Intron3'::3’ΔHygR + 5’ΔHygR::Intron5'::Read1::GACCTGTGTCATCAT::Read2::Intron3'::3’ΔHygR + 5’ΔHygR::Intron5'::Read1::TCACCATTTGACACA::Read2::Intron3'::3’ΔHygR + 5’ΔHygR::Intron5'::Read1::TGCCTATTTGATTTA::Read2::Intron3'::3’ΔHygR + 5’ΔHygR::Intron5'::Read1::ACACCATTTAACATG::Read2::Intron3'::3’ΔHygR + 5’ΔHygR::Intron5'::Read1::CATCATTCTCAGCTT::Read2::Intron3'::3’ΔHygR + 5’ΔHygR::Intron5'::Read1::CGGCATTCTTAAGTC::Read2::Intron3'::3’ΔHygR + 5’ΔHygR::Intron5'::Read1::AAGCGTTATAAAAAA::Read2::Intron3'::3’ΔHygR + 5’ΔHygR::Intron5'::Read1::CCCCAGTCTGAGTAC::Read2::Intron3'::3’ΔHygR + 5’ΔHygR::Intron5'::Read1::CAACAATGTTAAATT::Read2::Intron3'::3’ΔHygR + 5’ΔHygR::Intron5'::Read1::CATCGCTCTCATGCC::Read2::Intron3'::3’ΔHygR + 5’ΔHygR::Intron5'::Read1::GGTCTTTTTTAATAT::Read2::Intron3'::3’ΔHygR + 5’ΔHygR::Intron5'::Read1::ATACTCTTTGACGCA::Read2::Intron3'::3’ΔHygR + 5’ΔHygR::Intron5'::Read1::TTACTCTATGATCTC::Read2::Intron3'::3’ΔHygR + 5’ΔHygR::Intron5'::Read1::GCTCCATTTTATCCT::Read2::Intron3'::3’ΔHygR + 5’ΔHygR::Intron5'::Read1::TAACTATATCACACA::Read2::Intron3'::3’ΔHygR + 5’ΔHygR::Intron5'::Read1::ATCCCTTTTCATACC::Read2::Intron3'::3’ΔHygR + 5’ΔHygR::Intron5'::Read1::ACACTTTTTTACAGT::Read2::Intron3'::3’ΔHygR + 5’ΔHygR::Intron5'::Read1::TTACATTCTTATTCC::Read2::Intron3'::3’ΔHygR + 5’ΔHygR::Intron5'::Read1::CTGCTCTCTCAAAGT::Read2::Intron3'::3’ΔHygR + 5’ΔHygR::Intron5'::Read1::ATGCAATATCACACC::Read2::Intron3'::3’ΔHygR + 5’ΔHygR::Intron5'::Read1::CTACGATCTCACGAC::Read2::Intron3'::3’ΔHygR + 5’ΔHygR::Intron5'::Read1::TCCCCATTTAAAGCG::Read2::Intron3'::3’ΔHygR + 5’ΔHygR::Intron5'::Read1::CTCCCTTCTCATTCA::Read2::Intron3'::3’ΔHygR + 5’ΔHygR::Intron5'::Read1::GTTCACTATAACAAC::Read2::Intron3'::3’ΔHygR + 5’ΔHygR::Intron5'::Read1::GATCACTGTTAACAC::Read2::Intron3'::3’ΔHygR + 5’ΔHygR::Intron5'::Read1::TACCTGTGTGAATTC::Read2::Intron3'::3’ΔHygR + 5’ΔHygR::Intron5'::Read1::TTGCAATGTCATTTA::Read2::Intron3'::3’ΔHygR + 5’ΔHygR::Intron5'::Read1::CCACTCTCTCACTCA::Read2::Intron3'::3’ΔHygR + 5’ΔHygR::Intron5'::Read1::GTTCAATATTACGAC::Read2::Intron3'::3’ΔHygR + 5’ΔHygR::Intron5'::Read1::ATGCAGTCTCAGTGC::Read2::Intron3'::3’ΔHygR + 5’ΔHygR::Intron5'::Read1::TGTCTCTTTCAATAC::Read2::Intron3'::3’ΔHygR + 5’ΔHygR::Intron5'::Read1::CATCGGTGTGATGCC::Read2::Intron3'::3’ΔHygR + 5’ΔHygR::Intron5'::Read1::TTCCTCTTTAACTGC::Read2::Intron3'::3’ΔHygR + 5’ΔHygR::Intron5'::Read1::TGACCTTTTCAAAGA::Read2::Intron3'::3’ΔHygR + 5’ΔHygR::Intron5'::Read1::TAACATTTTAATCTG::Read2::Intron3'::3’ΔHygR + 5’ΔHygR::Intron5'::Read1::TCACGGTATCATATC::Read2::Intron3'::3’ΔHygR + 5’ΔHygR::Intron5'::Read1::TGGATCTATCACTTC::Read2::Intron3'::3’ΔHygR + 5’ΔHygR::Intron5'::Read1::CAACTGTCTAACTTT::Read2::Intron3'::3’ΔHygR + 5’ΔHygR::Intron5'::Read1::TACCTATGTTAAATT::Read2::Intron3'::3’ΔHygR + 5’ΔHygR::Intron5'::Read1::TCACAATCTAAATCA::Read2::Intron3'::3’ΔHygR + 5’ΔHygR::Intron5'::Read1::CCGCCCTCTTACTCT::Read2::Intron3'::3’ΔHygR + 5’ΔHygR::Intron5'::Read1::CCGCCTTGTAACGTG::Read2::Intron3'::3’ΔHygR + 5’ΔHygR::Intron5'::Read1::TTACTCTCTTAAGTC::Read2::Intron3'::3’ΔHygR + 5’ΔHygR::Intron5'::Read1::CATCCATATCACGTC::Read2::Intron3'::3’ΔHygR + 5’ΔHygR::Intron5'::Read1::TCCCTATCTCATTGA::Read2::Intron3'::3’ΔHygR + 5’ΔHygR::Intron5'::Read1::CCTCTATTTGATTGA::Read2::Intron3'::3’ΔHygR + 5’ΔHygR::Intron5'::Read1::TCTCACTATTACACA::Read2::Intron3'::3’ΔHygR + 5’ΔHygR::Intron5'::Read1::CTACCTTTTCACGAA::Read2::Intron3'::3’ΔHygR + 5’ΔHygR::Intron5'::Read1::GGCCGTTGTGAGCCT::Read2::Intron3'::3’ΔHygR + 5’ΔHygR::Intron5'::Read1::TTGCCCTGTAAGTCA::Read2::Intron3'::3’ΔHygR + 5’ΔHygR::Intron5'::Read1::GGACGTTGTAACTAT::Read2::Intron3'::3’ΔHygR + 5’ΔHygR::Intron5'::Read1::AACCTATATGATTGG::Read2::Intron3'::3’ΔHygR + 5’ΔHygR::Intron5'::Read1::GCTCTATCTAAAAGA::Read2::Intron3'::3’ΔHygR + 5’ΔHygR::Intron5'::Read1::ACTCCTTTTAAACAA::Read2::Intron3'::3’ΔHygR + 5’ΔHygR::Intron5'::Read1::CCCCCATTTAAACGC::Read2::Intron3'::3’ΔHygR + 5’ΔHygR::Intron5'::Read1::GCTCAGTGTTAAACA::Read2::Intron3'::3’ΔHygR + 5’ΔHygR::Intron5'::Read1::TATCGGTTTTATACA::Read2::Intron3'::3’ΔHygR + 5’ΔHygR::Intron5'::Read1::TGACTGTTTCAGTTC::Read2::Intron3'::3’ΔHygR + 5’ΔHygR::Intron5'::Read1::CACCCTTTTAATTGT::Read2::Intron3'::3’ΔHygR + 5’ΔHygR::Intron5'::Read1::ATACTTTATTATTCC::Read2::Intron3'::3’ΔHygR + 5’ΔHygR::Intron5'::Read1::TGACTGTGTAACTAC::Read2::Intron3'::3’ΔHygR + 5’ΔHygR::Intron5'::Read1::CAACAATCTTAACCT::Read2::Intron3'::3’ΔHygR + 5’ΔHygR::Intron5'::Read1::TCGCTGTATGATTAC::Read2::Intron3'::3’ΔHygR + 5’ΔHygR::Intron5'::Read1::TAACGATCTCATGTT::Read2::Intron3'::3’ΔHygR + 5’ΔHygR::Intron5'::Read1::TACCGGTATAACATA::Read2::Intron3'::3’ΔHygR +

5’ΔHygR::Intron5'::Read1::TTGCAGTCTGATCAT::Read2::Intron3'::3’ΔHygR + 5’ΔHygR::Intron5'::Read1::TATCTCTATCACGTG::Read2::Intron3'::3’ΔHygR + 5’ΔHygR::Intron5'::Read1::TCACGATGTAAAAAA::Read2::Intron3'::3’ΔHygR + 5’ΔHygR::Intron5'::Read1::TCGCTTTGTAATGGC::Read2::Intron3'::3’ΔHygR + 5’ΔHygR::Intron5'::Read1::AGCCATTATTACCGG::Read2::Intron3'::3’ΔHygR + 5’ΔHygR::Intron5'::Read1::AGTCATTCTAATATA::Read2::Intron3'::3’ΔHygR + 5’ΔHygR::Intron5'::Read1::GCGCCATTTTAACAG::Read2::Intron3'::3’ΔHygR + 5’ΔHygR::Intron5'::Read1::CTCCTTTTTTATTTT::Read2::Intron3'::3’ΔHygR + 5’ΔHygR::Intron5'::Read1::GACCGCTATCAATCC::Read2::Intron3'::3’ΔHygR + 5’ΔHygR::Intron5'::Read1::GCTCTGTTTGATTTC::Read2::Intron3'::3’ΔHygR + 5’ΔHygR::Intron5'::Read1::TACCTCTATAAGGAT::Read2::Intron3'::3’ΔHygR + 5’ΔHygR::Intron5'::Read1::ACTCTCTATTATGCC::Read2::Intron3'::3’ΔHygR + 5’ΔHygR::Intron5'::Read1::CTACTCTATAAACCC::Read2::Intron3'::3’ΔHygR + 5’ΔHygR::Intron5'::Read1::CCACGTTATTAATAC::Read2::Intron3'::3’ΔHygR + 5’ΔHygR::Intron5'::Read1::TATCTCTATTACCGG::Read2::Intron3'::3’ΔHygR + 5’ΔHygR::Intron5'::Read1::ATTCCCTATTACGAT::Read2::Intron3'::3’ΔHygR + 5’ΔHygR::Intron5'::Read1::GCGCTCTATCAACTA::Read2::Intron3'::3’ΔHygR + 5’ΔHygR::Intron5'::Read1::TATCTCTATCATGTT::Read2::Intron3'::3’ΔHygR + 5’ΔHygR::Intron5'::Read1::TCGCAATTTAATAGC::Read2::Intron3'::3’ΔHygR + 5’ΔHygR::Intron5'::Read1::AAACCATGTAACATT::Read2::Intron3'::3’ΔHygR + 5’ΔHygR::Intron5'::Read1::TCGCCGTATCAAGCC::Read2::Intron3'::3’ΔHygR + 5’ΔHygR::Intron5'::Read1::TCTCGCTATTACTGT::Read2::Intron3'::3’ΔHygR + 5’ΔHygR::Intron5'::Read1::CCCCTTTTTGATCAC::Read2::Intron3'::3’ΔHygR + 5’ΔHygR::Intron5'::Read1::CCACAGTTTAAGGCA::Read2::Intron3'::3’ΔHygR + 5’ΔHygR::Intron5'::Read1::AATCAATCTGAAAAT::Read2::Intron3'::3’ΔHygR + 5’ΔHygR::Intron5'::Read1::TATCAATCTAACAGC::Read2::Intron3'::3’ΔHygR + 5’ΔHygR::Intron5'::Read1::TTCCAATCTCACAAG::Read2::Intron3'::3’ΔHygR + 5’ΔHygR::Intron5'::Read1::TTCCAGTATCATTAC::Read2::Intron3'::3’ΔHygR + 5’ΔHygR::Intron5'::Read1::CAGCTGTCTGACTCA::Read2::Intron3'::3’ΔHygR + 5’ΔHygR::Intron5'::Read1::GCTCTTTGTTATTCC::Read2::Intron3'::3’ΔHygR + 5’ΔHygR::Intron5'::Read1::TCACTTTTTTATTTT::Read2::Intron3'::3’ΔHygR + 5’ΔHygR::Intron5'::Read1::TCACCTTCTAATTCA::Read2::Intron3'::3’ΔHygR + 5’ΔHygR::Intron5'::Read1::CATCCATCTAATTGC::Read2::Intron3'::3’ΔHygR + 5’ΔHygR::Intron5'::Read1::CAACCCTTTGACACC::Read2::Intron3'::3’ΔHygR + 5’ΔHygR::Intron5'::Read1::ACGCCCTATTATGAA::Read2::Intron3'::3’ΔHygR + 5’ΔHygR::Intron5'::Read1::GTCCCATGTTATGTA::Read2::Intron3'::3’ΔHygR + 5’ΔHygR::Intron5'::Read1::CGCCCCTATGATCCC::Read2::Intron3'::3’ΔHygR + 5’ΔHygR::Intron5'::Read1::GTGCTGTCTCATCCT::Read2::Intron3'::3’ΔHygR + 5’ΔHygR::Intron5'::Read1::TTGCACTTTGATTAC::Read2::Intron3'::3’ΔHygR + 5’ΔHygR::Intron5'::Read1::ACACCGTTTGAATTT::Read2::Intron3'::3’ΔHygR + 5’ΔHygR::Intron5'::Read1::TATTTAACATAGACG::Read2::Intron3'::3’ΔHygR + 5’ΔHygR::Intron5'::Read1::TTACGGTATTACCTA::Read2::Intron3'::3’ΔHygR + 5’ΔHygR::Intron5'::Read1::ATCCACTATAAAACC::Read2::Intron3'::3’ΔHygR + 5’ΔHygR::Intron5'::Read1::TGACCCTCTCACTGA::Read2::Intron3'::3’ΔHygR + 5’ΔHygR::Intron5'::Read1::TAACTATTTTATCCT::Read2::Intron3'::3’ΔHygR + 5’ΔHygR::Intron5'::Read1::CGTCCATTTCATTGA::Read2::Intron3'::3’ΔHygR + 5’ΔHygR::Intron5'::Read1::CTTCATTCTCACAAA::Read2::Intron3'::3’ΔHygR + 5’ΔHygR::Intron5'::Read1::GCTCTATTTCAGGTG::Read2::Intron3'::3’ΔHygR + 5’ΔHygR::Intron5'::Read1::CCACTGTTTCATATG::Read2::Intron3'::3’ΔHygR + 5’ΔHygR::Intron5'::Read1::AAACTATTTCAGCCA::Read2::Intron3'::3’ΔHygR + 5’ΔHygR::Intron5'::Read1::TCCCCATCTAAAGTC::Read2::Intron3'::3’ΔHygR + 5’ΔHygR::Intron5'::Read1::TTCCGATATCAGTCG::Read2::Intron3'::3’ΔHygR + 5’ΔHygR::Intron5'::Read1::ACGCTCTGTCAACCT::Read2::Intron3'::3’ΔHygR + 5’ΔHygR::Intron5'::Read1::ACTCTTTTTAAGTTC::Read2::Intron3'::3’ΔHygR + 5’ΔHygR::Intron5'::Read1::TGACATTTTGACGAA::Read2::Intron3'::3’ΔHygR + 5’ΔHygR::Intron5'::Read1::CTACATTTTTAGACT::Read2::Intron3'::3’ΔHygR + 5’ΔHygR::Intron5'::Read1::CTTCAATGTAAACGG::Read2::Intron3'::3’ΔHygR + 5’ΔHygR::Intron5'::Read1::AAACGGTCTTACAGT::Read2::Intron3'::3’ΔHygR + 5’ΔHygR::Intron5'::Read1::TCTCGATCTTAGCAC::Read2::Intron3'::3’ΔHygR + 5’ΔHygR::Intron5'::Read1::CGGCCTTGTGAGATC::Read2::Intron3'::3’ΔHygR + 5’ΔHygR::Intron5'::Read1::TTGCTATTTCAGCTT::Read2::Intron3'::3’ΔHygR + 5’ΔHygR::Intron5'::Read1::GATCACTATAATGTT::Read2::Intron3'::3’ΔHygR + 5’ΔHygR::Intron5'::Read1::AAACATTTTCAAAAT::Read2::Intron3'::3’ΔHygR + 5’ΔHygR::Intron5'::Read1::GGCCATTTTGATTGT::Read2::Intron3'::3’ΔHygR + 5’ΔHygR::Intron5'::Read1::GTCCTATGTCACTTT::Read2::Intron3'::3’ΔHygR + 5’ΔHygR::Intron5'::Read1::TTCCATTTTCACTGC::Read2::Intron3'::3’ΔHygR + 5’ΔHygR::Intron5'::Read1::CCACCTTATCACCCT::Read2::Intron3'::3’ΔHygR + 5’ΔHygR::Intron5'::Read1::TCACTCTTTAATCTC::Read2::Intron3'::3’ΔHygR + 5’ΔHygR::Intron5'::Read1::AAACATTCTCATCAG::Read2::Intron3'::3’ΔHygR + 5’ΔHygR::Intron5'::Read1::TTGCCATATCAGACG::Read2::Intron3'::3’ΔHygR +

5’ΔHygR::Intron5'::Read1::AATCAATTTTACGTA::Read2::Intron3'::3’ΔHygR + 5’ΔHygR::Intron5'::Read1::TCACTTTGTCAACGC::Read2::Intron3'::3’ΔHygR + 5’ΔHygR::Intron5'::Read1::CATCACTGTTACTCC::Read2::Intron3'::3’ΔHygR + 5’ΔHygR::Intron5'::Read1::TCGCCGTTTTATCGC::Read2::Intron3'::3’ΔHygR + 5’ΔHygR::Intron5'::Read1::GTTCTCTTTTATAAC::Read2::Intron3'::3’ΔHygR + 5’ΔHygR::Intron5'::Read1::GACCTATGTGATCCA::Read2::Intron3'::3’ΔHygR + 5’ΔHygR::Intron5'::Read1::CATCACTCTGAACAC::Read2::Intron3'::3’ΔHygR + 5’ΔHygR::Intron5'::Read1::CAACCCTCTAAGCTG::Read2::Intron3'::3’ΔHygR + 5’ΔHygR::Intron5'::Read1::AAACATTCTTACTGT::Read2::Intron3'::3’ΔHygR + 5’ΔHygR::Intron5'::Read1::GGTCCCTGTAACATA::Read2::Intron3'::3’ΔHygR + 5’ΔHygR::Intron5'::Read1::ACCCAGTGTTACTTT::Read2::Intron3'::3’ΔHygR + 5’ΔHygR::Intron5'::Read1::TAGCCTTTTAACTAA::Read2::Intron3'::3’ΔHygR + 5’ΔHygR::Intron5'::Read1::TCACACTTTAAGCTT::Read2::Intron3'::3’ΔHygR + 5’ΔHygR::Intron5'::Read1::AGGCCATTTTAGCAT::Read2::Intron3'::3’ΔHygR + 5’ΔHygR::Intron5'::Read1::CCTCGGTCTGAAACA::Read2::Intron3'::3’ΔHygR + 5’ΔHygR::Intron5'::Read1::GCACTGTCTCACTCA::Read2::Intron3'::3’ΔHygR + 5’ΔHygR::Intron5'::Read1::CTTCTTTCTAAGTCC::Read2::Intron3'::3’ΔHygR + 5’ΔHygR::Intron5'::Read1::GGACTATATAAGAGC::Read2::Intron3'::3’ΔHygR + 5’ΔHygR::Intron5'::Read1::CTCCAGTATGATAGC::Read2::Intron3'::3’ΔHygR + 5’ΔHygR::Intron5'::Read1::TCGCACTATTATGCC::Read2::Intron3'::3’ΔHygR + 5’ΔHygR::Intron5'::Read1::CTGCCGTATTAAAAA::Read2::Intron3'::3’ΔHygR + 5’ΔHygR::Intron5'::Read1::ACTCTTTTTCAGTTT::Read2::Intron3'::3’ΔHygR + 5’ΔHygR::Intron5'::Read1::ATCCATTATAAAGCA::Read2::Intron3'::3’ΔHygR + 5’ΔHygR::Intron5'::Read1::GTACTATATTAGTTT::Read2::Intron3'::3’ΔHygR + 5’ΔHygR::Intron5'::Read1::TTGCACTTTCACGTC::Read2::Intron3'::3’ΔHygR + 5’ΔHygR::Intron5'::Read1::ACCCCCTCTCAGCTA::Read2::Intron3'::3’ΔHygR + 5’ΔHygR::Intron5'::Read1::ACCCGCTATAATTAA::Read2::Intron3'::3’ΔHygR + 5’ΔHygR::Intron5'::Read1::CCCCGATGTTACGTC::Read2::Intron3'::3’ΔHygR + 5’ΔHygR::Intron5'::Read1::CTGCATTGTTAATTA::Read2::Intron3'::3’ΔHygR + 5’ΔHygR::Intron5'::Read1::CACCTTTTTGACGTC::Read2::Intron3'::3’ΔHygR + 5’ΔHygR::Intron5'::Read1::CTCCAGTATTAAGTC::Read2::Intron3'::3’ΔHygR + 5’ΔHygR::Intron5'::Read1::GAGCTTTTTCAAACA::Read2::Intron3'::3’ΔHygR + 5’ΔHygR::Intron5'::Read1::ATACTATCTCACAAT::Read2::Intron3'::3’ΔHygR + 5’ΔHygR::Intron5'::Read1::TACCTCTCTTAAGTC::Read2::Intron3'::3’ΔHygR + 5’ΔHygR::Intron5'::Read1::TTGCCCTATAACCTG::Read2::Intron3'::3’ΔHygR + 5’ΔHygR::Intron5'::Read1::GATCCATATTAACGG::Read2::Intron3'::3’ΔHygR + 5’ΔHygR::Intron5'::Read1::AAGCTCTGTTATTTG::Read2::Intron3'::3’ΔHygR + 5’ΔHygR::Intron5'::Read1::CAACCCTTTTACGAA::Read2::Intron3'::3’ΔHygR + 5’ΔHygR::Intron5'::Read1::GCGCTTTCTTATCCA::Read2::Intron3'::3’ΔHygR + 5’ΔHygR::Intron5'::Read1::TCTCCTTTTTACTCC::Read2::Intron3'::3’ΔHygR + 5’ΔHygR::Intron5'::Read1::GGACTTTTTGATCTC::Read2::Intron3'::3’ΔHygR + 5’ΔHygR::Intron5'::Read1::GTTCTCTGTAATATC::Read2::Intron3'::3’ΔHygR + 5’ΔHygR::Intron5'::Read1::TGTCTCTGTCAGGTG::Read2::Intron3'::3’ΔHygR + 5’ΔHygR::Intron5'::Read1::AATCGTTATTATTTT::Read2::Intron3'::3’ΔHygR + 5’ΔHygR::Intron5'::Read1::CTCCAATTTCAGCAA::Read2::Intron3'::3’ΔHygR + 5’ΔHygR::Intron5'::Read1::GGCCCCTTTGATTCA::Read2::Intron3'::3’ΔHygR + 5’ΔHygR::Intron5'::Read1::TCACTGTTTCAAGTC::Read2::Intron3'::3’ΔHygR + 5’ΔHygR::Intron5'::Read1::GCCCTATCTCATCCT::Read2::Intron3'::3’ΔHygR + 5’ΔHygR::Intron5'::Read1::CGACAATCTTATTCC::Read2::Intron3'::3’ΔHygR + 5’ΔHygR::Intron5'::Read1::TGTCGATATGAGGAG::Read2::Intron3'::3’ΔHygR + 5’ΔHygR::Intron5'::Read1::ACTCTCTCTTACCCT::Read2::Intron3'::3’ΔHygR + 5’ΔHygR::Intron5'::Read1::CGGCTATATGAATTT::Read2::Intron3'::3’ΔHygR + 5’ΔHygR::Intron5'::Read1::AGGCAATTTTACAAC::Read2::Intron3'::3’ΔHygR + 5’ΔHygR::Intron5'::Read1::AAGCCTTTTGAACCG::Read2::Intron3'::3’ΔHygR + 5’ΔHygR::Intron5'::Read1::AATCAGTCTGACATA::Read2::Intron3'::3’ΔHygR + 5’ΔHygR::Intron5'::Read1::ATTCTTTCTCACAAT::Read2::Intron3'::3’ΔHygR + 5’ΔHygR::Intron5'::Read1::CTTCAATTTTATTCG::Read2::Intron3'::3’ΔHygR + 5’ΔHygR::Intron5'::Read1::GTTCATTTTAACTGC::Read2::Intron3'::3’ΔHygR + 5’ΔHygR::Intron5'::Read1::TCACAATATTACTGC::Read2::Intron3'::3’ΔHygR + 5’ΔHygR::Intron5'::Read1::CGACATTTTTATGCT::Read2::Intron3'::3’ΔHygR + 5’ΔHygR::Intron5'::Read1::ATACTATCTTAATTA::Read2::Intron3'::3’ΔHygR + 5’ΔHygR::Intron5'::Read1::GCACAATGTGAACTA::Read2::Intron3'::3’ΔHygR + 5’ΔHygR::Intron5'::Read1::ACTCCTTCTCATAAT::Read2::Intron3'::3’ΔHygR + 5’ΔHygR::Intron5'::Read1::CCGCTGTTTTAAGTT::Read2::Intron3'::3’ΔHygR + 5’ΔHygR::Intron5'::Read1::CGACCATTTCATCTG::Read2::Intron3'::3’ΔHygR + 5’ΔHygR::Intron5'::Read1::TACCAATCTCATCTG::Read2::Intron3'::3’ΔHygR + 5’ΔHygR::Intron5'::Read1::GATCGCTTTCACTTT::Read2::Intron3'::3’ΔHygR + 5’ΔHygR::Intron5'::Read1::TTCCGTTATAAGCAT::Read2::Intron3'::3’ΔHygR + 5’ΔHygR::Intron5'::Read1::TCTCCATCTCACAAA::Read2::Intron3'::3’ΔHygR + 5’ΔHygR::Intron5'::Read1::AATCGATCTAAGCTT::Read2::Intron3'::3’ΔHygR +

5’ΔHygR::Intron5'::Read1::CAACAGTTTTACTAA::Read2::Intron3'::3’ΔHygR + 5’ΔHygR::Intron5'::Read1::ATACAGTATCATACG::Read2::Intron3'::3’ΔHygR + 5’ΔHygR::Intron5'::Read1::TTCCGCTATAACTTT::Read2::Intron3'::3’ΔHygR + 5’ΔHygR::Intron5'::Read1::CGCCTCTTTGATGCT::Read2::Intron3'::3’ΔHygR + 5’ΔHygR::Intron5'::Read1::GAACCATATCATGAC::Read2::Intron3'::3’ΔHygR + 5’ΔHygR::Intron5'::Read1::ATGCGGTATGACCGA::Read2::Intron3'::3’ΔHygR + 5’ΔHygR::Intron5'::Read1::ACACTATATGAGAAT::Read2::Intron3'::3’ΔHygR + 5’ΔHygR::Intron5'::Read1::GTTCGCTTTGACATT::Read2::Intron3'::3’ΔHygR + 5’ΔHygR::Intron5'::Read1::TAGCATTTTTATCAG::Read2::Intron3'::3’ΔHygR + 5’ΔHygR::Intron5'::Read1::ATACAATTTAATCTC::Read2::Intron3'::3’ΔHygR + 5’ΔHygR::Intron5'::Read1::CTACGTTCTCACGCG::Read2::Intron3'::3’ΔHygR + 5’ΔHygR::Intron5'::Read1::CAACCGTATGATTAC::Read2::Intron3'::3’ΔHygR + 5’ΔHygR::Intron5'::Read1::ACACCTTCTAACGCT::Read2::Intron3'::3’ΔHygR + 5’ΔHygR::Intron5'::Read1::CCACTTTATTAGAAA::Read2::Intron3'::3’ΔHygR + 5’ΔHygR::Intron5'::Read1::ACCCTTTCTAATTCA::Read2::Intron3'::3’ΔHygR + 5’ΔHygR::Intron5'::Read1::CATCGTTATGATACG::Read2::Intron3'::3’ΔHygR + 5’ΔHygR::Intron5'::Read1::GTACTTTTTGAAAAG::Read2::Intron3'::3’ΔHygR + 5’ΔHygR::Intron5'::Read1::TTTCCTTCTTACCAC::Read2::Intron3'::3’ΔHygR + 5’ΔHygR::Intron5'::Read1::TTACCTTATGACATC::Read2::Intron3'::3’ΔHygR + 5’ΔHygR::Intron5'::Read1::CACCGATCTTAGTTC::Read2::Intron3'::3’ΔHygR + 5’ΔHygR::Intron5'::Read1::ACTCTATGTCAACCA::Read2::Intron3'::3’ΔHygR + 5’ΔHygR::Intron5'::Read1::CCCCAATATGACGAA::Read2::Intron3'::3’ΔHygR + 5’ΔHygR::Intron5'::Read1::GAGCTCTCTTAAATC::Read2::Intron3'::3’ΔHygR + 5’ΔHygR::Intron5'::Read1::TGGCTGTCTTATCTT::Read2::Intron3'::3’ΔHygR + 5’ΔHygR::Intron5'::Read1::GCTCTATCTTAACCC::Read2::Intron3'::3’ΔHygR + 5’ΔHygR::Intron5'::Read1::GCGCCCTGTCAGTCA::Read2::Intron3'::3’ΔHygR + 5’ΔHygR::Intron5'::Read1::TGTCATTTTTATCCT::Read2::Intron3'::3’ΔHygR + 5’ΔHygR::Intron5'::Read1::TCTCTATGTTACTGA::Read2::Intron3'::3’ΔHygR + 5’ΔHygR::Intron5'::Read1::TGTCAATCTTACATA::Read2::Intron3'::3’ΔHygR + 5’ΔHygR::Intron5'::Read1::GCCCCTTTTAAGGAT::Read2::Intron3'::3’ΔHygR + 5’ΔHygR::Intron5'::Read1::CCCCATTCTTACCCT::Read2::Intron3'::3’ΔHygR + 5’ΔHygR::Intron5'::Read1::CCACATTCTGATGTC::Read2::Intron3'::3’ΔHygR + 5’ΔHygR::Intron5'::Read1::CCCCTCTCTAAGATG::Read2::Intron3'::3’ΔHygR + 5’ΔHygR::Intron5'::Read1::CTCCCCTTTTACATC::Read2::Intron3'::3’ΔHygR + 5’ΔHygR::Intron5'::Read1::AATCTTTGTGATCAG::Read2::Intron3'::3’ΔHygR + 5’ΔHygR::Intron5'::Read1::ACTCAATCTAATTGC::Read2::Intron3'::3’ΔHygR + 5’ΔHygR::Intron5'::Read1::ATATTAACAACGGGG::Read2::Intron3'::3’ΔHygR + 5’ΔHygR::Intron5'::Read1::TACCAATCTTAGTCG::Read2::Intron3'::3’ΔHygR + 5’ΔHygR::Intron5'::Read1::ACACACTATTAGTTG::Read2::Intron3'::3’ΔHygR + 5’ΔHygR::Intron5'::Read1::AGACATTTTCATACA::Read2::Intron3'::3’ΔHygR + 5’ΔHygR::Intron5'::Read1::TCACCATGTCATTCA::Read2::Intron3'::3’ΔHygR + 5’ΔHygR::Intron5'::Read1::GTTCAATGTCACCGC::Read2::Intron3'::3’ΔHygR + 5’ΔHygR::Intron5'::Read1::AGGCCTTATGAGCAT::Read2::Intron3'::3’ΔHygR + 5’ΔHygR::Intron5'::Read1::CGCCTCTTTTAAAGC::Read2::Intron3'::3’ΔHygR + 5’ΔHygR::Intron5'::Read1::CTTCCCTTTGAAAGA::Read2::Intron3'::3’ΔHygR + 5’ΔHygR::Intron5'::Read1::ACGCGTTATAACTCA::Read2::Intron3'::3’ΔHygR + 5’ΔHygR::Intron5'::Read1::CATCTTTATCATTGG::Read2::Intron3'::3’ΔHygR + 5’ΔHygR::Intron5'::Read1::ACACCCTTTCAGCCG::Read2::Intron3'::3’ΔHygR + 5’ΔHygR::Intron5'::Read1::GAACTATATCATCCG::Read2::Intron3'::3’ΔHygR + 5’ΔHygR::Intron5'::Read1::GTGCAGTCTTAAATT::Read2::Intron3'::3’ΔHygR + 5’ΔHygR::Intron5'::Read1::GTTCCCTCTAACGCA::Read2::Intron3'::3’ΔHygR + 5’ΔHygR::Intron5'::Read1::TGCCAATATCAGCAT::Read2::Intron3'::3’ΔHygR + 5’ΔHygR::Intron5'::Read1::CGGCTTTTTAATTTA::Read2::Intron3'::3’ΔHygR + 5’ΔHygR::Intron5'::Read1::CCCCTGTCTAAGACC::Read2::Intron3'::3’ΔHygR + 5’ΔHygR::Intron5'::Read1::ACGCTATGTCAGGCC::Read2::Intron3'::3’ΔHygR + 5’ΔHygR::Intron5'::Read1::TGACTATATAACGCG::Read2::Intron3'::3’ΔHygR + 5’ΔHygR::Intron5'::Read1::CTACACTCTGATTAA::Read2::Intron3'::3’ΔHygR + 5’ΔHygR::Intron5'::Read1::GGTCCTTTTAATCTG::Read2::Intron3'::3’ΔHygR + 5’ΔHygR::Intron5'::Read1::AAGCGGTTTTACCTA::Read2::Intron3'::3’ΔHygR + 5’ΔHygR::Intron5'::Read1::GCCCTGTCTAAGTAA::Read2::Intron3'::3’ΔHygR + 5’ΔHygR::Intron5'::Read1::GTTCAGTATCATCCC::Read2::Intron3'::3’ΔHygR + 5’ΔHygR::Intron5'::Read1::TGTCTCTCTCATAAT::Read2::Intron3'::3’ΔHygR + 5’ΔHygR::Intron5'::Read1::CTACTCTGTGATAGT::Read2::Intron3'::3’ΔHygR + 5’ΔHygR::Intron5'::Read1::GACCGCTTTCACTAA::Read2::Intron3'::3’ΔHygR + 5’ΔHygR::Intron5'::Read1::GGCCCTTATTAAATA::Read2::Intron3'::3’ΔHygR + 5’ΔHygR::Intron5'::Read1::GGCCCGTCTCACAGT::Read2::Intron3'::3’ΔHygR + 5’ΔHygR::Intron5'::Read1::TCCCGATATTACGTC::Read2::Intron3'::3’ΔHygR + 5’ΔHygR::Intron5'::Read1::ACACCTTATTAGATC::Read2::Intron3'::3’ΔHygR + 5’ΔHygR::Intron5'::Read1::ACGCAGTGTCAGACT::Read2::Intron3'::3’ΔHygR + 5’ΔHygR::Intron5'::Read1::AAACCTTTTCAACAA::Read2::Intron3'::3’ΔHygR +

5’ΔHygR::Intron5'::Read1::CCCCTTTCTTATGCA::Read2::Intron3'::3’ΔHygR + 5’ΔHygR::Intron5'::Read1::ATCCGTTATAAGCAA::Read2::Intron3'::3’ΔHygR + 5’ΔHygR::Intron5'::Read1::CGTCACTTTCAAAGA::Read2::Intron3'::3’ΔHygR + 5’ΔHygR::Intron5'::Read1::TCTCCCTTTAATTGT::Read2::Intron3'::3’ΔHygR + 5’ΔHygR::Intron5'::Read1::ACCCCTTTTGAAAAG::Read2::Intron3'::3’ΔHygR + 5’ΔHygR::Intron5'::Read1::ACCCACTGTCACTGC::Read2::Intron3'::3’ΔHygR + 5’ΔHygR::Intron5'::Read1::TCTCCTTCTAAGCCT::Read2::Intron3'::3’ΔHygR + 5’ΔHygR::Intron5'::Read1::CAACCATCTTAGCGC::Read2::Intron3'::3’ΔHygR + 5’ΔHygR::Intron5'::Read1::GGTCCCTTTAAATGA::Read2::Intron3'::3’ΔHygR + 5’ΔHygR::Intron5'::Read1::TACCGTTCTGAAAAC::Read2::Intron3'::3’ΔHygR + 5’ΔHygR::Intron5'::Read1::CGGCCCTATAAATAA::Read2::Intron3'::3’ΔHygR + 5’ΔHygR::Intron5'::Read1::TTCCATTTTAAGCCG::Read2::Intron3'::3’ΔHygR + 5’ΔHygR::Intron5'::Read1::TCCCCATTTCAACTA::Read2::Intron3'::3’ΔHygR + 5’ΔHygR::Intron5'::Read1::AAACTCTCTAACTAA::Read2::Intron3'::3’ΔHygR + 5’ΔHygR::Intron5'::Read1::TCTCGCTATCATTTA::Read2::Intron3'::3’ΔHygR + 5’ΔHygR::Intron5'::Read1::TTCCTTTTTCACCTT::Read2::Intron3'::3’ΔHygR + 5’ΔHygR::Intron5'::Read1::CACCTCTCTGAGGCC::Read2::Intron3'::3’ΔHygR + 5’ΔHygR::Intron5'::Read1::CGTCATTATAAACAG::Read2::Intron3'::3’ΔHygR + 5’ΔHygR::Intron5'::Read1::CTCCCTTTTAACACC::Read2::Intron3'::3’ΔHygR + 5’ΔHygR::Intron5'::Read1::CTTCTATTTCACCCA::Read2::Intron3'::3’ΔHygR + 5’ΔHygR::Intron5'::Read1::ATCCCATGTCAAACG::Read2::Intron3'::3’ΔHygR + 5’ΔHygR::Intron5'::Read1::TCCCGGTCTCAGTTG::Read2::Intron3'::3’ΔHygR + 5’ΔHygR::Intron5'::Read1::GTTCTATTTAAAACT::Read2::Intron3'::3’ΔHygR + 5’ΔHygR::Intron5'::Read1::GTCCAATCTTAGATC::Read2::Intron3'::3’ΔHygR + 5’ΔHygR::Intron5'::Read1::TTTCACTCTGACTTA::Read2::Intron3'::3’ΔHygR + 5’ΔHygR::Intron5'::Read1::AAACCTTTTAAACCA::Read2::Intron3'::3’ΔHygR + 5’ΔHygR::Intron5'::Read1::AGACCGTCTCATAAA::Read2::Intron3'::3’ΔHygR + 5’ΔHygR::Intron5'::Read1::CATCCCTCTAACACC::Read2::Intron3'::3’ΔHygR + 5’ΔHygR::Intron5'::Read1::TCTCACTATTACATG::Read2::Intron3'::3’ΔHygR + 5’ΔHygR::Intron5'::Read1::TAACAGTATCAACTC::Read2::Intron3'::3’ΔHygR + 5’ΔHygR::Intron5'::Read1::AACCAATATGACGGG::Read2::Intron3'::3’ΔHygR + 5’ΔHygR::Intron5'::Read1::TACCTATCTAACATC::Read2::Intron3'::3’ΔHygR + 5’ΔHygR::Intron5'::Read1::TGTCATTCTCATCCT::Read2::Intron3'::3’ΔHygR + 5’ΔHygR::Intron5'::Read1::ACCCGTTATGAGCAT::Read2::Intron3'::3’ΔHygR + 5’ΔHygR::Intron5'::Read1::ACGCACTATGATACC::Read2::Intron3'::3’ΔHygR + 5’ΔHygR::Intron5'::Read1::CATCGGTTTCATACA::Read2::Intron3'::3’ΔHygR + 5’ΔHygR::Intron5'::Read1::GAACTTTATAAACCC::Read2::Intron3'::3’ΔHygR + 5’ΔHygR::Intron5'::Read1::ACGCAATATAAACAA::Read2::Intron3'::3’ΔHygR + 5’ΔHygR::Intron5'::Read1::ACGCTATGTAAGCCT::Read2::Intron3'::3’ΔHygR + 5’ΔHygR::Intron5'::Read1::CAACTATATTATCTC::Read2::Intron3'::3’ΔHygR + 5’ΔHygR::Intron5'::Read1::CGGCTGTATTACACT::Read2::Intron3'::3’ΔHygR + 5’ΔHygR::Intron5'::Read1::CGTCATTTTGATTAC::Read2::Intron3'::3’ΔHygR + 5’ΔHygR::Intron5'::Read1::TTACGCTCTCAGTCT::Read2::Intron3'::3’ΔHygR + 5’ΔHygR::Intron5'::Read1::AACCAGTATCATATA::Read2::Intron3'::3’ΔHygR + 5’ΔHygR::Intron5'::Read1::CCACGGTATCAGAAA::Read2::Intron3'::3’ΔHygR + 5’ΔHygR::Intron5'::Read1::TACCATTATCATAAC::Read2::Intron3'::3’ΔHygR + 5’ΔHygR::Intron5'::Read1::TTCCTCTATTATCAA::Read2::Intron3'::3’ΔHygR + 5’ΔHygR::Intron5'::Read1::AAACTTTATTAAACA::Read2::Intron3'::3’ΔHygR + 5’ΔHygR::Intron5'::Read1::AGCCTGTGTAATTCC::Read2::Intron3'::3’ΔHygR + 5’ΔHygR::Intron5'::Read1::CCCCCTTTTGAAAAG::Read2::Intron3'::3’ΔHygR + 5’ΔHygR::Intron5'::Read1::CGCCAGTCTTAAAAC::Read2::Intron3'::3’ΔHygR + 5’ΔHygR::Intron5'::Read1::TCCCTTTTTTACAAC::Read2::Intron3'::3’ΔHygR + 5’ΔHygR::Intron5'::Read1::CACCTATTTAACAGA::Read2::Intron3'::3’ΔHygR + 5’ΔHygR::Intron5'::Read1::CCCCGTTCTAACCCT::Read2::Intron3'::3’ΔHygR + 5’ΔHygR::Intron5'::Read1::ACCCTATCTTACCAG::Read2::Intron3'::3’ΔHygR + 5’ΔHygR::Intron5'::Read1::ACTCTTTTTCATCCT::Read2::Intron3'::3’ΔHygR + 5’ΔHygR::Intron5'::Read1::GCACGCTCTAACAGT::Read2::Intron3'::3’ΔHygR + 5’ΔHygR::Intron5'::Read1::AAGCCCTATAAGCCA::Read2::Intron3'::3’ΔHygR + 5’ΔHygR::Intron5'::Read1::CTCCATTTTTACGAT::Read2::Intron3'::3’ΔHygR + 5’ΔHygR::Intron5'::Read1::CTCCTTTCTCATGCT::Read2::Intron3'::3’ΔHygR + 5’ΔHygR::Intron5'::Read1::CCTCGTTGTAACTTA::Read2::Intron3'::3’ΔHygR + 5’ΔHygR::Intron5'::Read1::TTCCGCTATGAACAG::Read2::Intron3'::3’ΔHygR + 5’ΔHygR::Intron5'::Read1::CAACTATATTATCGC::Read2::Intron3'::3’ΔHygR + 5’ΔHygR::Intron5'::Read1::GTTCAATCTTACCGT::Read2::Intron3'::3’ΔHygR + 5’ΔHygR::Intron5'::Read1::TAGCGCTGTAAGTCA::Read2::Intron3'::3’ΔHygR + 5’ΔHygR::Intron5'::Read1::AGACTGTCTTATCTC::Read2::Intron3'::3’ΔHygR + 5’ΔHygR::Intron5'::Read1::ATCCCCTCTTATTAC::Read2::Intron3'::3’ΔHygR + 5’ΔHygR::Intron5'::Read1::GATCCCTGTAACTCT::Read2::Intron3'::3’ΔHygR + 5’ΔHygR::Intron5'::Read1::TCACGCTGTTAAGAG::Read2::Intron3'::3’ΔHygR + 5’ΔHygR::Intron5'::Read1::TGTCCTTCTAATGCA::Read2::Intron3'::3’ΔHygR +

5’ΔHygR::Intron5'::Read1::CAGCGTTATGACATC::Read2::Intron3'::3’ΔHygR + 5’ΔHygR::Intron5'::Read1::CTCCGGTCTTACCAA::Read2::Intron3'::3’ΔHygR + 5’ΔHygR::Intron5'::Read1::TCTCCTTTTGACTGC::Read2::Intron3'::3’ΔHygR + 5’ΔHygR::Intron5'::Read1::ATTCTTTGTTATCCA::Read2::Intron3'::3’ΔHygR + 5’ΔHygR::Intron5'::Read1::TATCTCTCTCAGGAA::Read2::Intron3'::3’ΔHygR + 5’ΔHygR::Intron5'::Read1::GGCCATTCTGAAATT::Read2::Intron3'::3’ΔHygR + 5’ΔHygR::Intron5'::Read1::TGACCATATTAAATT::Read2::Intron3'::3’ΔHygR + 5’ΔHygR::Intron5'::Read1::ACTCTGTATTATCTT::Read2::Intron3'::3’ΔHygR + 5’ΔHygR::Intron5'::Read1::TTCCGATTTTATAAA::Read2::Intron3'::3’ΔHygR + 5’ΔHygR::Intron5'::Read1::AACCGTTGTTATAGT::Read2::Intron3'::3’ΔHygR + 5’ΔHygR::Intron5'::Read1::CATCTTTCTGAATTA::Read2::Intron3'::3’ΔHygR + 5’ΔHygR::Intron5'::Read1::ACCCCGTCTAAAGAC::Read2::Intron3'::3’ΔHygR + 5’ΔHygR::Intron5'::Read1::ACGCATTCTCAATCC::Read2::Intron3'::3’ΔHygR + 5’ΔHygR::Intron5'::Read1::TTTCCATTTCAGACC::Read2::Intron3'::3’ΔHygR + 5’ΔHygR::Intron5'::Read1::CATCAATATCATCCA::Read2::Intron3'::3’ΔHygR + 5’ΔHygR::Intron5'::Read1::CGCCTCTGTAATCAA::Read2::Intron3'::3’ΔHygR + 5’ΔHygR::Intron5'::Read1::TGACATTTTAATCTG::Read2::Intron3'::3’ΔHygR + 5’ΔHygR::Intron5'::Read1::TACCCATATCAGTTC::Read2::Intron3'::3’ΔHygR + 5’ΔHygR::Intron5'::Read1::TCTCTTTATTAATCA::Read2::Intron3'::3’ΔHygR + 5’ΔHygR::Intron5'::Read1::ACACGGTCTGATAAG::Read2::Intron3'::3’ΔHygR + 5’ΔHygR::Intron5'::Read1::CGTCCGTCTCAAAGT::Read2::Intron3'::3’ΔHygR + 5’ΔHygR::Intron5'::Read1::GCTCGGTTTCAACCT::Read2::Intron3'::3’ΔHygR + 5’ΔHygR::Intron5'::Read1::CTGCCATATCATTCT::Read2::Intron3'::3’ΔHygR + 5’ΔHygR::Intron5'::Read1::GCACCGTCTTAATCC::Read2::Intron3'::3’ΔHygR + 5’ΔHygR::Intron5'::Read1::TCTCTATTTTATATC::Read2::Intron3'::3’ΔHygR + 5’ΔHygR::Intron5'::Read1::TGGCCATCTTAAAGT::Read2::Intron3'::3’ΔHygR + 5’ΔHygR::Intron5'::Read1::ACCCGCTGTCAGTTC::Read2::Intron3'::3’ΔHygR + 5’ΔHygR::Intron5'::Read1::CAACGCTCTCACATC::Read2::Intron3'::3’ΔHygR + 5’ΔHygR::Intron5'::Read1::TGTCGCTTTTACTGT::Read2::Intron3'::3’ΔHygR + 5’ΔHygR::Intron5'::Read1::GACCGCTTTAACATT::Read2::Intron3'::3’ΔHygR + 5’ΔHygR::Intron5'::Read1::TTACCATGTAACCGA::Read2::Intron3'::3’ΔHygR + 5’ΔHygR::Intron5'::Read1::TTTCGCTGTTATGCA::Read2::Intron3'::3’ΔHygR + 5’ΔHygR::Intron5'::Read1::ACTCTTTGTGAACCA::Read2::Intron3'::3’ΔHygR + 5’ΔHygR::Intron5'::Read1::GCCCCTTTTAATGTT::Read2::Intron3'::3’ΔHygR + 5’ΔHygR::Intron5'::Read1::CGTCGTTCTTATTTC::Read2::Intron3'::3’ΔHygR + 5’ΔHygR::Intron5'::Read1::AAGCCATGTTATATA::Read2::Intron3'::3’ΔHygR + 5’ΔHygR::Intron5'::Read1::TCACAATATTACATA::Read2::Intron3'::3’ΔHygR + 5’ΔHygR::Intron5'::Read1::CTCCATTATCATATC::Read2::Intron3'::3’ΔHygR + 5’ΔHygR::Intron5'::Read1::TTCCGTTTTCACAGA::Read2::Intron3'::3’ΔHygR + 5’ΔHygR::Intron5'::Read1::TTGCTCTCTTACAAG::Read2::Intron3'::3’ΔHygR + 5’ΔHygR::Intron5'::Read1::CACCGTTCTCATGCT::Read2::Intron3'::3’ΔHygR + 5’ΔHygR::Intron5'::Read1::CTACTATGTGATGCT::Read2::Intron3'::3’ΔHygR + 5’ΔHygR::Intron5'::Read1::GCCCTATATGACCTG::Read2::Intron3'::3’ΔHygR + 5’ΔHygR::Intron5'::Read1::TAGCGATTTGATGAC::Read2::Intron3'::3’ΔHygR + 5’ΔHygR::Intron5'::Read1::AGTCACTGTTATGAC::Read2::Intron3'::3’ΔHygR + 5’ΔHygR::Intron5'::Read1::ATACCTTGTTATTAT::Read2::Intron3'::3’ΔHygR + 5’ΔHygR::Intron5'::Read1::ATTCGCTGTTATTAC::Read2::Intron3'::3’ΔHygR + 5’ΔHygR::Intron5'::Read1::CTCCTCTGTCAATTA::Read2::Intron3'::3’ΔHygR + 5’ΔHygR::Intron5'::Read1::GAACGATCTCAACAA::Read2::Intron3'::3’ΔHygR + 5’ΔHygR::Intron5'::Read1::GATCGTTCTCAACAG::Read2::Intron3'::3’ΔHygR + 5’ΔHygR::Intron5'::Read1::AGTCGATATAACGCC::Read2::Intron3'::3’ΔHygR + 5’ΔHygR::Intron5'::Read1::ACACCCTCTAAATTA::Read2::Intron3'::3’ΔHygR + 5’ΔHygR::Intron5'::Read1::CATCCTTATGAACGC::Read2::Intron3'::3’ΔHygR + 5’ΔHygR::Intron5'::Read1::CATCTTTATTAAATT::Read2::Intron3'::3’ΔHygR + 5’ΔHygR::Intron5'::Read1::TCCCGGTTTTACTGT::Read2::Intron3'::3’ΔHygR + 5’ΔHygR::Intron5'::Read1::CATCATTTTAAAACA::Read2::Intron3'::3’ΔHygR + 5’ΔHygR::Intron5'::Read1::GTACATTTTAATTTA::Read2::Intron3'::3’ΔHygR + 5’ΔHygR::Intron5'::Read1::AGACTTTTTGATACA::Read2::Intron3'::3’ΔHygR + 5’ΔHygR::Intron5'::Read1::ATTCAATATCACTGG::Read2::Intron3'::3’ΔHygR + 5’ΔHygR::Intron5'::Read1::ATCCTTTATTATACT::Read2::Intron3'::3’ΔHygR + 5’ΔHygR::Intron5'::Read1::CGACTTTATAAACTA::Read2::Intron3'::3’ΔHygR + 5’ΔHygR::Intron5'::Read1::CGGCAGTATCATCAA::Read2::Intron3'::3’ΔHygR + 5’ΔHygR::Intron5'::Read1::GCCCAGTCTTAGCGA::Read2::Intron3'::3’ΔHygR + 5’ΔHygR::Intron5'::Read1::TTGCAGTATGAGTGT::Read2::Intron3'::3’ΔHygR + 5’ΔHygR::Intron5'::Read1::GTAGTTAGAATGAAA::Read2::Intron3'::3’ΔHygR + 5’ΔHygR::Intron5'::Read1::TTTCTTTTTGACTTT::Read2::Intron3'::3’ΔHygR + 5’ΔHygR::Intron5'::Read1::TGCCACTATGACACC::Read2::Intron3'::3’ΔHygR + 5’ΔHygR::Intron5'::Read1::CGGCTCTGTAATCCA::Read2::Intron3'::3’ΔHygR + 5’ΔHygR::Intron5'::Read1::TAACATTCTTATAAT::Read2::Intron3'::3’ΔHygR + 5’ΔHygR::Intron5'::Read1::CGCCTTTATAATCCT::Read2::Intron3'::3’ΔHygR +

5’ΔHygR::Intron5'::Read1::GCGCAATATGAAGTC::Read2::Intron3'::3’ΔHygR + 5’ΔHygR::Intron5'::Read1::TACCTCTTTCAAACA::Read2::Intron3'::3’ΔHygR + 5’ΔHygR::Intron5'::Read1::CATCTTTTTAAAACC::Read2::Intron3'::3’ΔHygR + 5’ΔHygR::Intron5'::Read1::TACCAATTTAACAAC::Read2::Intron3'::3’ΔHygR + 5’ΔHygR::Intron5'::Read1::GTACATTATGACTTC::Read2::Intron3'::3’ΔHygR + 5’ΔHygR::Intron5'::Read1::TATCCGTTTTACGCA::Read2::Intron3'::3’ΔHygR + 5’ΔHygR::Intron5'::Read1::TCTCTTTATCACCAC::Read2::Intron3'::3’ΔHygR + 5’ΔHygR::Intron5'::Read1::CGCCAATTTGATTAT::Read2::Intron3'::3’ΔHygR + 5’ΔHygR::Intron5'::Read1::TGCCCCTTTAATATA::Read2::Intron3'::3’ΔHygR + 5’ΔHygR::Intron5'::Read1::TTCCTTTTTGACTAA::Read2::Intron3'::3’ΔHygR + 5’ΔHygR::Intron5'::Read1::ATACACTCTCAATTC::Read2::Intron3'::3’ΔHygR + 5’ΔHygR::Intron5'::Read1::AGCCGCTATCATACA::Read2::Intron3'::3’ΔHygR + 5’ΔHygR::Intron5'::Read1::TGTCAGTTTTAATAC::Read2::Intron3'::3’ΔHygR + 5’ΔHygR::Intron5'::Read1::CTTCATTATAAAAGC::Read2::Intron3'::3’ΔHygR + 5’ΔHygR::Intron5'::Read1::GCTCAATATCAGTAT::Read2::Intron3'::3’ΔHygR + 5’ΔHygR::Intron5'::Read1::CTACAATGTCAATTC::Read2::Intron3'::3’ΔHygR + 5’ΔHygR::Intron5'::Read1::AGGCAGTATAACACG::Read2::Intron3'::3’ΔHygR + 5’ΔHygR::Intron5'::Read1::CTCCGCTATAAAGAC::Read2::Intron3'::3’ΔHygR + 5’ΔHygR::Intron5'::Read1::TCCCTCTTTTACATC::Read2::Intron3'::3’ΔHygR + 5’ΔHygR::Intron5'::Read1::TAGCATTATTACCAT::Read2::Intron3'::3’ΔHygR + 5’ΔHygR::Intron5'::Read1::ATTCAGTCTGACAGC::Read2::Intron3'::3’ΔHygR + 5’ΔHygR::Intron5'::Read1::GTTCTATCTCAAACA::Read2::Intron3'::3’ΔHygR + 5’ΔHygR::Intron5'::Read1::TCGCTCTTTGATGTC::Read2::Intron3'::3’ΔHygR + 5’ΔHygR::Intron5'::Read1::CGACTGTGTCATGCA::Read2::Intron3'::3’ΔHygR + 5’ΔHygR::Intron5'::Read1::AACCTTTATTATCTA::Read2::Intron3'::3’ΔHygR + 5’ΔHygR::Intron5'::Read1::ATTCTATATTAAGAC::Read2::Intron3'::3’ΔHygR + 5’ΔHygR::Intron5'::Read1::GACCAATTTCATCCA::Read2::Intron3'::3’ΔHygR + 5’ΔHygR::Intron5'::Read1::TGTCAATGTTAAGTT::Read2::Intron3'::3’ΔHygR + 5’ΔHygR::Intron5'::Read1::GAACAATATAACAAA::Read2::Intron3'::3’ΔHygR + 5’ΔHygR::Intron5'::Read1::CCTCCATCTCATGCT::Read2::Intron3'::3’ΔHygR + 5’ΔHygR::Intron5'::Read1::CTCCGATTTAAACTT::Read2::Intron3'::3’ΔHygR + 5’ΔHygR::Intron5'::Read1::GCACTCTTTAAACGC::Read2::Intron3'::3’ΔHygR + 5’ΔHygR::Intron5'::Read1::TCGCACTCTCATCTA::Read2::Intron3'::3’ΔHygR + 5’ΔHygR::Intron5'::Read1::CTCCAATGTAAATTG::Read2::Intron3'::3’ΔHygR + 5’ΔHygR::Intron5'::Read1::TGACTTTTTTACGTC::Read2::Intron3'::3’ΔHygR + 5’ΔHygR::Intron5'::Read1::TTTCCATGTTAACTT::Read2::Intron3'::3’ΔHygR + 5’ΔHygR::Intron5'::Read1::CCGCGGTTTCATCAT::Read2::Intron3'::3’ΔHygR + 5’ΔHygR::Intron5'::Read1::GCACGTTGTCATCTT::Read2::Intron3'::3’ΔHygR + 5’ΔHygR::Intron5'::Read1::CATCAATATTATTTT::Read2::Intron3'::3’ΔHygR + 5’ΔHygR::Intron5'::Read1::GCCCTCTCTAACTTC::Read2::Intron3'::3’ΔHygR + 5’ΔHygR::Intron5'::Read1::GCTCAATTTTAGATG::Read2::Intron3'::3’ΔHygR + 5’ΔHygR::Intron5'::Read1::GTCCATTTTTAATCT::Read2::Intron3'::3’ΔHygR + 5’ΔHygR::Intron5'::Read1::ACGCTGTCTAAGCCT::Read2::Intron3'::3’ΔHygR + 5’ΔHygR::Intron5'::Read1::TGACTTTATAATGCC::Read2::Intron3'::3’ΔHygR + 5’ΔHygR::Intron5'::Read1::TGACCATTTTACGAC::Read2::Intron3'::3’ΔHygR + 5’ΔHygR::Intron5'::Read1::GCACTTTATCACGTA::Read2::Intron3'::3’ΔHygR + 5’ΔHygR::Intron5'::Read1::ATACGGTCTGAAAAT::Read2::Intron3'::3’ΔHygR + 5’ΔHygR::Intron5'::Read1::GATCCGTTTGATCGC::Read2::Intron3'::3’ΔHygR + 5’ΔHygR::Intron5'::Read1::CACCGTTGTCAAACT::Read2::Intron3'::3’ΔHygR + 5’ΔHygR::Intron5'::Read1::CCTCTATTTCATCCT::Read2::Intron3'::3’ΔHygR + 5’ΔHygR::Intron5'::Read1::CATCACTTTCATTCG::Read2::Intron3'::3’ΔHygR + 5’ΔHygR::Intron5'::Read1::CCGCCTTTTTACCGT::Read2::Intron3'::3’ΔHygR + 5’ΔHygR::Intron5'::Read1::CGCCAATTTAACTAC::Read2::Intron3'::3’ΔHygR + 5’ΔHygR::Intron5'::Read1::AACGTTACAAAGCGA::Read2::Intron3'::3’ΔHygR + 5’ΔHygR::Intron5'::Read1::CCGCCTTTTTAACCA::Read2::Intron3'::3’ΔHygR + 5’ΔHygR::Intron5'::Read1::TCTCGCTGTTAACCA::Read2::Intron3'::3’ΔHygR + 5’ΔHygR::Intron5'::Read1::AATAGTGTCACATGA::Read2::Intron3'::3’ΔHygR + 5’ΔHygR::Intron5'::Read1::TATCAATATTATTTT::Read2::Intron3'::3’ΔHygR + 5’ΔHygR::Intron5'::Read1::CATCTTGTTCCCATC::Read2::Intron3'::3’ΔHygR + 5’ΔHygR::Intron5'::Read1::TACCGATGTTACATC::Read2::Intron3'::3’ΔHygR + 5’ΔHygR::Intron5'::Read1::AACTCTATGGCACAG::Read2::Intron3'::3’ΔHygR + 5’ΔHygR::Intron5'::Read1::GGTCGCTTTGATATA::Read2::Intron3'::3’ΔHygR + 5’ΔHygR::Intron5'::Read1::TAAGTTATATCGTAC::Read2::Intron3'::3’ΔHygR + 5’ΔHygR::Intron5'::Read1::ATTATGAGAAGGAGT::Read2::Intron3'::3’ΔHygR + 5’ΔHygR::Intron5'::Read1::GGTGTCATAGTGGCA::Read2::Intron3'::3’ΔHygR + 5’ΔHygR::Intron5'::Read1::TACCTTCTGTCTGTT::Read2::Intron3'::3’ΔHygR + 5’ΔHygR::Intron5'::Read1::TTTCATTCTCATATC::Read2::Intron3'::3’ΔHygR + 5’ΔHygR::Intron5'::Read1::TAACGGTATGAATAA::Read2::Intron3'::3’ΔHygR + 5’ΔHygR::Intron5'::Read1::TGATTAATAAAGAGA::Read2::Intron3'::3’ΔHygR + 5’ΔHygR::Intron5'::Read1::ATACATTTTCAGCGC::Read2::Intron3'::3’ΔHygR +

5’ΔHygR::Intron5'::Read1::CACCTATTTCACAAC::Read2::Intron3'::3’ΔHygR + 5’ΔHygR::Intron5'::Read1::GCACGCTCTGATATT::Read2::Intron3'::3’ΔHygR + 5’ΔHygR::Intron5'::Read1::ATCCTATGTAAATTC::Read2::Intron3'::3’ΔHygR + 5’ΔHygR::Intron5'::Read1::CGACAGTATGACGCC::Read2::Intron3'::3’ΔHygR + 5’ΔHygR::Intron5'::Read1::CTGCAGTGTCAACCC::Read2::Intron3'::3’ΔHygR + 5’ΔHygR::Intron5'::Read1::GTAATGATAGCGATA::Read2::Intron3'::3’ΔHygR + 5’ΔHygR::Intron5'::Read1::TATTTGATAGGGGGA::Read2::Intron3'::3’ΔHygR + 5’ΔHygR::Intron5'::Read1::GAGCCTTGTAATCTA::Read2::Intron3'::3’ΔHygR + 5’ΔHygR::Intron5'::Read1::GTTCTATATTACGCT::Read2::Intron3'::3’ΔHygR + 5’ΔHygR::Intron5'::Read1::TCGCTTTGTAACGTT::Read2::Intron3'::3’ΔHygR + 5’ΔHygR::Intron5'::Read1::TTAATAACATCGGAC::Read2::Intron3'::3’ΔHygR + 5’ΔHygR::Intron5'::Read1::CGCACCTTGTTAGAG::Read2::Intron3'::3’ΔHygR + 5’ΔHygR::Intron5'::Read1::CTGCAGTGTTAGGCC::Read2::Intron3'::3’ΔHygR + 5’ΔHygR::Intron5'::Read1::GGTTTAATACTGTTA::Read2::Intron3'::3’ΔHygR + 5’ΔHygR::Intron5'::Read1::CACCCCTGTGACTAC::Read2::Intron3'::3’ΔHygR + 5’ΔHygR::Intron5'::Read1::AGACGGTTTTAACTC::Read2::Intron3'::3’ΔHygR + 5’ΔHygR::Intron5'::Read1::ATAGTAAGATAGTGT::Read2::Intron3'::3’ΔHygR + 5’ΔHygR::Intron5'::Read1::ATTCTGAAAGAGTAT::Read2::Intron3'::3’ΔHygR + 5’ΔHygR::Intron5'::Read1::CAACTATATCATTCC::Read2::Intron3'::3’ΔHygR + 5’ΔHygR::Intron5'::Read1::CATAGTTTTCCTTAT::Read2::Intron3'::3’ΔHygR + 5’ΔHygR::Intron5'::Read1::GAGCAATTTAATCAT::Read2::Intron3'::3’ΔHygR + 5’ΔHygR::Intron5'::Read1::GGTATTATAGGGCTT::Read2::Intron3'::3’ΔHygR + 5’ΔHygR::Intron5'::Read1::GGTATTATATGGTTA::Read2::Intron3'::3’ΔHygR + 5’ΔHygR::Intron5'::Read1::TATCCATTTTCATCA::Read2::Intron3'::3’ΔHygR + 5’ΔHygR::Intron5'::Read1::TTTGCATTTTCACTT::Read2::Intron3'::3’ΔHygR + 5’ΔHygR::Intron5'::Read1::AACCTTTATCATGGA::Read2::Intron3'::3’ΔHygR + 5’ΔHygR::Intron5'::Read1::ATTCCTTATAAACCT::Read2::Intron3'::3’ΔHygR + 5’ΔHygR::Intron5'::Read1::CCAGTCATAGTGAGT::Read2::Intron3'::3’ΔHygR + 5’ΔHygR::Intron5'::Read1::GTTGTCAAACTGTAG::Read2::Intron3'::3’ΔHygR + 5’ΔHygR::Intron5'::Read1::TAACTATTTCATTCA::Read2::Intron3'::3’ΔHygR + 5’ΔHygR::Intron5'::Read1::AAACTGATATGGAAC::Read2::Intron3'::3’ΔHygR + 5’ΔHygR::Intron5'::Read1::ACGGTGATAAGGGTA::Read2::Intron3'::3’ΔHygR + 5’ΔHygR::Intron5'::Read1::AGTATTACATCGATG::Read2::Intron3'::3’ΔHygR + 5’ΔHygR::Intron5'::Read1::CTGCGTTTTCATCAA::Read2::Intron3'::3’ΔHygR + 5’ΔHygR::Intron5'::Read1::GAACTTATAGAGCGT::Read2::Intron3'::3’ΔHygR + 5’ΔHygR::Intron5'::Read1::GATAATATACCTAGT::Read2::Intron3'::3’ΔHygR + 5’ΔHygR::Intron5'::Read1::GCAATTAGATTGAGT::Read2::Intron3'::3’ΔHygR + 5’ΔHygR::Intron5'::Read1::GCGCTGTGTGATCAA::Read2::Intron3'::3’ΔHygR + 5’ΔHygR::Intron5'::Read1::GTCGTGCAAGTTTGA::Read2::Intron3'::3’ΔHygR + 5’ΔHygR::Intron5'::Read1::TACCTATATTAGGCG::Read2::Intron3'::3’ΔHygR + 5’ΔHygR::Intron5'::Read1::TTATTGAGACAGGGA::Read2::Intron3'::3’ΔHygR + 5’ΔHygR::Intron5'::Read1::AATATGAGAATGGGT::Read2::Intron3'::3’ΔHygR + 5’ΔHygR::Intron5'::Read1::ACATTGACACTGAAG::Read2::Intron3'::3’ΔHygR + 5’ΔHygR::Intron5'::Read1::AGGTTGAGATAGCAT::Read2::Intron3'::3’ΔHygR + 5’ΔHygR::Intron5'::Read1::ATACATTCTTACACT::Read2::Intron3'::3’ΔHygR + 5’ΔHygR::Intron5'::Read1::ATGTTAATATTGAAC::Read2::Intron3'::3’ΔHygR + 5’ΔHygR::Intron5'::Read1::ATTTTCAGATTGATT::Read2::Intron3'::3’ΔHygR + 5’ΔHygR::Intron5'::Read1::CACCGCTTTTAACTT::Read2::Intron3'::3’ΔHygR + 5’ΔHygR::Intron5'::Read1::GAAATGATATTGAGA::Read2::Intron3'::3’ΔHygR + 5’ΔHygR::Intron5'::Read1::GACACATTGTTATTC::Read2::Intron3'::3’ΔHygR + 5’ΔHygR::Intron5'::Read1::GGAGTAAAAGGGGAA::Read2::Intron3'::3’ΔHygR + 5’ΔHygR::Intron5'::Read1::GTTGTGACACAGTAC::Read2::Intron3'::3’ΔHygR + 5’ΔHygR::Intron5'::Read1::TAAGTTAAATAGGCC::Read2::Intron3'::3’ΔHygR + 5’ΔHygR::Intron5'::Read1::TAGCTAAAAACGATT::Read2::Intron3'::3’ΔHygR + 5’ΔHygR::Intron5'::Read1::TATATGAGAAAGGGT::Read2::Intron3'::3’ΔHygR + 5’ΔHygR::Intron5'::Read1::TCACCCTAGTTAAGC::Read2::Intron3'::3’ΔHygR + 5’ΔHygR::Intron5'::Read1::TTGATTATAAAGGAA::Read2::Intron3'::3’ΔHygR + 5’ΔHygR::Intron5'::Read1::TTTATGATAATGTTC::Read2::Intron3'::3’ΔHygR + 5’ΔHygR::Intron5'::Read1::ACACTCTATCACCCC::Read2::Intron3'::3’ΔHygR + 5’ΔHygR::Intron5'::Read1::AGGATAAGAAGGACC::Read2::Intron3'::3’ΔHygR + 5’ΔHygR::Intron5'::Read1::AGTATCATAAGGACG::Read2::Intron3'::3’ΔHygR + 5’ΔHygR::Intron5'::Read1::ATACTAAGAATGCAA::Read2::Intron3'::3’ΔHygR + 5’ΔHygR::Intron5'::Read1::ATCCTTGTCACCTTA::Read2::Intron3'::3’ΔHygR + 5’ΔHygR::Intron5'::Read1::ATGATCAAATGGTGT::Read2::Intron3'::3’ΔHygR + 5’ΔHygR::Intron5'::Read1::ATGTTAAGAGCGAAA::Read2::Intron3'::3’ΔHygR + 5’ΔHygR::Intron5'::Read1::ATGTTGATAGGGGGA::Read2::Intron3'::3’ΔHygR + 5’ΔHygR::Intron5'::Read1::CCAGCCGTATCATTA::Read2::Intron3'::3’ΔHygR + 5’ΔHygR::Intron5'::Read1::CCGCTGTCTTAAATG::Read2::Intron3'::3’ΔHygR + 5’ΔHygR::Intron5'::Read1::CGCCATTCTTAATTT::Read2::Intron3'::3’ΔHygR + 5’ΔHygR::Intron5'::Read1::CTTATCAAAGTGTGA::Read2::Intron3'::3’ΔHygR +

5’ΔHygR::Intron5'::Read1::GAAATAACAAGGTGA::Read2::Intron3'::3’ΔHygR + 5’ΔHygR::Intron5'::Read1::GAACTAAGAGTGTCG::Read2::Intron3'::3’ΔHygR + 5’ΔHygR::Intron5'::Read1::GATATGAAAACGTTG::Read2::Intron3'::3’ΔHygR + 5’ΔHygR::Intron5'::Read1::GGAATAAGACAGTTT::Read2::Intron3'::3’ΔHygR + 5’ΔHygR::Intron5'::Read1::GGAATCATAACGGTA::Read2::Intron3'::3’ΔHygR + 5’ΔHygR::Intron5'::Read1::GGGATCATAGGGGCG::Read2::Intron3'::3’ΔHygR + 5’ΔHygR::Intron5'::Read1::GTCCTGTTTCATCCC::Read2::Intron3'::3’ΔHygR + 5’ΔHygR::Intron5'::Read1::GTGATTATACCGAGA::Read2::Intron3'::3’ΔHygR + 5’ΔHygR::Intron5'::Read1::TAATTGACATCGTTT::Read2::Intron3'::3’ΔHygR + 5’ΔHygR::Intron5'::Read1::TACATAAAATCGGTA::Read2::Intron3'::3’ΔHygR + 5’ΔHygR::Intron5'::Read1::TACCTAAAAGGGATC::Read2::Intron3'::3’ΔHygR + 5’ΔHygR::Intron5'::Read1::TAGACATTTTCCTGT::Read2::Intron3'::3’ΔHygR + 5’ΔHygR::Intron5'::Read1::TAGTTGAAATGGGGA::Read2::Intron3'::3’ΔHygR + 5’ΔHygR::Intron5'::Read1::TTATTTAGACAGAAT::Read2::Intron3'::3’ΔHygR + 5’ΔHygR::Intron5'::Read1::AAATTGAGACCGATA::Read2::Intron3'::3’ΔHygR + 5’ΔHygR::Intron5'::Read1::AATATGACAGCGAAG::Read2::Intron3'::3’ΔHygR + 5’ΔHygR::Intron5'::Read1::AATGTTATAGTGTGC::Read2::Intron3'::3’ΔHygR + 5’ΔHygR::Intron5'::Read1::AATTTGAAATAGACT::Read2::Intron3'::3’ΔHygR + 5’ΔHygR::Intron5'::Read1::ACTTTAAGATCGGTA::Read2::Intron3'::3’ΔHygR + 5’ΔHygR::Intron5'::Read1::AGTATGACAATGAGC::Read2::Intron3'::3’ΔHygR + 5’ΔHygR::Intron5'::Read1::AGTGTGAGAAAGCCA::Read2::Intron3'::3’ΔHygR + 5’ΔHygR::Intron5'::Read1::AGTGTTAGACAGGTT::Read2::Intron3'::3’ΔHygR + 5’ΔHygR::Intron5'::Read1::ATCGTAAGAAAGGTT::Read2::Intron3'::3’ΔHygR + 5’ΔHygR::Intron5'::Read1::CACCAATTTCACCGC::Read2::Intron3'::3’ΔHygR + 5’ΔHygR::Intron5'::Read1::CATATTAGATGGAAA::Read2::Intron3'::3’ΔHygR + 5’ΔHygR::Intron5'::Read1::CCACCATTTTATTTG::Read2::Intron3'::3’ΔHygR + 5’ΔHygR::Intron5'::Read1::CCCCGTTGTTAATAT::Read2::Intron3'::3’ΔHygR + 5’ΔHygR::Intron5'::Read1::CCTCTTTGTAAATCA::Read2::Intron3'::3’ΔHygR + 5’ΔHygR::Intron5'::Read1::CTATTGATACTGAGC::Read2::Intron3'::3’ΔHygR + 5’ΔHygR::Intron5'::Read1::CTCTTAAAAAGGAAC::Read2::Intron3'::3’ΔHygR + 5’ΔHygR::Intron5'::Read1::GAATTGAGATGGCTG::Read2::Intron3'::3’ΔHygR + 5’ΔHygR::Intron5'::Read1::GAGATAATAAGGCAA::Read2::Intron3'::3’ΔHygR + 5’ΔHygR::Intron5'::Read1::GAGCTAATACGGTTA::Read2::Intron3'::3’ΔHygR + 5’ΔHygR::Intron5'::Read1::GCCTTTATAAGGAGA::Read2::Intron3'::3’ΔHygR + 5’ΔHygR::Intron5'::Read1::GGCGTGAGACGGGCG::Read2::Intron3'::3’ΔHygR + 5’ΔHygR::Intron5'::Read1::GGGTTTAAACGGGTA::Read2::Intron3'::3’ΔHygR + 5’ΔHygR::Intron5'::Read1::GTAGTAAAAAGGGTG::Read2::Intron3'::3’ΔHygR + 5’ΔHygR::Intron5'::Read1::GTAGTAAAATGGGTC::Read2::Intron3'::3’ΔHygR + 5’ΔHygR::Intron5'::Read1::GTATGTGTTATTTTT::Read2::Intron3'::3’ΔHygR + 5’ΔHygR::Intron5'::Read1::GTGTTAACAGTGATC::Read2::Intron3'::3’ΔHygR + 5’ΔHygR::Intron5'::Read1::TAACTTTGTAATTCT::Read2::Intron3'::3’ΔHygR + 5’ΔHygR::Intron5'::Read1::TACTTAAAAGGGAAA::Read2::Intron3'::3’ΔHygR + 5’ΔHygR::Intron5'::Read1::TATGTTAAATGGTCG::Read2::Intron3'::3’ΔHygR + 5’ΔHygR::Intron5'::Read1::TGAATTAGAAAGGGT::Read2::Intron3'::3’ΔHygR + 5’ΔHygR::Intron5'::Read1::TGGATCAGAACGAGG::Read2::Intron3'::3’ΔHygR + 5’ΔHygR::Intron5'::Read1::TGTATTATAGGGTAA::Read2::Intron3'::3’ΔHygR + 5’ΔHygR::Intron5'::Read1::TGTTTTACAGAGTTT::Read2::Intron3'::3’ΔHygR + 5’ΔHygR::Intron5'::Read1::TTCGTTATAGTGTAC::Read2::Intron3'::3’ΔHygR + 5’ΔHygR::Intron5'::Read1::TTCTTGAGACAGTTG::Read2::Intron3'::3’ΔHygR + 5’ΔHygR::Intron5'::Read1::GGCCTATCTGACAGG::Read2::Intron3'::3’ΔHygR + 5’ΔHygR::Intron5'::Read1::TGCCCTTCTGACAGG::Read2::Intron3'::3’ΔHygR + 5’ΔHygR::Intron5'::Read1::CCCCTATCTTATTAA::Read2::Intron3'::3’ΔHygR + 5’ΔHygR::Intron5'::Read1::AATCGATATCACCGG::Read2::Intron3'::3’ΔHygR + 5’ΔHygR::Intron5'::Read1::GGGCGTTCTCACAGG::Read2::Intron3'::3’ΔHygR + 5’ΔHygR::Intron5'::Read1::ACACCATTTAACAGG::Read2::Intron3'::3’ΔHygR + 5’ΔHygR::Intron5'::Read1::TTGCCCTATAACCGG::Read2::Intron3'::3’ΔHygR + 5’ΔHygR::Intron5'::Read1::TACCAATCTCATCGG::Read2::Intron3'::3’ΔHygR + 5’ΔHygR::Intron5'::Read1::CGACCATTTCATCGG::Read2::Intron3'::3’ΔHygR + 5’ΔHygR::Intron5'::Read1::GCCCTATATGACCGG::Read2::Intron3'::3’ΔHygR + 5’ΔHygR::Intron5'::Read1::GTTCACTATAAGTCT::Read2::Intron3'::3’ΔHygR + 5’ΔHygR::Intron5'::Read1::ACGCCGATATTACGC::Read2::Intron3'::3’ΔHygR + 5’ΔHygR::Intron5'::Read1::GATATGATAATGGAG::Read2::Intron3'::3’ΔHygR + 5’ΔHygR::Intron5'::Read1::TCTATCATAAGGTGT::Read2::Intron3'::3’ΔHygR + 5’ΔHygR::Intron5'::Read1::TTCATTACAGAGTTA::Read2::Intron3'::3’ΔHygR + 5’ΔHygR::Intron5'::Read1::CGGCTAATAATGCTA::Read2::Intron3'::3’ΔHygR + 5’ΔHygR::Intron5'::Read1::GACCAGTCTCACTGG::Read2::Intron3'::3’ΔHygR + 5’ΔHygR::Intron5'::Read1::ACGCACTTTCATGGG::Read2::Intron3'::3’ΔHygR + 5’ΔHygR::Intron5'::Read1::TGGCACTATTACACG::Read2::Intron3'::3’ΔHygR + 5’ΔHygR::Intron5'::Read1::CTACGCTTTGACACG::Read2::Intron3'::3’ΔHygR + 5’ΔHygR::Intron5'::Read1::AACCCTTTTCACTGG::Read2::Intron3'::3’ΔHygR +

5’ΔHygR::Intron5'::Read1::GCCCGGTCTAATGGG::Read2::Intron3'::3’ΔHygR + 5’ΔHygR::Intron5'::Read1::CAACCCTCTAAGCGG::Read2::Intron3'::3’ΔHygR + 5’ΔHygR::Intron5'::Read1::CCGCCTTGTAACGGG::Read2::Intron3'::3’ΔHygR + 5’ΔHygR::Intron5'::Read1::TTTCCATCTAATAGG::Read2::Intron3'::3’ΔHygR + 5’ΔHygR::Intron5'::Read1::TTCCAATCTCACACG::Read2::Intron3'::3’ΔHygR + 5’ΔHygR::Intron5'::Read1::AGACGTTCTGATAGG::Read2::Intron3'::3’ΔHygR + 5’ΔHygR::Intron5'::Read1::CGGCCTTGGGAGATC::Read2::Intron3'::3’ΔHygR + 5’ΔHygR::Intron5'::Read1::CCACTGTTTCATAGG::Read2::Intron3'::3’ΔHygR + 5’ΔHygR::Intron5'::Read1::TTACACTATAATGGG::Read2::Intron3'::3’ΔHygR + 5’ΔHygR::Intron5'::Read1::TAACATTTTAATCGG::Read2::Intron3'::3’ΔHygR + 5’ΔHygR::Intron5'::Read1::CCCCTCTCTAAGAGG::Read2::Intron3'::3’ΔHygR + 5’ΔHygR::Intron5'::Read1::GGTCCTTTTAATCGG::Read2::Intron3'::3’ΔHygR + 5’ΔHygR::Intron5'::Read1::TTGCTCTCTTACACG::Read2::Intron3'::3’ΔHygR + 5’ΔHygR::Intron5'::Read1::GACCTCTCTAATATC::Read2::Intron3'::3’ΔHygR + 5’ΔHygR::Intron5'::Read1::GAATTGAAATGGCCG::Read2::Intron3'::3’ΔHygR + 5’ΔHygR::Intron5'::Read1::AATGTTAAAGCGGTC::Read2::Intron3'::3’ΔHygR + 5’ΔHygR::Intron5'::Read1::ACATTGATAGAGGGG::Read2::Intron3'::3’ΔHygR + 5’ΔHygR::Intron5'::Read1::ACGATTATAATGCTC::Read2::Intron3'::3’ΔHygR + 5’ΔHygR::Intron5'::Read1::AGGATAAGAGAGGAG::Read2::Intron3'::3’ΔHygR + 5’ΔHygR::Intron5'::Read1::CCCCTGTTTAAAGAT::Read2::Intron3'::3’ΔHygR + 5’ΔHygR::Intron5'::Read1::GTATTCAGACAGTTA::Read2::Intron3'::3’ΔHygR + 5’ΔHygR::Intron5'::Read1::TTTGTTATATTGTTC::Read2::Intron3'::3’ΔHygR + 5’ΔHygR::Intron5'::Read1::GACCTCTCTTTTCCA::Read2::Intron3'::3’ΔHygR + 5’ΔHygR::Intron5'::Read1::TGAATTACATAGCTA::Read2::Intron3'::3’ΔHygR + 5’ΔHygR::Intron5'::Read1::AACCTAACACTGATT::Read2::Intron3'::3’ΔHygR + 5’ΔHygR::Intron5'::Read1::ACACTATTTCATCTC::Read2::Intron3'::3’ΔHygR + 5’ΔHygR::Intron5'::Read1::CTCCCTTATCATTGC::Read2::Intron3'::3’ΔHygR + 5’ΔHygR::Intron5'::Read1::TATCGGTTTCAGTCA::Read2::Intron3'::3’ΔHygR + 5’ΔHygR::Intron5'::Read1::TTGCTCTCTTATCTG::Read2::Intron3'::3’ΔHygR + 5’ΔHygR::Intron5'::Read1::TACCCCTTTTAAACT::Read2::Intron3'::3’ΔHygR + 5’ΔHygR::Intron5'::Read1::GCTCATTCTGACGTA::Read2::Intron3'::3’ΔHygR + 5’ΔHygR::Intron5'::Read1::ACCCGTTTTAACTCT::Read2::Intron3'::3’ΔHygR + 5’ΔHygR::Intron5'::Read1::TTTCCTTGTCAGATT::Read2::Intron3'::3’ΔHygR + 5’ΔHygR::Intron5'::Read1::AGGCTTTGTCAAACG::Read2::Intron3'::3’ΔHygR + 5’ΔHygR::Intron5'::Read1::ACCCGCTTTTAGTTA::Read2::Intron3'::3’ΔHygR + 5’ΔHygR::Intron5'::Read1::TGCCCTTTTCAGTAC::Read2::Intron3'::3’ΔHygR + 5’ΔHygR::Intron5'::Read1::TACCAATTTGACCAT::Read2::Intron3'::3’ΔHygR + 5’ΔHygR::Intron5'::Read1::TTAAATTATCACATG::Read2::Intron3'::3’ΔHygR + 5’ΔHygR::Intron5'::Read1::GTACCGTTTAACGTC::Read2::Intron3'::3’ΔHygR + 5’ΔHygR::Intron5'::Read1::GCACCATCTGATGCC::Read2::Intron3'::3’ΔHygR +hsp-16.41p::Cas9(dpiRNA)::tbb-2 3’ UTR + prsp-27::NeoR::unc- 54 3' UTR + U6p:: GCGAAGTGACGGTAGACCGT]; fxSi47[ prsp-0p::5’ΔHygR:: GCGAAGTGACGGTAGACCGT ::3’ΔHygR::unc-54 3’::LoxP]

PX817 fxEx26[5’ΔHygR::Intron5'::Read1::CAACCTTCTCAACCA::Read2::Intron3'::3’ΔHygR + 5’ΔHygR::Intron5'::Read1::AACCATTTTCACTAT::Read2::Intron3'::3’ΔHygR + 5’ΔHygR::Intron5'::Read1::CTCCGATGTTACGTT::Read2::Intron3'::3’ΔHygR + 5’ΔHygR::Intron5'::Read1::TCGCGCTTTAATTCT::Read2::Intron3'::3’ΔHygR + 5’ΔHygR::Intron5'::Read1::GAGCTGTGTGACATC::Read2::Intron3'::3’ΔHygR + 5’ΔHygR::Intron5'::Read1::ACCCCCTTTGACCCA::Read2::Intron3'::3’ΔHygR + 5’ΔHygR::Intron5'::Read1::AACCTTTCTGACTTC::Read2::Intron3'::3’ΔHygR + 5’ΔHygR::Intron5'::Read1::ACACCTTCTTAGATC::Read2::Intron3'::3’ΔHygR + 5’ΔHygR::Intron5'::Read1::TAGCTGTCTGATTAC::Read2::Intron3'::3’ΔHygR + 5’ΔHygR::Intron5'::Read1::CATCCTTGTAACTCT::Read2::Intron3'::3’ΔHygR + 5’ΔHygR::Intron5'::Read1::TATCCGTCTCATAAC::Read2::Intron3'::3’ΔHygR + 5’ΔHygR::Intron5'::Read1::CTCCATTCTTAACTT::Read2::Intron3'::3’ΔHygR + 5’ΔHygR::Intron5'::Read1::CTGCACTTTTATTTC::Read2::Intron3'::3’ΔHygR + 5’ΔHygR::Intron5'::Read1::ACACTTTGTAATCTA::Read2::Intron3'::3’ΔHygR + 5’ΔHygR::Intron5'::Read1::TTTCACTATCAATCC::Read2::Intron3'::3’ΔHygR + 5’ΔHygR::Intron5'::Read1::TAGCTCTCTAATTCC::Read2::Intron3'::3’ΔHygR + 5’ΔHygR::Intron5'::Read1::CATCTATTTTAACTC::Read2::Intron3'::3’ΔHygR + 5’ΔHygR::Intron5'::Read1::TTACGGTATAAAGAT::Read2::Intron3'::3’ΔHygR + 5’ΔHygR::Intron5'::Read1::ACCCCTTGTCATACA::Read2::Intron3'::3’ΔHygR + 5’ΔHygR::Intron5'::Read1::CTTCAATATGATAAT::Read2::Intron3'::3’ΔHygR + 5’ΔHygR::Intron5'::Read1::CTCCGTTCTCACGCC::Read2::Intron3'::3’ΔHygR + 5’ΔHygR::Intron5'::Read1::CAACAATCTCAGGTT::Read2::Intron3'::3’ΔHygR + 5’ΔHygR::Intron5'::Read1::GTGCCATGTCAGTTA::Read2::Intron3'::3’ΔHygR + 5’ΔHygR::Intron5'::Read1::TATCGCTATGATAGC::Read2::Intron3'::3’ΔHygR + 5’ΔHygR::Intron5'::Read1::TGCCCCTATAAAGCC::Read2::Intron3'::3’ΔHygR + 5’ΔHygR::Intron5'::Read1::TTACCATGTTATCCA::Read2::Intron3'::3’ΔHygR +

5’ΔHygR::Intron5'::Read1::CGCCACTCTAATCCT::Read2::Intron3'::3’ΔHygR + 5’ΔHygR::Intron5'::Read1::CGTCACTATGATTAA::Read2::Intron3'::3’ΔHygR + 5’ΔHygR::Intron5'::Read1::CGACGATTTCACATC::Read2::Intron3'::3’ΔHygR + 5’ΔHygR::Intron5'::Read1::ACACCCTTTAAAACA::Read2::Intron3'::3’ΔHygR + 5’ΔHygR::Intron5'::Read1::TTTCATTCTTATCCT::Read2::Intron3'::3’ΔHygR + 5’ΔHygR::Intron5'::Read1::GCTCTATTTCATTCA::Read2::Intron3'::3’ΔHygR + 5’ΔHygR::Intron5'::Read1::GTCCCATGTCACGAA::Read2::Intron3'::3’ΔHygR + 5’ΔHygR::Intron5'::Read1::TCTCCGTATTACCTT::Read2::Intron3'::3’ΔHygR + 5’ΔHygR::Intron5'::Read1::ACACAATGTTAGTCG::Read2::Intron3'::3’ΔHygR + 5’ΔHygR::Intron5'::Read1::CCGCTTTTTGACACG::Read2::Intron3'::3’ΔHygR + 5’ΔHygR::Intron5'::Read1::CATCCGTTTGACGTA::Read2::Intron3'::3’ΔHygR + 5’ΔHygR::Intron5'::Read1::TGCCGGTCTTATCTA::Read2::Intron3'::3’ΔHygR + 5’ΔHygR::Intron5'::Read1::CAACGATTTAACAGC::Read2::Intron3'::3’ΔHygR + 5’ΔHygR::Intron5'::Read1::TCTCAATCTTATCTA::Read2::Intron3'::3’ΔHygR + 5’ΔHygR::Intron5'::Read1::TTGCCTTTTTATACA::Read2::Intron3'::3’ΔHygR + 5’ΔHygR::Intron5'::Read1::AGGCTTTATCAATCC::Read2::Intron3'::3’ΔHygR + 5’ΔHygR::Intron5'::Read1::TTACCATTTGAACCT::Read2::Intron3'::3’ΔHygR + 5’ΔHygR::Intron5'::Read1::CTGCCATTTGACCCC::Read2::Intron3'::3’ΔHygR + 5’ΔHygR::Intron5'::Read1::CACCGATGTAATTTA::Read2::Intron3'::3’ΔHygR + 5’ΔHygR::Intron5'::Read1::TAGCACTTTAACCCT::Read2::Intron3'::3’ΔHygR + 5’ΔHygR::Intron5'::Read1::CATCTTTTTGATAGT::Read2::Intron3'::3’ΔHygR + 5’ΔHygR::Intron5'::Read1::AGGCTCTATCAGTCG::Read2::Intron3'::3’ΔHygR + 5’ΔHygR::Intron5'::Read1::CTGCTTTCTCATCAT::Read2::Intron3'::3’ΔHygR + 5’ΔHygR::Intron5'::Read1::TTACGGTCTAATGCA::Read2::Intron3'::3’ΔHygR + 5’ΔHygR::Intron5'::Read1::CCTCTCTTTCAACCA::Read2::Intron3'::3’ΔHygR + 5’ΔHygR::Intron5'::Read1::AACCAGTATAATTCA::Read2::Intron3'::3’ΔHygR + 5’ΔHygR::Intron5'::Read1::GCCCTGTTTAAGGAA::Read2::Intron3'::3’ΔHygR + 5’ΔHygR::Intron5'::Read1::CGTCTTTCTCACCAT::Read2::Intron3'::3’ΔHygR + 5’ΔHygR::Intron5'::Read1::CGCCTGTCTAACTTT::Read2::Intron3'::3’ΔHygR + 5’ΔHygR::Intron5'::Read1::TAGCTATATTATCAG::Read2::Intron3'::3’ΔHygR + 5’ΔHygR::Intron5'::Read1::GAACCTTCTCAACAC::Read2::Intron3'::3’ΔHygR + 5’ΔHygR::Intron5'::Read1::TCACGATATCACCGT::Read2::Intron3'::3’ΔHygR + 5’ΔHygR::Intron5'::Read1::TTTCCCTTTTACCCT::Read2::Intron3'::3’ΔHygR + 5’ΔHygR::Intron5'::Read1::AGACTCTCTGATGAC::Read2::Intron3'::3’ΔHygR + 5’ΔHygR::Intron5'::Read1::ACCCTTTTTTAACTA::Read2::Intron3'::3’ΔHygR + 5’ΔHygR::Intron5'::Read1::GTACGCTTTAACCCG::Read2::Intron3'::3’ΔHygR + 5’ΔHygR::Intron5'::Read1::TGCCATTGTAAGCTT::Read2::Intron3'::3’ΔHygR + 5’ΔHygR::Intron5'::Read1::TACCACTCTAACCTG::Read2::Intron3'::3’ΔHygR + 5’ΔHygR::Intron5'::Read1::TTTCCCTTTGACTTC::Read2::Intron3'::3’ΔHygR + 5’ΔHygR::Intron5'::Read1::CGACACTTTCAATCA::Read2::Intron3'::3’ΔHygR + 5’ΔHygR::Intron5'::Read1::TCCCTCTCTAATCAT::Read2::Intron3'::3’ΔHygR + 5’ΔHygR::Intron5'::Read1::ATCCTGTTTTAGCCT::Read2::Intron3'::3’ΔHygR + 5’ΔHygR::Intron5'::Read1::CTTCTTTTTTATCGT::Read2::Intron3'::3’ΔHygR + 5’ΔHygR::Intron5'::Read1::ATCCCATTTGATCAA::Read2::Intron3'::3’ΔHygR + 5’ΔHygR::Intron5'::Read1::CCTCAATTTAAGACA::Read2::Intron3'::3’ΔHygR + 5’ΔHygR::Intron5'::Read1::GCGCTTTCTAACCCT::Read2::Intron3'::3’ΔHygR + 5’ΔHygR::Intron5'::Read1::CCACCTTCTTAATAC::Read2::Intron3'::3’ΔHygR + 5’ΔHygR::Intron5'::Read1::TACCACTATCACACA::Read2::Intron3'::3’ΔHygR + 5’ΔHygR::Intron5'::Read1::GGGCGATTTTAAAAA::Read2::Intron3'::3’ΔHygR + 5’ΔHygR::Intron5'::Read1::CAACTCTTTAAAACC::Read2::Intron3'::3’ΔHygR + 5’ΔHygR::Intron5'::Read1::TGACAATTTGAGTAC::Read2::Intron3'::3’ΔHygR + 5’ΔHygR::Intron5'::Read1::CACCTATTTCATTGG::Read2::Intron3'::3’ΔHygR + 5’ΔHygR::Intron5'::Read1::GCCCCATCTTAACTA::Read2::Intron3'::3’ΔHygR + 5’ΔHygR::Intron5'::Read1::CTTCCGTCTTAGCTC::Read2::Intron3'::3’ΔHygR + 5’ΔHygR::Intron5'::Read1::ATCCGTTTTGATCTT::Read2::Intron3'::3’ΔHygR + 5’ΔHygR::Intron5'::Read1::TGGCTCTTTCAATAC::Read2::Intron3'::3’ΔHygR + 5’ΔHygR::Intron5'::Read1::CATCACTGTTATGTT::Read2::Intron3'::3’ΔHygR + 5’ΔHygR::Intron5'::Read1::TTACGATGTCAGGTC::Read2::Intron3'::3’ΔHygR + 5’ΔHygR::Intron5'::Read1::GTTCGGTTTCAGACA::Read2::Intron3'::3’ΔHygR + 5’ΔHygR::Intron5'::Read1::CAACTCTTTGAAACC::Read2::Intron3'::3’ΔHygR + 5’ΔHygR::Intron5'::Read1::GGTCCTTTTCATCTC::Read2::Intron3'::3’ΔHygR + 5’ΔHygR::Intron5'::Read1::ACACGTTTTTACATC::Read2::Intron3'::3’ΔHygR + 5’ΔHygR::Intron5'::Read1::TCTCGATTTCAGTTG::Read2::Intron3'::3’ΔHygR + 5’ΔHygR::Intron5'::Read1::AAACGCTCTTACTAA::Read2::Intron3'::3’ΔHygR + 5’ΔHygR::Intron5'::Read1::GCCCTGTTTTATATA::Read2::Intron3'::3’ΔHygR + 5’ΔHygR::Intron5'::Read1::GTTCTATATTAAATC::Read2::Intron3'::3’ΔHygR + 5’ΔHygR::Intron5'::Read1::CTACCTTGTTAGAAC::Read2::Intron3'::3’ΔHygR + 5’ΔHygR::Intron5'::Read1::CGTCCTTTTCATATC::Read2::Intron3'::3’ΔHygR + 5’ΔHygR::Intron5'::Read1::TAGCGGTCTGAATTA::Read2::Intron3'::3’ΔHygR + 5’ΔHygR::Intron5'::Read1::AACCTTTTTAAATTG::Read2::Intron3'::3’ΔHygR +

5’ΔHygR::Intron5'::Read1::TTTCCGTATCATCTT::Read2::Intron3'::3’ΔHygR + 5’ΔHygR::Intron5'::Read1::CTGCTTTCTTACTTC::Read2::Intron3'::3’ΔHygR + 5’ΔHygR::Intron5'::Read1::TGGCCTTCTTATTAG::Read2::Intron3'::3’ΔHygR + 5’ΔHygR::Intron5'::Read1::AAACTTTGTCATGGA::Read2::Intron3'::3’ΔHygR + 5’ΔHygR::Intron5'::Read1::TTCCTTTGTAACCCT::Read2::Intron3'::3’ΔHygR + 5’ΔHygR::Intron5'::Read1::TTCCGATTTGATTGT::Read2::Intron3'::3’ΔHygR + 5’ΔHygR::Intron5'::Read1::GGCCCATGTTAAAGC::Read2::Intron3'::3’ΔHygR + 5’ΔHygR::Intron5'::Read1::TTTCCGTTTCACTCT::Read2::Intron3'::3’ΔHygR + 5’ΔHygR::Intron5'::Read1::ACACCTTTTTAATTA::Read2::Intron3'::3’ΔHygR + 5’ΔHygR::Intron5'::Read1::GCTCTTTCTAATTCA::Read2::Intron3'::3’ΔHygR + 5’ΔHygR::Intron5'::Read1::TAGCGTTCTCAGACT::Read2::Intron3'::3’ΔHygR + 5’ΔHygR::Intron5'::Read1::TTTCAATTTCAAAAG::Read2::Intron3'::3’ΔHygR + 5’ΔHygR::Intron5'::Read1::GAACTATCTTATACA::Read2::Intron3'::3’ΔHygR + 5’ΔHygR::Intron5'::Read1::AGACCATTTCAGCTC::Read2::Intron3'::3’ΔHygR + 5’ΔHygR::Intron5'::Read1::CAACTATGTAACTGA::Read2::Intron3'::3’ΔHygR + 5’ΔHygR::Intron5'::Read1::GCCCCCTGTAACCTC::Read2::Intron3'::3’ΔHygR + 5’ΔHygR::Intron5'::Read1::CAACTTTGTGAATAA::Read2::Intron3'::3’ΔHygR + 5’ΔHygR::Intron5'::Read1::AGCCTATCTTAATAC::Read2::Intron3'::3’ΔHygR + 5’ΔHygR::Intron5'::Read1::ATCCGTTGTCACACG::Read2::Intron3'::3’ΔHygR + 5’ΔHygR::Intron5'::Read1::CCTCAATTTCATACC::Read2::Intron3'::3’ΔHygR + 5’ΔHygR::Intron5'::Read1::TCGCCGTTTTAACCA::Read2::Intron3'::3’ΔHygR + 5’ΔHygR::Intron5'::Read1::CACCGATGTTACATC::Read2::Intron3'::3’ΔHygR + 5’ΔHygR::Intron5'::Read1::ATCCAATCTGAGCTT::Read2::Intron3'::3’ΔHygR + 5’ΔHygR::Intron5'::Read1::CCTCTGTATAACCCC::Read2::Intron3'::3’ΔHygR + 5’ΔHygR::Intron5'::Read1::AGCCTTTCTAAACGT::Read2::Intron3'::3’ΔHygR + 5’ΔHygR::Intron5'::Read1::TGACTATTTCAGTAC::Read2::Intron3'::3’ΔHygR + 5’ΔHygR::Intron5'::Read1::CACCGTTATGAACCC::Read2::Intron3'::3’ΔHygR + 5’ΔHygR::Intron5'::Read1::CCTCTATCTCACATA::Read2::Intron3'::3’ΔHygR + 5’ΔHygR::Intron5'::Read1::CAACACTATGACTTT::Read2::Intron3'::3’ΔHygR + 5’ΔHygR::Intron5'::Read1::AAGCTGTCTCATTGG::Read2::Intron3'::3’ΔHygR + 5’ΔHygR::Intron5'::Read1::TTGCATTTTTAAAAC::Read2::Intron3'::3’ΔHygR + 5’ΔHygR::Intron5'::Read1::TCGCAATTTCAGATA::Read2::Intron3'::3’ΔHygR + 5’ΔHygR::Intron5'::Read1::ATCCTGTCTTAATTT::Read2::Intron3'::3’ΔHygR + 5’ΔHygR::Intron5'::Read1::CGACACTTTCATATG::Read2::Intron3'::3’ΔHygR + 5’ΔHygR::Intron5'::Read1::ACGCTCTCTCAGGAA::Read2::Intron3'::3’ΔHygR + 5’ΔHygR::Intron5'::Read1::AAGCCATTTGAAAAC::Read2::Intron3'::3’ΔHygR + 5’ΔHygR::Intron5'::Read1::CGACCTTTTGAATGT::Read2::Intron3'::3’ΔHygR + 5’ΔHygR::Intron5'::Read1::TCGCGATCTTACATC::Read2::Intron3'::3’ΔHygR + 5’ΔHygR::Intron5'::Read1::GGCCTATTTCACTTA::Read2::Intron3'::3’ΔHygR + 5’ΔHygR::Intron5'::Read1::TACCGTTGTAACATT::Read2::Intron3'::3’ΔHygR + 5’ΔHygR::Intron5'::Read1::CAACTTTTTCACTCC::Read2::Intron3'::3’ΔHygR + 5’ΔHygR::Intron5'::Read1::TTCCACTCTTAGTTG::Read2::Intron3'::3’ΔHygR + 5’ΔHygR::Intron5'::Read1::CTGCATTTTGACTCC::Read2::Intron3'::3’ΔHygR + 5’ΔHygR::Intron5'::Read1::AGACCGTATAAACAT::Read2::Intron3'::3’ΔHygR + 5’ΔHygR::Intron5'::Read1::TGGCGATATGACTCA::Read2::Intron3'::3’ΔHygR + 5’ΔHygR::Intron5'::Read1::ATACCATATTACTCA::Read2::Intron3'::3’ΔHygR + 5’ΔHygR::Intron5'::Read1::TTACCGTGTGAGACA::Read2::Intron3'::3’ΔHygR + 5’ΔHygR::Intron5'::Read1::GTCCTATTTGATGCA::Read2::Intron3'::3’ΔHygR + 5’ΔHygR::Intron5'::Read1::TTTCGTTATGAATCT::Read2::Intron3'::3’ΔHygR + 5’ΔHygR::Intron5'::Read1::GTACAGTATGACGGA::Read2::Intron3'::3’ΔHygR + 5’ΔHygR::Intron5'::Read1::TGCCTTTTTGACTTA::Read2::Intron3'::3’ΔHygR + 5’ΔHygR::Intron5'::Read1::TCGCCATTTTATCAA::Read2::Intron3'::3’ΔHygR + 5’ΔHygR::Intron5'::Read1::CATCTATATGAATCA::Read2::Intron3'::3’ΔHygR + 5’ΔHygR::Intron5'::Read1::CCACTTTCTCACATT::Read2::Intron3'::3’ΔHygR + 5’ΔHygR::Intron5'::Read1::TTACACTATAACTAA::Read2::Intron3'::3’ΔHygR + 5’ΔHygR::Intron5'::Read1::TCACATTTTCACCAA::Read2::Intron3'::3’ΔHygR + 5’ΔHygR::Intron5'::Read1::GCACCGTCTGATTTG::Read2::Intron3'::3’ΔHygR + 5’ΔHygR::Intron5'::Read1::TGTCGATGTCATTAA::Read2::Intron3'::3’ΔHygR + 5’ΔHygR::Intron5'::Read1::AATCGCTTTTATATT::Read2::Intron3'::3’ΔHygR + 5’ΔHygR::Intron5'::Read1::CCACTATGTCATCCA::Read2::Intron3'::3’ΔHygR + 5’ΔHygR::Intron5'::Read1::CCTCTTTGTCATTCA::Read2::Intron3'::3’ΔHygR + 5’ΔHygR::Intron5'::Read1::GTACTTTTTAATGTT::Read2::Intron3'::3’ΔHygR + 5’ΔHygR::Intron5'::Read1::AAGCTCTATGATACT::Read2::Intron3'::3’ΔHygR + 5’ΔHygR::Intron5'::Read1::TAGCAGTTTAAATTC::Read2::Intron3'::3’ΔHygR + 5’ΔHygR::Intron5'::Read1::GATCCTTATTATAGC::Read2::Intron3'::3’ΔHygR + 5’ΔHygR::Intron5'::Read1::ACTCCCTTTAAGCTG::Read2::Intron3'::3’ΔHygR + 5’ΔHygR::Intron5'::Read1::CCTCCCTCTTAAACA::Read2::Intron3'::3’ΔHygR + 5’ΔHygR::Intron5'::Read1::CGTCTGTTTTAGTAC::Read2::Intron3'::3’ΔHygR + 5’ΔHygR::Intron5'::Read1::TTACCGTTTAAACTT::Read2::Intron3'::3’ΔHygR + 5’ΔHygR::Intron5'::Read1::CTCCTCTATTAGCTC::Read2::Intron3'::3’ΔHygR +

5’ΔHygR::Intron5'::Read1::GACCTCTTTTATACC::Read2::Intron3'::3’ΔHygR + 5’ΔHygR::Intron5'::Read1::ACTCGCTTTAAGGAC::Read2::Intron3'::3’ΔHygR + 5’ΔHygR::Intron5'::Read1::ACGCGTTTTAACTGA::Read2::Intron3'::3’ΔHygR + 5’ΔHygR::Intron5'::Read1::GCCCACTATTACTAA::Read2::Intron3'::3’ΔHygR + 5’ΔHygR::Intron5'::Read1::TAACATTATTAGCTT::Read2::Intron3'::3’ΔHygR + 5’ΔHygR::Intron5'::Read1::AAACTGTCTAAAACC::Read2::Intron3'::3’ΔHygR + 5’ΔHygR::Intron5'::Read1::CGCCTGTTTGAAGAT::Read2::Intron3'::3’ΔHygR + 5’ΔHygR::Intron5'::Read1::TCTCCGTCTTACTTC::Read2::Intron3'::3’ΔHygR + 5’ΔHygR::Intron5'::Read1::TTGCAGTATCACATC::Read2::Intron3'::3’ΔHygR + 5’ΔHygR::Intron5'::Read1::TAGCTATCTCATGAA::Read2::Intron3'::3’ΔHygR + 5’ΔHygR::Intron5'::Read1::CTGCTGTATCAATGA::Read2::Intron3'::3’ΔHygR + 5’ΔHygR::Intron5'::Read1::TGCCAATTTTATCGC::Read2::Intron3'::3’ΔHygR + 5’ΔHygR::Intron5'::Read1::TGACTGTATCATACT::Read2::Intron3'::3’ΔHygR + 5’ΔHygR::Intron5'::Read1::AAGCCCTTTCATTAC::Read2::Intron3'::3’ΔHygR + 5’ΔHygR::Intron5'::Read1::CGCCGCTATTATTTC::Read2::Intron3'::3’ΔHygR + 5’ΔHygR::Intron5'::Read1::GTTCCATTTCACACT::Read2::Intron3'::3’ΔHygR + 5’ΔHygR::Intron5'::Read1::ACACAATCTCACATA::Read2::Intron3'::3’ΔHygR + 5’ΔHygR::Intron5'::Read1::CGCCAATATTATGCT::Read2::Intron3'::3’ΔHygR + 5’ΔHygR::Intron5'::Read1::CCCCGTTATTAATCT::Read2::Intron3'::3’ΔHygR + 5’ΔHygR::Intron5'::Read1::GCACGTTTTCAAGCA::Read2::Intron3'::3’ΔHygR + 5’ΔHygR::Intron5'::Read1::ACCCTCTGTGACTAC::Read2::Intron3'::3’ΔHygR + 5’ΔHygR::Intron5'::Read1::CTGCTCTCTAACCCA::Read2::Intron3'::3’ΔHygR + 5’ΔHygR::Intron5'::Read1::GATCAGTCTTAGTGA::Read2::Intron3'::3’ΔHygR + 5’ΔHygR::Intron5'::Read1::CACCGTTTTCACCCC::Read2::Intron3'::3’ΔHygR + 5’ΔHygR::Intron5'::Read1::CACCTCTATTATTCC::Read2::Intron3'::3’ΔHygR + 5’ΔHygR::Intron5'::Read1::CTCCGATCTTAAACT::Read2::Intron3'::3’ΔHygR + 5’ΔHygR::Intron5'::Read1::GTACTTTCTCATCAC::Read2::Intron3'::3’ΔHygR + 5’ΔHygR::Intron5'::Read1::TCACCTTCTAACTGA::Read2::Intron3'::3’ΔHygR + 5’ΔHygR::Intron5'::Read1::TACCTTTTTGAACGC::Read2::Intron3'::3’ΔHygR + 5’ΔHygR::Intron5'::Read1::CGACTTTGTTAATGC::Read2::Intron3'::3’ΔHygR + 5’ΔHygR::Intron5'::Read1::CTGCCTTTTCATGTC::Read2::Intron3'::3’ΔHygR + 5’ΔHygR::Intron5'::Read1::TGGCCTTTTGATAAA::Read2::Intron3'::3’ΔHygR + 5’ΔHygR::Intron5'::Read1::GATCCGTTTTACGAT::Read2::Intron3'::3’ΔHygR + 5’ΔHygR::Intron5'::Read1::ATTCTCTTTTATCGC::Read2::Intron3'::3’ΔHygR + 5’ΔHygR::Intron5'::Read1::ACCCATTTTAATATC::Read2::Intron3'::3’ΔHygR + 5’ΔHygR::Intron5'::Read1::AGTCTCTGTTATTAT::Read2::Intron3'::3’ΔHygR + 5’ΔHygR::Intron5'::Read1::ACCCTCTGTAATTCC::Read2::Intron3'::3’ΔHygR + 5’ΔHygR::Intron5'::Read1::CTACGCTTTGAATCA::Read2::Intron3'::3’ΔHygR + 5’ΔHygR::Intron5'::Read1::TGCCATTATCACCGA::Read2::Intron3'::3’ΔHygR + 5’ΔHygR::Intron5'::Read1::AACCGTTCTCAGAGC::Read2::Intron3'::3’ΔHygR + 5’ΔHygR::Intron5'::Read1::GTTCGTTTTCACCAT::Read2::Intron3'::3’ΔHygR + 5’ΔHygR::Intron5'::Read1::CTCCTATTTCACATC::Read2::Intron3'::3’ΔHygR + 5’ΔHygR::Intron5'::Read1::CTGCATTCTCACCCA::Read2::Intron3'::3’ΔHygR + 5’ΔHygR::Intron5'::Read1::ATTCATTGTCACTAA::Read2::Intron3'::3’ΔHygR + 5’ΔHygR::Intron5'::Read1::CGTCGATGTGACTCA::Read2::Intron3'::3’ΔHygR + 5’ΔHygR::Intron5'::Read1::GAACATTGTAACCAT::Read2::Intron3'::3’ΔHygR + 5’ΔHygR::Intron5'::Read1::AACCCTTATCAGCGT::Read2::Intron3'::3’ΔHygR + 5’ΔHygR::Intron5'::Read1::TCCCGTTTTCATCCA::Read2::Intron3'::3’ΔHygR + 5’ΔHygR::Intron5'::Read1::GGCCTTTTTTACACT::Read2::Intron3'::3’ΔHygR + 5’ΔHygR::Intron5'::Read1::ATCCATTCTAACACC::Read2::Intron3'::3’ΔHygR + 5’ΔHygR::Intron5'::Read1::AAACATTATGAGCAA::Read2::Intron3'::3’ΔHygR + 5’ΔHygR::Intron5'::Read1::GGCCCTTATTACGTC::Read2::Intron3'::3’ΔHygR + 5’ΔHygR::Intron5'::Read1::TGGCATTCTAACGTA::Read2::Intron3'::3’ΔHygR + 5’ΔHygR::Intron5'::Read1::TACCTTTCTTAACAC::Read2::Intron3'::3’ΔHygR + 5’ΔHygR::Intron5'::Read1::TCCCATTTTTAAAAG::Read2::Intron3'::3’ΔHygR + 5’ΔHygR::Intron5'::Read1::TGATTTAGAACGGTA::Read2::Intron3'::3’ΔHygR + 5’ΔHygR::Intron5'::Read1::CCCCCCTATGACAAG::Read2::Intron3'::3’ΔHygR + 5’ΔHygR::Intron5'::Read1::AAGCTATTTCACGCC::Read2::Intron3'::3’ΔHygR + 5’ΔHygR::Intron5'::Read1::TTGCTTTATCATTCA::Read2::Intron3'::3’ΔHygR + 5’ΔHygR::Intron5'::Read1::TCCCTATCTGAAGCG::Read2::Intron3'::3’ΔHygR + 5’ΔHygR::Intron5'::Read1::TTGCAATATCAGTTG::Read2::Intron3'::3’ΔHygR + 5’ΔHygR::Intron5'::Read1::CCTCGGTTTCACTTA::Read2::Intron3'::3’ΔHygR + 5’ΔHygR::Intron5'::Read1::CGTCATTTTAACACC::Read2::Intron3'::3’ΔHygR + 5’ΔHygR::Intron5'::Read1::ATACATTCTAAACTC::Read2::Intron3'::3’ΔHygR + 5’ΔHygR::Intron5'::Read1::CATCCATCTTATACA::Read2::Intron3'::3’ΔHygR + 5’ΔHygR::Intron5'::Read1::TTGCATTCTAAGCTC::Read2::Intron3'::3’ΔHygR + 5’ΔHygR::Intron5'::Read1::CATCCTTTTCAATAG::Read2::Intron3'::3’ΔHygR + 5’ΔHygR::Intron5'::Read1::GGTCCATCTTAATAC::Read2::Intron3'::3’ΔHygR + 5’ΔHygR::Intron5'::Read1::GCGCGCTGTTATTCT::Read2::Intron3'::3’ΔHygR + 5’ΔHygR::Intron5'::Read1::AAACTATATGATTAC::Read2::Intron3'::3’ΔHygR +

5’ΔHygR::Intron5'::Read1::CACCGTTCTCACTAC::Read2::Intron3'::3’ΔHygR + 5’ΔHygR::Intron5'::Read1::TCTCTGTTTCATGCA::Read2::Intron3'::3’ΔHygR + 5’ΔHygR::Intron5'::Read1::CGTCTGTTTTACCGA::Read2::Intron3'::3’ΔHygR + 5’ΔHygR::Intron5'::Read1::ACTCTCTCTGAAACA::Read2::Intron3'::3’ΔHygR + 5’ΔHygR::Intron5'::Read1::CTTCATTGTTACTCA::Read2::Intron3'::3’ΔHygR + 5’ΔHygR::Intron5'::Read1::ACACAATTTAATGAC::Read2::Intron3'::3’ΔHygR + 5’ΔHygR::Intron5'::Read1::CTTCCCTTTGAATGT::Read2::Intron3'::3’ΔHygR + 5’ΔHygR::Intron5'::Read1::ATACGTTTTCATACG::Read2::Intron3'::3’ΔHygR + 5’ΔHygR::Intron5'::Read1::CCCCGGTATTACAAG::Read2::Intron3'::3’ΔHygR + 5’ΔHygR::Intron5'::Read1::GGTCTTTGTCAACGC::Read2::Intron3'::3’ΔHygR + 5’ΔHygR::Intron5'::Read1::GCCCCCTTTTAACGT::Read2::Intron3'::3’ΔHygR + 5’ΔHygR::Intron5'::Read1::TCTCTTTATTATAGC::Read2::Intron3'::3’ΔHygR + 5’ΔHygR::Intron5'::Read1::GCCCTATGTGATCAC::Read2::Intron3'::3’ΔHygR + 5’ΔHygR::Intron5'::Read1::ATTCTATCTCATGCC::Read2::Intron3'::3’ΔHygR + 5’ΔHygR::Intron5'::Read1::GCCCGTTCTCATTGA::Read2::Intron3'::3’ΔHygR + 5’ΔHygR::Intron5'::Read1::CGTCTTTTTAACACT::Read2::Intron3'::3’ΔHygR + 5’ΔHygR::Intron5'::Read1::GATCTCTTTTAAGGT::Read2::Intron3'::3’ΔHygR + 5’ΔHygR::Intron5'::Read1::TCACTCTCTAATATA::Read2::Intron3'::3’ΔHygR + 5’ΔHygR::Intron5'::Read1::GGACTTTGTAAACGG::Read2::Intron3'::3’ΔHygR + 5’ΔHygR::Intron5'::Read1::AGCCATTTTAAGCTT::Read2::Intron3'::3’ΔHygR + 5’ΔHygR::Intron5'::Read1::ATCCTTTGTCACATC::Read2::Intron3'::3’ΔHygR + 5’ΔHygR::Intron5'::Read1::GGTCATTATCACGAG::Read2::Intron3'::3’ΔHygR + 5’ΔHygR::Intron5'::Read1::GCACAATTTAACATC::Read2::Intron3'::3’ΔHygR + 5’ΔHygR::Intron5'::Read1::TTACCATTTAAATCG::Read2::Intron3'::3’ΔHygR + 5’ΔHygR::Intron5'::Read1::TGCCCTTTTCAAGTT::Read2::Intron3'::3’ΔHygR + 5’ΔHygR::Intron5'::Read1::TGACGTTATGAGTGT::Read2::Intron3'::3’ΔHygR + 5’ΔHygR::Intron5'::Read1::TATCGGTCTTACATC::Read2::Intron3'::3’ΔHygR + 5’ΔHygR::Intron5'::Read1::TGTCAGTCTCAAATC::Read2::Intron3'::3’ΔHygR + 5’ΔHygR::Intron5'::Read1::TTGCGTTGTCACAGA::Read2::Intron3'::3’ΔHygR + 5’ΔHygR::Intron5'::Read1::TTTCGGTCTCACAAT::Read2::Intron3'::3’ΔHygR + 5’ΔHygR::Intron5'::Read1::CTTCTGTGTTAATTA::Read2::Intron3'::3’ΔHygR + 5’ΔHygR::Intron5'::Read1::ACGCCGTTTGAGTTC::Read2::Intron3'::3’ΔHygR + 5’ΔHygR::Intron5'::Read1::GACCTATATAAATGC::Read2::Intron3'::3’ΔHygR + 5’ΔHygR::Intron5'::Read1::CCGCTCTATCACACT::Read2::Intron3'::3’ΔHygR + 5’ΔHygR::Intron5'::Read1::GGACCGTCTTAAAAT::Read2::Intron3'::3’ΔHygR + 5’ΔHygR::Intron5'::Read1::TTCCATTATAAGGTG::Read2::Intron3'::3’ΔHygR + 5’ΔHygR::Intron5'::Read1::GTGCTATGTCAGTAT::Read2::Intron3'::3’ΔHygR + 5’ΔHygR::Intron5'::Read1::TGTCGTTCTGACAGC::Read2::Intron3'::3’ΔHygR + 5’ΔHygR::Intron5'::Read1::CAGCGTTCTCAACAC::Read2::Intron3'::3’ΔHygR + 5’ΔHygR::Intron5'::Read1::AACCTTTCTCACACA::Read2::Intron3'::3’ΔHygR + 5’ΔHygR::Intron5'::Read1::CAACTCTCTCACGTC::Read2::Intron3'::3’ΔHygR + 5’ΔHygR::Intron5'::Read1::TTGCATTATGAAGCC::Read2::Intron3'::3’ΔHygR + 5’ΔHygR::Intron5'::Read1::AGCCATTCTTACGAG::Read2::Intron3'::3’ΔHygR + 5’ΔHygR::Intron5'::Read1::TTGCTATATTACCCT::Read2::Intron3'::3’ΔHygR + 5’ΔHygR::Intron5'::Read1::CATCTTTATCATTGG::Read2::Intron3'::3’ΔHygR + 5’ΔHygR::Intron5'::Read1::TACCGTTATCACCAT::Read2::Intron3'::3’ΔHygR + 5’ΔHygR::Intron5'::Read1::CAGCATTGTGACCAT::Read2::Intron3'::3’ΔHygR + 5’ΔHygR::Intron5'::Read1::ACCCGTTGTTACCCC::Read2::Intron3'::3’ΔHygR + 5’ΔHygR::Intron5'::Read1::TCCCTATTTAACGGC::Read2::Intron3'::3’ΔHygR + 5’ΔHygR::Intron5'::Read1::TGTCCATGTCACTTA::Read2::Intron3'::3’ΔHygR + 5’ΔHygR::Intron5'::Read1::ACACCATGTCACCTT::Read2::Intron3'::3’ΔHygR + 5’ΔHygR::Intron5'::Read1::GTTCCTTTTGAGATC::Read2::Intron3'::3’ΔHygR + 5’ΔHygR::Intron5'::Read1::ACCCGATGTTACTGC::Read2::Intron3'::3’ΔHygR + 5’ΔHygR::Intron5'::Read1::GAGCCATGTTACATC::Read2::Intron3'::3’ΔHygR + 5’ΔHygR::Intron5'::Read1::ACGCCCTCTAATGCT::Read2::Intron3'::3’ΔHygR + 5’ΔHygR::Intron5'::Read1::TATCATTCTAAGTTT::Read2::Intron3'::3’ΔHygR + 5’ΔHygR::Intron5'::Read1::TATCTATTTTACTTG::Read2::Intron3'::3’ΔHygR + 5’ΔHygR::Intron5'::Read1::TTGCACTATAAGGCC::Read2::Intron3'::3’ΔHygR + 5’ΔHygR::Intron5'::Read1::GATCTTTGTTACAAT::Read2::Intron3'::3’ΔHygR + 5’ΔHygR::Intron5'::Read1::CGTCCATTTGATCCT::Read2::Intron3'::3’ΔHygR + 5’ΔHygR::Intron5'::Read1::TCGCCTTATCAATGC::Read2::Intron3'::3’ΔHygR + 5’ΔHygR::Intron5'::Read1::TGTCCCTATAATTTT::Read2::Intron3'::3’ΔHygR + 5’ΔHygR::Intron5'::Read1::GATCGGTATTAATCA::Read2::Intron3'::3’ΔHygR + 5’ΔHygR::Intron5'::Read1::ATGCGCTCTGAACTA::Read2::Intron3'::3’ΔHygR + 5’ΔHygR::Intron5'::Read1::CGCCTCTATTACTAG::Read2::Intron3'::3’ΔHygR + 5’ΔHygR::Intron5'::Read1::TTGCAGTCTTATCGT::Read2::Intron3'::3’ΔHygR + 5’ΔHygR::Intron5'::Read1::AGTCGATATAAGTCT::Read2::Intron3'::3’ΔHygR + 5’ΔHygR::Intron5'::Read1::CATCTTTTTCACGTA::Read2::Intron3'::3’ΔHygR + 5’ΔHygR::Intron5'::Read1::TCGCAGTATAAGGCC::Read2::Intron3'::3’ΔHygR + 5’ΔHygR::Intron5'::Read1::TCTCAGTTTAATCCC::Read2::Intron3'::3’ΔHygR +

5’ΔHygR::Intron5'::Read1::CCACCGTGTGATTTA::Read2::Intron3'::3’ΔHygR + 5’ΔHygR::Intron5'::Read1::CAACTTTATAACACC::Read2::Intron3'::3’ΔHygR + 5’ΔHygR::Intron5'::Read1::CTGCATTATAATCCC::Read2::Intron3'::3’ΔHygR + 5’ΔHygR::Intron5'::Read1::GGACTTTATCACGCA::Read2::Intron3'::3’ΔHygR + 5’ΔHygR::Intron5'::Read1::GGACCCTGTCACTCG::Read2::Intron3'::3’ΔHygR + 5’ΔHygR::Intron5'::Read1::CTTCGTTTTTAAATA::Read2::Intron3'::3’ΔHygR + 5’ΔHygR::Intron5'::Read1::GCACACTCTGACAAA::Read2::Intron3'::3’ΔHygR + 5’ΔHygR::Intron5'::Read1::ACACTATTTTAGTAT::Read2::Intron3'::3’ΔHygR + 5’ΔHygR::Intron5'::Read1::CTGCCCTTTCATTCC::Read2::Intron3'::3’ΔHygR + 5’ΔHygR::Intron5'::Read1::GTTCACTCTTACTAA::Read2::Intron3'::3’ΔHygR + 5’ΔHygR::Intron5'::Read1::TTCCCATGTTAAAAG::Read2::Intron3'::3’ΔHygR + 5’ΔHygR::Intron5'::Read1::TATCCGTTTGACTTC::Read2::Intron3'::3’ΔHygR + 5’ΔHygR::Intron5'::Read1::ATCCGCTATAACAAT::Read2::Intron3'::3’ΔHygR + 5’ΔHygR::Intron5'::Read1::CCGCTCTATCAAGCA::Read2::Intron3'::3’ΔHygR + 5’ΔHygR::Intron5'::Read1::CGACAGTCTCATAAT::Read2::Intron3'::3’ΔHygR + 5’ΔHygR::Intron5'::Read1::TAACACTGTTAAACA::Read2::Intron3'::3’ΔHygR + 5’ΔHygR::Intron5'::Read1::AATCGTTTTAATTCC::Read2::Intron3'::3’ΔHygR + 5’ΔHygR::Intron5'::Read1::TCTCCTTGTGAGGCT::Read2::Intron3'::3’ΔHygR + 5’ΔHygR::Intron5'::Read1::TCCCCCTCTGAGAAT::Read2::Intron3'::3’ΔHygR + 5’ΔHygR::Intron5'::Read1::TTTCAATCTGATTCC::Read2::Intron3'::3’ΔHygR + 5’ΔHygR::Intron5'::Read1::ATGCCCTTTAAAAGA::Read2::Intron3'::3’ΔHygR + 5’ΔHygR::Intron5'::Read1::TCCCTCTTTTAACCC::Read2::Intron3'::3’ΔHygR + 5’ΔHygR::Intron5'::Read1::TTTCCCTATAATATC::Read2::Intron3'::3’ΔHygR + 5’ΔHygR::Intron5'::Read1::ACACTGTCTCAGGTT::Read2::Intron3'::3’ΔHygR + 5’ΔHygR::Intron5'::Read1::ACTCATTTTGATACA::Read2::Intron3'::3’ΔHygR + 5’ΔHygR::Intron5'::Read1::CAACTTTATCACCCT::Read2::Intron3'::3’ΔHygR + 5’ΔHygR::Intron5'::Read1::TAGCAGTTTGATTCA::Read2::Intron3'::3’ΔHygR + 5’ΔHygR::Intron5'::Read1::ACGCGATATGATATT::Read2::Intron3'::3’ΔHygR + 5’ΔHygR::Intron5'::Read1::TACCAGTCTAAAGCA::Read2::Intron3'::3’ΔHygR + 5’ΔHygR::Intron5'::Read1::ACTCTTTGTCAGGTT::Read2::Intron3'::3’ΔHygR + 5’ΔHygR::Intron5'::Read1::AAGCCGTTTTAGATA::Read2::Intron3'::3’ΔHygR + 5’ΔHygR::Intron5'::Read1::ACTCCTTGTTATCGA::Read2::Intron3'::3’ΔHygR + 5’ΔHygR::Intron5'::Read1::CAACACTATTAACGG::Read2::Intron3'::3’ΔHygR + 5’ΔHygR::Intron5'::Read1::CGGCACTATTAAACG::Read2::Intron3'::3’ΔHygR + 5’ΔHygR::Intron5'::Read1::CGGCATTATCATCAC::Read2::Intron3'::3’ΔHygR + 5’ΔHygR::Intron5'::Read1::TTACCTTATGAAAAT::Read2::Intron3'::3’ΔHygR + 5’ΔHygR::Intron5'::Read1::ATTCGTTCTGAATAC::Read2::Intron3'::3’ΔHygR + 5’ΔHygR::Intron5'::Read1::TCTCCTTCTTACGTT::Read2::Intron3'::3’ΔHygR + 5’ΔHygR::Intron5'::Read1::ACCCCCTATTAACTA::Read2::Intron3'::3’ΔHygR + 5’ΔHygR::Intron5'::Read1::GCTCATTATTATACC::Read2::Intron3'::3’ΔHygR + 5’ΔHygR::Intron5'::Read1::TGACTTTTTAAAGTG::Read2::Intron3'::3’ΔHygR + 5’ΔHygR::Intron5'::Read1::TTCCTGTTTAACGTT::Read2::Intron3'::3’ΔHygR + 5’ΔHygR::Intron5'::Read1::AAGCACTATGACTCC::Read2::Intron3'::3’ΔHygR + 5’ΔHygR::Intron5'::Read1::AATCTGTATAACCGA::Read2::Intron3'::3’ΔHygR + 5’ΔHygR::Intron5'::Read1::CTTCTGTCTCATTCC::Read2::Intron3'::3’ΔHygR + 5’ΔHygR::Intron5'::Read1::CATCGATGTCACAAC::Read2::Intron3'::3’ΔHygR + 5’ΔHygR::Intron5'::Read1::ATACATTCTCATGAG::Read2::Intron3'::3’ΔHygR + 5’ΔHygR::Intron5'::Read1::TCCCGCTATCACTTG::Read2::Intron3'::3’ΔHygR + 5’ΔHygR::Intron5'::Read1::CTCCTGTATCACCTA::Read2::Intron3'::3’ΔHygR + 5’ΔHygR::Intron5'::Read1::TAGCTGTGTCACATG::Read2::Intron3'::3’ΔHygR + 5’ΔHygR::Intron5'::Read1::GCACCATCTTAATAA::Read2::Intron3'::3’ΔHygR + 5’ΔHygR::Intron5'::Read1::TGGCTCTTTCATTCC::Read2::Intron3'::3’ΔHygR + 5’ΔHygR::Intron5'::Read1::AATCACTCTAAGCAC::Read2::Intron3'::3’ΔHygR + 5’ΔHygR::Intron5'::Read1::CATCTATTTAAGTCC::Read2::Intron3'::3’ΔHygR + 5’ΔHygR::Intron5'::Read1::CTACGCTTTAATCAC::Read2::Intron3'::3’ΔHygR + 5’ΔHygR::Intron5'::Read1::TCACTTTGTAACGAT::Read2::Intron3'::3’ΔHygR + 5’ΔHygR::Intron5'::Read1::TTCCCTTTTGACTCT::Read2::Intron3'::3’ΔHygR + 5’ΔHygR::Intron5'::Read1::CGACAGTATAACCCT::Read2::Intron3'::3’ΔHygR + 5’ΔHygR::Intron5'::Read1::ACTCATTATAAAAAC::Read2::Intron3'::3’ΔHygR + 5’ΔHygR::Intron5'::Read1::GGTCCTTGTTACATA::Read2::Intron3'::3’ΔHygR + 5’ΔHygR::Intron5'::Read1::TATCTTTCTCACCTC::Read2::Intron3'::3’ΔHygR + 5’ΔHygR::Intron5'::Read1::ACGCTATCTCAGTTT::Read2::Intron3'::3’ΔHygR + 5’ΔHygR::Intron5'::Read1::ATACTGTCTGAACCT::Read2::Intron3'::3’ΔHygR + 5’ΔHygR::Intron5'::Read1::AGACGCTATAACAAA::Read2::Intron3'::3’ΔHygR + 5’ΔHygR::Intron5'::Read1::AAGCGATCTCATTGG::Read2::Intron3'::3’ΔHygR + 5’ΔHygR::Intron5'::Read1::CCTCCGTATCAGACA::Read2::Intron3'::3’ΔHygR + 5’ΔHygR::Intron5'::Read1::TCTCATTTTCACAGT::Read2::Intron3'::3’ΔHygR + 5’ΔHygR::Intron5'::Read1::CACCGGTCTCAACCA::Read2::Intron3'::3’ΔHygR + 5’ΔHygR::Intron5'::Read1::CGTCAATCTAACGGA::Read2::Intron3'::3’ΔHygR + 5’ΔHygR::Intron5'::Read1::GCACTATCTAAGGCA::Read2::Intron3'::3’ΔHygR +

5’ΔHygR::Intron5'::Read1::TATCCCTTTGAGATT::Read2::Intron3'::3’ΔHygR + 5’ΔHygR::Intron5'::Read1::AAGCAATTTCAGCCC::Read2::Intron3'::3’ΔHygR + 5’ΔHygR::Intron5'::Read1::CCTCCTTGTCACGTA::Read2::Intron3'::3’ΔHygR + 5’ΔHygR::Intron5'::Read1::GCGCTGTGTCAATCA::Read2::Intron3'::3’ΔHygR + 5’ΔHygR::Intron5'::Read1::CATCTTTATCAATGA::Read2::Intron3'::3’ΔHygR + 5’ΔHygR::Intron5'::Read1::TGACAGTGTCACCCG::Read2::Intron3'::3’ΔHygR + 5’ΔHygR::Intron5'::Read1::CTTCTATCTCAGCCT::Read2::Intron3'::3’ΔHygR + 5’ΔHygR::Intron5'::Read1::CTCCGTTTTAACCTT::Read2::Intron3'::3’ΔHygR + 5’ΔHygR::Intron5'::Read1::ACGCCATGTTACTAT::Read2::Intron3'::3’ΔHygR + 5’ΔHygR::Intron5'::Read1::GAACTTTGTCATTTT::Read2::Intron3'::3’ΔHygR + 5’ΔHygR::Intron5'::Read1::CGACCGTCTCATCCT::Read2::Intron3'::3’ΔHygR + 5’ΔHygR::Intron5'::Read1::TACCGTTCTCATACA::Read2::Intron3'::3’ΔHygR + 5’ΔHygR::Intron5'::Read1::TGTCGCTCTTAATGC::Read2::Intron3'::3’ΔHygR + 5’ΔHygR::Intron5'::Read1::CCACGATCTTATCAA::Read2::Intron3'::3’ΔHygR + 5’ΔHygR::Intron5'::Read1::GGACAGTCTCAAATT::Read2::Intron3'::3’ΔHygR + 5’ΔHygR::Intron5'::Read1::TGTCGTTCTGACGTC::Read2::Intron3'::3’ΔHygR + 5’ΔHygR::Intron5'::Read1::CTCCCATTTGATCTT::Read2::Intron3'::3’ΔHygR + 5’ΔHygR::Intron5'::Read1::AGGCCTTATTATGTA::Read2::Intron3'::3’ΔHygR + 5’ΔHygR::Intron5'::Read1::TTCCTTTTTCATATA::Read2::Intron3'::3’ΔHygR + 5’ΔHygR::Intron5'::Read1::GTCCGCTCTAACATC::Read2::Intron3'::3’ΔHygR + 5’ΔHygR::Intron5'::Read1::TTACACTATCATTCG::Read2::Intron3'::3’ΔHygR + 5’ΔHygR::Intron5'::Read1::TAACGTTCTTATATG::Read2::Intron3'::3’ΔHygR + 5’ΔHygR::Intron5'::Read1::TCACTTTCTAACTAA::Read2::Intron3'::3’ΔHygR + 5’ΔHygR::Intron5'::Read1::TTGCCATTTGACGTC::Read2::Intron3'::3’ΔHygR + 5’ΔHygR::Intron5'::Read1::ATGCTATCTTAATAT::Read2::Intron3'::3’ΔHygR + 5’ΔHygR::Intron5'::Read1::ATTCCATGTTATTTA::Read2::Intron3'::3’ΔHygR + 5’ΔHygR::Intron5'::Read1::CCACAATTTCAACGT::Read2::Intron3'::3’ΔHygR + 5’ΔHygR::Intron5'::Read1::CTTCTCTGTGACTGC::Read2::Intron3'::3’ΔHygR + 5’ΔHygR::Intron5'::Read1::GCACACTTTAAACTA::Read2::Intron3'::3’ΔHygR + 5’ΔHygR::Intron5'::Read1::GGGCCCTATGAAGCA::Read2::Intron3'::3’ΔHygR + 5’ΔHygR::Intron5'::Read1::CTTCTCTTTCATCAC::Read2::Intron3'::3’ΔHygR + 5’ΔHygR::Intron5'::Read1::GTCCCCTATCAAATC::Read2::Intron3'::3’ΔHygR + 5’ΔHygR::Intron5'::Read1::TCTCAATTTTATTCC::Read2::Intron3'::3’ΔHygR + 5’ΔHygR::Intron5'::Read1::GATCTCTGTCACCGA::Read2::Intron3'::3’ΔHygR + 5’ΔHygR::Intron5'::Read1::GTCCTATTTTATCCC::Read2::Intron3'::3’ΔHygR + 5’ΔHygR::Intron5'::Read1::GGTCAGTATTATGCT::Read2::Intron3'::3’ΔHygR + 5’ΔHygR::Intron5'::Read1::TTTCCATGTCACATC::Read2::Intron3'::3’ΔHygR + 5’ΔHygR::Intron5'::Read1::GGGCAATCTTATCCG::Read2::Intron3'::3’ΔHygR + 5’ΔHygR::Intron5'::Read1::TTCCGCTTTCAATTG::Read2::Intron3'::3’ΔHygR + 5’ΔHygR::Intron5'::Read1::GTCCGGTTTCAAGTG::Read2::Intron3'::3’ΔHygR + 5’ΔHygR::Intron5'::Read1::GTGCAGTATTACTGG::Read2::Intron3'::3’ΔHygR + 5’ΔHygR::Intron5'::Read1::ACTCATTTTAACGGA::Read2::Intron3'::3’ΔHygR + 5’ΔHygR::Intron5'::Read1::CGTTAGTATTAAAAC::Read2::Intron3'::3’ΔHygR + 5’ΔHygR::Intron5'::Read1::GACCCGTATTACCTT::Read2::Intron3'::3’ΔHygR + 5’ΔHygR::Intron5'::Read1::TGCCCTTCTAACTGC::Read2::Intron3'::3’ΔHygR + 5’ΔHygR::Intron5'::Read1::CACCGATGTCAAGTA::Read2::Intron3'::3’ΔHygR + 5’ΔHygR::Intron5'::Read1::TCACAATCTCATAGA::Read2::Intron3'::3’ΔHygR + 5’ΔHygR::Intron5'::Read1::GCACCTTCTTAGCAC::Read2::Intron3'::3’ΔHygR + 5’ΔHygR::Intron5'::Read1::GTTCACTATCACACG::Read2::Intron3'::3’ΔHygR + 5’ΔHygR::Intron5'::Read1::TACCGGTATCACTCA::Read2::Intron3'::3’ΔHygR + 5’ΔHygR::Intron5'::Read1::ACTCAGTTTCAAATG::Read2::Intron3'::3’ΔHygR + 5’ΔHygR::Intron5'::Read1::CAGCCTTTTGATCAC::Read2::Intron3'::3’ΔHygR + 5’ΔHygR::Intron5'::Read1::CCCCGCTGTCACTAG::Read2::Intron3'::3’ΔHygR + 5’ΔHygR::Intron5'::Read1::TCACCATTTGAACCT::Read2::Intron3'::3’ΔHygR + 5’ΔHygR::Intron5'::Read1::TCGCCATATAATCTC::Read2::Intron3'::3’ΔHygR + 5’ΔHygR::Intron5'::Read1::CCACTCTTTTAACCT::Read2::Intron3'::3’ΔHygR + 5’ΔHygR::Intron5'::Read1::AAACCTTATGAACTC::Read2::Intron3'::3’ΔHygR + 5’ΔHygR::Intron5'::Read1::ATCCCTTTTAAGTCC::Read2::Intron3'::3’ΔHygR + 5’ΔHygR::Intron5'::Read1::ATTCGCTATTAGTCA::Read2::Intron3'::3’ΔHygR + 5’ΔHygR::Intron5'::Read1::ATCCTATTTAAGCTT::Read2::Intron3'::3’ΔHygR + 5’ΔHygR::Intron5'::Read1::TAACCGTGTCATCCT::Read2::Intron3'::3’ΔHygR + 5’ΔHygR::Intron5'::Read1::CGTCGTTGTTATCTA::Read2::Intron3'::3’ΔHygR + 5’ΔHygR::Intron5'::Read1::TGTCATTTTGAGCCT::Read2::Intron3'::3’ΔHygR + 5’ΔHygR::Intron5'::Read1::GACCCGTTTTATCAA::Read2::Intron3'::3’ΔHygR + 5’ΔHygR::Intron5'::Read1::GCTCACTATCACGAT::Read2::Intron3'::3’ΔHygR + 5’ΔHygR::Intron5'::Read1::TTTCTATTTCACATA::Read2::Intron3'::3’ΔHygR + 5’ΔHygR::Intron5'::Read1::TTGCAATCTAAAACA::Read2::Intron3'::3’ΔHygR + 5’ΔHygR::Intron5'::Read1::ACACTCTGTCAGCAA::Read2::Intron3'::3’ΔHygR + 5’ΔHygR::Intron5'::Read1::CTTCATTCTCAAGCG::Read2::Intron3'::3’ΔHygR + 5’ΔHygR::Intron5'::Read1::GCTCATTATAACGTC::Read2::Intron3'::3’ΔHygR +

5’ΔHygR::Intron5'::Read1::ATACAATATTAGGAT::Read2::Intron3'::3’ΔHygR + 5’ΔHygR::Intron5'::Read1::CTCCCTTATCACTTC::Read2::Intron3'::3’ΔHygR + 5’ΔHygR::Intron5'::Read1::CAGCCGTGTTAATCA::Read2::Intron3'::3’ΔHygR + 5’ΔHygR::Intron5'::Read1::GCACGATCTCATGGC::Read2::Intron3'::3’ΔHygR + 5’ΔHygR::Intron5'::Read1::TATCCATCTCACCCT::Read2::Intron3'::3’ΔHygR + 5’ΔHygR::Intron5'::Read1::ACACACTCTGACAAC::Read2::Intron3'::3’ΔHygR + 5’ΔHygR::Intron5'::Read1::AGTCTGTTTAAACTC::Read2::Intron3'::3’ΔHygR + 5’ΔHygR::Intron5'::Read1::GTTCGCTTTGAACAT::Read2::Intron3'::3’ΔHygR + 5’ΔHygR::Intron5'::Read1::CGCCCTTTTCACTAT::Read2::Intron3'::3’ΔHygR + 5’ΔHygR::Intron5'::Read1::CAACTGTTTTATCAC::Read2::Intron3'::3’ΔHygR + 5’ΔHygR::Intron5'::Read1::CTACTTTATAAAGCA::Read2::Intron3'::3’ΔHygR + 5’ΔHygR::Intron5'::Read1::CAGCTCTCTTATTCT::Read2::Intron3'::3’ΔHygR + 5’ΔHygR::Intron5'::Read1::GTGCAATCTTACATA::Read2::Intron3'::3’ΔHygR + 5’ΔHygR::Intron5'::Read1::ATACTTTGTCAAGCA::Read2::Intron3'::3’ΔHygR + 5’ΔHygR::Intron5'::Read1::CCCCCTTATTACCGC::Read2::Intron3'::3’ΔHygR + 5’ΔHygR::Intron5'::Read1::GCTCACTTTAACTTC::Read2::Intron3'::3’ΔHygR + 5’ΔHygR::Intron5'::Read1::ACCCTCTGTCATGCA::Read2::Intron3'::3’ΔHygR + 5’ΔHygR::Intron5'::Read1::CAACTCTGTGAGCTT::Read2::Intron3'::3’ΔHygR + 5’ΔHygR::Intron5'::Read1::CATCCCTGTAATTCT::Read2::Intron3'::3’ΔHygR + 5’ΔHygR::Intron5'::Read1::ATTCAATTTTAACCG::Read2::Intron3'::3’ΔHygR + 5’ΔHygR::Intron5'::Read1::GACCCATTTGAACGC::Read2::Intron3'::3’ΔHygR + 5’ΔHygR::Intron5'::Read1::GATCGGTGTCAGCTT::Read2::Intron3'::3’ΔHygR + 5’ΔHygR::Intron5'::Read1::GCTCTTTCTCATCCA::Read2::Intron3'::3’ΔHygR + 5’ΔHygR::Intron5'::Read1::ATACCGTTTCATGTC::Read2::Intron3'::3’ΔHygR + 5’ΔHygR::Intron5'::Read1::CAGCTTTATAACGAA::Read2::Intron3'::3’ΔHygR + 5’ΔHygR::Intron5'::Read1::GTTCGTTTTCAGCTC::Read2::Intron3'::3’ΔHygR + 5’ΔHygR::Intron5'::Read1::TTCCCTTCTTACTTG::Read2::Intron3'::3’ΔHygR + 5’ΔHygR::Intron5'::Read1::TAACAATTTCACCAC::Read2::Intron3'::3’ΔHygR + 5’ΔHygR::Intron5'::Read1::AACCAGTTTCACTTA::Read2::Intron3'::3’ΔHygR + 5’ΔHygR::Intron5'::Read1::ATTCGGTTTTAACTC::Read2::Intron3'::3’ΔHygR + 5’ΔHygR::Intron5'::Read1::TTCCGTTTTCAGTGG::Read2::Intron3'::3’ΔHygR + 5’ΔHygR::Intron5'::Read1::CGGCCATATAAAAGC::Read2::Intron3'::3’ΔHygR + 5’ΔHygR::Intron5'::Read1::TCGCCTTGTCACTCT::Read2::Intron3'::3’ΔHygR + 5’ΔHygR::Intron5'::Read1::TGTCGGTCTTAACTG::Read2::Intron3'::3’ΔHygR + 5’ΔHygR::Intron5'::Read1::CAACCTTATGACACA::Read2::Intron3'::3’ΔHygR + 5’ΔHygR::Intron5'::Read1::CCCCGATGTGAGGTC::Read2::Intron3'::3’ΔHygR + 5’ΔHygR::Intron5'::Read1::TTCCATTGTCACAGA::Read2::Intron3'::3’ΔHygR + 5’ΔHygR::Intron5'::Read1::CAACAGTTTCAGTTA::Read2::Intron3'::3’ΔHygR + 5’ΔHygR::Intron5'::Read1::CATCTCTGTTAGTTC::Read2::Intron3'::3’ΔHygR + 5’ΔHygR::Intron5'::Read1::ACGCCCTCTTAACTC::Read2::Intron3'::3’ΔHygR + 5’ΔHygR::Intron5'::Read1::CCTCTCTTTTACACC::Read2::Intron3'::3’ΔHygR + 5’ΔHygR::Intron5'::Read1::TCGCATTCTAAGTTA::Read2::Intron3'::3’ΔHygR + 5’ΔHygR::Intron5'::Read1::AACCGGTTTCAACAT::Read2::Intron3'::3’ΔHygR + 5’ΔHygR::Intron5'::Read1::GTACCTTCTAATTCA::Read2::Intron3'::3’ΔHygR + 5’ΔHygR::Intron5'::Read1::TCACACTATCATGCA::Read2::Intron3'::3’ΔHygR + 5’ΔHygR::Intron5'::Read1::ACTCCCTCTCACACT::Read2::Intron3'::3’ΔHygR + 5’ΔHygR::Intron5'::Read1::TTGCACTGTTAACAC::Read2::Intron3'::3’ΔHygR + 5’ΔHygR::Intron5'::Read1::GGCCGTTGTGAAATC::Read2::Intron3'::3’ΔHygR + 5’ΔHygR::Intron5'::Read1::TAGCTCTCTTATTAC::Read2::Intron3'::3’ΔHygR + 5’ΔHygR::Intron5'::Read1::GCGCTCTTTCACACT::Read2::Intron3'::3’ΔHygR + 5’ΔHygR::Intron5'::Read1::TTCCTTTCTTATGAT::Read2::Intron3'::3’ΔHygR + 5’ΔHygR::Intron5'::Read1::GGACAATGTAATCCT::Read2::Intron3'::3’ΔHygR + 5’ΔHygR::Intron5'::Read1::GTGCATTATCAGGTT::Read2::Intron3'::3’ΔHygR + 5’ΔHygR::Intron5'::Read1::TTGCTGTATGACTAT::Read2::Intron3'::3’ΔHygR + 5’ΔHygR::Intron5'::Read1::ACTCTCTTTCACTAG::Read2::Intron3'::3’ΔHygR + 5’ΔHygR::Intron5'::Read1::CCTCAGTATTAGCAT::Read2::Intron3'::3’ΔHygR + 5’ΔHygR::Intron5'::Read1::TCTCCTTCTAAACTA::Read2::Intron3'::3’ΔHygR + 5’ΔHygR::Intron5'::Read1::AGTCGCTCTTATACA::Read2::Intron3'::3’ΔHygR + 5’ΔHygR::Intron5'::Read1::CCTCATTGTCAGGCT::Read2::Intron3'::3’ΔHygR + 5’ΔHygR::Intron5'::Read1::TACCCATATGACATC::Read2::Intron3'::3’ΔHygR + 5’ΔHygR::Intron5'::Read1::TTTCCATTTAAGACA::Read2::Intron3'::3’ΔHygR + 5’ΔHygR::Intron5'::Read1::AGGCGCTGTTACCTC::Read2::Intron3'::3’ΔHygR + 5’ΔHygR::Intron5'::Read1::CCACTTTCTCATTGC::Read2::Intron3'::3’ΔHygR + 5’ΔHygR::Intron5'::Read1::TACCATTTTAATTCC::Read2::Intron3'::3’ΔHygR + 5’ΔHygR::Intron5'::Read1::TGCCTGTGTTATTTA::Read2::Intron3'::3’ΔHygR + 5’ΔHygR::Intron5'::Read1::ATTCTCTATTATCCA::Read2::Intron3'::3’ΔHygR + 5’ΔHygR::Intron5'::Read1::TGCCATTCTCAACAG::Read2::Intron3'::3’ΔHygR + 5’ΔHygR::Intron5'::Read1::ACTCCATTTTATTTG::Read2::Intron3'::3’ΔHygR + 5’ΔHygR::Intron5'::Read1::GTACATTCTTATCCT::Read2::Intron3'::3’ΔHygR + 5’ΔHygR::Intron5'::Read1::TACCTTTCTTAGGCC::Read2::Intron3'::3’ΔHygR +

5’ΔHygR::Intron5'::Read1::TGTCCCTCTTACCAC::Read2::Intron3'::3’ΔHygR + 5’ΔHygR::Intron5'::Read1::ACACTATATGATACT::Read2::Intron3'::3’ΔHygR + 5’ΔHygR::Intron5'::Read1::ATACCCTGTCAGGTC::Read2::Intron3'::3’ΔHygR + 5’ΔHygR::Intron5'::Read1::ACCCGTTCTCATCTC::Read2::Intron3'::3’ΔHygR + 5’ΔHygR::Intron5'::Read1::CGGCTTTGTCACATC::Read2::Intron3'::3’ΔHygR + 5’ΔHygR::Intron5'::Read1::TGGCGTTATTACTCC::Read2::Intron3'::3’ΔHygR + 5’ΔHygR::Intron5'::Read1::TCGCAATCTCACCAA::Read2::Intron3'::3’ΔHygR + 5’ΔHygR::Intron5'::Read1::AGCCCTTTTCACTGA::Read2::Intron3'::3’ΔHygR + 5’ΔHygR::Intron5'::Read1::ACGCGATCTAACAAA::Read2::Intron3'::3’ΔHygR + 5’ΔHygR::Intron5'::Read1::GTTCAATATCAGGTT::Read2::Intron3'::3’ΔHygR + 5’ΔHygR::Intron5'::Read1::CGCCACTATGACATT::Read2::Intron3'::3’ΔHygR + 5’ΔHygR::Intron5'::Read1::GAACGGTATTATGTC::Read2::Intron3'::3’ΔHygR + 5’ΔHygR::Intron5'::Read1::GTCCCATATGAACAT::Read2::Intron3'::3’ΔHygR + 5’ΔHygR::Intron5'::Read1::TTGCTCTTTAATTTT::Read2::Intron3'::3’ΔHygR + 5’ΔHygR::Intron5'::Read1::ACTCGCTGTGACGGG::Read2::Intron3'::3’ΔHygR + 5’ΔHygR::Intron5'::Read1::TAGCTATGTAATGAG::Read2::Intron3'::3’ΔHygR + 5’ΔHygR::Intron5'::Read1::TATCCCTTTCAAGTC::Read2::Intron3'::3’ΔHygR + 5’ΔHygR::Intron5'::Read1::TCACAATCTTAGATA::Read2::Intron3'::3’ΔHygR + 5’ΔHygR::Intron5'::Read1::TCGCATTGTTAATAT::Read2::Intron3'::3’ΔHygR + 5’ΔHygR::Intron5'::Read1::TGCCGCTCTAATCAA::Read2::Intron3'::3’ΔHygR + 5’ΔHygR::Intron5'::Read1::GGACACTCTAAGAAA::Read2::Intron3'::3’ΔHygR + 5’ΔHygR::Intron5'::Read1::TTGCCATCTTAAAGT::Read2::Intron3'::3’ΔHygR + 5’ΔHygR::Intron5'::Read1::CACCATTTTAAGAAC::Read2::Intron3'::3’ΔHygR + 5’ΔHygR::Intron5'::Read1::ACCCGTTCTAACCCT::Read2::Intron3'::3’ΔHygR + 5’ΔHygR::Intron5'::Read1::TAACTTTGTAAATTC::Read2::Intron3'::3’ΔHygR + 5’ΔHygR::Intron5'::Read1::AATCTTTTTCAGGAT::Read2::Intron3'::3’ΔHygR + 5’ΔHygR::Intron5'::Read1::CCACCCTCTGATTTT::Read2::Intron3'::3’ΔHygR + 5’ΔHygR::Intron5'::Read1::TATCTGTCTCACTAG::Read2::Intron3'::3’ΔHygR + 5’ΔHygR::Intron5'::Read1::TGACCTTCTTAACCT::Read2::Intron3'::3’ΔHygR + 5’ΔHygR::Intron5'::Read1::ACGCTTTATCAACAC::Read2::Intron3'::3’ΔHygR + 5’ΔHygR::Intron5'::Read1::CTACTTTCTAATTTC::Read2::Intron3'::3’ΔHygR + 5’ΔHygR::Intron5'::Read1::TTTCCGTCTAATAAT::Read2::Intron3'::3’ΔHygR + 5’ΔHygR::Intron5'::Read1::TCGCCCTGTTAGTAC::Read2::Intron3'::3’ΔHygR + 5’ΔHygR::Intron5'::Read1::TGCCTCTTTCAACTT::Read2::Intron3'::3’ΔHygR + 5’ΔHygR::Intron5'::Read1::CCGCTATTTGAACTT::Read2::Intron3'::3’ΔHygR + 5’ΔHygR::Intron5'::Read1::AGGCGTTGTTATAAT::Read2::Intron3'::3’ΔHygR + 5’ΔHygR::Intron5'::Read1::CATCACTTTTAACTA::Read2::Intron3'::3’ΔHygR + 5’ΔHygR::Intron5'::Read1::ATTCTTTATCATTTA::Read2::Intron3'::3’ΔHygR + 5’ΔHygR::Intron5'::Read1::CCACCTTCTAAGCTA::Read2::Intron3'::3’ΔHygR + 5’ΔHygR::Intron5'::Read1::AGTCTCTCTGAAGTC::Read2::Intron3'::3’ΔHygR + 5’ΔHygR::Intron5'::Read1::TTTCTCTTTAACTTT::Read2::Intron3'::3’ΔHygR + 5’ΔHygR::Intron5'::Read1::TTACCTTTTTAATGA::Read2::Intron3'::3’ΔHygR + 5’ΔHygR::Intron5'::Read1::GCTCGCTTTGACACT::Read2::Intron3'::3’ΔHygR + 5’ΔHygR::Intron5'::Read1::TGTCCATGTAACATA::Read2::Intron3'::3’ΔHygR + 5’ΔHygR::Intron5'::Read1::TTACCATTTTAAACT::Read2::Intron3'::3’ΔHygR + 5’ΔHygR::Intron5'::Read1::AACCCGTCTCATTGT::Read2::Intron3'::3’ΔHygR + 5’ΔHygR::Intron5'::Read1::ATTCGCTTTTAATAC::Read2::Intron3'::3’ΔHygR + 5’ΔHygR::Intron5'::Read1::TCACTATGTTACCTC::Read2::Intron3'::3’ΔHygR + 5’ΔHygR::Intron5'::Read1::ATTCGATTTCACCCC::Read2::Intron3'::3’ΔHygR + 5’ΔHygR::Intron5'::Read1::TAACTATATTATTAG::Read2::Intron3'::3’ΔHygR + 5’ΔHygR::Intron5'::Read1::CGGCTGTTTAACCTC::Read2::Intron3'::3’ΔHygR + 5’ΔHygR::Intron5'::Read1::GGACTGTCTTAATCA::Read2::Intron3'::3’ΔHygR + 5’ΔHygR::Intron5'::Read1::ACCCACTATCATCAC::Read2::Intron3'::3’ΔHygR + 5’ΔHygR::Intron5'::Read1::AGTCAATATAACGTA::Read2::Intron3'::3’ΔHygR + 5’ΔHygR::Intron5'::Read1::TACCGGTCTGAGAAC::Read2::Intron3'::3’ΔHygR + 5’ΔHygR::Intron5'::Read1::ACCCAGTATTAACTC::Read2::Intron3'::3’ΔHygR + 5’ΔHygR::Intron5'::Read1::GTTCACTTTTAGGAG::Read2::Intron3'::3’ΔHygR + 5’ΔHygR::Intron5'::Read1::CATCGGTATAACACC::Read2::Intron3'::3’ΔHygR + 5’ΔHygR::Intron5'::Read1::CTGCCCTATCATCAA::Read2::Intron3'::3’ΔHygR + 5’ΔHygR::Intron5'::Read1::TCGCCTTCTGAATTC::Read2::Intron3'::3’ΔHygR + 5’ΔHygR::Intron5'::Read1::TTACCATTTCATAAT::Read2::Intron3'::3’ΔHygR + 5’ΔHygR::Intron5'::Read1::ACTCGATATCAGTGC::Read2::Intron3'::3’ΔHygR + 5’ΔHygR::Intron5'::Read1::TCTCAGTGTAAACGA::Read2::Intron3'::3’ΔHygR + 5’ΔHygR::Intron5'::Read1::TTCCGGTGTCATCCT::Read2::Intron3'::3’ΔHygR + 5’ΔHygR::Intron5'::Read1::TATCCCTATAAATCA::Read2::Intron3'::3’ΔHygR + 5’ΔHygR::Intron5'::Read1::TCCCCTTATAAACGA::Read2::Intron3'::3’ΔHygR + 5’ΔHygR::Intron5'::Read1::ACACAATATTACATT::Read2::Intron3'::3’ΔHygR + 5’ΔHygR::Intron5'::Read1::AGTCCGTCTAACTAC::Read2::Intron3'::3’ΔHygR + 5’ΔHygR::Intron5'::Read1::GAACTTTATCAGCAC::Read2::Intron3'::3’ΔHygR + 5’ΔHygR::Intron5'::Read1::GCTCTTTGTCACAAC::Read2::Intron3'::3’ΔHygR +

5’ΔHygR::Intron5'::Read1::TTTCGTTATTAGGGT::Read2::Intron3'::3’ΔHygR + 5’ΔHygR::Intron5'::Read1::AGTCTCTATAATCAC::Read2::Intron3'::3’ΔHygR + 5’ΔHygR::Intron5'::Read1::ACCCAATATTATTTA::Read2::Intron3'::3’ΔHygR + 5’ΔHygR::Intron5'::Read1::GAACACTTTCAGCCC::Read2::Intron3'::3’ΔHygR + 5’ΔHygR::Intron5'::Read1::AATCCGTGTCATACA::Read2::Intron3'::3’ΔHygR + 5’ΔHygR::Intron5'::Read1::ACACTTTCTAACGTC::Read2::Intron3'::3’ΔHygR + 5’ΔHygR::Intron5'::Read1::GAACTCTCTCAGCAT::Read2::Intron3'::3’ΔHygR + 5’ΔHygR::Intron5'::Read1::ACGCTCTTTTATTTA::Read2::Intron3'::3’ΔHygR + 5’ΔHygR::Intron5'::Read1::CCGCTATGTTAGATT::Read2::Intron3'::3’ΔHygR + 5’ΔHygR::Intron5'::Read1::TCACGGTGTCAATCA::Read2::Intron3'::3’ΔHygR + 5’ΔHygR::Intron5'::Read1::TTGCTGTTTTAGTCA::Read2::Intron3'::3’ΔHygR + 5’ΔHygR::Intron5'::Read1::CTGCAATATCAAGCG::Read2::Intron3'::3’ΔHygR + 5’ΔHygR::Intron5'::Read1::AAACAATCTTATGTG::Read2::Intron3'::3’ΔHygR + 5’ΔHygR::Intron5'::Read1::CCGCATTATGACATT::Read2::Intron3'::3’ΔHygR + 5’ΔHygR::Intron5'::Read1::TAACATTTTTAACAT::Read2::Intron3'::3’ΔHygR + 5’ΔHygR::Intron5'::Read1::TCACAGTTTTACATT::Read2::Intron3'::3’ΔHygR + 5’ΔHygR::Intron5'::Read1::GCGCTGTTTAATCCT::Read2::Intron3'::3’ΔHygR + 5’ΔHygR::Intron5'::Read1::GGTCGTTGTTAACCT::Read2::Intron3'::3’ΔHygR + 5’ΔHygR::Intron5'::Read1::TCACTATATCAAGTT::Read2::Intron3'::3’ΔHygR + 5’ΔHygR::Intron5'::Read1::TTTCTGTTTCAATCG::Read2::Intron3'::3’ΔHygR + 5’ΔHygR::Intron5'::Read1::GTCCTTTGTTAATCG::Read2::Intron3'::3’ΔHygR + 5’ΔHygR::Intron5'::Read1::TCCCGCTGTCATTCG::Read2::Intron3'::3’ΔHygR + 5’ΔHygR::Intron5'::Read1::CACCATTCTCATATT::Read2::Intron3'::3’ΔHygR + 5’ΔHygR::Intron5'::Read1::TCTCCATGTAAAGTA::Read2::Intron3'::3’ΔHygR + 5’ΔHygR::Intron5'::Read1::AACCGCTCTGACATT::Read2::Intron3'::3’ΔHygR + 5’ΔHygR::Intron5'::Read1::ACTCCATATCAAAAT::Read2::Intron3'::3’ΔHygR + 5’ΔHygR::Intron5'::Read1::CACCGATCTCATCAC::Read2::Intron3'::3’ΔHygR + 5’ΔHygR::Intron5'::Read1::CAGCCCTTTGAATAG::Read2::Intron3'::3’ΔHygR + 5’ΔHygR::Intron5'::Read1::CCACCTTGTAAACTC::Read2::Intron3'::3’ΔHygR + 5’ΔHygR::Intron5'::Read1::CAGCCTTTTCACAAA::Read2::Intron3'::3’ΔHygR + 5’ΔHygR::Intron5'::Read1::CTTCAGTCTTAACCC::Read2::Intron3'::3’ΔHygR + 5’ΔHygR::Intron5'::Read1::GGTCTATATAAGTTG::Read2::Intron3'::3’ΔHygR + 5’ΔHygR::Intron5'::Read1::TCGCTGTATAAGACC::Read2::Intron3'::3’ΔHygR + 5’ΔHygR::Intron5'::Read1::GATCCTTATGACCTA::Read2::Intron3'::3’ΔHygR + 5’ΔHygR::Intron5'::Read1::GAGCCTTCTGACATC::Read2::Intron3'::3’ΔHygR + 5’ΔHygR::Intron5'::Read1::CCACCTTGTTAACAA::Read2::Intron3'::3’ΔHygR + 5’ΔHygR::Intron5'::Read1::TGACGTTGTTATGGT::Read2::Intron3'::3’ΔHygR + 5’ΔHygR::Intron5'::Read1::AAGCAATTTTACCAC::Read2::Intron3'::3’ΔHygR + 5’ΔHygR::Intron5'::Read1::TCTCTATATGACATC::Read2::Intron3'::3’ΔHygR + 5’ΔHygR::Intron5'::Read1::ACGCCTTATCAGCTC::Read2::Intron3'::3’ΔHygR + 5’ΔHygR::Intron5'::Read1::ATCCTCTTTGAATCT::Read2::Intron3'::3’ΔHygR + 5’ΔHygR::Intron5'::Read1::TCCCTATTTAAATGT::Read2::Intron3'::3’ΔHygR + 5’ΔHygR::Intron5'::Read1::ACGCACTCTCACTTC::Read2::Intron3'::3’ΔHygR + 5’ΔHygR::Intron5'::Read1::GATCTCTCTGACCGT::Read2::Intron3'::3’ΔHygR + 5’ΔHygR::Intron5'::Read1::CTTCCCTCTTATGTC::Read2::Intron3'::3’ΔHygR + 5’ΔHygR::Intron5'::Read1::TAACAATGTCAAATA::Read2::Intron3'::3’ΔHygR + 5’ΔHygR::Intron5'::Read1::ATTCTATATGACGCC::Read2::Intron3'::3’ΔHygR + 5’ΔHygR::Intron5'::Read1::CTACGATTTCAGCCT::Read2::Intron3'::3’ΔHygR + 5’ΔHygR::Intron5'::Read1::CATCCGTATTATTAA::Read2::Intron3'::3’ΔHygR + 5’ΔHygR::Intron5'::Read1::AATCACTATAACCCG::Read2::Intron3'::3’ΔHygR + 5’ΔHygR::Intron5'::Read1::ATTCGCTTTAAAAAC::Read2::Intron3'::3’ΔHygR + 5’ΔHygR::Intron5'::Read1::CATCTTTTTAACATC::Read2::Intron3'::3’ΔHygR + 5’ΔHygR::Intron5'::Read1::GATCTATATGATAAA::Read2::Intron3'::3’ΔHygR + 5’ΔHygR::Intron5'::Read1::TTCCATTCTGAATTC::Read2::Intron3'::3’ΔHygR + 5’ΔHygR::Intron5'::Read1::TCTCCATGTTAGCAA::Read2::Intron3'::3’ΔHygR + 5’ΔHygR::Intron5'::Read1::TTACCATTTAATACT::Read2::Intron3'::3’ΔHygR + 5’ΔHygR::Intron5'::Read1::TCACCATATCATTGT::Read2::Intron3'::3’ΔHygR + 5’ΔHygR::Intron5'::Read1::ATCCTTTGTTAACAA::Read2::Intron3'::3’ΔHygR + 5’ΔHygR::Intron5'::Read1::GCACCATCTCAGTAT::Read2::Intron3'::3’ΔHygR + 5’ΔHygR::Intron5'::Read1::TTGCGATTTAAAGCC::Read2::Intron3'::3’ΔHygR + 5’ΔHygR::Intron5'::Read1::AGCCAGTCTTACAAA::Read2::Intron3'::3’ΔHygR + 5’ΔHygR::Intron5'::Read1::GCACTGTTTTATGCA::Read2::Intron3'::3’ΔHygR + 5’ΔHygR::Intron5'::Read1::TACCGTTTTGAGCAT::Read2::Intron3'::3’ΔHygR + 5’ΔHygR::Intron5'::Read1::TCACTTTTTCACAGT::Read2::Intron3'::3’ΔHygR + 5’ΔHygR::Intron5'::Read1::ATTCTATCTTACACC::Read2::Intron3'::3’ΔHygR + 5’ΔHygR::Intron5'::Read1::TTTCGATTTAACTCG::Read2::Intron3'::3’ΔHygR + 5’ΔHygR::Intron5'::Read1::TCTCGCTGTGATTAA::Read2::Intron3'::3’ΔHygR + 5’ΔHygR::Intron5'::Read1::GCGCTTTGTGACGTT::Read2::Intron3'::3’ΔHygR + 5’ΔHygR::Intron5'::Read1::TGGCAGTATGACTCC::Read2::Intron3'::3’ΔHygR + 5’ΔHygR::Intron5'::Read1::CTCCTGTTTAATAAT::Read2::Intron3'::3’ΔHygR +

5’ΔHygR::Intron5'::Read1::TCGCTATCTTATTAT::Read2::Intron3'::3’ΔHygR + 5’ΔHygR::Intron5'::Read1::AAGCTCTGTAACCTC::Read2::Intron3'::3’ΔHygR + 5’ΔHygR::Intron5'::Read1::CCTCGTTCTTAGTAT::Read2::Intron3'::3’ΔHygR + 5’ΔHygR::Intron5'::Read1::AAGCTTTTTGACTCC::Read2::Intron3'::3’ΔHygR + 5’ΔHygR::Intron5'::Read1::CGTCGATCTGAACAC::Read2::Intron3'::3’ΔHygR + 5’ΔHygR::Intron5'::Read1::TCTCTCTATAATTCC::Read2::Intron3'::3’ΔHygR + 5’ΔHygR::Intron5'::Read1::CATCGATTTCATTCT::Read2::Intron3'::3’ΔHygR + 5’ΔHygR::Intron5'::Read1::CCGCCATCTTAGGTC::Read2::Intron3'::3’ΔHygR + 5’ΔHygR::Intron5'::Read1::CTCCAGTCTTACCTC::Read2::Intron3'::3’ΔHygR + 5’ΔHygR::Intron5'::Read1::TAACGGTATGAGGAT::Read2::Intron3'::3’ΔHygR + 5’ΔHygR::Intron5'::Read1::TATCTGTGTGATCTC::Read2::Intron3'::3’ΔHygR + 5’ΔHygR::Intron5'::Read1::ATTCAATCTAAGCGC::Read2::Intron3'::3’ΔHygR + 5’ΔHygR::Intron5'::Read1::CACCTCTGTCAACTT::Read2::Intron3'::3’ΔHygR + 5’ΔHygR::Intron5'::Read1::ACGCCCTTTAACCAA::Read2::Intron3'::3’ΔHygR + 5’ΔHygR::Intron5'::Read1::TCCCAATATCAGCAA::Read2::Intron3'::3’ΔHygR + 5’ΔHygR::Intron5'::Read1::ATACGGTTTCAGTGC::Read2::Intron3'::3’ΔHygR + 5’ΔHygR::Intron5'::Read1::CCACAATTTCACCAT::Read2::Intron3'::3’ΔHygR + 5’ΔHygR::Intron5'::Read1::GCTCCCTCTCAACGT::Read2::Intron3'::3’ΔHygR + 5’ΔHygR::Intron5'::Read1::TTACGGTATTAAGAC::Read2::Intron3'::3’ΔHygR + 5’ΔHygR::Intron5'::Read1::GCTCACTGTAACACC::Read2::Intron3'::3’ΔHygR + 5’ΔHygR::Intron5'::Read1::ATTCCGTATGATTCA::Read2::Intron3'::3’ΔHygR + 5’ΔHygR::Intron5'::Read1::AACCGCTTTGAGTTT::Read2::Intron3'::3’ΔHygR + 5’ΔHygR::Intron5'::Read1::AATCCCTCTAACCCC::Read2::Intron3'::3’ΔHygR + 5’ΔHygR::Intron5'::Read1::GGGCCCTGTGAAAGG::Read2::Intron3'::3’ΔHygR + 5’ΔHygR::Intron5'::Read1::TACCGTTTTGAATTT::Read2::Intron3'::3’ΔHygR + 5’ΔHygR::Intron5'::Read1::TTACTCTCTAACACG::Read2::Intron3'::3’ΔHygR + 5’ΔHygR::Intron5'::Read1::AAGCCATCTAACCTA::Read2::Intron3'::3’ΔHygR + 5’ΔHygR::Intron5'::Read1::CCACGTTCTGACCCC::Read2::Intron3'::3’ΔHygR + 5’ΔHygR::Intron5'::Read1::TTGCGCTATCATCGT::Read2::Intron3'::3’ΔHygR + 5’ΔHygR::Intron5'::Read1::GCACTCTCTGACCTA::Read2::Intron3'::3’ΔHygR + 5’ΔHygR::Intron5'::Read1::AAACTCTATAATCAA::Read2::Intron3'::3’ΔHygR + 5’ΔHygR::Intron5'::Read1::ACTCCATGTCATATC::Read2::Intron3'::3’ΔHygR + 5’ΔHygR::Intron5'::Read1::AGTCCTTTTTAGCCA::Read2::Intron3'::3’ΔHygR + 5’ΔHygR::Intron5'::Read1::CCCCTTTTTGAACTC::Read2::Intron3'::3’ΔHygR + 5’ΔHygR::Intron5'::Read1::ATCCACTATTAAGCA::Read2::Intron3'::3’ΔHygR + 5’ΔHygR::Intron5'::Read1::CTCCCTTCTGAGATC::Read2::Intron3'::3’ΔHygR + 5’ΔHygR::Intron5'::Read1::GCTCCTTCTTATAAC::Read2::Intron3'::3’ΔHygR + 5’ΔHygR::Intron5'::Read1::TGGCAATCTTAAAAT::Read2::Intron3'::3’ΔHygR + 5’ΔHygR::Intron5'::Read1::CACCTATTTTACTTA::Read2::Intron3'::3’ΔHygR + 5’ΔHygR::Intron5'::Read1::AACCTTTTTAAGTTA::Read2::Intron3'::3’ΔHygR + 5’ΔHygR::Intron5'::Read1::CGGCGTTGTAAGTGC::Read2::Intron3'::3’ΔHygR + 5’ΔHygR::Intron5'::Read1::GACCAGTTTTAGACA::Read2::Intron3'::3’ΔHygR + 5’ΔHygR::Intron5'::Read1::CGTCCTTCTCAGCGC::Read2::Intron3'::3’ΔHygR + 5’ΔHygR::Intron5'::Read1::TATCACTGTTAATCT::Read2::Intron3'::3’ΔHygR + 5’ΔHygR::Intron5'::Read1::TATCTGTTTGATGTC::Read2::Intron3'::3’ΔHygR + 5’ΔHygR::Intron5'::Read1::GATCATTCTTACATG::Read2::Intron3'::3’ΔHygR + 5’ΔHygR::Intron5'::Read1::GTCCATTTTCAATCT::Read2::Intron3'::3’ΔHygR + 5’ΔHygR::Intron5'::Read1::CTACTTTTTCAATCC::Read2::Intron3'::3’ΔHygR + 5’ΔHygR::Intron5'::Read1::GATCCATTTCACATT::Read2::Intron3'::3’ΔHygR + 5’ΔHygR::Intron5'::Read1::TTCCGTTGTTACAAT::Read2::Intron3'::3’ΔHygR + 5’ΔHygR::Intron5'::Read1::ATTCGATCTCACTTC::Read2::Intron3'::3’ΔHygR + 5’ΔHygR::Intron5'::Read1::CCCCTATGTAACCTT::Read2::Intron3'::3’ΔHygR + 5’ΔHygR::Intron5'::Read1::TACCAGTCTCAACTT::Read2::Intron3'::3’ΔHygR + 5’ΔHygR::Intron5'::Read1::GACCTGTTTAACTTC::Read2::Intron3'::3’ΔHygR + 5’ΔHygR::Intron5'::Read1::TGACTCTCTCAAGCC::Read2::Intron3'::3’ΔHygR + 5’ΔHygR::Intron5'::Read1::ACCCCGTATCACCAC::Read2::Intron3'::3’ΔHygR + 5’ΔHygR::Intron5'::Read1::GATCGGTGTCAAGCT::Read2::Intron3'::3’ΔHygR + 5’ΔHygR::Intron5'::Read1::GTTCGATGTTAACTA::Read2::Intron3'::3’ΔHygR + 5’ΔHygR::Intron5'::Read1::AAGCGATCTCATCTC::Read2::Intron3'::3’ΔHygR + 5’ΔHygR::Intron5'::Read1::GATCCATATTACACC::Read2::Intron3'::3’ΔHygR + 5’ΔHygR::Intron5'::Read1::TCACAATATCACTAA::Read2::Intron3'::3’ΔHygR + 5’ΔHygR::Intron5'::Read1::ATACCATGTTATGCA::Read2::Intron3'::3’ΔHygR + 5’ΔHygR::Intron5'::Read1::CGGCCTTCTCAATCA::Read2::Intron3'::3’ΔHygR + 5’ΔHygR::Intron5'::Read1::GTCCATTTTTATGTT::Read2::Intron3'::3’ΔHygR + 5’ΔHygR::Intron5'::Read1::GTACCCTATTACACG::Read2::Intron3'::3’ΔHygR + 5’ΔHygR::Intron5'::Read1::ACTCCATTTAAAACC::Read2::Intron3'::3’ΔHygR + 5’ΔHygR::Intron5'::Read1::CTTCGTTGTTAACTT::Read2::Intron3'::3’ΔHygR + 5’ΔHygR::Intron5'::Read1::TTTCGCTCTCACTCC::Read2::Intron3'::3’ΔHygR + 5’ΔHygR::Intron5'::Read1::ACACTCTATAACATA::Read2::Intron3'::3’ΔHygR + 5’ΔHygR::Intron5'::Read1::CTTCAATGTGATACT::Read2::Intron3'::3’ΔHygR +

5’ΔHygR::Intron5'::Read1::ATTCCTTTTAAACCG::Read2::Intron3'::3’ΔHygR + 5’ΔHygR::Intron5'::Read1::ATCCGGTGTCAATAC::Read2::Intron3'::3’ΔHygR + 5’ΔHygR::Intron5'::Read1::TCGCTGTGTTACAAC::Read2::Intron3'::3’ΔHygR + 5’ΔHygR::Intron5'::Read1::TTACAGTCTTAAACA::Read2::Intron3'::3’ΔHygR + 5’ΔHygR::Intron5'::Read1::CCTCCTTCTAACTAC::Read2::Intron3'::3’ΔHygR + 5’ΔHygR::Intron5'::Read1::AATCCCTCTAATAGC::Read2::Intron3'::3’ΔHygR + 5’ΔHygR::Intron5'::Read1::ATCCCATTTTACACA::Read2::Intron3'::3’ΔHygR + 5’ΔHygR::Intron5'::Read1::CTACCGTTTAATTCG::Read2::Intron3'::3’ΔHygR + 5’ΔHygR::Intron5'::Read1::TACCGTTTTAACGTT::Read2::Intron3'::3’ΔHygR + 5’ΔHygR::Intron5'::Read1::TCCCAGTCTCAAGCA::Read2::Intron3'::3’ΔHygR + 5’ΔHygR::Intron5'::Read1::ATGCTCTCTCAACGC::Read2::Intron3'::3’ΔHygR + 5’ΔHygR::Intron5'::Read1::CGGCCATCTCAGTAA::Read2::Intron3'::3’ΔHygR + 5’ΔHygR::Intron5'::Read1::CTTCTATCTAATTGT::Read2::Intron3'::3’ΔHygR + 5’ΔHygR::Intron5'::Read1::GCTCACTCTGAAACT::Read2::Intron3'::3’ΔHygR + 5’ΔHygR::Intron5'::Read1::TCGCGGTCTAAAGAC::Read2::Intron3'::3’ΔHygR + 5’ΔHygR::Intron5'::Read1::CACCATTTTAAACGA::Read2::Intron3'::3’ΔHygR + 5’ΔHygR::Intron5'::Read1::CCACGTTATAATGAA::Read2::Intron3'::3’ΔHygR + 5’ΔHygR::Intron5'::Read1::CCCCAGTTTAACTGA::Read2::Intron3'::3’ΔHygR + 5’ΔHygR::Intron5'::Read1::ACGCAGTTTCATGCA::Read2::Intron3'::3’ΔHygR + 5’ΔHygR::Intron5'::Read1::CTCCTTTCTCATAAG::Read2::Intron3'::3’ΔHygR + 5’ΔHygR::Intron5'::Read1::TATCCATATCATTTT::Read2::Intron3'::3’ΔHygR + 5’ΔHygR::Intron5'::Read1::ACACGTTCTCATTCT::Read2::Intron3'::3’ΔHygR + 5’ΔHygR::Intron5'::Read1::GCCCCTTCTGAACTT::Read2::Intron3'::3’ΔHygR + 5’ΔHygR::Intron5'::Read1::GTACATTTTTAAACT::Read2::Intron3'::3’ΔHygR + 5’ΔHygR::Intron5'::Read1::GTCCTTTTTTAGTCC::Read2::Intron3'::3’ΔHygR + 5’ΔHygR::Intron5'::Read1::AGCCACTCTTATCAT::Read2::Intron3'::3’ΔHygR + 5’ΔHygR::Intron5'::Read1::GCACCCTCTTACATC::Read2::Intron3'::3’ΔHygR + 5’ΔHygR::Intron5'::Read1::GGTCCCTGTTACAAT::Read2::Intron3'::3’ΔHygR + 5’ΔHygR::Intron5'::Read1::TAGCCCTTTGAAAGC::Read2::Intron3'::3’ΔHygR + 5’ΔHygR::Intron5'::Read1::GTCCTGTATAAACCT::Read2::Intron3'::3’ΔHygR + 5’ΔHygR::Intron5'::Read1::CAGCTCTATTATGCA::Read2::Intron3'::3’ΔHygR + 5’ΔHygR::Intron5'::Read1::TCCCTTTCTCAGCAC::Read2::Intron3'::3’ΔHygR + 5’ΔHygR::Intron5'::Read1::CCCCATTTTAAGCGC::Read2::Intron3'::3’ΔHygR + 5’ΔHygR::Intron5'::Read1::TTTCCGTTTTATTTT::Read2::Intron3'::3’ΔHygR + 5’ΔHygR::Intron5'::Read1::GGCCGTTCTTAAATA::Read2::Intron3'::3’ΔHygR + 5’ΔHygR::Intron5'::Read1::AACCTCTCTTACGGC::Read2::Intron3'::3’ΔHygR + 5’ΔHygR::Intron5'::Read1::CAGCGATCTTAATGC::Read2::Intron3'::3’ΔHygR + 5’ΔHygR::Intron5'::Read1::TGTCCCTCTCAACCG::Read2::Intron3'::3’ΔHygR + 5’ΔHygR::Intron5'::Read1::AGTCGCTCTTAAAGC::Read2::Intron3'::3’ΔHygR + 5’ΔHygR::Intron5'::Read1::CACCTTTTTTATTTA::Read2::Intron3'::3’ΔHygR + 5’ΔHygR::Intron5'::Read1::GTGCAATATTAGGCG::Read2::Intron3'::3’ΔHygR + 5’ΔHygR::Intron5'::Read1::TCACGTTATTACCGA::Read2::Intron3'::3’ΔHygR + 5’ΔHygR::Intron5'::Read1::GCTCCATATGACTTC::Read2::Intron3'::3’ΔHygR + 5’ΔHygR::Intron5'::Read1::TCGCTTTGTAAAAAA::Read2::Intron3'::3’ΔHygR + 5’ΔHygR::Intron5'::Read1::CTACAATGTCAACAC::Read2::Intron3'::3’ΔHygR + 5’ΔHygR::Intron5'::Read1::GACCATTCTCATCAC::Read2::Intron3'::3’ΔHygR + 5’ΔHygR::Intron5'::Read1::AAACATTATTATTGT::Read2::Intron3'::3’ΔHygR + 5’ΔHygR::Intron5'::Read1::CATCATTTTCAAGGC::Read2::Intron3'::3’ΔHygR + 5’ΔHygR::Intron5'::Read1::GGCCATTATCATACG::Read2::Intron3'::3’ΔHygR + 5’ΔHygR::Intron5'::Read1::AGCCAGTCTAACTAT::Read2::Intron3'::3’ΔHygR + 5’ΔHygR::Intron5'::Read1::AGGCTATCTCACCTG::Read2::Intron3'::3’ΔHygR + 5’ΔHygR::Intron5'::Read1::ACTCCCTTTTATGTA::Read2::Intron3'::3’ΔHygR + 5’ΔHygR::Intron5'::Read1::CCCCATTTTCAACTA::Read2::Intron3'::3’ΔHygR + 5’ΔHygR::Intron5'::Read1::AAACTGTCTGACGTG::Read2::Intron3'::3’ΔHygR + 5’ΔHygR::Intron5'::Read1::AAACTTTTTGAGTAG::Read2::Intron3'::3’ΔHygR + 5’ΔHygR::Intron5'::Read1::CATCCCTATTAGGCA::Read2::Intron3'::3’ΔHygR + 5’ΔHygR::Intron5'::Read1::GTCCACTTTTACTCG::Read2::Intron3'::3’ΔHygR + 5’ΔHygR::Intron5'::Read1::AGCCAATTTGACTAG::Read2::Intron3'::3’ΔHygR + 5’ΔHygR::Intron5'::Read1::CTCCTGTTTCATTTA::Read2::Intron3'::3’ΔHygR + 5’ΔHygR::Intron5'::Read1::CGGCAGTCTAAGCAC::Read2::Intron3'::3’ΔHygR + 5’ΔHygR::Intron5'::Read1::CTGCTTTCTTATGAA::Read2::Intron3'::3’ΔHygR + 5’ΔHygR::Intron5'::Read1::TTTCGATGTTAACGA::Read2::Intron3'::3’ΔHygR + 5’ΔHygR::Intron5'::Read1::ATTCGTTCTCAGAGC::Read2::Intron3'::3’ΔHygR + 5’ΔHygR::Intron5'::Read1::CCGCTGTATTACCTC::Read2::Intron3'::3’ΔHygR + 5’ΔHygR::Intron5'::Read1::TAACCGTCTCATTAT::Read2::Intron3'::3’ΔHygR + 5’ΔHygR::Intron5'::Read1::CCGCTATTTTACCCC::Read2::Intron3'::3’ΔHygR + 5’ΔHygR::Intron5'::Read1::CTACCCTCTTATCTC::Read2::Intron3'::3’ΔHygR + 5’ΔHygR::Intron5'::Read1::TCCCCATCTGAAGAC::Read2::Intron3'::3’ΔHygR + 5’ΔHygR::Intron5'::Read1::TTCCAGTGTCAAGAT::Read2::Intron3'::3’ΔHygR + 5’ΔHygR::Intron5'::Read1::AGTCCATCTTAGGAA::Read2::Intron3'::3’ΔHygR +

5’ΔHygR::Intron5'::Read1::AAACGTTATAACCAT::Read2::Intron3'::3’ΔHygR + 5’ΔHygR::Intron5'::Read1::AGACTCTGTCACGAT::Read2::Intron3'::3’ΔHygR + 5’ΔHygR::Intron5'::Read1::ATACATTTTTAGACC::Read2::Intron3'::3’ΔHygR + 5’ΔHygR::Intron5'::Read1::CCACCATATAAATTC::Read2::Intron3'::3’ΔHygR + 5’ΔHygR::Intron5'::Read1::GCTCAGTCTAATCTC::Read2::Intron3'::3’ΔHygR + 5’ΔHygR::Intron5'::Read1::CAACATTGTTACCAC::Read2::Intron3'::3’ΔHygR + 5’ΔHygR::Intron5'::Read1::AACCTCTTTAATTAA::Read2::Intron3'::3’ΔHygR + 5’ΔHygR::Intron5'::Read1::GTGCTATGTTATAGG::Read2::Intron3'::3’ΔHygR + 5’ΔHygR::Intron5'::Read1::CTCCGATCTTAGAGC::Read2::Intron3'::3’ΔHygR + 5’ΔHygR::Intron5'::Read1::TCGCATTGTAATACT::Read2::Intron3'::3’ΔHygR + 5’ΔHygR::Intron5'::Read1::TTCCGTTGTAACACT::Read2::Intron3'::3’ΔHygR + 5’ΔHygR::Intron5'::Read1::TATCTATGTGATACA::Read2::Intron3'::3’ΔHygR + 5’ΔHygR::Intron5'::Read1::TCACCCTATAAACTG::Read2::Intron3'::3’ΔHygR + 5’ΔHygR::Intron5'::Read1::TTCCGATCTGACGAC::Read2::Intron3'::3’ΔHygR + 5’ΔHygR::Intron5'::Read1::TAACCATCTGATCAT::Read2::Intron3'::3’ΔHygR + 5’ΔHygR::Intron5'::Read1::TATCAATATTAATCT::Read2::Intron3'::3’ΔHygR + 5’ΔHygR::Intron5'::Read1::AAACGATTTCAAGTA::Read2::Intron3'::3’ΔHygR + 5’ΔHygR::Intron5'::Read1::CACCGCTCTTAGCAA::Read2::Intron3'::3’ΔHygR + 5’ΔHygR::Intron5'::Read1::ACTCTTTCTCAGCAA::Read2::Intron3'::3’ΔHygR + 5’ΔHygR::Intron5'::Read1::TTCCGATATTAGTTA::Read2::Intron3'::3’ΔHygR + 5’ΔHygR::Intron5'::Read1::TCGCTCTATAAGATT::Read2::Intron3'::3’ΔHygR + 5’ΔHygR::Intron5'::Read1::TGACCATCTAATAGG::Read2::Intron3'::3’ΔHygR + 5’ΔHygR::Intron5'::Read1::ACTCGCTTTAATACT::Read2::Intron3'::3’ΔHygR + 5’ΔHygR::Intron5'::Read1::ATCCTATTTGAAGTA::Read2::Intron3'::3’ΔHygR + 5’ΔHygR::Intron5'::Read1::TCTCTATATAACCAA::Read2::Intron3'::3’ΔHygR + 5’ΔHygR::Intron5'::Read1::AGGCGCTCTTAATCG::Read2::Intron3'::3’ΔHygR + 5’ΔHygR::Intron5'::Read1::CGTCACTATTAGCTT::Read2::Intron3'::3’ΔHygR + 5’ΔHygR::Intron5'::Read1::GTCCCTTTTAAGAAG::Read2::Intron3'::3’ΔHygR + 5’ΔHygR::Intron5'::Read1::TGGCAATCTTATTGG::Read2::Intron3'::3’ΔHygR + 5’ΔHygR::Intron5'::Read1::GGCCAATATGATAGA::Read2::Intron3'::3’ΔHygR + 5’ΔHygR::Intron5'::Read1::CGTCCTTGTCATGAT::Read2::Intron3'::3’ΔHygR + 5’ΔHygR::Intron5'::Read1::CTCCTTTGTTATCGC::Read2::Intron3'::3’ΔHygR + 5’ΔHygR::Intron5'::Read1::GAGCTTTTTGATCTT::Read2::Intron3'::3’ΔHygR + 5’ΔHygR::Intron5'::Read1::GCACCATTTCACAGA::Read2::Intron3'::3’ΔHygR + 5’ΔHygR::Intron5'::Read1::TGTCACTCTTACCTG::Read2::Intron3'::3’ΔHygR + 5’ΔHygR::Intron5'::Read1::AACCCATGTAAACAG::Read2::Intron3'::3’ΔHygR + 5’ΔHygR::Intron5'::Read1::TGGCTATCTCACATC::Read2::Intron3'::3’ΔHygR + 5’ΔHygR::Intron5'::Read1::CCTCAGTTTTAATCA::Read2::Intron3'::3’ΔHygR + 5’ΔHygR::Intron5'::Read1::TCACATTCTTACAAC::Read2::Intron3'::3’ΔHygR + 5’ΔHygR::Intron5'::Read1::ACACGTTGTAAACAC::Read2::Intron3'::3’ΔHygR + 5’ΔHygR::Intron5'::Read1::ATGCATTCTAATACT::Read2::Intron3'::3’ΔHygR + 5’ΔHygR::Intron5'::Read1::CATCCCTCTTACCTT::Read2::Intron3'::3’ΔHygR + 5’ΔHygR::Intron5'::Read1::CCTCTATCTCACTCA::Read2::Intron3'::3’ΔHygR + 5’ΔHygR::Intron5'::Read1::TATCCATTTAATTTC::Read2::Intron3'::3’ΔHygR + 5’ΔHygR::Intron5'::Read1::TATCGGTATCACTCT::Read2::Intron3'::3’ΔHygR + 5’ΔHygR::Intron5'::Read1::TCTCGGTATTAACCC::Read2::Intron3'::3’ΔHygR + 5’ΔHygR::Intron5'::Read1::ATCCGTTTTAAGCCG::Read2::Intron3'::3’ΔHygR + 5’ΔHygR::Intron5'::Read1::AGTCCCTATCAACTA::Read2::Intron3'::3’ΔHygR + 5’ΔHygR::Intron5'::Read1::ATACAATCTCACTGC::Read2::Intron3'::3’ΔHygR + 5’ΔHygR::Intron5'::Read1::TTCCGCTATCAGTTC::Read2::Intron3'::3’ΔHygR + 5’ΔHygR::Intron5'::Read1::TTGCCCTTTCATCGT::Read2::Intron3'::3’ΔHygR + 5’ΔHygR::Intron5'::Read1::CCCCGTTCTTATTTT::Read2::Intron3'::3’ΔHygR + 5’ΔHygR::Intron5'::Read1::CGGCCTTCTCACGAT::Read2::Intron3'::3’ΔHygR + 5’ΔHygR::Intron5'::Read1::CACCACTCTTAATCA::Read2::Intron3'::3’ΔHygR + 5’ΔHygR::Intron5'::Read1::GCACTGTATTATTCT::Read2::Intron3'::3’ΔHygR + 5’ΔHygR::Intron5'::Read1::GGCCAGTATGACGCA::Read2::Intron3'::3’ΔHygR + 5’ΔHygR::Intron5'::Read1::ACGCGCTATGAGACT::Read2::Intron3'::3’ΔHygR + 5’ΔHygR::Intron5'::Read1::CCTCATTATGAGTTT::Read2::Intron3'::3’ΔHygR + 5’ΔHygR::Intron5'::Read1::CCTCTTTGTCACAAC::Read2::Intron3'::3’ΔHygR + 5’ΔHygR::Intron5'::Read1::CTTCACTATAAGACT::Read2::Intron3'::3’ΔHygR + 5’ΔHygR::Intron5'::Read1::GGTCTTTATGAGAGG::Read2::Intron3'::3’ΔHygR + 5’ΔHygR::Intron5'::Read1::AACCGTTCTCACCTC::Read2::Intron3'::3’ΔHygR + 5’ΔHygR::Intron5'::Read1::GTACTTTGTTATCTA::Read2::Intron3'::3’ΔHygR + 5’ΔHygR::Intron5'::Read1::CATCCTTTTCAAGAC::Read2::Intron3'::3’ΔHygR + 5’ΔHygR::Intron5'::Read1::GTACTGTGTCACAAT::Read2::Intron3'::3’ΔHygR + 5’ΔHygR::Intron5'::Read1::TACCTATCTTAAGTT::Read2::Intron3'::3’ΔHygR + 5’ΔHygR::Intron5'::Read1::AAACCCTATCAAGAG::Read2::Intron3'::3’ΔHygR + 5’ΔHygR::Intron5'::Read1::AATCTCTCTAAATCG::Read2::Intron3'::3’ΔHygR + 5’ΔHygR::Intron5'::Read1::GCTCTATTTTACCTC::Read2::Intron3'::3’ΔHygR + 5’ΔHygR::Intron5'::Read1::CGACCATTTTATAGT::Read2::Intron3'::3’ΔHygR +

5’ΔHygR::Intron5'::Read1::GTCCTGTTTGATGGA::Read2::Intron3'::3’ΔHygR + 5’ΔHygR::Intron5'::Read1::TTTCCGTCTAACTTT::Read2::Intron3'::3’ΔHygR + 5’ΔHygR::Intron5'::Read1::ATTCCCTTTAAGTTC::Read2::Intron3'::3’ΔHygR + 5’ΔHygR::Intron5'::Read1::CTTCCATCTTACCGC::Read2::Intron3'::3’ΔHygR + 5’ΔHygR::Intron5'::Read1::TACCAATATCAAGAC::Read2::Intron3'::3’ΔHygR + 5’ΔHygR::Intron5'::Read1::TTCCCTTTTAACCAC::Read2::Intron3'::3’ΔHygR + 5’ΔHygR::Intron5'::Read1::TTTCACTTTGACTTC::Read2::Intron3'::3’ΔHygR + 5’ΔHygR::Intron5'::Read1::ACACTATATCACAAG::Read2::Intron3'::3’ΔHygR + 5’ΔHygR::Intron5'::Read1::GTCCAATTTAACGCA::Read2::Intron3'::3’ΔHygR + 5’ΔHygR::Intron5'::Read1::TACCTTTTTTACGTT::Read2::Intron3'::3’ΔHygR + 5’ΔHygR::Intron5'::Read1::TGACTTTATAACCAC::Read2::Intron3'::3’ΔHygR + 5’ΔHygR::Intron5'::Read1::TTACACTCTCAAATC::Read2::Intron3'::3’ΔHygR + 5’ΔHygR::Intron5'::Read1::CCTCCATTTCAAGTC::Read2::Intron3'::3’ΔHygR + 5’ΔHygR::Intron5'::Read1::CGACTCTATTAAATT::Read2::Intron3'::3’ΔHygR + 5’ΔHygR::Intron5'::Read1::GACCCCTCTCACCAT::Read2::Intron3'::3’ΔHygR + 5’ΔHygR::Intron5'::Read1::TAACAATGTTATTCC::Read2::Intron3'::3’ΔHygR + 5’ΔHygR::Intron5'::Read1::CCACCCTTTTAGTCA::Read2::Intron3'::3’ΔHygR + 5’ΔHygR::Intron5'::Read1::TATCATTTTGACGAA::Read2::Intron3'::3’ΔHygR + 5’ΔHygR::Intron5'::Read1::TTGCACTTTGACATT::Read2::Intron3'::3’ΔHygR + 5’ΔHygR::Intron5'::Read1::TGGCCCTATTAGCTT::Read2::Intron3'::3’ΔHygR + 5’ΔHygR::Intron5'::Read1::CTGCTTTTTCATACC::Read2::Intron3'::3’ΔHygR + 5’ΔHygR::Intron5'::Read1::CTACAATATGACAAA::Read2::Intron3'::3’ΔHygR + 5’ΔHygR::Intron5'::Read1::CAACTCTTTCATTGG::Read2::Intron3'::3’ΔHygR + 5’ΔHygR::Intron5'::Read1::CCCCCTTTTCACAAT::Read2::Intron3'::3’ΔHygR + 5’ΔHygR::Intron5'::Read1::ACCCTGTTTTACTTC::Read2::Intron3'::3’ΔHygR + 5’ΔHygR::Intron5'::Read1::CGTCAATCTCACCAC::Read2::Intron3'::3’ΔHygR + 5’ΔHygR::Intron5'::Read1::GCACCTTGTTATAAA::Read2::Intron3'::3’ΔHygR + 5’ΔHygR::Intron5'::Read1::CGACATTATTAGCAT::Read2::Intron3'::3’ΔHygR + 5’ΔHygR::Intron5'::Read1::CTGCGGTCTCACCTC::Read2::Intron3'::3’ΔHygR + 5’ΔHygR::Intron5'::Read1::TCACCTTCTAACTCC::Read2::Intron3'::3’ΔHygR + 5’ΔHygR::Intron5'::Read1::TGTCGTTTTAAACAT::Read2::Intron3'::3’ΔHygR + 5’ΔHygR::Intron5'::Read1::ATTCCATATGACAAC::Read2::Intron3'::3’ΔHygR + 5’ΔHygR::Intron5'::Read1::ACACGGTCTCAAAGG::Read2::Intron3'::3’ΔHygR + 5’ΔHygR::Intron5'::Read1::CAGCCGTTTTATCAT::Read2::Intron3'::3’ΔHygR + 5’ΔHygR::Intron5'::Read1::CGGCTATTTGAAGCA::Read2::Intron3'::3’ΔHygR + 5’ΔHygR::Intron5'::Read1::TTCCGATTTCACTTA::Read2::Intron3'::3’ΔHygR + 5’ΔHygR::Intron5'::Read1::ATTCGATTTTAGCAA::Read2::Intron3'::3’ΔHygR + 5’ΔHygR::Intron5'::Read1::CATCACTTTAACTCC::Read2::Intron3'::3’ΔHygR + 5’ΔHygR::Intron5'::Read1::CTGCGCTTTAAACTC::Read2::Intron3'::3’ΔHygR + 5’ΔHygR::Intron5'::Read1::TAACAATTTCATCGT::Read2::Intron3'::3’ΔHygR + 5’ΔHygR::Intron5'::Read1::TCTCAGTCTCACTTC::Read2::Intron3'::3’ΔHygR + 5’ΔHygR::Intron5'::Read1::CTACTTTCTTACTTC::Read2::Intron3'::3’ΔHygR + 5’ΔHygR::Intron5'::Read1::GATCCGTCTCAAATG::Read2::Intron3'::3’ΔHygR + 5’ΔHygR::Intron5'::Read1::TGTCGTTATAAGCGA::Read2::Intron3'::3’ΔHygR + 5’ΔHygR::Intron5'::Read1::AACCTATGTGAGACC::Read2::Intron3'::3’ΔHygR + 5’ΔHygR::Intron5'::Read1::GCTCCTTGTAAACAC::Read2::Intron3'::3’ΔHygR + 5’ΔHygR::Intron5'::Read1::GCTCTTTGTTATCCG::Read2::Intron3'::3’ΔHygR + 5’ΔHygR::Intron5'::Read1::TATCGTTGTTAATCG::Read2::Intron3'::3’ΔHygR + 5’ΔHygR::Intron5'::Read1::GGCCTTTATCAAGCG::Read2::Intron3'::3’ΔHygR + 5’ΔHygR::Intron5'::Read1::CTACCTTATAATCCA::Read2::Intron3'::3’ΔHygR + 5’ΔHygR::Intron5'::Read1::CTGCTATTTAATGAC::Read2::Intron3'::3’ΔHygR + 5’ΔHygR::Intron5'::Read1::AATCTTTTTCAACTA::Read2::Intron3'::3’ΔHygR + 5’ΔHygR::Intron5'::Read1::AGTCTTTCTAAGCCA::Read2::Intron3'::3’ΔHygR + 5’ΔHygR::Intron5'::Read1::AACCCGTTTCACGAA::Read2::Intron3'::3’ΔHygR + 5’ΔHygR::Intron5'::Read1::AAGCATTATTATCCT::Read2::Intron3'::3’ΔHygR + 5’ΔHygR::Intron5'::Read1::CTCCCATATTAGGCC::Read2::Intron3'::3’ΔHygR + 5’ΔHygR::Intron5'::Read1::GCACCATTTCATCAA::Read2::Intron3'::3’ΔHygR + 5’ΔHygR::Intron5'::Read1::TAGCTGTATCAGCTC::Read2::Intron3'::3’ΔHygR + 5’ΔHygR::Intron5'::Read1::TATCTGTCTAATATC::Read2::Intron3'::3’ΔHygR + 5’ΔHygR::Intron5'::Read1::TCGCACTCTAATCCT::Read2::Intron3'::3’ΔHygR + 5’ΔHygR::Intron5'::Read1::AAGCCTTCTTAGTAC::Read2::Intron3'::3’ΔHygR + 5’ΔHygR::Intron5'::Read1::ACCCTATTTAAGATC::Read2::Intron3'::3’ΔHygR + 5’ΔHygR::Intron5'::Read1::CACCATTATGAGTAC::Read2::Intron3'::3’ΔHygR + 5’ΔHygR::Intron5'::Read1::GGTCGCTATAACCAA::Read2::Intron3'::3’ΔHygR + 5’ΔHygR::Intron5'::Read1::TGCCTCTATAAAATT::Read2::Intron3'::3’ΔHygR + 5’ΔHygR::Intron5'::Read1::ACTCCTTATCAATCC::Read2::Intron3'::3’ΔHygR + 5’ΔHygR::Intron5'::Read1::CGCCTTTGTGAAGTT::Read2::Intron3'::3’ΔHygR + 5’ΔHygR::Intron5'::Read1::CTGCTTTTTTATCTA::Read2::Intron3'::3’ΔHygR + 5’ΔHygR::Intron5'::Read1::GTCCCATGTAAGCCT::Read2::Intron3'::3’ΔHygR + 5’ΔHygR::Intron5'::Read1::TCTCACTCTAAGCGT::Read2::Intron3'::3’ΔHygR +

5’ΔHygR::Intron5'::Read1::TTACCATCTCACGAG::Read2::Intron3'::3’ΔHygR + 5’ΔHygR::Intron5'::Read1::CGACGATGTGACCCT::Read2::Intron3'::3’ΔHygR + 5’ΔHygR::Intron5'::Read1::GGACCATATTAATCA::Read2::Intron3'::3’ΔHygR + 5’ΔHygR::Intron5'::Read1::TCTCAATTTTACTAA::Read2::Intron3'::3’ΔHygR + 5’ΔHygR::Intron5'::Read1::AAACGTTGTGACTTT::Read2::Intron3'::3’ΔHygR + 5’ΔHygR::Intron5'::Read1::AATCCTTATCACCCA::Read2::Intron3'::3’ΔHygR + 5’ΔHygR::Intron5'::Read1::GCTCCATTTTAAAGT::Read2::Intron3'::3’ΔHygR + 5’ΔHygR::Intron5'::Read1::ACTCCCTATTAATTC::Read2::Intron3'::3’ΔHygR + 5’ΔHygR::Intron5'::Read1::AGCCGGTATTATCCG::Read2::Intron3'::3’ΔHygR + 5’ΔHygR::Intron5'::Read1::TATCAATTTCACATG::Read2::Intron3'::3’ΔHygR + 5’ΔHygR::Intron5'::Read1::TCACCATTTAACAGT::Read2::Intron3'::3’ΔHygR + 5’ΔHygR::Intron5'::Read1::TCCCAATATGACGTC::Read2::Intron3'::3’ΔHygR + 5’ΔHygR::Intron5'::Read1::GCACCGTTTGAGACC::Read2::Intron3'::3’ΔHygR + 5’ΔHygR::Intron5'::Read1::GCTCTTTTTCACCAC::Read2::Intron3'::3’ΔHygR + 5’ΔHygR::Intron5'::Read1::GTCCTATGTCACTCT::Read2::Intron3'::3’ΔHygR + 5’ΔHygR::Intron5'::Read1::AGGCATTTTCACGTC::Read2::Intron3'::3’ΔHygR + 5’ΔHygR::Intron5'::Read1::CACCGGTTTCATTTC::Read2::Intron3'::3’ΔHygR + 5’ΔHygR::Intron5'::Read1::CCACGTTGTTACTAT::Read2::Intron3'::3’ΔHygR + 5’ΔHygR::Intron5'::Read1::CCGCCATATCATTAC::Read2::Intron3'::3’ΔHygR + 5’ΔHygR::Intron5'::Read1::CGACGTTGTTAACTA::Read2::Intron3'::3’ΔHygR + 5’ΔHygR::Intron5'::Read1::GGTCCGTCTAACTTG::Read2::Intron3'::3’ΔHygR + 5’ΔHygR::Intron5'::Read1::CGCCCCTGTTAAACC::Read2::Intron3'::3’ΔHygR + 5’ΔHygR::Intron5'::Read1::TTACACTTTCAACAC::Read2::Intron3'::3’ΔHygR + 5’ΔHygR::Intron5'::Read1::CACCAATCTTAGCAC::Read2::Intron3'::3’ΔHygR + 5’ΔHygR::Intron5'::Read1::CATCCATCTTACACT::Read2::Intron3'::3’ΔHygR + 5’ΔHygR::Intron5'::Read1::GAACCTTCTTATTCG::Read2::Intron3'::3’ΔHygR + 5’ΔHygR::Intron5'::Read1::TCCCGCTTTAAATGT::Read2::Intron3'::3’ΔHygR + 5’ΔHygR::Intron5'::Read1::TGTCAATCTCACACG::Read2::Intron3'::3’ΔHygR + 5’ΔHygR::Intron5'::Read1::CGGCAGTTTTACTCC::Read2::Intron3'::3’ΔHygR + 5’ΔHygR::Intron5'::Read1::GTGCTATCTCAGCGT::Read2::Intron3'::3’ΔHygR + 5’ΔHygR::Intron5'::Read1::ATCCATTATAAACCC::Read2::Intron3'::3’ΔHygR + 5’ΔHygR::Intron5'::Read1::TTACATTTTGATCAC::Read2::Intron3'::3’ΔHygR + 5’ΔHygR::Intron5'::Read1::ACTCTGTCTTAACCA::Read2::Intron3'::3’ΔHygR + 5’ΔHygR::Intron5'::Read1::CCTCGATCTCATGAC::Read2::Intron3'::3’ΔHygR + 5’ΔHygR::Intron5'::Read1::CGACGGTGTCAATTT::Read2::Intron3'::3’ΔHygR + 5’ΔHygR::Intron5'::Read1::TGCCTTTATGATGGC::Read2::Intron3'::3’ΔHygR + 5’ΔHygR::Intron5'::Read1::TGTCTATCTGAATTG::Read2::Intron3'::3’ΔHygR + 5’ΔHygR::Intron5'::Read1::TTCCGTTGTCAGACA::Read2::Intron3'::3’ΔHygR + 5’ΔHygR::Intron5'::Read1::AATCACTTTCATTTC::Read2::Intron3'::3’ΔHygR + 5’ΔHygR::Intron5'::Read1::ACGCATTCTGACAGG::Read2::Intron3'::3’ΔHygR + 5’ΔHygR::Intron5'::Read1::AGACGCTGTAACACA::Read2::Intron3'::3’ΔHygR + 5’ΔHygR::Intron5'::Read1::CTACCATATAAGAGC::Read2::Intron3'::3’ΔHygR + 5’ΔHygR::Intron5'::Read1::CTTCGATTTCATCAT::Read2::Intron3'::3’ΔHygR + 5’ΔHygR::Intron5'::Read1::GCTCCGTTTCATTCT::Read2::Intron3'::3’ΔHygR + 5’ΔHygR::Intron5'::Read1::ATACCCTCTAACAGA::Read2::Intron3'::3’ΔHygR + 5’ΔHygR::Intron5'::Read1::TGCCTGTCTCACAAC::Read2::Intron3'::3’ΔHygR + 5’ΔHygR::Intron5'::Read1::ATCCAGTATCACTCC::Read2::Intron3'::3’ΔHygR + 5’ΔHygR::Intron5'::Read1::CGGCCATATCATCTA::Read2::Intron3'::3’ΔHygR + 5’ΔHygR::Intron5'::Read1::GAACCTTATCACCGA::Read2::Intron3'::3’ΔHygR + 5’ΔHygR::Intron5'::Read1::GCCCACTTTTAGTTC::Read2::Intron3'::3’ΔHygR + 5’ΔHygR::Intron5'::Read1::GTTCCCTATTAATTC::Read2::Intron3'::3’ΔHygR + 5’ΔHygR::Intron5'::Read1::AGGCCGTCTTACTTG::Read2::Intron3'::3’ΔHygR + 5’ΔHygR::Intron5'::Read1::CTCCATTCTTACTAC::Read2::Intron3'::3’ΔHygR + 5’ΔHygR::Intron5'::Read1::TGCCCTTGTCATGGA::Read2::Intron3'::3’ΔHygR + 5’ΔHygR::Intron5'::Read1::AGGCTTTCTAATAGC::Read2::Intron3'::3’ΔHygR + 5’ΔHygR::Intron5'::Read1::AGTCGATCTTACCCA::Read2::Intron3'::3’ΔHygR + 5’ΔHygR::Intron5'::Read1::ATCCTCTCTAACGTC::Read2::Intron3'::3’ΔHygR + 5’ΔHygR::Intron5'::Read1::GTGCTCTTTAACTTT::Read2::Intron3'::3’ΔHygR + 5’ΔHygR::Intron5'::Read1::TCGCCATATGAATCA::Read2::Intron3'::3’ΔHygR + 5’ΔHygR::Intron5'::Read1::AATCACTTTCATACA::Read2::Intron3'::3’ΔHygR + 5’ΔHygR::Intron5'::Read1::CGGCATTTTAACCCC::Read2::Intron3'::3’ΔHygR + 5’ΔHygR::Intron5'::Read1::GAACGATTTCAAATG::Read2::Intron3'::3’ΔHygR + 5’ΔHygR::Intron5'::Read1::GATCGGTGTTACTAT::Read2::Intron3'::3’ΔHygR + 5’ΔHygR::Intron5'::Read1::GCCCTCTATCATCTT::Read2::Intron3'::3’ΔHygR + 5’ΔHygR::Intron5'::Read1::GCTCCATGTTACGAC::Read2::Intron3'::3’ΔHygR + 5’ΔHygR::Intron5'::Read1::GTCCCCTATTAGACT::Read2::Intron3'::3’ΔHygR + 5’ΔHygR::Intron5'::Read1::TTCCTCTTTTACGCC::Read2::Intron3'::3’ΔHygR + 5’ΔHygR::Intron5'::Read1::TGCCTCTTTAAACGC::Read2::Intron3'::3’ΔHygR + 5’ΔHygR::Intron5'::Read1::AACCGCTATTACAAC::Read2::Intron3'::3’ΔHygR + 5’ΔHygR::Intron5'::Read1::CACCAATATTACCTC::Read2::Intron3'::3’ΔHygR +

5’ΔHygR::Intron5'::Read1::CGTCCATTTGAGTCT::Read2::Intron3'::3’ΔHygR + 5’ΔHygR::Intron5'::Read1::TCACCTTCTAACCTA::Read2::Intron3'::3’ΔHygR + 5’ΔHygR::Intron5'::Read1::ACGCGGTTTAATCCG::Read2::Intron3'::3’ΔHygR + 5’ΔHygR::Intron5'::Read1::ATACGTTATTAATAC::Read2::Intron3'::3’ΔHygR + 5’ΔHygR::Intron5'::Read1::TCTCATTCTAACTCA::Read2::Intron3'::3’ΔHygR + 5’ΔHygR::Intron5'::Read1::TGGCACTCTCATGCT::Read2::Intron3'::3’ΔHygR + 5’ΔHygR::Intron5'::Read1::ACGCTTTCTCATAAC::Read2::Intron3'::3’ΔHygR + 5’ΔHygR::Intron5'::Read1::GAGCAATTTTATAGT::Read2::Intron3'::3’ΔHygR + 5’ΔHygR::Intron5'::Read1::TGGCAATCTAATGGT::Read2::Intron3'::3’ΔHygR + 5’ΔHygR::Intron5'::Read1::CCACATTTTCACAGC::Read2::Intron3'::3’ΔHygR + 5’ΔHygR::Intron5'::Read1::GAGCAGTATCATCCG::Read2::Intron3'::3’ΔHygR + 5’ΔHygR::Intron5'::Read1::GGCCTTTGTTACGCA::Read2::Intron3'::3’ΔHygR + 5’ΔHygR::Intron5'::Read1::TCTCATTTTCAGGCT::Read2::Intron3'::3’ΔHygR + 5’ΔHygR::Intron5'::Read1::ATACACTGTCAACTC::Read2::Intron3'::3’ΔHygR + 5’ΔHygR::Intron5'::Read1::ATCCCATATGAGATT::Read2::Intron3'::3’ΔHygR + 5’ΔHygR::Intron5'::Read1::TCACTCTCTCACCAT::Read2::Intron3'::3’ΔHygR + 5’ΔHygR::Intron5'::Read1::AAACACTGTCATTTT::Read2::Intron3'::3’ΔHygR + 5’ΔHygR::Intron5'::Read1::ACTCATTCTGAGCCG::Read2::Intron3'::3’ΔHygR + 5’ΔHygR::Intron5'::Read1::ATACGCTATCAACAC::Read2::Intron3'::3’ΔHygR + 5’ΔHygR::Intron5'::Read1::ATGCCCTATCAGATT::Read2::Intron3'::3’ΔHygR + 5’ΔHygR::Intron5'::Read1::ATTCGTTCTCACGAG::Read2::Intron3'::3’ΔHygR + 5’ΔHygR::Intron5'::Read1::GTCCCATCTTAGACT::Read2::Intron3'::3’ΔHygR + 5’ΔHygR::Intron5'::Read1::TCGCGATATAAACTA::Read2::Intron3'::3’ΔHygR + 5’ΔHygR::Intron5'::Read1::TCCCGCTCTAAAGTA::Read2::Intron3'::3’ΔHygR + 5’ΔHygR::Intron5'::Read1::GCGCAATTTCAACAG::Read2::Intron3'::3’ΔHygR + 5’ΔHygR::Intron5'::Read1::TCTCCCTTTGACTCA::Read2::Intron3'::3’ΔHygR + 5’ΔHygR::Intron5'::Read1::AATCTATCTCATCAT::Read2::Intron3'::3’ΔHygR + 5’ΔHygR::Intron5'::Read1::CGCCACTCTGACTTA::Read2::Intron3'::3’ΔHygR + 5’ΔHygR::Intron5'::Read1::CGCCCTTCTTATCTA::Read2::Intron3'::3’ΔHygR + 5’ΔHygR::Intron5'::Read1::GCTCCATCTTATTCA::Read2::Intron3'::3’ΔHygR + 5’ΔHygR::Intron5'::Read1::TGACCCTGTCACAAA::Read2::Intron3'::3’ΔHygR + 5’ΔHygR::Intron5'::Read1::TTACATTTTTAGGCT::Read2::Intron3'::3’ΔHygR + 5’ΔHygR::Intron5'::Read1::ACACCATTTCACACA::Read2::Intron3'::3’ΔHygR + 5’ΔHygR::Intron5'::Read1::ACTCATTGTTACCCT::Read2::Intron3'::3’ΔHygR + 5’ΔHygR::Intron5'::Read1::ATTCCATTTTAACGT::Read2::Intron3'::3’ΔHygR + 5’ΔHygR::Intron5'::Read1::AGCCCCTTTGAACGT::Read2::Intron3'::3’ΔHygR + 5’ΔHygR::Intron5'::Read1::GGGCAGTGTAATCAA::Read2::Intron3'::3’ΔHygR + 5’ΔHygR::Intron5'::Read1::GTACTTTTTTATTCA::Read2::Intron3'::3’ΔHygR + 5’ΔHygR::Intron5'::Read1::TATCAATTTGACTTA::Read2::Intron3'::3’ΔHygR + 5’ΔHygR::Intron5'::Read1::TTCCCTTTTAACCCC::Read2::Intron3'::3’ΔHygR + 5’ΔHygR::Intron5'::Read1::ACGCCCTCTTAAGCC::Read2::Intron3'::3’ΔHygR + 5’ΔHygR::Intron5'::Read1::ACTCGATCTCAAACC::Read2::Intron3'::3’ΔHygR + 5’ΔHygR::Intron5'::Read1::CACCGTTCTTAACTG::Read2::Intron3'::3’ΔHygR + 5’ΔHygR::Intron5'::Read1::CATCAATATGAGTTC::Read2::Intron3'::3’ΔHygR + 5’ΔHygR::Intron5'::Read1::CCCCAGTTTAATTTA::Read2::Intron3'::3’ΔHygR + 5’ΔHygR::Intron5'::Read1::CTTCAGTATAAGTTT::Read2::Intron3'::3’ΔHygR + 5’ΔHygR::Intron5'::Read1::GATCCTTCTTACAGC::Read2::Intron3'::3’ΔHygR + 5’ΔHygR::Intron5'::Read1::TTCCACTATCAAGTC::Read2::Intron3'::3’ΔHygR + 5’ΔHygR::Intron5'::Read1::GGACGTTCTGAGATA::Read2::Intron3'::3’ΔHygR + 5’ΔHygR::Intron5'::Read1::TAACTATATCATACG::Read2::Intron3'::3’ΔHygR + 5’ΔHygR::Intron5'::Read1::CGCCTTTATCAGCAA::Read2::Intron3'::3’ΔHygR + 5’ΔHygR::Intron5'::Read1::TACCCTTTTGATTAC::Read2::Intron3'::3’ΔHygR + 5’ΔHygR::Intron5'::Read1::TATCCGTCTAATACT::Read2::Intron3'::3’ΔHygR + 5’ΔHygR::Intron5'::Read1::TCGCAGTTTGAGATA::Read2::Intron3'::3’ΔHygR + 5’ΔHygR::Intron5'::Read1::TGCCCGTGTTATTAA::Read2::Intron3'::3’ΔHygR + 5’ΔHygR::Intron5'::Read1::TGGCTGTATTAAATT::Read2::Intron3'::3’ΔHygR + 5’ΔHygR::Intron5'::Read1::TTCCGGTCTGATACC::Read2::Intron3'::3’ΔHygR + 5’ΔHygR::Intron5'::Read1::TAACCCTATAAAAGC::Read2::Intron3'::3’ΔHygR + 5’ΔHygR::Intron5'::Read1::TCCCTTTTTAATACT::Read2::Intron3'::3’ΔHygR + 5’ΔHygR::Intron5'::Read1::GCCCGCTATGACTTC::Read2::Intron3'::3’ΔHygR + 5’ΔHygR::Intron5'::Read1::GTTCGATGTAAGGAA::Read2::Intron3'::3’ΔHygR + 5’ΔHygR::Intron5'::Read1::TCTCATTTTAACCCC::Read2::Intron3'::3’ΔHygR + 5’ΔHygR::Intron5'::Read1::TTTCCTTCTGATCAC::Read2::Intron3'::3’ΔHygR + 5’ΔHygR::Intron5'::Read1::CCACGATCTAATATT::Read2::Intron3'::3’ΔHygR + 5’ΔHygR::Intron5'::Read1::TCGCAATATCAATGG::Read2::Intron3'::3’ΔHygR + 5’ΔHygR::Intron5'::Read1::TTCCCGTCTAAAGTC::Read2::Intron3'::3’ΔHygR + 5’ΔHygR::Intron5'::Read1::TTCCGCTATTAGTCC::Read2::Intron3'::3’ΔHygR + 5’ΔHygR::Intron5'::Read1::TTCCTATCTAACCAC::Read2::Intron3'::3’ΔHygR + 5’ΔHygR::Intron5'::Read1::AATCCTTATCATGCA::Read2::Intron3'::3’ΔHygR + 5’ΔHygR::Intron5'::Read1::AATCTCTATTACTTT::Read2::Intron3'::3’ΔHygR +

5’ΔHygR::Intron5'::Read1::CAACCCTTTAACTTC::Read2::Intron3'::3’ΔHygR + 5’ΔHygR::Intron5'::Read1::CCTCTGTCTAATCGA::Read2::Intron3'::3’ΔHygR + 5’ΔHygR::Intron5'::Read1::TAACTTTATCATTAT::Read2::Intron3'::3’ΔHygR + 5’ΔHygR::Intron5'::Read1::TCTCCTTATTAAGCC::Read2::Intron3'::3’ΔHygR + 5’ΔHygR::Intron5'::Read1::CTACGCTTTAACTAC::Read2::Intron3'::3’ΔHygR + 5’ΔHygR::Intron5'::Read1::GAGCAATGTAACGGA::Read2::Intron3'::3’ΔHygR + 5’ΔHygR::Intron5'::Read1::TAACTTTATAAAGGC::Read2::Intron3'::3’ΔHygR + 5’ΔHygR::Intron5'::Read1::TACCGATATCACACG::Read2::Intron3'::3’ΔHygR + 5’ΔHygR::Intron5'::Read1::TCCCTATGTTATCGC::Read2::Intron3'::3’ΔHygR + 5’ΔHygR::Intron5'::Read1::CTCCGTTCTGAACAA::Read2::Intron3'::3’ΔHygR + 5’ΔHygR::Intron5'::Read1::CTCCAATCTGATATA::Read2::Intron3'::3’ΔHygR + 5’ΔHygR::Intron5'::Read1::GCCCACTGTCAGTCA::Read2::Intron3'::3’ΔHygR + 5’ΔHygR::Intron5'::Read1::TGTCTTTCTTAAGTT::Read2::Intron3'::3’ΔHygR + 5’ΔHygR::Intron5'::Read1::GGCCTATATAACTGA::Read2::Intron3'::3’ΔHygR + 5’ΔHygR::Intron5'::Read1::CTACAATCTTATGTT::Read2::Intron3'::3’ΔHygR + 5’ΔHygR::Intron5'::Read1::GGACCATATGATTCA::Read2::Intron3'::3’ΔHygR + 5’ΔHygR::Intron5'::Read1::GGGCGATCTCATCAC::Read2::Intron3'::3’ΔHygR + 5’ΔHygR::Intron5'::Read1::TCCCCTTGTAACGAT::Read2::Intron3'::3’ΔHygR + 5’ΔHygR::Intron5'::Read1::AGCCTGTGTAAATCA::Read2::Intron3'::3’ΔHygR + 5’ΔHygR::Intron5'::Read1::ACACGTTATGACTCA::Read2::Intron3'::3’ΔHygR + 5’ΔHygR::Intron5'::Read1::CAGCTATATCACTCA::Read2::Intron3'::3’ΔHygR + 5’ΔHygR::Intron5'::Read1::CATCACTCTAATTCA::Read2::Intron3'::3’ΔHygR + 5’ΔHygR::Intron5'::Read1::CTTCGGTTTGAAGCT::Read2::Intron3'::3’ΔHygR + 5’ΔHygR::Intron5'::Read1::GCCCTTTCTCATATA::Read2::Intron3'::3’ΔHygR + 5’ΔHygR::Intron5'::Read1::TCACATTCTTATAAG::Read2::Intron3'::3’ΔHygR + 5’ΔHygR::Intron5'::Read1::TTGCGGTGTTAATTA::Read2::Intron3'::3’ΔHygR + 5’ΔHygR::Intron5'::Read1::ACTCTATGTCAATCG::Read2::Intron3'::3’ΔHygR + 5’ΔHygR::Intron5'::Read1::CATCAGTCTGAAAGA::Read2::Intron3'::3’ΔHygR + 5’ΔHygR::Intron5'::Read1::GTACTTTTTCAACTT::Read2::Intron3'::3’ΔHygR + 5’ΔHygR::Intron5'::Read1::TCTCGTTTTAAACAT::Read2::Intron3'::3’ΔHygR + 5’ΔHygR::Intron5'::Read1::TTACAGTGTTAATGC::Read2::Intron3'::3’ΔHygR + 5’ΔHygR::Intron5'::Read1::AAACGCTGTAACAAC::Read2::Intron3'::3’ΔHygR + 5’ΔHygR::Intron5'::Read1::ACTCTTTTTAACACC::Read2::Intron3'::3’ΔHygR + 5’ΔHygR::Intron5'::Read1::TCCCCCTATTAGGCA::Read2::Intron3'::3’ΔHygR + 5’ΔHygR::Intron5'::Read1::TTACTCTATCACTGC::Read2::Intron3'::3’ΔHygR + 5’ΔHygR::Intron5'::Read1::ACTCAGTATGAAGGC::Read2::Intron3'::3’ΔHygR + 5’ΔHygR::Intron5'::Read1::CCCCAGTATCATTTC::Read2::Intron3'::3’ΔHygR + 5’ΔHygR::Intron5'::Read1::GGCCGGTTTCAAAAA::Read2::Intron3'::3’ΔHygR + 5’ΔHygR::Intron5'::Read1::GTTCTGTTTAAACCT::Read2::Intron3'::3’ΔHygR + 5’ΔHygR::Intron5'::Read1::TCACCGTTTCAACAG::Read2::Intron3'::3’ΔHygR + 5’ΔHygR::Intron5'::Read1::ACTCCGTATCAACCC::Read2::Intron3'::3’ΔHygR + 5’ΔHygR::Intron5'::Read1::CTACAATATTACGTC::Read2::Intron3'::3’ΔHygR + 5’ΔHygR::Intron5'::Read1::GTCCTCTCTCATCTG::Read2::Intron3'::3’ΔHygR + 5’ΔHygR::Intron5'::Read1::TTGCTTTCTCATTCA::Read2::Intron3'::3’ΔHygR + 5’ΔHygR::Intron5'::Read1::TTTCCTTATTAGCAC::Read2::Intron3'::3’ΔHygR + 5’ΔHygR::Intron5'::Read1::GAACTATCTTATTCG::Read2::Intron3'::3’ΔHygR + 5’ΔHygR::Intron5'::Read1::TGGCCCTATTATCGA::Read2::Intron3'::3’ΔHygR + 5’ΔHygR::Intron5'::Read1::ACTCGGTCTCATTTA::Read2::Intron3'::3’ΔHygR + 5’ΔHygR::Intron5'::Read1::AGTCAATATTAAATG::Read2::Intron3'::3’ΔHygR + 5’ΔHygR::Intron5'::Read1::ATCCCGTTTAACGAA::Read2::Intron3'::3’ΔHygR + 5’ΔHygR::Intron5'::Read1::GATCCTTCTCAAAAC::Read2::Intron3'::3’ΔHygR + 5’ΔHygR::Intron5'::Read1::GGCCTTTGTCAGTCT::Read2::Intron3'::3’ΔHygR + 5’ΔHygR::Intron5'::Read1::GGTCACTGTCAAAGA::Read2::Intron3'::3’ΔHygR + 5’ΔHygR::Intron5'::Read1::TTACTCTGTGACCGC::Read2::Intron3'::3’ΔHygR + 5’ΔHygR::Intron5'::Read1::AACCTCTATCAACAT::Read2::Intron3'::3’ΔHygR + 5’ΔHygR::Intron5'::Read1::ATCCGTTCTCAGCCT::Read2::Intron3'::3’ΔHygR + 5’ΔHygR::Intron5'::Read1::CTCCTATGTGAGTTG::Read2::Intron3'::3’ΔHygR + 5’ΔHygR::Intron5'::Read1::TCACGGTCTAAATAT::Read2::Intron3'::3’ΔHygR + 5’ΔHygR::Intron5'::Read1::TGCCTTTTTTAACAT::Read2::Intron3'::3’ΔHygR + 5’ΔHygR::Intron5'::Read1::AAACTATTTAAGCCG::Read2::Intron3'::3’ΔHygR + 5’ΔHygR::Intron5'::Read1::AGTCAGTCTCACCGA::Read2::Intron3'::3’ΔHygR + 5’ΔHygR::Intron5'::Read1::ATACAATCTCATCTT::Read2::Intron3'::3’ΔHygR + 5’ΔHygR::Intron5'::Read1::CATCGTTGTGAGACG::Read2::Intron3'::3’ΔHygR + 5’ΔHygR::Intron5'::Read1::CATCTCTTTCACCCG::Read2::Intron3'::3’ΔHygR + 5’ΔHygR::Intron5'::Read1::CCCCAGTCTAAGTCC::Read2::Intron3'::3’ΔHygR + 5’ΔHygR::Intron5'::Read1::CCCCCTTATTAATTC::Read2::Intron3'::3’ΔHygR + 5’ΔHygR::Intron5'::Read1::CGCCGGTCTTAATCG::Read2::Intron3'::3’ΔHygR + 5’ΔHygR::Intron5'::Read1::TAACAATATGACTAC::Read2::Intron3'::3’ΔHygR + 5’ΔHygR::Intron5'::Read1::AAACTATTTTATCTT::Read2::Intron3'::3’ΔHygR + 5’ΔHygR::Intron5'::Read1::ATACTTTTTCATCCC::Read2::Intron3'::3’ΔHygR +

5’ΔHygR::Intron5'::Read1::ATCCTGTATCAATCT::Read2::Intron3'::3’ΔHygR + 5’ΔHygR::Intron5'::Read1::ATTCTGTTTTAAGAT::Read2::Intron3'::3’ΔHygR + 5’ΔHygR::Intron5'::Read1::CCTCTATTTTACAAC::Read2::Intron3'::3’ΔHygR + 5’ΔHygR::Intron5'::Read1::CGACTTTTTAAGATA::Read2::Intron3'::3’ΔHygR + 5’ΔHygR::Intron5'::Read1::GATCCTTATGAGTAC::Read2::Intron3'::3’ΔHygR + 5’ΔHygR::Intron5'::Read1::TATCCATCTAAACCC::Read2::Intron3'::3’ΔHygR + 5’ΔHygR::Intron5'::Read1::TCGCTCTTTAATAGG::Read2::Intron3'::3’ΔHygR + 5’ΔHygR::Intron5'::Read1::TGGCGTTGTGAACCG::Read2::Intron3'::3’ΔHygR + 5’ΔHygR::Intron5'::Read1::TGGCTTTATGACCGT::Read2::Intron3'::3’ΔHygR + 5’ΔHygR::Intron5'::Read1::TTCCAATTTCAGCCA::Read2::Intron3'::3’ΔHygR + 5’ΔHygR::Intron5'::Read1::AGCCGTTATAATTAC::Read2::Intron3'::3’ΔHygR + 5’ΔHygR::Intron5'::Read1::CTTCCATCTCAAGTT::Read2::Intron3'::3’ΔHygR + 5’ΔHygR::Intron5'::Read1::AAGCACTTTTACCTC::Read2::Intron3'::3’ΔHygR + 5’ΔHygR::Intron5'::Read1::AATCACTGTGACATA::Read2::Intron3'::3’ΔHygR + 5’ΔHygR::Intron5'::Read1::AGGCGGTGTAAGTTA::Read2::Intron3'::3’ΔHygR + 5’ΔHygR::Intron5'::Read1::GACCTCTCTCAACCA::Read2::Intron3'::3’ΔHygR + 5’ΔHygR::Intron5'::Read1::GATCGTTTTCACAGC::Read2::Intron3'::3’ΔHygR + 5’ΔHygR::Intron5'::Read1::TTTCTCTCTCAACCG::Read2::Intron3'::3’ΔHygR + 5’ΔHygR::Intron5'::Read1::CACCAGTATCACTCT::Read2::Intron3'::3’ΔHygR + 5’ΔHygR::Intron5'::Read1::CATCAGTCTCAGTTT::Read2::Intron3'::3’ΔHygR + 5’ΔHygR::Intron5'::Read1::CCACTCTATGACCAA::Read2::Intron3'::3’ΔHygR + 5’ΔHygR::Intron5'::Read1::CCACTTTCTCACCTT::Read2::Intron3'::3’ΔHygR + 5’ΔHygR::Intron5'::Read1::CTTCACTATTACCAT::Read2::Intron3'::3’ΔHygR + 5’ΔHygR::Intron5'::Read1::GAACGTTCTCATTAT::Read2::Intron3'::3’ΔHygR + 5’ΔHygR::Intron5'::Read1::AGTCCGTCTAAGCAT::Read2::Intron3'::3’ΔHygR + 5’ΔHygR::Intron5'::Read1::ATACTCTGTAATTAC::Read2::Intron3'::3’ΔHygR + 5’ΔHygR::Intron5'::Read1::CCTCTCTCTGAAACT::Read2::Intron3'::3’ΔHygR + 5’ΔHygR::Intron5'::Read1::GCCCAATTTTAACAG::Read2::Intron3'::3’ΔHygR + 5’ΔHygR::Intron5'::Read1::AATCATTTTGAACAC::Read2::Intron3'::3’ΔHygR + 5’ΔHygR::Intron5'::Read1::CGGCAATCTTAGCCT::Read2::Intron3'::3’ΔHygR + 5’ΔHygR::Intron5'::Read1::TATCACTGTAAACCA::Read2::Intron3'::3’ΔHygR + 5’ΔHygR::Intron5'::Read1::AGTCATTTTCAATTC::Read2::Intron3'::3’ΔHygR + 5’ΔHygR::Intron5'::Read1::GACCTTTCTTAATCC::Read2::Intron3'::3’ΔHygR + 5’ΔHygR::Intron5'::Read1::GCACCTTTTAAACAT::Read2::Intron3'::3’ΔHygR + 5’ΔHygR::Intron5'::Read1::AGACTTTTTAACCAT::Read2::Intron3'::3’ΔHygR + 5’ΔHygR::Intron5'::Read1::AGGCCATTTCATTTC::Read2::Intron3'::3’ΔHygR + 5’ΔHygR::Intron5'::Read1::CCGCTGTGTCAACTA::Read2::Intron3'::3’ΔHygR + 5’ΔHygR::Intron5'::Read1::TTTCCATCTGAAAAC::Read2::Intron3'::3’ΔHygR + 5’ΔHygR::Intron5'::Read1::ACTCTTTATCAGTCA::Read2::Intron3'::3’ΔHygR + 5’ΔHygR::Intron5'::Read1::AGACCATCTAACTCT::Read2::Intron3'::3’ΔHygR + 5’ΔHygR::Intron5'::Read1::CATCATTCTAACTCG::Read2::Intron3'::3’ΔHygR + 5’ΔHygR::Intron5'::Read1::CCCCGTTGTTACATC::Read2::Intron3'::3’ΔHygR + 5’ΔHygR::Intron5'::Read1::CTTCCTTGTAACACA::Read2::Intron3'::3’ΔHygR + 5’ΔHygR::Intron5'::Read1::GAGCGATTTCATCAA::Read2::Intron3'::3’ΔHygR + 5’ΔHygR::Intron5'::Read1::GGGCATTGTCATTCT::Read2::Intron3'::3’ΔHygR + 5’ΔHygR::Intron5'::Read1::GAACATTTTCAAGTC::Read2::Intron3'::3’ΔHygR + 5’ΔHygR::Intron5'::Read1::TACCGATCTGACATC::Read2::Intron3'::3’ΔHygR + 5’ΔHygR::Intron5'::Read1::TGACTATCTTAGCAC::Read2::Intron3'::3’ΔHygR + 5’ΔHygR::Intron5'::Read1::AACCATTCTTAACTC::Read2::Intron3'::3’ΔHygR + 5’ΔHygR::Intron5'::Read1::GGGCCATGTTAAAAC::Read2::Intron3'::3’ΔHygR + 5’ΔHygR::Intron5'::Read1::TGTCATTTTCAGTGT::Read2::Intron3'::3’ΔHygR + 5’ΔHygR::Intron5'::Read1::CATCGCTCTAATACC::Read2::Intron3'::3’ΔHygR + 5’ΔHygR::Intron5'::Read1::CGCCCTTCTAATCTG::Read2::Intron3'::3’ΔHygR + 5’ΔHygR::Intron5'::Read1::GAGCCCTCTAATAAT::Read2::Intron3'::3’ΔHygR + 5’ΔHygR::Intron5'::Read1::GTGCAGTATCAAAAA::Read2::Intron3'::3’ΔHygR + 5’ΔHygR::Intron5'::Read1::TTTCGCTGTTATAGT::Read2::Intron3'::3’ΔHygR + 5’ΔHygR::Intron5'::Read1::CAGCTCTGTAATCCC::Read2::Intron3'::3’ΔHygR + 5’ΔHygR::Intron5'::Read1::GATCACTTTCAATGC::Read2::Intron3'::3’ΔHygR + 5’ΔHygR::Intron5'::Read1::GGGCTCTGTTACCTG::Read2::Intron3'::3’ΔHygR + 5’ΔHygR::Intron5'::Read1::TCACCCTTTAATGTT::Read2::Intron3'::3’ΔHygR + 5’ΔHygR::Intron5'::Read1::TTGCGGTCTTACGTT::Read2::Intron3'::3’ΔHygR + 5’ΔHygR::Intron5'::Read1::AAGCAATCTGAATTG::Read2::Intron3'::3’ΔHygR + 5’ΔHygR::Intron5'::Read1::CCACCCTGTTATTTC::Read2::Intron3'::3’ΔHygR + 5’ΔHygR::Intron5'::Read1::GATCCGTCTAAGGGC::Read2::Intron3'::3’ΔHygR + 5’ΔHygR::Intron5'::Read1::TTACCGTTTCACCTG::Read2::Intron3'::3’ΔHygR + 5’ΔHygR::Intron5'::Read1::TTTCCTTTTCAACAC::Read2::Intron3'::3’ΔHygR + 5’ΔHygR::Intron5'::Read1::CGACCATCTCACGAC::Read2::Intron3'::3’ΔHygR + 5’ΔHygR::Intron5'::Read1::TGACCATGTTACTCG::Read2::Intron3'::3’ΔHygR + 5’ΔHygR::Intron5'::Read1::ATGCCATTTAACTCA::Read2::Intron3'::3’ΔHygR + 5’ΔHygR::Intron5'::Read1::ATTCAATATCATTCG::Read2::Intron3'::3’ΔHygR +

5’ΔHygR::Intron5'::Read1::CCTCCTTATCAGCTC::Read2::Intron3'::3’ΔHygR + 5’ΔHygR::Intron5'::Read1::ATTCCGTCTAAATGC::Read2::Intron3'::3’ΔHygR + 5’ΔHygR::Intron5'::Read1::CCTCTTTATAATCAT::Read2::Intron3'::3’ΔHygR + 5’ΔHygR::Intron5'::Read1::CGTCTATATGATCAC::Read2::Intron3'::3’ΔHygR + 5’ΔHygR::Intron5'::Read1::GAACTGTCTCAAACA::Read2::Intron3'::3’ΔHygR + 5’ΔHygR::Intron5'::Read1::TCACCCTATAACCCA::Read2::Intron3'::3’ΔHygR + 5’ΔHygR::Intron5'::Read1::ACGCCATATCAACAT::Read2::Intron3'::3’ΔHygR + 5’ΔHygR::Intron5'::Read1::ATTCTATCTAAGCGT::Read2::Intron3'::3’ΔHygR + 5’ΔHygR::Intron5'::Read1::CGGCTCTTTGAATCT::Read2::Intron3'::3’ΔHygR + 5’ΔHygR::Intron5'::Read1::CTACATTGTCATCAC::Read2::Intron3'::3’ΔHygR + 5’ΔHygR::Intron5'::Read1::TTCCGTTATAAACCT::Read2::Intron3'::3’ΔHygR + 5’ΔHygR::Intron5'::Read1::TTTCCGTGTTACAGT::Read2::Intron3'::3’ΔHygR + 5’ΔHygR::Intron5'::Read1::AGCCTTTCTCAAACC::Read2::Intron3'::3’ΔHygR + 5’ΔHygR::Intron5'::Read1::CTACGGTATAACTTT::Read2::Intron3'::3’ΔHygR + 5’ΔHygR::Intron5'::Read1::TACCATTTTAACACT::Read2::Intron3'::3’ΔHygR + 5’ΔHygR::Intron5'::Read1::ACCCTATTTAAATCT::Read2::Intron3'::3’ΔHygR + 5’ΔHygR::Intron5'::Read1::CCGCTTTCTAATTCT::Read2::Intron3'::3’ΔHygR + 5’ΔHygR::Intron5'::Read1::TCCCTCTATTACTAT::Read2::Intron3'::3’ΔHygR + 5’ΔHygR::Intron5'::Read1::TGTCGGTCTTAGGCC::Read2::Intron3'::3’ΔHygR + 5’ΔHygR::Intron5'::Read1::AATCTATTTGACGAT::Read2::Intron3'::3’ΔHygR + 5’ΔHygR::Intron5'::Read1::CAACACTTTAAATAC::Read2::Intron3'::3’ΔHygR + 5’ΔHygR::Intron5'::Read1::GCGCAGTATAAGTCT::Read2::Intron3'::3’ΔHygR + 5’ΔHygR::Intron5'::Read1::TATCAATTTGATAGT::Read2::Intron3'::3’ΔHygR + 5’ΔHygR::Intron5'::Read1::TCACCGTCTCATCTA::Read2::Intron3'::3’ΔHygR + 5’ΔHygR::Intron5'::Read1::ATTCGTTCTTATCAA::Read2::Intron3'::3’ΔHygR + 5’ΔHygR::Intron5'::Read1::GATCCGTCTGATGTC::Read2::Intron3'::3’ΔHygR + 5’ΔHygR::Intron5'::Read1::GCACCTTTTAAGAAA::Read2::Intron3'::3’ΔHygR + 5’ΔHygR::Intron5'::Read1::AATCCTTATGAACTA::Read2::Intron3'::3’ΔHygR + 5’ΔHygR::Intron5'::Read1::ATACGTTTTAAATAT::Read2::Intron3'::3’ΔHygR + 5’ΔHygR::Intron5'::Read1::CATCCCTATCATAGT::Read2::Intron3'::3’ΔHygR + 5’ΔHygR::Intron5'::Read1::CATCCGTTTCAGGCA::Read2::Intron3'::3’ΔHygR + 5’ΔHygR::Intron5'::Read1::GACCCATTTTACTAG::Read2::Intron3'::3’ΔHygR + 5’ΔHygR::Intron5'::Read1::TTACCCTATAATTCC::Read2::Intron3'::3’ΔHygR + 5’ΔHygR::Intron5'::Read1::ACACAATCTAATCAA::Read2::Intron3'::3’ΔHygR + 5’ΔHygR::Intron5'::Read1::ACCCGCTCTCAAATC::Read2::Intron3'::3’ΔHygR + 5’ΔHygR::Intron5'::Read1::ATGCGTTCTCAGTAC::Read2::Intron3'::3’ΔHygR + 5’ΔHygR::Intron5'::Read1::CCTCATTGTTATTAT::Read2::Intron3'::3’ΔHygR + 5’ΔHygR::Intron5'::Read1::CGCCTGTTTCAACCT::Read2::Intron3'::3’ΔHygR + 5’ΔHygR::Intron5'::Read1::GGCCCTTGTCATGCG::Read2::Intron3'::3’ΔHygR + 5’ΔHygR::Intron5'::Read1::TACCTTTGTGATGCG::Read2::Intron3'::3’ΔHygR + 5’ΔHygR::Intron5'::Read1::TGCCATTATTACACA::Read2::Intron3'::3’ΔHygR + 5’ΔHygR::Intron5'::Read1::ACCCACTGTGATGAG::Read2::Intron3'::3’ΔHygR + 5’ΔHygR::Intron5'::Read1::CAACACTGTTACTAA::Read2::Intron3'::3’ΔHygR + 5’ΔHygR::Intron5'::Read1::GCACGTTGTCAATCA::Read2::Intron3'::3’ΔHygR + 5’ΔHygR::Intron5'::Read1::TCGCGTTCTTAGGCA::Read2::Intron3'::3’ΔHygR + 5’ΔHygR::Intron5'::Read1::TGTCGATGTTATGTC::Read2::Intron3'::3’ΔHygR + 5’ΔHygR::Intron5'::Read1::CATCACTGTCAACTG::Read2::Intron3'::3’ΔHygR + 5’ΔHygR::Intron5'::Read1::CCACAGTGTTATCGA::Read2::Intron3'::3’ΔHygR + 5’ΔHygR::Intron5'::Read1::GCTCTCTGTCACCTA::Read2::Intron3'::3’ΔHygR + 5’ΔHygR::Intron5'::Read1::TAACCCTTTTAGCAA::Read2::Intron3'::3’ΔHygR + 5’ΔHygR::Intron5'::Read1::TTACCCTATGAGCTA::Read2::Intron3'::3’ΔHygR + 5’ΔHygR::Intron5'::Read1::AATCCATCTAATACC::Read2::Intron3'::3’ΔHygR + 5’ΔHygR::Intron5'::Read1::ATTCTTTGTGATCTT::Read2::Intron3'::3’ΔHygR + 5’ΔHygR::Intron5'::Read1::CATCCCTTTCAGTCT::Read2::Intron3'::3’ΔHygR + 5’ΔHygR::Intron5'::Read1::CGGCGATCTGATTCC::Read2::Intron3'::3’ΔHygR + 5’ΔHygR::Intron5'::Read1::GCGCCATGTGAATAT::Read2::Intron3'::3’ΔHygR + 5’ΔHygR::Intron5'::Read1::AAACGTTATTATATC::Read2::Intron3'::3’ΔHygR + 5’ΔHygR::Intron5'::Read1::CCACGTTATCACATT::Read2::Intron3'::3’ΔHygR + 5’ΔHygR::Intron5'::Read1::GAACATTATAATCAA::Read2::Intron3'::3’ΔHygR + 5’ΔHygR::Intron5'::Read1::GACCACTGTAAACTC::Read2::Intron3'::3’ΔHygR + 5’ΔHygR::Intron5'::Read1::AGTCTATATCAATAC::Read2::Intron3'::3’ΔHygR + 5’ΔHygR::Intron5'::Read1::AATCGCTATTAAGGC::Read2::Intron3'::3’ΔHygR + 5’ΔHygR::Intron5'::Read1::GACCCTTGTTAAGAC::Read2::Intron3'::3’ΔHygR + 5’ΔHygR::Intron5'::Read1::GTCCTATATCATTAT::Read2::Intron3'::3’ΔHygR + 5’ΔHygR::Intron5'::Read1::TACCCCTTTGATAGT::Read2::Intron3'::3’ΔHygR + 5’ΔHygR::Intron5'::Read1::ACGCCCTCTCAACGT::Read2::Intron3'::3’ΔHygR + 5’ΔHygR::Intron5'::Read1::CGACAATATGACTAG::Read2::Intron3'::3’ΔHygR + 5’ΔHygR::Intron5'::Read1::GATTTAATAGAGTAG::Read2::Intron3'::3’ΔHygR + 5’ΔHygR::Intron5'::Read1::GTCCCCTCTAATCTA::Read2::Intron3'::3’ΔHygR + 5’ΔHygR::Intron5'::Read1::TCTCCATATCAATGC::Read2::Intron3'::3’ΔHygR +

5’ΔHygR::Intron5'::Read1::ACACCCTATGAAGCC::Read2::Intron3'::3’ΔHygR + 5’ΔHygR::Intron5'::Read1::ACGCCTTGTAACGCA::Read2::Intron3'::3’ΔHygR + 5’ΔHygR::Intron5'::Read1::CCACTCTATCAGCAT::Read2::Intron3'::3’ΔHygR + 5’ΔHygR::Intron5'::Read1::CTACTATATCACTAA::Read2::Intron3'::3’ΔHygR + 5’ΔHygR::Intron5'::Read1::CTCCCGTCTGAGTCC::Read2::Intron3'::3’ΔHygR + 5’ΔHygR::Intron5'::Read1::GTGCCTTTTCACCGT::Read2::Intron3'::3’ΔHygR + 5’ΔHygR::Intron5'::Read1::TATCCGTATTAACCC::Read2::Intron3'::3’ΔHygR + 5’ΔHygR::Intron5'::Read1::TGCCCTTATTAATTC::Read2::Intron3'::3’ΔHygR + 5’ΔHygR::Intron5'::Read1::TGTCCTTCTCAATAA::Read2::Intron3'::3’ΔHygR + 5’ΔHygR::Intron5'::Read1::TGTCTTTTTCATTTC::Read2::Intron3'::3’ΔHygR + 5’ΔHygR::Intron5'::Read1::AAACCGTATGATCAT::Read2::Intron3'::3’ΔHygR + 5’ΔHygR::Intron5'::Read1::ACACATTCTAATTAA::Read2::Intron3'::3’ΔHygR + 5’ΔHygR::Intron5'::Read1::CCTCACTTTGATTGA::Read2::Intron3'::3’ΔHygR + 5’ΔHygR::Intron5'::Read1::TCACTCTTTGACGCG::Read2::Intron3'::3’ΔHygR + 5’ΔHygR::Intron5'::Read1::CAACTATATAAGGCT::Read2::Intron3'::3’ΔHygR + 5’ΔHygR::Intron5'::Read1::GCACACTATAAGGGT::Read2::Intron3'::3’ΔHygR + 5’ΔHygR::Intron5'::Read1::GTTCCTTGTCATGAA::Read2::Intron3'::3’ΔHygR + 5’ΔHygR::Intron5'::Read1::TATCACTTTCACTTC::Read2::Intron3'::3’ΔHygR + 5’ΔHygR::Intron5'::Read1::AGAGTCAAATAGGCA::Read2::Intron3'::3’ΔHygR + 5’ΔHygR::Intron5'::Read1::CATCGCTTTTACAAC::Read2::Intron3'::3’ΔHygR + 5’ΔHygR::Intron5'::Read1::CGTCATTCTCACTTT::Read2::Intron3'::3’ΔHygR + 5’ΔHygR::Intron5'::Read1::GTACTGTCTCACCAA::Read2::Intron3'::3’ΔHygR + 5’ΔHygR::Intron5'::Read1::TTACTATCTAAATGC::Read2::Intron3'::3’ΔHygR + 5’ΔHygR::Intron5'::Read1::AATCACTTTCACGCT::Read2::Intron3'::3’ΔHygR + 5’ΔHygR::Intron5'::Read1::CACCCATATAAGCTT::Read2::Intron3'::3’ΔHygR + 5’ΔHygR::Intron5'::Read1::CACCGATATTATACC::Read2::Intron3'::3’ΔHygR + 5’ΔHygR::Intron5'::Read1::CTTCTCTATGAGCGT::Read2::Intron3'::3’ΔHygR + 5’ΔHygR::Intron5'::Read1::GGCCGCTATTACTGA::Read2::Intron3'::3’ΔHygR + 5’ΔHygR::Intron5'::Read1::TGGCGTTGTGACATG::Read2::Intron3'::3’ΔHygR + 5’ΔHygR::Intron5'::Read1::TTACTATTTAACTTT::Read2::Intron3'::3’ΔHygR + 5’ΔHygR::Intron5'::Read1::ACTCCATGTCAACTC::Read2::Intron3'::3’ΔHygR + 5’ΔHygR::Intron5'::Read1::CAACTATGTAAGGTC::Read2::Intron3'::3’ΔHygR + 5’ΔHygR::Intron5'::Read1::GCTCTCTATAATGCG::Read2::Intron3'::3’ΔHygR + 5’ΔHygR::Intron5'::Read1::AATCGTTCTAAAATA::Read2::Intron3'::3’ΔHygR + 5’ΔHygR::Intron5'::Read1::AGACCATGTAATTTT::Read2::Intron3'::3’ΔHygR + 5’ΔHygR::Intron5'::Read1::AGTCAATATCAGCCC::Read2::Intron3'::3’ΔHygR + 5’ΔHygR::Intron5'::Read1::ATTCTATTTGACTAC::Read2::Intron3'::3’ΔHygR + 5’ΔHygR::Intron5'::Read1::CACCCTTATTACTTT::Read2::Intron3'::3’ΔHygR + 5’ΔHygR::Intron5'::Read1::CCCCTTTCTCAAGCT::Read2::Intron3'::3’ΔHygR + 5’ΔHygR::Intron5'::Read1::CGACCCTGTGACTTC::Read2::Intron3'::3’ΔHygR + 5’ΔHygR::Intron5'::Read1::CGGCCTTCTGAGGAT::Read2::Intron3'::3’ΔHygR + 5’ΔHygR::Intron5'::Read1::CTCCCTTGTCATCCC::Read2::Intron3'::3’ΔHygR + 5’ΔHygR::Intron5'::Read1::ATTCATTTTTACTAC::Read2::Intron3'::3’ΔHygR + 5’ΔHygR::Intron5'::Read1::GGACTCTATAAAACT::Read2::Intron3'::3’ΔHygR + 5’ΔHygR::Intron5'::Read1::AAGCGATATTAATCA::Read2::Intron3'::3’ΔHygR + 5’ΔHygR::Intron5'::Read1::ACCCATTGTTATCCC::Read2::Intron3'::3’ΔHygR + 5’ΔHygR::Intron5'::Read1::ATTCGATATCAGCAA::Read2::Intron3'::3’ΔHygR + 5’ΔHygR::Intron5'::Read1::CAACAGTGTCATTCC::Read2::Intron3'::3’ΔHygR + 5’ΔHygR::Intron5'::Read1::CTACTGTTTGAAACT::Read2::Intron3'::3’ΔHygR + 5’ΔHygR::Intron5'::Read1::GGTCATTATCATATC::Read2::Intron3'::3’ΔHygR + 5’ΔHygR::Intron5'::Read1::GCACCTTTTCATTTG::Read2::Intron3'::3’ΔHygR + 5’ΔHygR::Intron5'::Read1::TCCCCTTCTGAACCC::Read2::Intron3'::3’ΔHygR + 5’ΔHygR::Intron5'::Read1::TCGCTCTGTGAAGTC::Read2::Intron3'::3’ΔHygR + 5’ΔHygR::Intron5'::Read1::TTACAATGTTACATG::Read2::Intron3'::3’ΔHygR + 5’ΔHygR::Intron5'::Read1::TTGCACTCTGACTCC::Read2::Intron3'::3’ΔHygR + 5’ΔHygR::Intron5'::Read1::ATCCGATATAACATC::Read2::Intron3'::3’ΔHygR + 5’ΔHygR::Intron5'::Read1::CTCCTATTTAATGTG::Read2::Intron3'::3’ΔHygR + 5’ΔHygR::Intron5'::Read1::TACCCATCTCAAATG::Read2::Intron3'::3’ΔHygR + 5’ΔHygR::Intron5'::Read1::TTTCTATTTGAATGC::Read2::Intron3'::3’ΔHygR + 5’ΔHygR::Intron5'::Read1::AACCCTTGTCACCGC::Read2::Intron3'::3’ΔHygR + 5’ΔHygR::Intron5'::Read1::ACTCACTCTTAATTA::Read2::Intron3'::3’ΔHygR + 5’ΔHygR::Intron5'::Read1::CAGCTTTGTAAAACT::Read2::Intron3'::3’ΔHygR + 5’ΔHygR::Intron5'::Read1::CCACCTTTTCAACCC::Read2::Intron3'::3’ΔHygR + 5’ΔHygR::Intron5'::Read1::CTACTGTTTAATTCG::Read2::Intron3'::3’ΔHygR + 5’ΔHygR::Intron5'::Read1::CTCCTGTGTTATTCC::Read2::Intron3'::3’ΔHygR + 5’ΔHygR::Intron5'::Read1::GCTCATTTTAATTTA::Read2::Intron3'::3’ΔHygR + 5’ΔHygR::Intron5'::Read1::GTTCATTCTTAAGCC::Read2::Intron3'::3’ΔHygR + 5’ΔHygR::Intron5'::Read1::TATCCTTCTAACCTC::Read2::Intron3'::3’ΔHygR + 5’ΔHygR::Intron5'::Read1::TCTCCTTGTCATGCA::Read2::Intron3'::3’ΔHygR + 5’ΔHygR::Intron5'::Read1::TTCCACTATAACACC::Read2::Intron3'::3’ΔHygR +

5’ΔHygR::Intron5'::Read1::AAGCTATTTCAGTCC::Read2::Intron3'::3’ΔHygR + 5’ΔHygR::Intron5'::Read1::ATCCTCTTTGAACCA::Read2::Intron3'::3’ΔHygR + 5’ΔHygR::Intron5'::Read1::CTCCCATTTGAATCA::Read2::Intron3'::3’ΔHygR + 5’ΔHygR::Intron5'::Read1::GCGCAATGTTATGGA::Read2::Intron3'::3’ΔHygR + 5’ΔHygR::Intron5'::Read1::GTACTATGTCAACAG::Read2::Intron3'::3’ΔHygR + 5’ΔHygR::Intron5'::Read1::GTGCCCTCTCAGTGA::Read2::Intron3'::3’ΔHygR + 5’ΔHygR::Intron5'::Read1::TGGCCATATCACGGC::Read2::Intron3'::3’ΔHygR + 5’ΔHygR::Intron5'::Read1::TTACAATATCAACGT::Read2::Intron3'::3’ΔHygR + 5’ΔHygR::Intron5'::Read1::AAACCCTCTAAAATC::Read2::Intron3'::3’ΔHygR + 5’ΔHygR::Intron5'::Read1::ATCCTCTGTAATAGC::Read2::Intron3'::3’ΔHygR + 5’ΔHygR::Intron5'::Read1::CCTCCCTTTCATGAA::Read2::Intron3'::3’ΔHygR + 5’ΔHygR::Intron5'::Read1::CTACGTTGTTAAGCT::Read2::Intron3'::3’ΔHygR + 5’ΔHygR::Intron5'::Read1::GATCATTTTGAGCAC::Read2::Intron3'::3’ΔHygR + 5’ΔHygR::Intron5'::Read1::AGACCATTTCATCGT::Read2::Intron3'::3’ΔHygR + 5’ΔHygR::Intron5'::Read1::ATGCAGTTTTAACCG::Read2::Intron3'::3’ΔHygR + 5’ΔHygR::Intron5'::Read1::CACCCTTATTACAAC::Read2::Intron3'::3’ΔHygR + 5’ΔHygR::Intron5'::Read1::GACCTGTGTCAGAAC::Read2::Intron3'::3’ΔHygR + 5’ΔHygR::Intron5'::Read1::GATCCATTTAAGAAC::Read2::Intron3'::3’ΔHygR + 5’ΔHygR::Intron5'::Read1::GTCCGGTTTTAACGT::Read2::Intron3'::3’ΔHygR + 5’ΔHygR::Intron5'::Read1::GTGCCCTGTTATTAC::Read2::Intron3'::3’ΔHygR + 5’ΔHygR::Intron5'::Read1::TACCATTTTGATCAC::Read2::Intron3'::3’ΔHygR + 5’ΔHygR::Intron5'::Read1::TAGCTTTTTAATTTT::Read2::Intron3'::3’ΔHygR + 5’ΔHygR::Intron5'::Read1::TTACAATCTTATATG::Read2::Intron3'::3’ΔHygR + 5’ΔHygR::Intron5'::Read1::AACCTCTCTAAGGCA::Read2::Intron3'::3’ΔHygR + 5’ΔHygR::Intron5'::Read1::ATGCTCTATGAATTC::Read2::Intron3'::3’ΔHygR + 5’ΔHygR::Intron5'::Read1::ATTCCGTTTCATAAT::Read2::Intron3'::3’ΔHygR + 5’ΔHygR::Intron5'::Read1::TAGCCTTTTGATTAC::Read2::Intron3'::3’ΔHygR + 5’ΔHygR::Intron5'::Read1::TTACTTTGTGAATAC::Read2::Intron3'::3’ΔHygR + 5’ΔHygR::Intron5'::Read1::TTCCATTATTACCAA::Read2::Intron3'::3’ΔHygR + 5’ΔHygR::Intron5'::Read1::AGGCATTTTGATAAA::Read2::Intron3'::3’ΔHygR + 5’ΔHygR::Intron5'::Read1::CAGCGGTGTCATTAA::Read2::Intron3'::3’ΔHygR + 5’ΔHygR::Intron5'::Read1::CTACAATTTCACTTT::Read2::Intron3'::3’ΔHygR + 5’ΔHygR::Intron5'::Read1::CTTCTGTATCATTGC::Read2::Intron3'::3’ΔHygR + 5’ΔHygR::Intron5'::Read1::TTGCCTTGTCAACGT::Read2::Intron3'::3’ΔHygR + 5’ΔHygR::Intron5'::Read1::AATCTGTGTTACCCG::Read2::Intron3'::3’ΔHygR + 5’ΔHygR::Intron5'::Read1::CATCATTTTAAATGA::Read2::Intron3'::3’ΔHygR + 5’ΔHygR::Intron5'::Read1::GTTCTTTTTTATAAT::Read2::Intron3'::3’ΔHygR + 5’ΔHygR::Intron5'::Read1::TGTCCTTGTCATTCC::Read2::Intron3'::3’ΔHygR + 5’ΔHygR::Intron5'::Read1::TTACTATATCAGTCG::Read2::Intron3'::3’ΔHygR + 5’ΔHygR::Intron5'::Read1::ATTCAGTTTTATCTC::Read2::Intron3'::3’ΔHygR + 5’ΔHygR::Intron5'::Read1::CAGCGCTGTTAACCC::Read2::Intron3'::3’ΔHygR + 5’ΔHygR::Intron5'::Read1::TAGCTGTTTAATCAA::Read2::Intron3'::3’ΔHygR + 5’ΔHygR::Intron5'::Read1::TGGCTCTTTAAACCC::Read2::Intron3'::3’ΔHygR + 5’ΔHygR::Intron5'::Read1::TTACTCTTTTATTGC::Read2::Intron3'::3’ΔHygR + 5’ΔHygR::Intron5'::Read1::ATACTATTTTACTTC::Read2::Intron3'::3’ΔHygR + 5’ΔHygR::Intron5'::Read1::CAACAATTTAAATCC::Read2::Intron3'::3’ΔHygR + 5’ΔHygR::Intron5'::Read1::CAACCTTCTAATATC::Read2::Intron3'::3’ΔHygR + 5’ΔHygR::Intron5'::Read1::GAGCAGTGTTACATG::Read2::Intron3'::3’ΔHygR + 5’ΔHygR::Intron5'::Read1::TACCAGTATAATATC::Read2::Intron3'::3’ΔHygR + 5’ΔHygR::Intron5'::Read1::TACCATTGTTACTAA::Read2::Intron3'::3’ΔHygR + 5’ΔHygR::Intron5'::Read1::ATCCAGTATCACCTC::Read2::Intron3'::3’ΔHygR + 5’ΔHygR::Intron5'::Read1::ATTCTTTTTTATGAA::Read2::Intron3'::3’ΔHygR + 5’ΔHygR::Intron5'::Read1::CCACGATATTACCAT::Read2::Intron3'::3’ΔHygR + 5’ΔHygR::Intron5'::Read1::GAACGATTTTATCGA::Read2::Intron3'::3’ΔHygR + 5’ΔHygR::Intron5'::Read1::AGCCAGTTTCATTAC::Read2::Intron3'::3’ΔHygR + 5’ΔHygR::Intron5'::Read1::CCTCGCTATTACTCA::Read2::Intron3'::3’ΔHygR + 5’ΔHygR::Intron5'::Read1::CGACTTTATTATCTC::Read2::Intron3'::3’ΔHygR + 5’ΔHygR::Intron5'::Read1::GAGCGTTGTCATAGT::Read2::Intron3'::3’ΔHygR + 5’ΔHygR::Intron5'::Read1::GCCCCATTTAAATCA::Read2::Intron3'::3’ΔHygR + 5’ΔHygR::Intron5'::Read1::AAACTATATTATCTC::Read2::Intron3'::3’ΔHygR + 5’ΔHygR::Intron5'::Read1::CCTCGCTGTCATTTC::Read2::Intron3'::3’ΔHygR + 5’ΔHygR::Intron5'::Read1::TCCCTATTTCATATC::Read2::Intron3'::3’ΔHygR + 5’ΔHygR::Intron5'::Read1::ACGCAGTGTAAGAGC::Read2::Intron3'::3’ΔHygR + 5’ΔHygR::Intron5'::Read1::ATACTCTATGATTGC::Read2::Intron3'::3’ΔHygR + 5’ΔHygR::Intron5'::Read1::CCCCATTGTAATGCA::Read2::Intron3'::3’ΔHygR + 5’ΔHygR::Intron5'::Read1::TATCTTTGTTACTAG::Read2::Intron3'::3’ΔHygR + 5’ΔHygR::Intron5'::Read1::CCTCACTATTATCTT::Read2::Intron3'::3’ΔHygR + 5’ΔHygR::Intron5'::Read1::CTCCCTTTTTAGAGC::Read2::Intron3'::3’ΔHygR + 5’ΔHygR::Intron5'::Read1::GAACAATCTTACGTT::Read2::Intron3'::3’ΔHygR + 5’ΔHygR::Intron5'::Read1::GATCCGTTTTAGGCC::Read2::Intron3'::3’ΔHygR +

5’ΔHygR::Intron5'::Read1::GCGCAATCTGATCTA::Read2::Intron3'::3’ΔHygR + 5’ΔHygR::Intron5'::Read1::TACCCTTCTAATGAT::Read2::Intron3'::3’ΔHygR + 5’ΔHygR::Intron5'::Read1::TCCCTATTTGAGAAG::Read2::Intron3'::3’ΔHygR + 5’ΔHygR::Intron5'::Read1::AGACCATTTTACACT::Read2::Intron3'::3’ΔHygR + 5’ΔHygR::Intron5'::Read1::GTGCATTATCAATCA::Read2::Intron3'::3’ΔHygR + 5’ΔHygR::Intron5'::Read1::GTGCCTTGTTACACG::Read2::Intron3'::3’ΔHygR + 5’ΔHygR::Intron5'::Read1::TATCTGTGTCAGAAC::Read2::Intron3'::3’ΔHygR + 5’ΔHygR::Intron5'::Read1::TGCCTATATTAGATT::Read2::Intron3'::3’ΔHygR + 5’ΔHygR::Intron5'::Read1::TTGCTTTCTAACGAC::Read2::Intron3'::3’ΔHygR + 5’ΔHygR::Intron5'::Read1::CAACAATATAAGTAA::Read2::Intron3'::3’ΔHygR + 5’ΔHygR::Intron5'::Read1::CAACCTTATCAAAAG::Read2::Intron3'::3’ΔHygR + 5’ΔHygR::Intron5'::Read1::CTCCGTTATCAGATT::Read2::Intron3'::3’ΔHygR + 5’ΔHygR::Intron5'::Read1::CGACATTCTAAAAAT::Read2::Intron3'::3’ΔHygR + 5’ΔHygR::Intron5'::Read1::GGCCTATTTCACTTC::Read2::Intron3'::3’ΔHygR + 5’ΔHygR::Intron5'::Read1::GTACCTTCTCAATCC::Read2::Intron3'::3’ΔHygR + 5’ΔHygR::Intron5'::Read1::TCTCGTTTTCAAGAC::Read2::Intron3'::3’ΔHygR + 5’ΔHygR::Intron5'::Read1::TTTCTTTGTAAAAAC::Read2::Intron3'::3’ΔHygR + 5’ΔHygR::Intron5'::Read1::ACCCGATCTGATGTT::Read2::Intron3'::3’ΔHygR + 5’ΔHygR::Intron5'::Read1::GGCCACTATCAGATT::Read2::Intron3'::3’ΔHygR + 5’ΔHygR::Intron5'::Read1::CGGCCTTGTTAATAA::Read2::Intron3'::3’ΔHygR + 5’ΔHygR::Intron5'::Read1::GATCATTCTGAACTA::Read2::Intron3'::3’ΔHygR + 5’ΔHygR::Intron5'::Read1::GCCCCGTATGACCCA::Read2::Intron3'::3’ΔHygR + 5’ΔHygR::Intron5'::Read1::TATCGGTCTAACTAG::Read2::Intron3'::3’ΔHygR + 5’ΔHygR::Intron5'::Read1::TCCCCCTATGATGGT::Read2::Intron3'::3’ΔHygR + 5’ΔHygR::Intron5'::Read1::AAGCACTCTTATTAA::Read2::Intron3'::3’ΔHygR + 5’ΔHygR::Intron5'::Read1::CGGCGCTCTAATCTC::Read2::Intron3'::3’ΔHygR + 5’ΔHygR::Intron5'::Read1::CTACATTCTCATTCC::Read2::Intron3'::3’ΔHygR + 5’ΔHygR::Intron5'::Read1::TATCTTTATCACTGA::Read2::Intron3'::3’ΔHygR + 5’ΔHygR::Intron5'::Read1::TTACAGTTTAAACTA::Read2::Intron3'::3’ΔHygR + 5’ΔHygR::Intron5'::Read1::ACTCTTTCTTAACAC::Read2::Intron3'::3’ΔHygR + 5’ΔHygR::Intron5'::Read1::CACCCATGTTAAGAA::Read2::Intron3'::3’ΔHygR + 5’ΔHygR::Intron5'::Read1::CTCCGTTGTGACCGC::Read2::Intron3'::3’ΔHygR + 5’ΔHygR::Intron5'::Read1::GGGCTCTTTCAAGTC::Read2::Intron3'::3’ΔHygR + 5’ΔHygR::Intron5'::Read1::TGGCCATATCACACG::Read2::Intron3'::3’ΔHygR + 5’ΔHygR::Intron5'::Read1::CGACAGTTTTACGCA::Read2::Intron3'::3’ΔHygR + 5’ΔHygR::Intron5'::Read1::GTCCGGTGTCAGTCT::Read2::Intron3'::3’ΔHygR + 5’ΔHygR::Intron5'::Read1::TACCTCTGTAAGCAC::Read2::Intron3'::3’ΔHygR + 5’ΔHygR::Intron5'::Read1::TTGCCATATAACTCT::Read2::Intron3'::3’ΔHygR + 5’ΔHygR::Intron5'::Read1::AAACCTTATAATCTG::Read2::Intron3'::3’ΔHygR + 5’ΔHygR::Intron5'::Read1::AGGCTCTATGACCTT::Read2::Intron3'::3’ΔHygR + 5’ΔHygR::Intron5'::Read1::CTCCTTTTTAATCCA::Read2::Intron3'::3’ΔHygR + 5’ΔHygR::Intron5'::Read1::GTTCCATCTTACTAT::Read2::Intron3'::3’ΔHygR + 5’ΔHygR::Intron5'::Read1::TTACTTTATAAGAGA::Read2::Intron3'::3’ΔHygR + 5’ΔHygR::Intron5'::Read1::TTTCTTTTTCATCTG::Read2::Intron3'::3’ΔHygR + 5’ΔHygR::Intron5'::Read1::CGACGTTTTAACAAC::Read2::Intron3'::3’ΔHygR + 5’ΔHygR::Intron5'::Read1::CTGCTCTGTAAGAGA::Read2::Intron3'::3’ΔHygR + 5’ΔHygR::Intron5'::Read1::CTTCCTTATCATCCA::Read2::Intron3'::3’ΔHygR + 5’ΔHygR::Intron5'::Read1::GCGCGATTTTAGTAC::Read2::Intron3'::3’ΔHygR + 5’ΔHygR::Intron5'::Read1::GCTCCATCTAACCCA::Read2::Intron3'::3’ΔHygR + 5’ΔHygR::Intron5'::Read1::TAGCAATGTAACACT::Read2::Intron3'::3’ΔHygR + 5’ΔHygR::Intron5'::Read1::TCGCTGTTTTACGGT::Read2::Intron3'::3’ΔHygR + 5’ΔHygR::Intron5'::Read1::TTTCACTTTAATTGC::Read2::Intron3'::3’ΔHygR + 5’ΔHygR::Intron5'::Read1::ATACTCTATTAACCA::Read2::Intron3'::3’ΔHygR + 5’ΔHygR::Intron5'::Read1::ATTCGCTATAAACAT::Read2::Intron3'::3’ΔHygR + 5’ΔHygR::Intron5'::Read1::CAACCTTATTAAAAC::Read2::Intron3'::3’ΔHygR + 5’ΔHygR::Intron5'::Read1::CCACGCTTTGATAAA::Read2::Intron3'::3’ΔHygR + 5’ΔHygR::Intron5'::Read1::GGCCCTTCTTACCGC::Read2::Intron3'::3’ΔHygR + 5’ΔHygR::Intron5'::Read1::GACCCATTTTAAACA::Read2::Intron3'::3’ΔHygR + 5’ΔHygR::Intron5'::Read1::GCTCACTATTAAACT::Read2::Intron3'::3’ΔHygR + 5’ΔHygR::Intron5'::Read1::GCGCTCTGTTAGTTG::Read2::Intron3'::3’ΔHygR + 5’ΔHygR::Intron5'::Read1::TCTCGGTGTTACCAG::Read2::Intron3'::3’ΔHygR + 5’ΔHygR::Intron5'::Read1::CCACACTATCATTAC::Read2::Intron3'::3’ΔHygR + 5’ΔHygR::Intron5'::Read1::CCTCATTTTGACCTC::Read2::Intron3'::3’ΔHygR + 5’ΔHygR::Intron5'::Read1::CCTCGTTATCACTTA::Read2::Intron3'::3’ΔHygR + 5’ΔHygR::Intron5'::Read1::GGACATTATTAGCCT::Read2::Intron3'::3’ΔHygR + 5’ΔHygR::Intron5'::Read1::TATCACTATGATCCA::Read2::Intron3'::3’ΔHygR + 5’ΔHygR::Intron5'::Read1::AAACTCTATGAAATC::Read2::Intron3'::3’ΔHygR + 5’ΔHygR::Intron5'::Read1::ACACCATTTTAGCCC::Read2::Intron3'::3’ΔHygR + 5’ΔHygR::Intron5'::Read1::ACTCCATCTTACAAG::Read2::Intron3'::3’ΔHygR + 5’ΔHygR::Intron5'::Read1::AGCCGCTATGAAATC::Read2::Intron3'::3’ΔHygR +

5’ΔHygR::Intron5'::Read1::CTACTCTGTTAGGCC::Read2::Intron3'::3’ΔHygR + 5’ΔHygR::Intron5'::Read1::GACCCATTTAATAAC::Read2::Intron3'::3’ΔHygR + 5’ΔHygR::Intron5'::Read1::GATCTTTCTGAAAAA::Read2::Intron3'::3’ΔHygR + 5’ΔHygR::Intron5'::Read1::GCCCACTATTACTTA::Read2::Intron3'::3’ΔHygR + 5’ΔHygR::Intron5'::Read1::TACCCATTTCAGCAA::Read2::Intron3'::3’ΔHygR + 5’ΔHygR::Intron5'::Read1::AATCTTTATGAGAAG::Read2::Intron3'::3’ΔHygR + 5’ΔHygR::Intron5'::Read1::AGTCTTTTTCACGTA::Read2::Intron3'::3’ΔHygR + 5’ΔHygR::Intron5'::Read1::ATTCGGTGTTACTAT::Read2::Intron3'::3’ΔHygR + 5’ΔHygR::Intron5'::Read1::CACCTTTTTAAGATC::Read2::Intron3'::3’ΔHygR + 5’ΔHygR::Intron5'::Read1::CTTCGTTCTTAATGC::Read2::Intron3'::3’ΔHygR + 5’ΔHygR::Intron5'::Read1::TATCTTTTTAACTGA::Read2::Intron3'::3’ΔHygR + 5’ΔHygR::Intron5'::Read1::AATCGATCTTAGCTT::Read2::Intron3'::3’ΔHygR + 5’ΔHygR::Intron5'::Read1::ATACGATTTTACTAC::Read2::Intron3'::3’ΔHygR + 5’ΔHygR::Intron5'::Read1::ATCCTTTGTTAATAG::Read2::Intron3'::3’ΔHygR + 5’ΔHygR::Intron5'::Read1::CTCCTTTCTTACCCT::Read2::Intron3'::3’ΔHygR + 5’ΔHygR::Intron5'::Read1::TAGCGTTTTAACAAA::Read2::Intron3'::3’ΔHygR + 5’ΔHygR::Intron5'::Read1::TCCCCCTCTTAATGC::Read2::Intron3'::3’ΔHygR + 5’ΔHygR::Intron5'::Read1::TGACTATCTGAATCC::Read2::Intron3'::3’ΔHygR + 5’ΔHygR::Intron5'::Read1::TTACGTTCTAATAAC::Read2::Intron3'::3’ΔHygR + 5’ΔHygR::Intron5'::Read1::ACACTCTGTAAGCCT::Read2::Intron3'::3’ΔHygR + 5’ΔHygR::Intron5'::Read1::AGACACTATCACCTC::Read2::Intron3'::3’ΔHygR + 5’ΔHygR::Intron5'::Read1::GATCATTGTGATTTG::Read2::Intron3'::3’ΔHygR + 5’ΔHygR::Intron5'::Read1::TTCCACTTTCACGCC::Read2::Intron3'::3’ΔHygR + 5’ΔHygR::Intron5'::Read1::ACTCCTTTTAACTAT::Read2::Intron3'::3’ΔHygR + 5’ΔHygR::Intron5'::Read1::AGTCTATTTTAGTAA::Read2::Intron3'::3’ΔHygR + 5’ΔHygR::Intron5'::Read1::ATCCGTTTTCAGTTC::Read2::Intron3'::3’ΔHygR + 5’ΔHygR::Intron5'::Read1::CCCCCGTTTAATGAA::Read2::Intron3'::3’ΔHygR + 5’ΔHygR::Intron5'::Read1::CTCCTATTTAAAGTC::Read2::Intron3'::3’ΔHygR + 5’ΔHygR::Intron5'::Read1::GCACACTCTTAAGTT::Read2::Intron3'::3’ΔHygR + 5’ΔHygR::Intron5'::Read1::GGTCGTTCTCACTCG::Read2::Intron3'::3’ΔHygR + 5’ΔHygR::Intron5'::Read1::TACCCTTTTCATTAC::Read2::Intron3'::3’ΔHygR + 5’ΔHygR::Intron5'::Read1::ATCCAATGTTACCAA::Read2::Intron3'::3’ΔHygR + 5’ΔHygR::Intron5'::Read1::CCACAGTTTCATTCA::Read2::Intron3'::3’ΔHygR + 5’ΔHygR::Intron5'::Read1::GACCCATCTCACCGA::Read2::Intron3'::3’ΔHygR + 5’ΔHygR::Intron5'::Read1::GTACGCTATTACTTT::Read2::Intron3'::3’ΔHygR + 5’ΔHygR::Intron5'::Read1::TAGCCGTATTATTAC::Read2::Intron3'::3’ΔHygR + 5’ΔHygR::Intron5'::Read1::ATACTGTGTAAAACT::Read2::Intron3'::3’ΔHygR + 5’ΔHygR::Intron5'::Read1::CATCTTTGTTAATCG::Read2::Intron3'::3’ΔHygR + 5’ΔHygR::Intron5'::Read1::CTGCATTGTCACCCA::Read2::Intron3'::3’ΔHygR + 5’ΔHygR::Intron5'::Read1::GTTCTCTCTAAATAC::Read2::Intron3'::3’ΔHygR + 5’ΔHygR::Intron5'::Read1::TCACAGTATTAGTGC::Read2::Intron3'::3’ΔHygR + 5’ΔHygR::Intron5'::Read1::TGCCTTTGTCATTCC::Read2::Intron3'::3’ΔHygR + 5’ΔHygR::Intron5'::Read1::TTACTTTATGAACAC::Read2::Intron3'::3’ΔHygR + 5’ΔHygR::Intron5'::Read1::CGCCATCATTACCAC::Read2::Intron3'::3’ΔHygR + 5’ΔHygR::Intron5'::Read1::CTTCGGTATAACGTA::Read2::Intron3'::3’ΔHygR + 5’ΔHygR::Intron5'::Read1::CTTCGTTATGATAGT::Read2::Intron3'::3’ΔHygR + 5’ΔHygR::Intron5'::Read1::TATCCCTTTGAGACC::Read2::Intron3'::3’ΔHygR + 5’ΔHygR::Intron5'::Read1::TGGCCCTCTAACTCA::Read2::Intron3'::3’ΔHygR + 5’ΔHygR::Intron5'::Read1::TGTCGTTATAATCCG::Read2::Intron3'::3’ΔHygR + 5’ΔHygR::Intron5'::Read1::TTACCTTTTTACGAT::Read2::Intron3'::3’ΔHygR + 5’ΔHygR::Intron5'::Read1::TTTCAATGTCATCAG::Read2::Intron3'::3’ΔHygR + 5’ΔHygR::Intron5'::Read1::AATCGATTTGAGCAA::Read2::Intron3'::3’ΔHygR + 5’ΔHygR::Intron5'::Read1::AATCTATTTAATTCT::Read2::Intron3'::3’ΔHygR + 5’ΔHygR::Intron5'::Read1::AATCTGTTTAACTTA::Read2::Intron3'::3’ΔHygR + 5’ΔHygR::Intron5'::Read1::GTACTTTCTTAGACC::Read2::Intron3'::3’ΔHygR + 5’ΔHygR::Intron5'::Read1::ATACCTTTTTATCCC::Read2::Intron3'::3’ΔHygR + 5’ΔHygR::Intron5'::Read1::CTTCGGTGTAACGCA::Read2::Intron3'::3’ΔHygR + 5’ΔHygR::Intron5'::Read1::GTTCTATTTAAAACG::Read2::Intron3'::3’ΔHygR + 5’ΔHygR::Intron5'::Read1::TCGCAGTTTAACTAG::Read2::Intron3'::3’ΔHygR + 5’ΔHygR::Intron5'::Read1::TGCATAAGAACGTTA::Read2::Intron3'::3’ΔHygR + 5’ΔHygR::Intron5'::Read1::TTTCTATTTAAATGT::Read2::Intron3'::3’ΔHygR + 5’ΔHygR::Intron5'::Read1::ACCCACTATTACGTC::Read2::Intron3'::3’ΔHygR + 5’ΔHygR::Intron5'::Read1::ATACTCTCTAATCTA::Read2::Intron3'::3’ΔHygR + 5’ΔHygR::Intron5'::Read1::CTCCGTTGTAAGGCA::Read2::Intron3'::3’ΔHygR + 5’ΔHygR::Intron5'::Read1::GACCATTATAACAAC::Read2::Intron3'::3’ΔHygR + 5’ΔHygR::Intron5'::Read1::TACCCATATAAATCT::Read2::Intron3'::3’ΔHygR + 5’ΔHygR::Intron5'::Read1::TGTCCGTCTAACTCT::Read2::Intron3'::3’ΔHygR + 5’ΔHygR::Intron5'::Read1::TTGCGTTTTGATTTG::Read2::Intron3'::3’ΔHygR + 5’ΔHygR::Intron5'::Read1::TTTCAGTATTACGAT::Read2::Intron3'::3’ΔHygR + 5’ΔHygR::Intron5'::Read1::ATACCCTGTGACTCT::Read2::Intron3'::3’ΔHygR +

5’ΔHygR::Intron5'::Read1::TTCCTATTTCAGACC::Read2::Intron3'::3’ΔHygR + 5’ΔHygR::Intron5'::Read1::AATCGCTATGATGCA::Read2::Intron3'::3’ΔHygR + 5’ΔHygR::Intron5'::Read1::AGTCAGTGTTACCGA::Read2::Intron3'::3’ΔHygR + 5’ΔHygR::Intron5'::Read1::CCTCTATATGAGTGC::Read2::Intron3'::3’ΔHygR + 5’ΔHygR::Intron5'::Read1::CGTCGTTATCACCTC::Read2::Intron3'::3’ΔHygR + 5’ΔHygR::Intron5'::Read1::GTTCAGTGTAATCCA::Read2::Intron3'::3’ΔHygR + 5’ΔHygR::Intron5'::Read1::TATCCTTTTTAACAG::Read2::Intron3'::3’ΔHygR + 5’ΔHygR::Intron5'::Read1::ATTCTCTATCACCTC::Read2::Intron3'::3’ΔHygR + 5’ΔHygR::Intron5'::Read1::GATCCCTATTAAGAT::Read2::Intron3'::3’ΔHygR + 5’ΔHygR::Intron5'::Read1::TACCTCTCTCATCGC::Read2::Intron3'::3’ΔHygR + 5’ΔHygR::Intron5'::Read1::TCTCAATGTTAGTTC::Read2::Intron3'::3’ΔHygR + 5’ΔHygR::Intron5'::Read1::TGGCGCTCTTAATTA::Read2::Intron3'::3’ΔHygR + 5’ΔHygR::Intron5'::Read1::AGGCAATGTTAATCG::Read2::Intron3'::3’ΔHygR + 5’ΔHygR::Intron5'::Read1::CTACTTTGTCACAAG::Read2::Intron3'::3’ΔHygR + 5’ΔHygR::Intron5'::Read1::GCTCTATTTTAATCC::Read2::Intron3'::3’ΔHygR + 5’ΔHygR::Intron5'::Read1::ACGCAGTGTTACTTA::Read2::Intron3'::3’ΔHygR + 5’ΔHygR::Intron5'::Read1::CATCGATTTAACTTA::Read2::Intron3'::3’ΔHygR + 5’ΔHygR::Intron5'::Read1::CGTCCATTTAATAGT::Read2::Intron3'::3’ΔHygR + 5’ΔHygR::Intron5'::Read1::GCTCGTTGTTACATT::Read2::Intron3'::3’ΔHygR + 5’ΔHygR::Intron5'::Read1::TAACTATCTTACAAC::Read2::Intron3'::3’ΔHygR + 5’ΔHygR::Intron5'::Read1::TGCCGCTATAATACA::Read2::Intron3'::3’ΔHygR + 5’ΔHygR::Intron5'::Read1::CCACGGTATAATATT::Read2::Intron3'::3’ΔHygR + 5’ΔHygR::Intron5'::Read1::AAACGCTGTAAACTA::Read2::Intron3'::3’ΔHygR + 5’ΔHygR::Intron5'::Read1::ACACGCTATTATCGT::Read2::Intron3'::3’ΔHygR + 5’ΔHygR::Intron5'::Read1::CCTCCGTGTTACACA::Read2::Intron3'::3’ΔHygR + 5’ΔHygR::Intron5'::Read1::GGTCAATGTAAACAA::Read2::Intron3'::3’ΔHygR + 5’ΔHygR::Intron5'::Read1::TGACATTCTTACGTC::Read2::Intron3'::3’ΔHygR + 5’ΔHygR::Intron5'::Read1::TTTCAATATCAGCAC::Read2::Intron3'::3’ΔHygR + 5’ΔHygR::Intron5'::Read1::AGCCCGTCTAATACA::Read2::Intron3'::3’ΔHygR + 5’ΔHygR::Intron5'::Read1::CGGCAATATCAGCCT::Read2::Intron3'::3’ΔHygR + 5’ΔHygR::Intron5'::Read1::TAACGTTTTAACTAA::Read2::Intron3'::3’ΔHygR + 5’ΔHygR::Intron5'::Read1::TGCCTGTTTAAATGC::Read2::Intron3'::3’ΔHygR + 5’ΔHygR::Intron5'::Read1::ATCCTCTGTAAAATT::Read2::Intron3'::3’ΔHygR + 5’ΔHygR::Intron5'::Read1::GTACCGTTTTACAAT::Read2::Intron3'::3’ΔHygR + 5’ΔHygR::Intron5'::Read1::TCGCGATATTAGTGT::Read2::Intron3'::3’ΔHygR + 5’ΔHygR::Intron5'::Read1::TGGCTTTTTTAACTA::Read2::Intron3'::3’ΔHygR + 5’ΔHygR::Intron5'::Read1::AATCCCTTTGATATC::Read2::Intron3'::3’ΔHygR + 5’ΔHygR::Intron5'::Read1::TAACGTTTTCAACTC::Read2::Intron3'::3’ΔHygR + 5’ΔHygR::Intron5'::Read1::TTACTTTCTTACACC::Read2::Intron3'::3’ΔHygR + 5’ΔHygR::Intron5'::Read1::TTTCGTTTTCAATCT::Read2::Intron3'::3’ΔHygR + 5’ΔHygR::Intron5'::Read1::ACCCTTTATGATGAC::Read2::Intron3'::3’ΔHygR + 5’ΔHygR::Intron5'::Read1::TAACCTTTTTACTAT::Read2::Intron3'::3’ΔHygR + 5’ΔHygR::Intron5'::Read1::TCCCTATCTCAGCGT::Read2::Intron3'::3’ΔHygR + 5’ΔHygR::Intron5'::Read1::GACCCGTGTCAAGAA::Read2::Intron3'::3’ΔHygR + 5’ΔHygR::Intron5'::Read1::GCCCTCTCTAACAAC::Read2::Intron3'::3’ΔHygR + 5’ΔHygR::Intron5'::Read1::TTGCCGTTTAACCTC::Read2::Intron3'::3’ΔHygR + 5’ΔHygR::Intron5'::Read1::TTTCCCTATCACCGC::Read2::Intron3'::3’ΔHygR + 5’ΔHygR::Intron5'::Read1::AACCAGTGTAATATC::Read2::Intron3'::3’ΔHygR + 5’ΔHygR::Intron5'::Read1::ACACCGTCTCACTGA::Read2::Intron3'::3’ΔHygR + 5’ΔHygR::Intron5'::Read1::GCCCGCTTTAACTAA::Read2::Intron3'::3’ΔHygR + 5’ΔHygR::Intron5'::Read1::TACCTATTTCACAGA::Read2::Intron3'::3’ΔHygR + 5’ΔHygR::Intron5'::Read1::TACCAGTTTAACTCG::Read2::Intron3'::3’ΔHygR + 5’ΔHygR::Intron5'::Read1::TATCCCTTTTAATAG::Read2::Intron3'::3’ΔHygR + 5’ΔHygR::Intron5'::Read1::TGACACTCTGATGTA::Read2::Intron3'::3’ΔHygR + 5’ΔHygR::Intron5'::Read1::TTGCTTTCTCAAGCC::Read2::Intron3'::3’ΔHygR + 5’ΔHygR::Intron5'::Read1::ATACGTTTTTATCAG::Read2::Intron3'::3’ΔHygR + 5’ΔHygR::Intron5'::Read1::ACGCTTTATGATGCA::Read2::Intron3'::3’ΔHygR + 5’ΔHygR::Intron5'::Read1::CGTCGTTCTAATTTT::Read2::Intron3'::3’ΔHygR + 5’ΔHygR::Intron5'::Read1::TATCACTGTCACGTT::Read2::Intron3'::3’ΔHygR + 5’ΔHygR::Intron5'::Read1::TCGCTGTATTAACCC::Read2::Intron3'::3’ΔHygR + 5’ΔHygR::Intron5'::Read1::TTCCTCTATTACTAT::Read2::Intron3'::3’ΔHygR + 5’ΔHygR::Intron5'::Read1::ATGCTGTCTCAGGGC::Read2::Intron3'::3’ΔHygR + 5’ΔHygR::Intron5'::Read1::ATTCGTTATGATAAC::Read2::Intron3'::3’ΔHygR + 5’ΔHygR::Intron5'::Read1::CACCGATATCAAAAC::Read2::Intron3'::3’ΔHygR + 5’ΔHygR::Intron5'::Read1::CCACCGTTTTAATAT::Read2::Intron3'::3’ΔHygR + 5’ΔHygR::Intron5'::Read1::CGCCACTGTTATATA::Read2::Intron3'::3’ΔHygR + 5’ΔHygR::Intron5'::Read1::GAACGCTATAAATGA::Read2::Intron3'::3’ΔHygR + 5’ΔHygR::Intron5'::Read1::GGGCTCTGTCATCTA::Read2::Intron3'::3’ΔHygR + 5’ΔHygR::Intron5'::Read1::TACCGGTGTTACGAT::Read2::Intron3'::3’ΔHygR + 5’ΔHygR::Intron5'::Read1::TAGCCCTATAAAAGC::Read2::Intron3'::3’ΔHygR +

5’ΔHygR::Intron5'::Read1::TGTCATTTTTAGTTC::Read2::Intron3'::3’ΔHygR + 5’ΔHygR::Intron5'::Read1::TTGCACTCTCACCCT::Read2::Intron3'::3’ΔHygR + 5’ΔHygR::Intron5'::Read1::ACACTTTTTAACTCA::Read2::Intron3'::3’ΔHygR + 5’ΔHygR::Intron5'::Read1::AGCCCTTATAAGATG::Read2::Intron3'::3’ΔHygR + 5’ΔHygR::Intron5'::Read1::ATTCCATATGACCCA::Read2::Intron3'::3’ΔHygR + 5’ΔHygR::Intron5'::Read1::CTTCTGTATCATTCA::Read2::Intron3'::3’ΔHygR + 5’ΔHygR::Intron5'::Read1::GGTCAATATCAAACT::Read2::Intron3'::3’ΔHygR + 5’ΔHygR::Intron5'::Read1::GTTCGGTGTTATATT::Read2::Intron3'::3’ΔHygR + 5’ΔHygR::Intron5'::Read1::ACACCTTGTTAACGG::Read2::Intron3'::3’ΔHygR + 5’ΔHygR::Intron5'::Read1::TGCCACTGTTATAAC::Read2::Intron3'::3’ΔHygR + 5’ΔHygR::Intron5'::Read1::TGCCGGTGTGAACAT::Read2::Intron3'::3’ΔHygR + 5’ΔHygR::Intron5'::Read1::GTACCTTTTGACTAT::Read2::Intron3'::3’ΔHygR + 5’ΔHygR::Intron5'::Read1::GTTCCCTATAAGCAA::Read2::Intron3'::3’ΔHygR + 5’ΔHygR::Intron5'::Read1::TAGCCATTTGAAGCA::Read2::Intron3'::3’ΔHygR + 5’ΔHygR::Intron5'::Read1::TATCGCTTTTAATGT::Read2::Intron3'::3’ΔHygR + 5’ΔHygR::Intron5'::Read1::TCACATTTTCATACG::Read2::Intron3'::3’ΔHygR + 5’ΔHygR::Intron5'::Read1::TCACGCTCTCAGGCA::Read2::Intron3'::3’ΔHygR + 5’ΔHygR::Intron5'::Read1::ATTCGCTTTCATCTA::Read2::Intron3'::3’ΔHygR + 5’ΔHygR::Intron5'::Read1::CCGCCTTATAATCAA::Read2::Intron3'::3’ΔHygR + 5’ΔHygR::Intron5'::Read1::CGACTATATCATCCC::Read2::Intron3'::3’ΔHygR + 5’ΔHygR::Intron5'::Read1::GAGCTTTTTAACACC::Read2::Intron3'::3’ΔHygR + 5’ΔHygR::Intron5'::Read1::GTACACTCTTATACA::Read2::Intron3'::3’ΔHygR + 5’ΔHygR::Intron5'::Read1::ACACATTTTCAGACA::Read2::Intron3'::3’ΔHygR + 5’ΔHygR::Intron5'::Read1::AGGCAATCTAATTGA::Read2::Intron3'::3’ΔHygR + 5’ΔHygR::Intron5'::Read1::AGGCATTTTCACATA::Read2::Intron3'::3’ΔHygR + 5’ΔHygR::Intron5'::Read1::ATCCCGTTTGACATC::Read2::Intron3'::3’ΔHygR + 5’ΔHygR::Intron5'::Read1::TCCCTTTATAATACA::Read2::Intron3'::3’ΔHygR + 5’ΔHygR::Intron5'::Read1::TGACTCTGTTATCCC::Read2::Intron3'::3’ΔHygR + 5’ΔHygR::Intron5'::Read1::ATTCTATGTGATCAT::Read2::Intron3'::3’ΔHygR + 5’ΔHygR::Intron5'::Read1::CAGCCTTTTTACTAC::Read2::Intron3'::3’ΔHygR + 5’ΔHygR::Intron5'::Read1::TAACCTTATGATCTC::Read2::Intron3'::3’ΔHygR + 5’ΔHygR::Intron5'::Read1::TAACTTTTTTATGTG::Read2::Intron3'::3’ΔHygR + 5’ΔHygR::Intron5'::Read1::TCTCTGTATCACTTA::Read2::Intron3'::3’ΔHygR + 5’ΔHygR::Intron5'::Read1::TTTCTTTGTAAACGT::Read2::Intron3'::3’ΔHygR + 5’ΔHygR::Intron5'::Read1::AGCCTGTTTTATCTC::Read2::Intron3'::3’ΔHygR + 5’ΔHygR::Intron5'::Read1::TTCCTGTATTAGTCC::Read2::Intron3'::3’ΔHygR + 5’ΔHygR::Intron5'::Read1::ACACACTCTCACATA::Read2::Intron3'::3’ΔHygR + 5’ΔHygR::Intron5'::Read1::ATCCAATCTAATGGT::Read2::Intron3'::3’ΔHygR + 5’ΔHygR::Intron5'::Read1::CGACTTTATCATCCA::Read2::Intron3'::3’ΔHygR + 5’ΔHygR::Intron5'::Read1::CTACAGTCTTATATT::Read2::Intron3'::3’ΔHygR + 5’ΔHygR::Intron5'::Read1::GTACGATTTTACGGG::Read2::Intron3'::3’ΔHygR + 5’ΔHygR::Intron5'::Read1::AGACCATCTCATGCA::Read2::Intron3'::3’ΔHygR + 5’ΔHygR::Intron5'::Read1::ATGCTATCTCATAAC::Read2::Intron3'::3’ΔHygR + 5’ΔHygR::Intron5'::Read1::CAGCAATGTTATCAA::Read2::Intron3'::3’ΔHygR + 5’ΔHygR::Intron5'::Read1::GCGCCCTATTAAGTC::Read2::Intron3'::3’ΔHygR + 5’ΔHygR::Intron5'::Read1::GGACACTATGATTCG::Read2::Intron3'::3’ΔHygR + 5’ΔHygR::Intron5'::Read1::TCCCTGTATTAACAA::Read2::Intron3'::3’ΔHygR + 5’ΔHygR::Intron5'::Read1::ATACCTTTTAACCCG::Read2::Intron3'::3’ΔHygR + 5’ΔHygR::Intron5'::Read1::CAACGTTCTTACAAC::Read2::Intron3'::3’ΔHygR + 5’ΔHygR::Intron5'::Read1::CACCGTTTTAAAAAC::Read2::Intron3'::3’ΔHygR + 5’ΔHygR::Intron5'::Read1::CTGCTATCTAAACAC::Read2::Intron3'::3’ΔHygR + 5’ΔHygR::Intron5'::Read1::TTCCCTTGTTACATT::Read2::Intron3'::3’ΔHygR + 5’ΔHygR::Intron5'::Read1::TTTCCTTCTTATGCC::Read2::Intron3'::3’ΔHygR + 5’ΔHygR::Intron5'::Read1::TTTCTATTTGACCCA::Read2::Intron3'::3’ΔHygR + 5’ΔHygR::Intron5'::Read1::ATGCGTTATGACTAT::Read2::Intron3'::3’ΔHygR + 5’ΔHygR::Intron5'::Read1::TCCCGATCTAATACT::Read2::Intron3'::3’ΔHygR + 5’ΔHygR::Intron5'::Read1::TCGCGTTCTAACAAG::Read2::Intron3'::3’ΔHygR + 5’ΔHygR::Intron5'::Read1::TCTCTGTTTGATTAA::Read2::Intron3'::3’ΔHygR + 5’ΔHygR::Intron5'::Read1::ACACCCTCTCAGCCT::Read2::Intron3'::3’ΔHygR + 5’ΔHygR::Intron5'::Read1::ATGCTATATTACTTC::Read2::Intron3'::3’ΔHygR + 5’ΔHygR::Intron5'::Read1::CAACCCTATAAACGA::Read2::Intron3'::3’ΔHygR + 5’ΔHygR::Intron5'::Read1::CAGCAATTTCATCAC::Read2::Intron3'::3’ΔHygR + 5’ΔHygR::Intron5'::Read1::CGTCTCTCTTAATTT::Read2::Intron3'::3’ΔHygR + 5’ΔHygR::Intron5'::Read1::GAACCATATAAGCTC::Read2::Intron3'::3’ΔHygR + 5’ΔHygR::Intron5'::Read1::GCACACTTTTAGTTT::Read2::Intron3'::3’ΔHygR + 5’ΔHygR::Intron5'::Read1::GTTCGTTGTAACTAT::Read2::Intron3'::3’ΔHygR + 5’ΔHygR::Intron5'::Read1::TTCCAATGTTAATCA::Read2::Intron3'::3’ΔHygR + 5’ΔHygR::Intron5'::Read1::AAACTCTTTGAAAGG::Read2::Intron3'::3’ΔHygR + 5’ΔHygR::Intron5'::Read1::ACCCCATGTAAACCG::Read2::Intron3'::3’ΔHygR + 5’ΔHygR::Intron5'::Read1::CTACAATCTGACCGC::Read2::Intron3'::3’ΔHygR +

5’ΔHygR::Intron5'::Read1::GATCAATCTTATAAA::Read2::Intron3'::3’ΔHygR + 5’ΔHygR::Intron5'::Read1::GCCCATTATAAATAT::Read2::Intron3'::3’ΔHygR + 5’ΔHygR::Intron5'::Read1::ACACCCTCTCATCAC::Read2::Intron3'::3’ΔHygR + 5’ΔHygR::Intron5'::Read1::ATTCTGTTTGATTCG::Read2::Intron3'::3’ΔHygR + 5’ΔHygR::Intron5'::Read1::CGACCGTTTTAGGTC::Read2::Intron3'::3’ΔHygR + 5’ΔHygR::Intron5'::Read1::CGACTTTGTTATAAC::Read2::Intron3'::3’ΔHygR + 5’ΔHygR::Intron5'::Read1::GGCCGCTCTCACTTC::Read2::Intron3'::3’ΔHygR + 5’ΔHygR::Intron5'::Read1::GGTCTCTATGAAACG::Read2::Intron3'::3’ΔHygR + 5’ΔHygR::Intron5'::Read1::TAACGGTTTGATCAG::Read2::Intron3'::3’ΔHygR + 5’ΔHygR::Intron5'::Read1::ACGCAGTGTGACATC::Read2::Intron3'::3’ΔHygR + 5’ΔHygR::Intron5'::Read1::AGCCCATATGAGCCC::Read2::Intron3'::3’ΔHygR + 5’ΔHygR::Intron5'::Read1::CAACATTCTCACACT::Read2::Intron3'::3’ΔHygR + 5’ΔHygR::Intron5'::Read1::CATCGATTTCACAAC::Read2::Intron3'::3’ΔHygR + 5’ΔHygR::Intron5'::Read1::GCTCACTTTTACACA::Read2::Intron3'::3’ΔHygR + 5’ΔHygR::Intron5'::Read1::TCACGGTTTAATTAA::Read2::Intron3'::3’ΔHygR + 5’ΔHygR::Intron5'::Read1::GATCGCTCTAATTAT::Read2::Intron3'::3’ΔHygR + 5’ΔHygR::Intron5'::Read1::GCTCGGTCTTATAAC::Read2::Intron3'::3’ΔHygR + 5’ΔHygR::Intron5'::Read1::GTTCCCTTTGACCTA::Read2::Intron3'::3’ΔHygR + 5’ΔHygR::Intron5'::Read1::TGCCCATCTCATTAA::Read2::Intron3'::3’ΔHygR + 5’ΔHygR::Intron5'::Read1::TGCCCATGTTACCCT::Read2::Intron3'::3’ΔHygR + 5’ΔHygR::Intron5'::Read1::ACACGCTGTGAACGA::Read2::Intron3'::3’ΔHygR + 5’ΔHygR::Intron5'::Read1::AGCCAGTATTAATAA::Read2::Intron3'::3’ΔHygR + 5’ΔHygR::Intron5'::Read1::CCTCACTGTCAAGTT::Read2::Intron3'::3’ΔHygR + 5’ΔHygR::Intron5'::Read1::GATCTATTTTAATCC::Read2::Intron3'::3’ΔHygR + 5’ΔHygR::Intron5'::Read1::GGGCCCTTTCAACTC::Read2::Intron3'::3’ΔHygR + 5’ΔHygR::Intron5'::Read1::TCCCGTTATCACGCT::Read2::Intron3'::3’ΔHygR + 5’ΔHygR::Intron5'::Read1::TCGCTTTTTCATTCT::Read2::Intron3'::3’ΔHygR + 5’ΔHygR::Intron5'::Read1::ATTCCCTATTAACAT::Read2::Intron3'::3’ΔHygR + 5’ΔHygR::Intron5'::Read1::ATTCCTTATGATTTC::Read2::Intron3'::3’ΔHygR + 5’ΔHygR::Intron5'::Read1::CATCGCTCTTAATCT::Read2::Intron3'::3’ΔHygR + 5’ΔHygR::Intron5'::Read1::CGACCCTTTCATTTC::Read2::Intron3'::3’ΔHygR + 5’ΔHygR::Intron5'::Read1::CGCCTGTTTTACATC::Read2::Intron3'::3’ΔHygR + 5’ΔHygR::Intron5'::Read1::AAACAGTATTATTTT::Read2::Intron3'::3’ΔHygR + 5’ΔHygR::Intron5'::Read1::CATCGATCTGACGGC::Read2::Intron3'::3’ΔHygR + 5’ΔHygR::Intron5'::Read1::CGTCTCTCTGAAGTC::Read2::Intron3'::3’ΔHygR + 5’ΔHygR::Intron5'::Read1::GACCGTTCTAAATCC::Read2::Intron3'::3’ΔHygR + 5’ΔHygR::Intron5'::Read1::GTTCGATCTTAGTCC::Read2::Intron3'::3’ΔHygR + 5’ΔHygR::Intron5'::Read1::ACACTCTTTAAGACC::Read2::Intron3'::3’ΔHygR + 5’ΔHygR::Intron5'::Read1::ACCCATTCTTATCAC::Read2::Intron3'::3’ΔHygR + 5’ΔHygR::Intron5'::Read1::ATGCTATGTCAGTTG::Read2::Intron3'::3’ΔHygR + 5’ΔHygR::Intron5'::Read1::CAACGCTATAAACTT::Read2::Intron3'::3’ΔHygR + 5’ΔHygR::Intron5'::Read1::TGCCCGTGTTACTAT::Read2::Intron3'::3’ΔHygR + 5’ΔHygR::Intron5'::Read1::TGGCCTTTTAACTTT::Read2::Intron3'::3’ΔHygR + 5’ΔHygR::Intron5'::Read1::CAACGTTATCACTAA::Read2::Intron3'::3’ΔHygR + 5’ΔHygR::Intron5'::Read1::CAGCAGTTTTATCCA::Read2::Intron3'::3’ΔHygR + 5’ΔHygR::Intron5'::Read1::CATCAATTTGACCCC::Read2::Intron3'::3’ΔHygR + 5’ΔHygR::Intron5'::Read1::CATCCATATCAACAC::Read2::Intron3'::3’ΔHygR + 5’ΔHygR::Intron5'::Read1::CCACCCTATTACGTT::Read2::Intron3'::3’ΔHygR + 5’ΔHygR::Intron5'::Read1::CTTCCGTCTGACCAC::Read2::Intron3'::3’ΔHygR + 5’ΔHygR::Intron5'::Read1::CTTCTATCTAATACC::Read2::Intron3'::3’ΔHygR + 5’ΔHygR::Intron5'::Read1::GCACTATTTAAGAAT::Read2::Intron3'::3’ΔHygR + 5’ΔHygR::Intron5'::Read1::GTTCAATCTAACTAG::Read2::Intron3'::3’ΔHygR + 5’ΔHygR::Intron5'::Read1::TATCGATCTTATCTT::Read2::Intron3'::3’ΔHygR + 5’ΔHygR::Intron5'::Read1::TCGCATTTTAAATGC::Read2::Intron3'::3’ΔHygR + 5’ΔHygR::Intron5'::Read1::TCGCATTTTAACTGT::Read2::Intron3'::3’ΔHygR + 5’ΔHygR::Intron5'::Read1::TTACAATTTGACTTA::Read2::Intron3'::3’ΔHygR + 5’ΔHygR::Intron5'::Read1::AACCGGTGTCACACC::Read2::Intron3'::3’ΔHygR + 5’ΔHygR::Intron5'::Read1::AGTCACTTTTAGCCC::Read2::Intron3'::3’ΔHygR + 5’ΔHygR::Intron5'::Read1::CCCCAATCTAACTAG::Read2::Intron3'::3’ΔHygR + 5’ΔHygR::Intron5'::Read1::CGACGTTTTTACCTT::Read2::Intron3'::3’ΔHygR + 5’ΔHygR::Intron5'::Read1::CGGCTATATCAACTA::Read2::Intron3'::3’ΔHygR + 5’ΔHygR::Intron5'::Read1::CTGCCATGTTACCAA::Read2::Intron3'::3’ΔHygR + 5’ΔHygR::Intron5'::Read1::CTTCCTTTTTAACCC::Read2::Intron3'::3’ΔHygR + 5’ΔHygR::Intron5'::Read1::TTTCATTATAATGGG::Read2::Intron3'::3’ΔHygR + 5’ΔHygR::Intron5'::Read1::ACCCGTTTTAATTAC::Read2::Intron3'::3’ΔHygR + 5’ΔHygR::Intron5'::Read1::CAACATTATTATTGT::Read2::Intron3'::3’ΔHygR + 5’ΔHygR::Intron5'::Read1::CACCATTTTAAGTGC::Read2::Intron3'::3’ΔHygR + 5’ΔHygR::Intron5'::Read1::CTTCTATGTAATATC::Read2::Intron3'::3’ΔHygR + 5’ΔHygR::Intron5'::Read1::GTTCACTATGACAAC::Read2::Intron3'::3’ΔHygR + 5’ΔHygR::Intron5'::Read1::CACCCCTTTAATCAC::Read2::Intron3'::3’ΔHygR +

5’ΔHygR::Intron5'::Read1::TCGCACTCTAATCGT::Read2::Intron3'::3’ΔHygR + 5’ΔHygR::Intron5'::Read1::AACCCTTATGACTCA::Read2::Intron3'::3’ΔHygR + 5’ΔHygR::Intron5'::Read1::CCGCACTCTAAGATC::Read2::Intron3'::3’ΔHygR + 5’ΔHygR::Intron5'::Read1::TGCCCATCTTATCAT::Read2::Intron3'::3’ΔHygR + 5’ΔHygR::Intron5'::Read1::TTACAATGTCACTCC::Read2::Intron3'::3’ΔHygR + 5’ΔHygR::Intron5'::Read1::AGACTTTTTTAGATT::Read2::Intron3'::3’ΔHygR + 5’ΔHygR::Intron5'::Read1::ATTCCTTATTACGCA::Read2::Intron3'::3’ΔHygR + 5’ΔHygR::Intron5'::Read1::ATTCTGTCTTATCCT::Read2::Intron3'::3’ΔHygR + 5’ΔHygR::Intron5'::Read1::GCCCTTTGTCAGTGA::Read2::Intron3'::3’ΔHygR + 5’ΔHygR::Intron5'::Read1::TCCCGGTCTAAGTCG::Read2::Intron3'::3’ΔHygR + 5’ΔHygR::Intron5'::Read1::TCGCGGTCTTAAAAA::Read2::Intron3'::3’ΔHygR + 5’ΔHygR::Intron5'::Read1::CTCCATTGTGATCAA::Read2::Intron3'::3’ΔHygR + 5’ΔHygR::Intron5'::Read1::CTGCTGTCTTAATGT::Read2::Intron3'::3’ΔHygR + 5’ΔHygR::Intron5'::Read1::GACCATTATAACAGT::Read2::Intron3'::3’ΔHygR + 5’ΔHygR::Intron5'::Read1::TGTCTATATCATCCA::Read2::Intron3'::3’ΔHygR + 5’ΔHygR::Intron5'::Read1::AGGCGTTTTAATTTA::Read2::Intron3'::3’ΔHygR + 5’ΔHygR::Intron5'::Read1::CCCCGCTTTCAAGTA::Read2::Intron3'::3’ΔHygR + 5’ΔHygR::Intron5'::Read1::CTCCGTTGTAATCAT::Read2::Intron3'::3’ΔHygR + 5’ΔHygR::Intron5'::Read1::GATCAATCTAAACGC::Read2::Intron3'::3’ΔHygR + 5’ΔHygR::Intron5'::Read1::TATCAATTTAAGTGC::Read2::Intron3'::3’ΔHygR + 5’ΔHygR::Intron5'::Read1::TTACACTTTCAACCG::Read2::Intron3'::3’ΔHygR + 5’ΔHygR::Intron5'::Read1::ACGCACTGTTATACT::Read2::Intron3'::3’ΔHygR + 5’ΔHygR::Intron5'::Read1::CAACCCTTTTACTGC::Read2::Intron3'::3’ΔHygR + 5’ΔHygR::Intron5'::Read1::CGGCTGTGTCAATTA::Read2::Intron3'::3’ΔHygR + 5’ΔHygR::Intron5'::Read1::GGTCCCTTTCACCAG::Read2::Intron3'::3’ΔHygR + 5’ΔHygR::Intron5'::Read1::TCTCATTCTTATACG::Read2::Intron3'::3’ΔHygR + 5’ΔHygR::Intron5'::Read1::ATGCCCTCTTAAGCG::Read2::Intron3'::3’ΔHygR + 5’ΔHygR::Intron5'::Read1::TACCTTTATGATCTT::Read2::Intron3'::3’ΔHygR + 5’ΔHygR::Intron5'::Read1::CTACAGTCTAACAGT::Read2::Intron3'::3’ΔHygR + 5’ΔHygR::Intron5'::Read1::CTGCTTTATGAATCC::Read2::Intron3'::3’ΔHygR + 5’ΔHygR::Intron5'::Read1::CTTCCCTATAACGAT::Read2::Intron3'::3’ΔHygR + 5’ΔHygR::Intron5'::Read1::GGTCGCTCTAATGCG::Read2::Intron3'::3’ΔHygR + 5’ΔHygR::Intron5'::Read1::TCCCCGTCTTACCCC::Read2::Intron3'::3’ΔHygR + 5’ΔHygR::Intron5'::Read1::TGACTTTCTCATGTT::Read2::Intron3'::3’ΔHygR + 5’ΔHygR::Intron5'::Read1::TTTCCTTATAAGCGC::Read2::Intron3'::3’ΔHygR + 5’ΔHygR::Intron5'::Read1::AAACCTTGTTACACA::Read2::Intron3'::3’ΔHygR + 5’ΔHygR::Intron5'::Read1::AAGCACTATCACCTA::Read2::Intron3'::3’ΔHygR + 5’ΔHygR::Intron5'::Read1::AGCCAATCTCATCTA::Read2::Intron3'::3’ΔHygR + 5’ΔHygR::Intron5'::Read1::CATCGTTTTTACTAC::Read2::Intron3'::3’ΔHygR + 5’ΔHygR::Intron5'::Read1::GAACTCTTTGACCTA::Read2::Intron3'::3’ΔHygR + 5’ΔHygR::Intron5'::Read1::GGTCTCTATTAAACA::Read2::Intron3'::3’ΔHygR + 5’ΔHygR::Intron5'::Read1::ATTCCCTCTGATTCC::Read2::Intron3'::3’ΔHygR + 5’ΔHygR::Intron5'::Read1::CATCTTTCTCACCTG::Read2::Intron3'::3’ΔHygR + 5’ΔHygR::Intron5'::Read1::CCGCAATCTTACTGT::Read2::Intron3'::3’ΔHygR + 5’ΔHygR::Intron5'::Read1::GATCTTTGTCAGTAC::Read2::Intron3'::3’ΔHygR + 5’ΔHygR::Intron5'::Read1::ATGCTTTTTCAATAA::Read2::Intron3'::3’ΔHygR + 5’ΔHygR::Intron5'::Read1::TAGCTGTTTCACAAT::Read2::Intron3'::3’ΔHygR + 5’ΔHygR::Intron5'::Read1::TATCCGTTTAACCAT::Read2::Intron3'::3’ΔHygR + 5’ΔHygR::Intron5'::Read1::TTTCGCTGTTAGCCT::Read2::Intron3'::3’ΔHygR + 5’ΔHygR::Intron5'::Read1::ATCCTTTCTCAGCAG::Read2::Intron3'::3’ΔHygR + 5’ΔHygR::Intron5'::Read1::ATTCTCTGTTACAGC::Read2::Intron3'::3’ΔHygR + 5’ΔHygR::Intron5'::Read1::CATCGTTCTTAACTT::Read2::Intron3'::3’ΔHygR + 5’ΔHygR::Intron5'::Read1::CCCCGATATTACTTC::Read2::Intron3'::3’ΔHygR + 5’ΔHygR::Intron5'::Read1::CTTCCATCTTAGTTT::Read2::Intron3'::3’ΔHygR + 5’ΔHygR::Intron5'::Read1::GTGCACTTTTAACAA::Read2::Intron3'::3’ΔHygR + 5’ΔHygR::Intron5'::Read1::AAGCACTCTTAGACC::Read2::Intron3'::3’ΔHygR + 5’ΔHygR::Intron5'::Read1::ATTCCTTATCAGATC::Read2::Intron3'::3’ΔHygR + 5’ΔHygR::Intron5'::Read1::CAACGGTCTGAACTC::Read2::Intron3'::3’ΔHygR + 5’ΔHygR::Intron5'::Read1::CATCCCTCTTATTCA::Read2::Intron3'::3’ΔHygR + 5’ΔHygR::Intron5'::Read1::CCACAGTGTAAGGCA::Read2::Intron3'::3’ΔHygR + 5’ΔHygR::Intron5'::Read1::TCACCTTTTTAGCAC::Read2::Intron3'::3’ΔHygR + 5’ΔHygR::Intron5'::Read1::CGTCTATTTAATTCT::Read2::Intron3'::3’ΔHygR + 5’ΔHygR::Intron5'::Read1::CTACCCTTTCAACTC::Read2::Intron3'::3’ΔHygR + 5’ΔHygR::Intron5'::Read1::CTGCCATCTGATAAC::Read2::Intron3'::3’ΔHygR + 5’ΔHygR::Intron5'::Read1::GGTCTATTTGAAACA::Read2::Intron3'::3’ΔHygR + 5’ΔHygR::Intron5'::Read1::TTCCGTTTTAACTTT::Read2::Intron3'::3’ΔHygR + 5’ΔHygR::Intron5'::Read1::TTGCCATATCATTTA::Read2::Intron3'::3’ΔHygR + 5’ΔHygR::Intron5'::Read1::CAACCATGTAAAAAT::Read2::Intron3'::3’ΔHygR + 5’ΔHygR::Intron5'::Read1::CTTCGCTCTTAACAC::Read2::Intron3'::3’ΔHygR + 5’ΔHygR::Intron5'::Read1::TTACTCTCTAAGTGC::Read2::Intron3'::3’ΔHygR +

5’ΔHygR::Intron5'::Read1::AGCCCGTATCACCTG::Read2::Intron3'::3’ΔHygR + 5’ΔHygR::Intron5'::Read1::AGTCCTTCTAACCCC::Read2::Intron3'::3’ΔHygR + 5’ΔHygR::Intron5'::Read1::CCACACTATGATCTA::Read2::Intron3'::3’ΔHygR + 5’ΔHygR::Intron5'::Read1::CCGCCATGTCAATAG::Read2::Intron3'::3’ΔHygR + 5’ΔHygR::Intron5'::Read1::TTACTGTTTCAGTTA::Read2::Intron3'::3’ΔHygR + 5’ΔHygR::Intron5'::Read1::ACTCCTTCTAACTTC::Read2::Intron3'::3’ΔHygR + 5’ΔHygR::Intron5'::Read1::CATCAGTATGAGTTT::Read2::Intron3'::3’ΔHygR + 5’ΔHygR::Intron5'::Read1::TAACGCTATTAATGC::Read2::Intron3'::3’ΔHygR + 5’ΔHygR::Intron5'::Read1::TTGCGCTTTCAATCG::Read2::Intron3'::3’ΔHygR + 5’ΔHygR::Intron5'::Read1::TTGCTCTATTACTTC::Read2::Intron3'::3’ΔHygR + 5’ΔHygR::Intron5'::Read1::TTTCCTTCTCACCTG::Read2::Intron3'::3’ΔHygR + 5’ΔHygR::Intron5'::Read1::CACCCATATCACGCA::Read2::Intron3'::3’ΔHygR + 5’ΔHygR::Intron5'::Read1::CCTCAATCTAAATAG::Read2::Intron3'::3’ΔHygR + 5’ΔHygR::Intron5'::Read1::CTCCTATATCAACTT::Read2::Intron3'::3’ΔHygR + 5’ΔHygR::Intron5'::Read1::GAACCCTTTTAAGCA::Read2::Intron3'::3’ΔHygR + 5’ΔHygR::Intron5'::Read1::TATCATTGTTACTGG::Read2::Intron3'::3’ΔHygR + 5’ΔHygR::Intron5'::Read1::ACCCTATTTCACGTC::Read2::Intron3'::3’ΔHygR + 5’ΔHygR::Intron5'::Read1::GAGCATTATTATTCG::Read2::Intron3'::3’ΔHygR + 5’ΔHygR::Intron5'::Read1::GTTCTATTTTAATCA::Read2::Intron3'::3’ΔHygR + 5’ΔHygR::Intron5'::Read1::TCCCATTCTAAAACT::Read2::Intron3'::3’ΔHygR + 5’ΔHygR::Intron5'::Read1::TCGCACTTTGACTCG::Read2::Intron3'::3’ΔHygR + 5’ΔHygR::Intron5'::Read1::TTACTCTGTTATTAG::Read2::Intron3'::3’ΔHygR + 5’ΔHygR::Intron5'::Read1::ATTCATTTTCACCTA::Read2::Intron3'::3’ΔHygR + 5’ΔHygR::Intron5'::Read1::CGACTCTCTCAGTCC::Read2::Intron3'::3’ΔHygR + 5’ΔHygR::Intron5'::Read1::CTTCCCTTTTATTAC::Read2::Intron3'::3’ΔHygR + 5’ΔHygR::Intron5'::Read1::AGACCATTTTAAAAC::Read2::Intron3'::3’ΔHygR + 5’ΔHygR::Intron5'::Read1::AGACTCTATCAGCTT::Read2::Intron3'::3’ΔHygR + 5’ΔHygR::Intron5'::Read1::CTGCTATATCAAAAT::Read2::Intron3'::3’ΔHygR + 5’ΔHygR::Intron5'::Read1::GCACGTTTTAACATA::Read2::Intron3'::3’ΔHygR + 5’ΔHygR::Intron5'::Read1::GGACACTCTAATGAT::Read2::Intron3'::3’ΔHygR + 5’ΔHygR::Intron5'::Read1::CTACAATATTAACCA::Read2::Intron3'::3’ΔHygR + 5’ΔHygR::Intron5'::Read1::GACCACTATAAGATT::Read2::Intron3'::3’ΔHygR + 5’ΔHygR::Intron5'::Read1::GCTCAATGTCATCTT::Read2::Intron3'::3’ΔHygR + 5’ΔHygR::Intron5'::Read1::CCCCTTTTTGAAAAT::Read2::Intron3'::3’ΔHygR + 5’ΔHygR::Intron5'::Read1::CGACAATTTTACGAC::Read2::Intron3'::3’ΔHygR + 5’ΔHygR::Intron5'::Read1::TGGCGCTTTCAAACA::Read2::Intron3'::3’ΔHygR + 5’ΔHygR::Intron5'::Read1::TTCCATTCTGACAAC::Read2::Intron3'::3’ΔHygR + 5’ΔHygR::Intron5'::Read1::TTCCTTTATTACTAC::Read2::Intron3'::3’ΔHygR + 5’ΔHygR::Intron5'::Read1::AAACGCTTTCATGTC::Read2::Intron3'::3’ΔHygR + 5’ΔHygR::Intron5'::Read1::ATTCACTCTGACACT::Read2::Intron3'::3’ΔHygR + 5’ΔHygR::Intron5'::Read1::CAACCATATCATGCT::Read2::Intron3'::3’ΔHygR + 5’ΔHygR::Intron5'::Read1::CAACCTTTTAAGAAA::Read2::Intron3'::3’ΔHygR + 5’ΔHygR::Intron5'::Read1::GAGCAATGTTATTAA::Read2::Intron3'::3’ΔHygR + 5’ΔHygR::Intron5'::Read1::TTGCCGTGTCATTGC::Read2::Intron3'::3’ΔHygR + 5’ΔHygR::Intron5'::Read1::ACGCTATATTATCCA::Read2::Intron3'::3’ΔHygR + 5’ΔHygR::Intron5'::Read1::ATACCTTATCAGCGA::Read2::Intron3'::3’ΔHygR + 5’ΔHygR::Intron5'::Read1::CAGCTATGTGATAAT::Read2::Intron3'::3’ΔHygR + 5’ΔHygR::Intron5'::Read1::CCCCTTTTTAACCAA::Read2::Intron3'::3’ΔHygR + 5’ΔHygR::Intron5'::Read1::CTGCCATGTTAGTCC::Read2::Intron3'::3’ΔHygR + 5’ΔHygR::Intron5'::Read1::TATCGTTTTCATCCG::Read2::Intron3'::3’ΔHygR + 5’ΔHygR::Intron5'::Read1::TGCCTTTTTGATAAC::Read2::Intron3'::3’ΔHygR + 5’ΔHygR::Intron5'::Read1::AAACCCTTTAACATA::Read2::Intron3'::3’ΔHygR + 5’ΔHygR::Intron5'::Read1::CAGCCCTTTTACCCA::Read2::Intron3'::3’ΔHygR + 5’ΔHygR::Intron5'::Read1::CCTCATTATAATTTC::Read2::Intron3'::3’ΔHygR + 5’ΔHygR::Intron5'::Read1::GGCCACTCTAATATT::Read2::Intron3'::3’ΔHygR + 5’ΔHygR::Intron5'::Read1::GGGCACTTTTAGTGC::Read2::Intron3'::3’ΔHygR + 5’ΔHygR::Intron5'::Read1::TAACCATCTAAAACT::Read2::Intron3'::3’ΔHygR + 5’ΔHygR::Intron5'::Read1::TAGCTATCTAATATG::Read2::Intron3'::3’ΔHygR + 5’ΔHygR::Intron5'::Read1::TCACGCTGTCATCCA::Read2::Intron3'::3’ΔHygR + 5’ΔHygR::Intron5'::Read1::TTACTATTTAAGCAG::Read2::Intron3'::3’ΔHygR + 5’ΔHygR::Intron5'::Read1::TTCCAATTTCAATCA::Read2::Intron3'::3’ΔHygR + 5’ΔHygR::Intron5'::Read1::TTTCACTCTCAATTA::Read2::Intron3'::3’ΔHygR + 5’ΔHygR::Intron5'::Read1::ATACCCTTTAATTCG::Read2::Intron3'::3’ΔHygR + 5’ΔHygR::Intron5'::Read1::CTTCGATCTCACTAG::Read2::Intron3'::3’ΔHygR + 5’ΔHygR::Intron5'::Read1::GGCCGCTTTAATAAC::Read2::Intron3'::3’ΔHygR + 5’ΔHygR::Intron5'::Read1::GTGCGTTATAATTAT::Read2::Intron3'::3’ΔHygR + 5’ΔHygR::Intron5'::Read1::TCGCTCTATCATCTA::Read2::Intron3'::3’ΔHygR + 5’ΔHygR::Intron5'::Read1::TTCCCCTGTAACTTC::Read2::Intron3'::3’ΔHygR + 5’ΔHygR::Intron5'::Read1::ACACTTTGTGATTAG::Read2::Intron3'::3’ΔHygR + 5’ΔHygR::Intron5'::Read1::ACGCTATATTAAGGC::Read2::Intron3'::3’ΔHygR +

5’ΔHygR::Intron5'::Read1::ATTCGATGTGACAGC::Read2::Intron3'::3’ΔHygR + 5’ΔHygR::Intron5'::Read1::CGGCGGTATCAGATT::Read2::Intron3'::3’ΔHygR + 5’ΔHygR::Intron5'::Read1::CTCCCTTCTAAGCTT::Read2::Intron3'::3’ΔHygR + 5’ΔHygR::Intron5'::Read1::TAACCATCTTATCCC::Read2::Intron3'::3’ΔHygR + 5’ΔHygR::Intron5'::Read1::TTTCTGTCTAATATC::Read2::Intron3'::3’ΔHygR + 5’ΔHygR::Intron5'::Read1::ACGCAGTCTAAATTA::Read2::Intron3'::3’ΔHygR + 5’ΔHygR::Intron5'::Read1::ATCCAATATCACGGG::Read2::Intron3'::3’ΔHygR + 5’ΔHygR::Intron5'::Read1::CTACCCTATCATGCA::Read2::Intron3'::3’ΔHygR + 5’ΔHygR::Intron5'::Read1::GTCCATTCTAATGTC::Read2::Intron3'::3’ΔHygR + 5’ΔHygR::Intron5'::Read1::TCCCAATGTTAACGA::Read2::Intron3'::3’ΔHygR + 5’ΔHygR::Intron5'::Read1::TGACATTCTTAATGA::Read2::Intron3'::3’ΔHygR + 5’ΔHygR::Intron5'::Read1::TTTCCGTATGATACG::Read2::Intron3'::3’ΔHygR + 5’ΔHygR::Intron5'::Read1::AAGCCTTTTAATCCC::Read2::Intron3'::3’ΔHygR + 5’ΔHygR::Intron5'::Read1::ATACAATTTTACGAC::Read2::Intron3'::3’ΔHygR + 5’ΔHygR::Intron5'::Read1::CAACAATCTAATCTA::Read2::Intron3'::3’ΔHygR + 5’ΔHygR::Intron5'::Read1::CATCACTTTCAGCGA::Read2::Intron3'::3’ΔHygR + 5’ΔHygR::Intron5'::Read1::CGCCCATCTTACTTG::Read2::Intron3'::3’ΔHygR + 5’ΔHygR::Intron5'::Read1::CTACTATGTCAAATT::Read2::Intron3'::3’ΔHygR + 5’ΔHygR::Intron5'::Read1::CTACTCTATGATGGA::Read2::Intron3'::3’ΔHygR + 5’ΔHygR::Intron5'::Read1::GTCCCTTATAAGCTG::Read2::Intron3'::3’ΔHygR + 5’ΔHygR::Intron5'::Read1::TCGCCTTGTCAGATC::Read2::Intron3'::3’ΔHygR + 5’ΔHygR::Intron5'::Read1::ACTCATTGTAAGGGC::Read2::Intron3'::3’ΔHygR + 5’ΔHygR::Intron5'::Read1::CATCCTTGTCATAAT::Read2::Intron3'::3’ΔHygR + 5’ΔHygR::Intron5'::Read1::GATCTGTTTCAATAA::Read2::Intron3'::3’ΔHygR + 5’ΔHygR::Intron5'::Read1::GCTCAATGTCAGACT::Read2::Intron3'::3’ΔHygR + 5’ΔHygR::Intron5'::Read1::GGACTGTATTACCAT::Read2::Intron3'::3’ΔHygR + 5’ΔHygR::Intron5'::Read1::TGACCCTTTTAATAC::Read2::Intron3'::3’ΔHygR + 5’ΔHygR::Intron5'::Read1::CACCTCTTTAACGCC::Read2::Intron3'::3’ΔHygR + 5’ΔHygR::Intron5'::Read1::CATCTATTTTACACA::Read2::Intron3'::3’ΔHygR + 5’ΔHygR::Intron5'::Read1::CCGCAGTCTAACGCA::Read2::Intron3'::3’ΔHygR + 5’ΔHygR::Intron5'::Read1::GCGCGTTCTAAGTAA::Read2::Intron3'::3’ΔHygR + 5’ΔHygR::Intron5'::Read1::TTGCCATCTTAGACT::Read2::Intron3'::3’ΔHygR + 5’ΔHygR::Intron5'::Read1::AAACATTTTAAGACT::Read2::Intron3'::3’ΔHygR + 5’ΔHygR::Intron5'::Read1::AACCCCTGTGATAAT::Read2::Intron3'::3’ΔHygR + 5’ΔHygR::Intron5'::Read1::CTTCTCTTTCAAATA::Read2::Intron3'::3’ΔHygR + 5’ΔHygR::Intron5'::Read1::GTACATTATAAGACT::Read2::Intron3'::3’ΔHygR + 5’ΔHygR::Intron5'::Read1::TACCCGTTTCACGCA::Read2::Intron3'::3’ΔHygR + 5’ΔHygR::Intron5'::Read1::TCTCAGTTTGACAAA::Read2::Intron3'::3’ΔHygR + 5’ΔHygR::Intron5'::Read1::AAACTATATCACTTT::Read2::Intron3'::3’ΔHygR + 5’ΔHygR::Intron5'::Read1::GAGCGCTATGAATAC::Read2::Intron3'::3’ΔHygR + 5’ΔHygR::Intron5'::Read1::GTACCCTTTTAAGTT::Read2::Intron3'::3’ΔHygR + 5’ΔHygR::Intron5'::Read1::GTCCCCTTTCACAGT::Read2::Intron3'::3’ΔHygR + 5’ΔHygR::Intron5'::Read1::GTCCCGTGTGATTGC::Read2::Intron3'::3’ΔHygR + 5’ΔHygR::Intron5'::Read1::TCACCTTATAAGTAC::Read2::Intron3'::3’ΔHygR + 5’ΔHygR::Intron5'::Read1::TTGCTCTATCACCCG::Read2::Intron3'::3’ΔHygR + 5’ΔHygR::Intron5'::Read1::AATCGGTCTGATACC::Read2::Intron3'::3’ΔHygR + 5’ΔHygR::Intron5'::Read1::ACTCTTTATGATACG::Read2::Intron3'::3’ΔHygR + 5’ΔHygR::Intron5'::Read1::AGGCGTTGTCAATAA::Read2::Intron3'::3’ΔHygR + 5’ΔHygR::Intron5'::Read1::ATACTATCTAACAAC::Read2::Intron3'::3’ΔHygR + 5’ΔHygR::Intron5'::Read1::ATTCCTTGTAAATCA::Read2::Intron3'::3’ΔHygR + 5’ΔHygR::Intron5'::Read1::CAACATTATCAATAA::Read2::Intron3'::3’ΔHygR + 5’ΔHygR::Intron5'::Read1::CAGCAGTTTTAAACG::Read2::Intron3'::3’ΔHygR + 5’ΔHygR::Intron5'::Read1::CAGCCTTCTGACAGA::Read2::Intron3'::3’ΔHygR + 5’ΔHygR::Intron5'::Read1::CATCTATATTATCAC::Read2::Intron3'::3’ΔHygR + 5’ΔHygR::Intron5'::Read1::CCCCTCTTTCATTAT::Read2::Intron3'::3’ΔHygR + 5’ΔHygR::Intron5'::Read1::CTTCTATTTTATTAA::Read2::Intron3'::3’ΔHygR + 5’ΔHygR::Intron5'::Read1::GCTCAATATCATCAT::Read2::Intron3'::3’ΔHygR + 5’ΔHygR::Intron5'::Read1::GCTCCATTTCAGGTC::Read2::Intron3'::3’ΔHygR + 5’ΔHygR::Intron5'::Read1::GTACTCTTTGATCCT::Read2::Intron3'::3’ΔHygR + 5’ΔHygR::Intron5'::Read1::TATCAATCTCAATTA::Read2::Intron3'::3’ΔHygR + 5’ΔHygR::Intron5'::Read1::TATCGGTTTTACTGC::Read2::Intron3'::3’ΔHygR + 5’ΔHygR::Intron5'::Read1::CACCCATTTGACTGA::Read2::Intron3'::3’ΔHygR + 5’ΔHygR::Intron5'::Read1::CATCGCTGTGATACC::Read2::Intron3'::3’ΔHygR + 5’ΔHygR::Intron5'::Read1::CGCCGATTTCATGCC::Read2::Intron3'::3’ΔHygR + 5’ΔHygR::Intron5'::Read1::CGTCCGTTTTATTCA::Read2::Intron3'::3’ΔHygR + 5’ΔHygR::Intron5'::Read1::GTTCCATTTAAGTCG::Read2::Intron3'::3’ΔHygR + 5’ΔHygR::Intron5'::Read1::TAACCATGTTAAACC::Read2::Intron3'::3’ΔHygR + 5’ΔHygR::Intron5'::Read1::TATCGTTTTTAGCAT::Read2::Intron3'::3’ΔHygR + 5’ΔHygR::Intron5'::Read1::TCACTATGTAACATT::Read2::Intron3'::3’ΔHygR + 5’ΔHygR::Intron5'::Read1::TGCCATTATTACTAT::Read2::Intron3'::3’ΔHygR +

5’ΔHygR::Intron5'::Read1::ACCCTATCTGAACCC::Read2::Intron3'::3’ΔHygR + 5’ΔHygR::Intron5'::Read1::CACCTTTCTCAATTA::Read2::Intron3'::3’ΔHygR + 5’ΔHygR::Intron5'::Read1::CCACCCTTTCACTTA::Read2::Intron3'::3’ΔHygR + 5’ΔHygR::Intron5'::Read1::CTCCCCTCTAACTCT::Read2::Intron3'::3’ΔHygR + 5’ΔHygR::Intron5'::Read1::CTTCGTTCTTACACA::Read2::Intron3'::3’ΔHygR + 5’ΔHygR::Intron5'::Read1::GAACTATTTCACACT::Read2::Intron3'::3’ΔHygR + 5’ΔHygR::Intron5'::Read1::GTACACTTTTACTTC::Read2::Intron3'::3’ΔHygR + 5’ΔHygR::Intron5'::Read1::TAACTCTGTAAGTCT::Read2::Intron3'::3’ΔHygR + 5’ΔHygR::Intron5'::Read1::TCCCTGTGTAAAACC::Read2::Intron3'::3’ΔHygR + 5’ΔHygR::Intron5'::Read1::TCGCCATGTTAAGGT::Read2::Intron3'::3’ΔHygR + 5’ΔHygR::Intron5'::Read1::TTTCACTCTGAAATC::Read2::Intron3'::3’ΔHygR + 5’ΔHygR::Intron5'::Read1::AATCTTTGTTACCGT::Read2::Intron3'::3’ΔHygR + 5’ΔHygR::Intron5'::Read1::ATACAGTTTCACTTA::Read2::Intron3'::3’ΔHygR + 5’ΔHygR::Intron5'::Read1::ATCCGTTATTAATAC::Read2::Intron3'::3’ΔHygR + 5’ΔHygR::Intron5'::Read1::ATTCATTATCAGAGC::Read2::Intron3'::3’ΔHygR + 5’ΔHygR::Intron5'::Read1::GATCACTGTGATGTA::Read2::Intron3'::3’ΔHygR + 5’ΔHygR::Intron5'::Read1::GCACACTCTGATCGT::Read2::Intron3'::3’ΔHygR + 5’ΔHygR::Intron5'::Read1::GCACTGTGTCACATA::Read2::Intron3'::3’ΔHygR + 5’ΔHygR::Intron5'::Read1::GCGCAGTTTTATCCA::Read2::Intron3'::3’ΔHygR + 5’ΔHygR::Intron5'::Read1::GTGCTTTTTGACTGG::Read2::Intron3'::3’ΔHygR + 5’ΔHygR::Intron5'::Read1::TAACCCTGTCACTCT::Read2::Intron3'::3’ΔHygR + 5’ΔHygR::Intron5'::Read1::TACCAATCTCACTCA::Read2::Intron3'::3’ΔHygR + 5’ΔHygR::Intron5'::Read1::TAGCACTTTCACTAT::Read2::Intron3'::3’ΔHygR + 5’ΔHygR::Intron5'::Read1::TCACCTTGTAATACT::Read2::Intron3'::3’ΔHygR + 5’ΔHygR::Intron5'::Read1::AAACAATATAACGGA::Read2::Intron3'::3’ΔHygR + 5’ΔHygR::Intron5'::Read1::AATCATTTTCATATG::Read2::Intron3'::3’ΔHygR + 5’ΔHygR::Intron5'::Read1::ATGCTGTGTTAACCA::Read2::Intron3'::3’ΔHygR + 5’ΔHygR::Intron5'::Read1::CCCCAGTCTAATCAA::Read2::Intron3'::3’ΔHygR + 5’ΔHygR::Intron5'::Read1::GATCTATATCAACTA::Read2::Intron3'::3’ΔHygR + 5’ΔHygR::Intron5'::Read1::GTACAATATTATCCT::Read2::Intron3'::3’ΔHygR + 5’ΔHygR::Intron5'::Read1::TAACATTTTTAGTCA::Read2::Intron3'::3’ΔHygR + 5’ΔHygR::Intron5'::Read1::TCTCAGTCTGAACGC::Read2::Intron3'::3’ΔHygR + 5’ΔHygR::Intron5'::Read1::TGCCTGTTTAAATTA::Read2::Intron3'::3’ΔHygR + 5’ΔHygR::Intron5'::Read1::AACCCTTATAAGCCA::Read2::Intron3'::3’ΔHygR + 5’ΔHygR::Intron5'::Read1::CACCAGTGTGATAAT::Read2::Intron3'::3’ΔHygR + 5’ΔHygR::Intron5'::Read1::CCACCTTTTCACTAT::Read2::Intron3'::3’ΔHygR + 5’ΔHygR::Intron5'::Read1::CTTCACTTTAATTCT::Read2::Intron3'::3’ΔHygR + 5’ΔHygR::Intron5'::Read1::TTCCAATGTTACATT::Read2::Intron3'::3’ΔHygR + 5’ΔHygR::Intron5'::Read1::CACCCGTTTGACTAA::Read2::Intron3'::3’ΔHygR + 5’ΔHygR::Intron5'::Read1::CGACATTCTCACTTT::Read2::Intron3'::3’ΔHygR + 5’ΔHygR::Intron5'::Read1::GGTCTCTGTTAACTG::Read2::Intron3'::3’ΔHygR + 5’ΔHygR::Intron5'::Read1::TAACCTTTTTAGCAT::Read2::Intron3'::3’ΔHygR + 5’ΔHygR::Intron5'::Read1::TACCTCTATTACTGT::Read2::Intron3'::3’ΔHygR + 5’ΔHygR::Intron5'::Read1::TGGCCTTTTAAATCC::Read2::Intron3'::3’ΔHygR + 5’ΔHygR::Intron5'::Read1::TTCCACTATTAGTCT::Read2::Intron3'::3’ΔHygR + 5’ΔHygR::Intron5'::Read1::CCTCAATATTACTTT::Read2::Intron3'::3’ΔHygR + 5’ΔHygR::Intron5'::Read1::GGACTCTTTCACGCT::Read2::Intron3'::3’ΔHygR + 5’ΔHygR::Intron5'::Read1::GTGCTTTATGATATT::Read2::Intron3'::3’ΔHygR + 5’ΔHygR::Intron5'::Read1::TGACCTTTTCATCGT::Read2::Intron3'::3’ΔHygR + 5’ΔHygR::Intron5'::Read1::CTGCAATCTTATGCC::Read2::Intron3'::3’ΔHygR + 5’ΔHygR::Intron5'::Read1::GATCATTATGACTAA::Read2::Intron3'::3’ΔHygR + 5’ΔHygR::Intron5'::Read1::TAGCTTTTTAACATA::Read2::Intron3'::3’ΔHygR + 5’ΔHygR::Intron5'::Read1::TCACCCTATCAAGAT::Read2::Intron3'::3’ΔHygR + 5’ΔHygR::Intron5'::Read1::TTACATTGTAACATA::Read2::Intron3'::3’ΔHygR + 5’ΔHygR::Intron5'::Read1::AGCCCATATGATGTA::Read2::Intron3'::3’ΔHygR + 5’ΔHygR::Intron5'::Read1::CGCCCATCTCACTAC::Read2::Intron3'::3’ΔHygR + 5’ΔHygR::Intron5'::Read1::TAACCATTTGACTGC::Read2::Intron3'::3’ΔHygR + 5’ΔHygR::Intron5'::Read1::TATCTTTTTAACCCA::Read2::Intron3'::3’ΔHygR + 5’ΔHygR::Intron5'::Read1::TTTCCGTTTTACATT::Read2::Intron3'::3’ΔHygR + 5’ΔHygR::Intron5'::Read1::AATCCTTATCACTGG::Read2::Intron3'::3’ΔHygR + 5’ΔHygR::Intron5'::Read1::ATACTATTTTATTCA::Read2::Intron3'::3’ΔHygR + 5’ΔHygR::Intron5'::Read1::CCACTTTTTCAATCG::Read2::Intron3'::3’ΔHygR + 5’ΔHygR::Intron5'::Read1::CCGCCATCTTAACAA::Read2::Intron3'::3’ΔHygR + 5’ΔHygR::Intron5'::Read1::CCTCTGTCTGACATC::Read2::Intron3'::3’ΔHygR + 5’ΔHygR::Intron5'::Read1::CGACCCTGTAAGCCC::Read2::Intron3'::3’ΔHygR + 5’ΔHygR::Intron5'::Read1::GCACTGTTTCGTTAT::Read2::Intron3'::3’ΔHygR + 5’ΔHygR::Intron5'::Read1::GTACCTTATCACAAT::Read2::Intron3'::3’ΔHygR + 5’ΔHygR::Intron5'::Read1::TCCCTCTGTTATTTA::Read2::Intron3'::3’ΔHygR + 5’ΔHygR::Intron5'::Read1::TGACTGTTTAATAAA::Read2::Intron3'::3’ΔHygR + 5’ΔHygR::Intron5'::Read1::CTACTGTCTAAAGGG::Read2::Intron3'::3’ΔHygR +

5’ΔHygR::Intron5'::Read1::GATCCTTCTAAAATC::Read2::Intron3'::3’ΔHygR + 5’ΔHygR::Intron5'::Read1::GGTCAATTTCACTCT::Read2::Intron3'::3’ΔHygR + 5’ΔHygR::Intron5'::Read1::GTCCTGTGTAACTCC::Read2::Intron3'::3’ΔHygR + 5’ΔHygR::Intron5'::Read1::TGCCTTTCTTATAAA::Read2::Intron3'::3’ΔHygR + 5’ΔHygR::Intron5'::Read1::TTACGCTTTGAAATC::Read2::Intron3'::3’ΔHygR + 5’ΔHygR::Intron5'::Read1::AAACTCTTTAAGCAA::Read2::Intron3'::3’ΔHygR + 5’ΔHygR::Intron5'::Read1::GCACTATTTTAGATC::Read2::Intron3'::3’ΔHygR + 5’ΔHygR::Intron5'::Read1::GCTCCGTCTCAGTGA::Read2::Intron3'::3’ΔHygR + 5’ΔHygR::Intron5'::Read1::TTCCTTTATCAATGT::Read2::Intron3'::3’ΔHygR + 5’ΔHygR::Intron5'::Read1::AGCCTGTGTCAAGCA::Read2::Intron3'::3’ΔHygR + 5’ΔHygR::Intron5'::Read1::ATACATTTTGACCCC::Read2::Intron3'::3’ΔHygR + 5’ΔHygR::Intron5'::Read1::ATACCATGTGACTTC::Read2::Intron3'::3’ΔHygR + 5’ΔHygR::Intron5'::Read1::CCACGCTCTCAATAA::Read2::Intron3'::3’ΔHygR + 5’ΔHygR::Intron5'::Read1::CTCCAATTTTATAAG::Read2::Intron3'::3’ΔHygR + 5’ΔHygR::Intron5'::Read1::GCACTTTCTTACTTG::Read2::Intron3'::3’ΔHygR + 5’ΔHygR::Intron5'::Read1::GCGCTCTTTTAAAAT::Read2::Intron3'::3’ΔHygR + 5’ΔHygR::Intron5'::Read1::GGCCGGTATTAACCA::Read2::Intron3'::3’ΔHygR + 5’ΔHygR::Intron5'::Read1::GGTCTTTATTATTTG::Read2::Intron3'::3’ΔHygR + 5’ΔHygR::Intron5'::Read1::CGCCAATTTAATTGC::Read2::Intron3'::3’ΔHygR + 5’ΔHygR::Intron5'::Read1::GAACACTGTCAAATG::Read2::Intron3'::3’ΔHygR + 5’ΔHygR::Intron5'::Read1::GAGCCCTTTCACAGT::Read2::Intron3'::3’ΔHygR + 5’ΔHygR::Intron5'::Read1::GATCCATATCACTTA::Read2::Intron3'::3’ΔHygR + 5’ΔHygR::Intron5'::Read1::TCACGCTTTGAGCAT::Read2::Intron3'::3’ΔHygR + 5’ΔHygR::Intron5'::Read1::TCTCATTGTCAGTGC::Read2::Intron3'::3’ΔHygR + 5’ΔHygR::Intron5'::Read1::ACACGCTATCACAGA::Read2::Intron3'::3’ΔHygR + 5’ΔHygR::Intron5'::Read1::ACGCTTTCTCATTAC::Read2::Intron3'::3’ΔHygR + 5’ΔHygR::Intron5'::Read1::ATTCGTTCTTACACC::Read2::Intron3'::3’ΔHygR + 5’ΔHygR::Intron5'::Read1::CAACTTTCTCATAAC::Read2::Intron3'::3’ΔHygR + 5’ΔHygR::Intron5'::Read1::CCACCCTTTCATCCG::Read2::Intron3'::3’ΔHygR + 5’ΔHygR::Intron5'::Read1::TATCGATCTCATAGG::Read2::Intron3'::3’ΔHygR + 5’ΔHygR::Intron5'::Read1::TATCTGTCTAAACCC::Read2::Intron3'::3’ΔHygR + 5’ΔHygR::Intron5'::Read1::TCCCCATGTCAATAC::Read2::Intron3'::3’ΔHygR + 5’ΔHygR::Intron5'::Read1::TCGCATTTTTATTTC::Read2::Intron3'::3’ΔHygR + 5’ΔHygR::Intron5'::Read1::TGGCAATATTAGCTC::Read2::Intron3'::3’ΔHygR + 5’ΔHygR::Intron5'::Read1::CAACACTATTATATA::Read2::Intron3'::3’ΔHygR + 5’ΔHygR::Intron5'::Read1::CACCCCTGTAAAACC::Read2::Intron3'::3’ΔHygR + 5’ΔHygR::Intron5'::Read1::CCCCAGTCTCACCCT::Read2::Intron3'::3’ΔHygR + 5’ΔHygR::Intron5'::Read1::CGCCAGTTTAATTCC::Read2::Intron3'::3’ΔHygR + 5’ΔHygR::Intron5'::Read1::CTCCGCTATTACATC::Read2::Intron3'::3’ΔHygR + 5’ΔHygR::Intron5'::Read1::GATCAATCTCAACCA::Read2::Intron3'::3’ΔHygR + 5’ΔHygR::Intron5'::Read1::GCTCACTTTGAATTG::Read2::Intron3'::3’ΔHygR + 5’ΔHygR::Intron5'::Read1::GTTCGCTCTCAGGAC::Read2::Intron3'::3’ΔHygR + 5’ΔHygR::Intron5'::Read1::TGGCCTTTTCATTCA::Read2::Intron3'::3’ΔHygR + 5’ΔHygR::Intron5'::Read1::ACGCAATATCAAATT::Read2::Intron3'::3’ΔHygR + 5’ΔHygR::Intron5'::Read1::AGACGTTATAACACG::Read2::Intron3'::3’ΔHygR + 5’ΔHygR::Intron5'::Read1::ATCCAGTTTCATACG::Read2::Intron3'::3’ΔHygR + 5’ΔHygR::Intron5'::Read1::GAACCGTTTAATTAA::Read2::Intron3'::3’ΔHygR + 5’ΔHygR::Intron5'::Read1::GCGCAATCTCATCAA::Read2::Intron3'::3’ΔHygR + 5’ΔHygR::Intron5'::Read1::GCTCTGTTTGACGTA::Read2::Intron3'::3’ΔHygR + 5’ΔHygR::Intron5'::Read1::GGGCACTCTCAACTT::Read2::Intron3'::3’ΔHygR + 5’ΔHygR::Intron5'::Read1::GTCCTTTGTTAAAAA::Read2::Intron3'::3’ΔHygR + 5’ΔHygR::Intron5'::Read1::ACCCGCTTTAAACGT::Read2::Intron3'::3’ΔHygR + 5’ΔHygR::Intron5'::Read1::AGACGTTGTCATAGT::Read2::Intron3'::3’ΔHygR + 5’ΔHygR::Intron5'::Read1::CTACCCTGTAATAAA::Read2::Intron3'::3’ΔHygR + 5’ΔHygR::Intron5'::Read1::TCCCCCTGTAATAGT::Read2::Intron3'::3’ΔHygR + 5’ΔHygR::Intron5'::Read1::AATCACTGTTACGCA::Read2::Intron3'::3’ΔHygR + 5’ΔHygR::Intron5'::Read1::ACGCGGTATAAAATT::Read2::Intron3'::3’ΔHygR + 5’ΔHygR::Intron5'::Read1::ATTCTCTCTCATAAA::Read2::Intron3'::3’ΔHygR + 5’ΔHygR::Intron5'::Read1::CAGCCCTTTGAGTGA::Read2::Intron3'::3’ΔHygR + 5’ΔHygR::Intron5'::Read1::CATCGCTTTGATTAC::Read2::Intron3'::3’ΔHygR + 5’ΔHygR::Intron5'::Read1::CCACCGTATAACACT::Read2::Intron3'::3’ΔHygR + 5’ΔHygR::Intron5'::Read1::CCTCGGTCTCAAGGA::Read2::Intron3'::3’ΔHygR + 5’ΔHygR::Intron5'::Read1::CTCCATTATGAGCTA::Read2::Intron3'::3’ΔHygR + 5’ΔHygR::Intron5'::Read1::GCGCCCTTTTATAGT::Read2::Intron3'::3’ΔHygR + 5’ΔHygR::Intron5'::Read1::AACCGCTATAAGGCT::Read2::Intron3'::3’ΔHygR + 5’ΔHygR::Intron5'::Read1::AATCGTTGTCAATCG::Read2::Intron3'::3’ΔHygR + 5’ΔHygR::Intron5'::Read1::ACTCGATGTTAGTTT::Read2::Intron3'::3’ΔHygR + 5’ΔHygR::Intron5'::Read1::CCCCCTTATCACGAC::Read2::Intron3'::3’ΔHygR + 5’ΔHygR::Intron5'::Read1::TCGCCCTCTAATGAG::Read2::Intron3'::3’ΔHygR + 5’ΔHygR::Intron5'::Read1::AAACTCTCTCATCCA::Read2::Intron3'::3’ΔHygR +

5’ΔHygR::Intron5'::Read1::ACCCGTTTTTAACCG::Read2::Intron3'::3’ΔHygR + 5’ΔHygR::Intron5'::Read1::ATGCCTTATTATTCC::Read2::Intron3'::3’ΔHygR + 5’ΔHygR::Intron5'::Read1::CTCCACTTTTATGTG::Read2::Intron3'::3’ΔHygR + 5’ΔHygR::Intron5'::Read1::CTCCTCTCTAAAACT::Read2::Intron3'::3’ΔHygR + 5’ΔHygR::Intron5'::Read1::TATCACTTTTACACC::Read2::Intron3'::3’ΔHygR + 5’ΔHygR::Intron5'::Read1::TGACCATGTTATTGC::Read2::Intron3'::3’ΔHygR + 5’ΔHygR::Intron5'::Read1::AATCTTTGTAATTCG::Read2::Intron3'::3’ΔHygR + 5’ΔHygR::Intron5'::Read1::TACCCATATAAGACG::Read2::Intron3'::3’ΔHygR + 5’ΔHygR::Intron5'::Read1::TCCCTTTGTCATCCT::Read2::Intron3'::3’ΔHygR + 5’ΔHygR::Intron5'::Read1::TGTCGCTATTATCTA::Read2::Intron3'::3’ΔHygR + 5’ΔHygR::Intron5'::Read1::TTCCCTTATTAAACA::Read2::Intron3'::3’ΔHygR + 5’ΔHygR::Intron5'::Read1::CATCACTTTCACTAA::Read2::Intron3'::3’ΔHygR + 5’ΔHygR::Intron5'::Read1::CTCCGCTTTAAATAT::Read2::Intron3'::3’ΔHygR + 5’ΔHygR::Intron5'::Read1::GATTTGACAGTGGGT::Read2::Intron3'::3’ΔHygR + 5’ΔHygR::Intron5'::Read1::GGACATTATCAATCC::Read2::Intron3'::3’ΔHygR + 5’ΔHygR::Intron5'::Read1::TCTCCATATCATACC::Read2::Intron3'::3’ΔHygR + 5’ΔHygR::Intron5'::Read1::TCTCCATCTGACCAA::Read2::Intron3'::3’ΔHygR + 5’ΔHygR::Intron5'::Read1::TCTCCGTTTAACCAT::Read2::Intron3'::3’ΔHygR + 5’ΔHygR::Intron5'::Read1::TGTCCCTATAACAAC::Read2::Intron3'::3’ΔHygR + 5’ΔHygR::Intron5'::Read1::TTACCTTATCATTGA::Read2::Intron3'::3’ΔHygR + 5’ΔHygR::Intron5'::Read1::AATCTATTTTAAGCA::Read2::Intron3'::3’ΔHygR + 5’ΔHygR::Intron5'::Read1::ACCCGTTTTTATTGA::Read2::Intron3'::3’ΔHygR + 5’ΔHygR::Intron5'::Read1::ATTCCGTTTAATCAC::Read2::Intron3'::3’ΔHygR + 5’ΔHygR::Intron5'::Read1::CTTCTATTTCACTCC::Read2::Intron3'::3’ΔHygR + 5’ΔHygR::Intron5'::Read1::GCCCAATTTAACAAG::Read2::Intron3'::3’ΔHygR + 5’ΔHygR::Intron5'::Read1::GCCCGGTATCACTCC::Read2::Intron3'::3’ΔHygR + 5’ΔHygR::Intron5'::Read1::TCTCACTTTGACTCC::Read2::Intron3'::3’ΔHygR + 5’ΔHygR::Intron5'::Read1::TGCCTGTATCATAGG::Read2::Intron3'::3’ΔHygR + 5’ΔHygR::Intron5'::Read1::TGGCGCTATAATGTC::Read2::Intron3'::3’ΔHygR + 5’ΔHygR::Intron5'::Read1::TGTCCCTCTTAGAAG::Read2::Intron3'::3’ΔHygR + 5’ΔHygR::Intron5'::Read1::TTCCATTATCAACCC::Read2::Intron3'::3’ΔHygR + 5’ΔHygR::Intron5'::Read1::TTTCTATCTCATCGT::Read2::Intron3'::3’ΔHygR + 5’ΔHygR::Intron5'::Read1::AAGCAGTTTGAAGTC::Read2::Intron3'::3’ΔHygR + 5’ΔHygR::Intron5'::Read1::TCCCCTTCTTACCGT::Read2::Intron3'::3’ΔHygR + 5’ΔHygR::Intron5'::Read1::ACGCAGTGTGACGTA::Read2::Intron3'::3’ΔHygR + 5’ΔHygR::Intron5'::Read1::AGCCCTTCTCAACTA::Read2::Intron3'::3’ΔHygR + 5’ΔHygR::Intron5'::Read1::CCACACTCTCACCTC::Read2::Intron3'::3’ΔHygR + 5’ΔHygR::Intron5'::Read1::CTGCGCTGTCAGCCC::Read2::Intron3'::3’ΔHygR + 5’ΔHygR::Intron5'::Read1::CTTCTCTATCACCAT::Read2::Intron3'::3’ΔHygR + 5’ΔHygR::Intron5'::Read1::GCTCAGTTTTACACG::Read2::Intron3'::3’ΔHygR + 5’ΔHygR::Intron5'::Read1::GTCCAATGTAATGTA::Read2::Intron3'::3’ΔHygR + 5’ΔHygR::Intron5'::Read1::GTCCGATCTAATCCA::Read2::Intron3'::3’ΔHygR + 5’ΔHygR::Intron5'::Read1::TGGCAATATCAACAC::Read2::Intron3'::3’ΔHygR + 5’ΔHygR::Intron5'::Read1::AAGCTGTCTCAGAAT::Read2::Intron3'::3’ΔHygR + 5’ΔHygR::Intron5'::Read1::ATTCTCTTTGAACTT::Read2::Intron3'::3’ΔHygR + 5’ΔHygR::Intron5'::Read1::CGTCGATTTTACAAT::Read2::Intron3'::3’ΔHygR + 5’ΔHygR::Intron5'::Read1::CTTCTCTCTTAGATC::Read2::Intron3'::3’ΔHygR + 5’ΔHygR::Intron5'::Read1::GTTCTATCTTATTCA::Read2::Intron3'::3’ΔHygR + 5’ΔHygR::Intron5'::Read1::TCACGGTGTCATAGT::Read2::Intron3'::3’ΔHygR + 5’ΔHygR::Intron5'::Read1::TCACTCTATCATTCC::Read2::Intron3'::3’ΔHygR + 5’ΔHygR::Intron5'::Read1::TCTCGCTATCATACG::Read2::Intron3'::3’ΔHygR + 5’ΔHygR::Intron5'::Read1::TTGCACTATCATACA::Read2::Intron3'::3’ΔHygR + 5’ΔHygR::Intron5'::Read1::AAGCACTTTTAAGTG::Read2::Intron3'::3’ΔHygR + 5’ΔHygR::Intron5'::Read1::ACCCCATGTCACAAC::Read2::Intron3'::3’ΔHygR + 5’ΔHygR::Intron5'::Read1::ACTCTTTATAATGCC::Read2::Intron3'::3’ΔHygR + 5’ΔHygR::Intron5'::Read1::CAACCGTTTAAACTC::Read2::Intron3'::3’ΔHygR + 5’ΔHygR::Intron5'::Read1::CGCCCTTATCAAAAT::Read2::Intron3'::3’ΔHygR + 5’ΔHygR::Intron5'::Read1::TAGCTTTCTTACCAG::Read2::Intron3'::3’ΔHygR + 5’ΔHygR::Intron5'::Read1::AACCACTATCACCGT::Read2::Intron3'::3’ΔHygR + 5’ΔHygR::Intron5'::Read1::AATCATTTTTACACG::Read2::Intron3'::3’ΔHygR + 5’ΔHygR::Intron5'::Read1::AATCGGTCTGATGAA::Read2::Intron3'::3’ΔHygR + 5’ΔHygR::Intron5'::Read1::ACGCCGTTTCACATG::Read2::Intron3'::3’ΔHygR + 5’ΔHygR::Intron5'::Read1::CCGCCCTATTACAAT::Read2::Intron3'::3’ΔHygR + 5’ΔHygR::Intron5'::Read1::CGACAATCTTATTCA::Read2::Intron3'::3’ΔHygR + 5’ΔHygR::Intron5'::Read1::CTTCAATATCACTCA::Read2::Intron3'::3’ΔHygR + 5’ΔHygR::Intron5'::Read1::TAGCCGTATGAGGAA::Read2::Intron3'::3’ΔHygR + 5’ΔHygR::Intron5'::Read1::TATCTCTCTCAATTA::Read2::Intron3'::3’ΔHygR + 5’ΔHygR::Intron5'::Read1::TCACCCTCTTAACAA::Read2::Intron3'::3’ΔHygR + 5’ΔHygR::Intron5'::Read1::TTCCATTTTGATTTC::Read2::Intron3'::3’ΔHygR + 5’ΔHygR::Intron5'::Read1::CCACCATATTACGTT::Read2::Intron3'::3’ΔHygR +

5’ΔHygR::Intron5'::Read1::GAGCAGTCTAAGTCC::Read2::Intron3'::3’ΔHygR + 5’ΔHygR::Intron5'::Read1::GCGCACTTTCAACAT::Read2::Intron3'::3’ΔHygR + 5’ΔHygR::Intron5'::Read1::GCGCATTATCACAAG::Read2::Intron3'::3’ΔHygR + 5’ΔHygR::Intron5'::Read1::TGCCAATATAAATCA::Read2::Intron3'::3’ΔHygR + 5’ΔHygR::Intron5'::Read1::TTTCTGTCTTAAGCC::Read2::Intron3'::3’ΔHygR + 5’ΔHygR::Intron5'::Read1::AAGCCTTTTAACTCT::Read2::Intron3'::3’ΔHygR + 5’ΔHygR::Intron5'::Read1::AGCCATTTTAACAAA::Read2::Intron3'::3’ΔHygR + 5’ΔHygR::Intron5'::Read1::CTCCTTTCTCACAGA::Read2::Intron3'::3’ΔHygR + 5’ΔHygR::Intron5'::Read1::TAACACTTTAATTAC::Read2::Intron3'::3’ΔHygR + 5’ΔHygR::Intron5'::Read1::TCACTATTTAAGCTA::Read2::Intron3'::3’ΔHygR + 5’ΔHygR::Intron5'::Read1::ACTCCATATCAACAG::Read2::Intron3'::3’ΔHygR + 5’ΔHygR::Intron5'::Read1::ACTCGCTATCATGAC::Read2::Intron3'::3’ΔHygR + 5’ΔHygR::Intron5'::Read1::ATACCCTCTGAAATA::Read2::Intron3'::3’ΔHygR + 5’ΔHygR::Intron5'::Read1::GTACTATATTAGGTT::Read2::Intron3'::3’ΔHygR + 5’ΔHygR::Intron5'::Read1::TAACTATATCAGGAT::Read2::Intron3'::3’ΔHygR + 5’ΔHygR::Intron5'::Read1::TATCCCTCTGACAAG::Read2::Intron3'::3’ΔHygR + 5’ΔHygR::Intron5'::Read1::TTCCAGTGTTAAATC::Read2::Intron3'::3’ΔHygR + 5’ΔHygR::Intron5'::Read1::ACCCCCTGTTAAATT::Read2::Intron3'::3’ΔHygR + 5’ΔHygR::Intron5'::Read1::ACCCTGTGTCATCAC::Read2::Intron3'::3’ΔHygR + 5’ΔHygR::Intron5'::Read1::CATCTGTCTCACATT::Read2::Intron3'::3’ΔHygR + 5’ΔHygR::Intron5'::Read1::CCACCCTCTTATTAA::Read2::Intron3'::3’ΔHygR + 5’ΔHygR::Intron5'::Read1::CCTCAATTTAAAGCT::Read2::Intron3'::3’ΔHygR + 5’ΔHygR::Intron5'::Read1::CCTCCATCTCATGCT::Read2::Intron3'::3’ΔHygR + 5’ΔHygR::Intron5'::Read1::CCTCTTTCTAATCTT::Read2::Intron3'::3’ΔHygR + 5’ΔHygR::Intron5'::Read1::CTCCAGTATAAACCG::Read2::Intron3'::3’ΔHygR + 5’ΔHygR::Intron5'::Read1::GAGCAGTATTATCCA::Read2::Intron3'::3’ΔHygR + 5’ΔHygR::Intron5'::Read1::TCACCCTGTCATCAA::Read2::Intron3'::3’ΔHygR + 5’ΔHygR::Intron5'::Read1::TGGCGCTGTCAATCG::Read2::Intron3'::3’ΔHygR + 5’ΔHygR::Intron5'::Read1::TTCCAGTCTAAATAT::Read2::Intron3'::3’ΔHygR + 5’ΔHygR::Intron5'::Read1::AAGCGATTTCATCAT::Read2::Intron3'::3’ΔHygR + 5’ΔHygR::Intron5'::Read1::AATCTCTTTAAATCA::Read2::Intron3'::3’ΔHygR + 5’ΔHygR::Intron5'::Read1::ATTCCCTTTCACACG::Read2::Intron3'::3’ΔHygR + 5’ΔHygR::Intron5'::Read1::ATTCTTTGTCAACAA::Read2::Intron3'::3’ΔHygR + 5’ΔHygR::Intron5'::Read1::CCGCATTTTCATGCA::Read2::Intron3'::3’ΔHygR + 5’ΔHygR::Intron5'::Read1::CCTCACTCTTATTTC::Read2::Intron3'::3’ΔHygR + 5’ΔHygR::Intron5'::Read1::CTACGTTTTTACCAA::Read2::Intron3'::3’ΔHygR + 5’ΔHygR::Intron5'::Read1::CTCCAATATAACTCT::Read2::Intron3'::3’ΔHygR + 5’ΔHygR::Intron5'::Read1::CTTCCTTATAAATGG::Read2::Intron3'::3’ΔHygR + 5’ΔHygR::Intron5'::Read1::GGTCGCTCTCAGTAT::Read2::Intron3'::3’ΔHygR + 5’ΔHygR::Intron5'::Read1::TTTCAATATCATAGC::Read2::Intron3'::3’ΔHygR + 5’ΔHygR::Intron5'::Read1::AACCTCTGTTAATTG::Read2::Intron3'::3’ΔHygR + 5’ΔHygR::Intron5'::Read1::ATCCGATCTAAGTGC::Read2::Intron3'::3’ΔHygR + 5’ΔHygR::Intron5'::Read1::CAACCGTGTTACTAA::Read2::Intron3'::3’ΔHygR + 5’ΔHygR::Intron5'::Read1::GGCCTTTATAATCGC::Read2::Intron3'::3’ΔHygR + 5’ΔHygR::Intron5'::Read1::TAGCATTTTAATAAT::Read2::Intron3'::3’ΔHygR + 5’ΔHygR::Intron5'::Read1::TATCACTGTGAGAAA::Read2::Intron3'::3’ΔHygR + 5’ΔHygR::Intron5'::Read1::ACTCACTCTTACTCA::Read2::Intron3'::3’ΔHygR + 5’ΔHygR::Intron5'::Read1::CACCGCTTTGACATA::Read2::Intron3'::3’ΔHygR + 5’ΔHygR::Intron5'::Read1::CCGCAATTTAATCCT::Read2::Intron3'::3’ΔHygR + 5’ΔHygR::Intron5'::Read1::CTTCTCTATAAACCA::Read2::Intron3'::3’ΔHygR + 5’ΔHygR::Intron5'::Read1::GGGCTTTTTGAAGGC::Read2::Intron3'::3’ΔHygR + 5’ΔHygR::Intron5'::Read1::TAACAATTTAAGGGA::Read2::Intron3'::3’ΔHygR + 5’ΔHygR::Intron5'::Read1::TACCTTTCTTACTAG::Read2::Intron3'::3’ΔHygR + 5’ΔHygR::Intron5'::Read1::TGGCTGTCTTAGTCT::Read2::Intron3'::3’ΔHygR + 5’ΔHygR::Intron5'::Read1::TTGCGGTATAACAAA::Read2::Intron3'::3’ΔHygR + 5’ΔHygR::Intron5'::Read1::ACACCTTTTCATCAG::Read2::Intron3'::3’ΔHygR + 5’ΔHygR::Intron5'::Read1::ACACTTTATAAAAAA::Read2::Intron3'::3’ΔHygR + 5’ΔHygR::Intron5'::Read1::ACCCCTTTTTAGCAG::Read2::Intron3'::3’ΔHygR + 5’ΔHygR::Intron5'::Read1::TCACGTTATGAACCA::Read2::Intron3'::3’ΔHygR + 5’ΔHygR::Intron5'::Read1::TCACTATCTCAAACA::Read2::Intron3'::3’ΔHygR + 5’ΔHygR::Intron5'::Read1::ATACCTTATAAGATG::Read2::Intron3'::3’ΔHygR + 5’ΔHygR::Intron5'::Read1::CCCCATTCTCACTTG::Read2::Intron3'::3’ΔHygR + 5’ΔHygR::Intron5'::Read1::CGTCCATCTCAACTT::Read2::Intron3'::3’ΔHygR + 5’ΔHygR::Intron5'::Read1::TACCGATTTCACACC::Read2::Intron3'::3’ΔHygR + 5’ΔHygR::Intron5'::Read1::TCTCGCTATGACACC::Read2::Intron3'::3’ΔHygR + 5’ΔHygR::Intron5'::Read1::TCTCGGTCTAACCCC::Read2::Intron3'::3’ΔHygR + 5’ΔHygR::Intron5'::Read1::TGTCACTTTCATTAT::Read2::Intron3'::3’ΔHygR + 5’ΔHygR::Intron5'::Read1::TGTCATTCTCAAACT::Read2::Intron3'::3’ΔHygR + 5’ΔHygR::Intron5'::Read1::AAACCGTTTAATCAT::Read2::Intron3'::3’ΔHygR + 5’ΔHygR::Intron5'::Read1::AAGCAGTGTGACTTG::Read2::Intron3'::3’ΔHygR +

5’ΔHygR::Intron5'::Read1::GCGCTATATCAATAC::Read2::Intron3'::3’ΔHygR + 5’ΔHygR::Intron5'::Read1::TAACAGTATAACCTG::Read2::Intron3'::3’ΔHygR + 5’ΔHygR::Intron5'::Read1::TCGCCCTATAATGAA::Read2::Intron3'::3’ΔHygR + 5’ΔHygR::Intron5'::Read1::TTACATTGTTAATCC::Read2::Intron3'::3’ΔHygR + 5’ΔHygR::Intron5'::Read1::TTTCAATATCAGCTC::Read2::Intron3'::3’ΔHygR + 5’ΔHygR::Intron5'::Read1::TTTCTTTTTGATACC::Read2::Intron3'::3’ΔHygR + 5’ΔHygR::Intron5'::Read1::AAACCCTTTAATGCC::Read2::Intron3'::3’ΔHygR + 5’ΔHygR::Intron5'::Read1::ACGCCATGTTATCCC::Read2::Intron3'::3’ΔHygR + 5’ΔHygR::Intron5'::Read1::ATTCTGTTTGAACAC::Read2::Intron3'::3’ΔHygR + 5’ΔHygR::Intron5'::Read1::CATCATTATAATTGA::Read2::Intron3'::3’ΔHygR + 5’ΔHygR::Intron5'::Read1::CCCCAATTTGAGACC::Read2::Intron3'::3’ΔHygR + 5’ΔHygR::Intron5'::Read1::CTACGTTGTTAGTCA::Read2::Intron3'::3’ΔHygR + 5’ΔHygR::Intron5'::Read1::GCCCCTTGTTATTCA::Read2::Intron3'::3’ΔHygR + 5’ΔHygR::Intron5'::Read1::GTTCTTTTTTAATTT::Read2::Intron3'::3’ΔHygR + 5’ΔHygR::Intron5'::Read1::AGGCTATCTCACATC::Read2::Intron3'::3’ΔHygR + 5’ΔHygR::Intron5'::Read1::GCCCACTCTGAATCA::Read2::Intron3'::3’ΔHygR + 5’ΔHygR::Intron5'::Read1::GTTCTCTATAACTTA::Read2::Intron3'::3’ΔHygR + 5’ΔHygR::Intron5'::Read1::TCACGGTTTGAAGTT::Read2::Intron3'::3’ΔHygR + 5’ΔHygR::Intron5'::Read1::TCCCCTTTTTAACCG::Read2::Intron3'::3’ΔHygR + 5’ΔHygR::Intron5'::Read1::TCGCTGTTTGATTTA::Read2::Intron3'::3’ΔHygR + 5’ΔHygR::Intron5'::Read1::AAACAATTTCAACTC::Read2::Intron3'::3’ΔHygR + 5’ΔHygR::Intron5'::Read1::ATACCATATGAAAAG::Read2::Intron3'::3’ΔHygR + 5’ΔHygR::Intron5'::Read1::CCTCACTGTTACCTA::Read2::Intron3'::3’ΔHygR + 5’ΔHygR::Intron5'::Read1::CTGCTATGTTACACC::Read2::Intron3'::3’ΔHygR + 5’ΔHygR::Intron5'::Read1::GTTCAGTGTTATCCG::Read2::Intron3'::3’ΔHygR + 5’ΔHygR::Intron5'::Read1::GTTCGGTTTAATAAC::Read2::Intron3'::3’ΔHygR + 5’ΔHygR::Intron5'::Read1::TAACGTTCTCAACTA::Read2::Intron3'::3’ΔHygR + 5’ΔHygR::Intron5'::Read1::TCCCGCTGTAATCAG::Read2::Intron3'::3’ΔHygR + 5’ΔHygR::Intron5'::Read1::TCGCCTTTTTAGCTT::Read2::Intron3'::3’ΔHygR + 5’ΔHygR::Intron5'::Read1::TGGCCCTATTACTTC::Read2::Intron3'::3’ΔHygR + 5’ΔHygR::Intron5'::Read1::TTGCACTCTCAAGTC::Read2::Intron3'::3’ΔHygR + 5’ΔHygR::Intron5'::Read1::TTTCCGTCTTAACCG::Read2::Intron3'::3’ΔHygR + 5’ΔHygR::Intron5'::Read1::AACCACTATTAAGTT::Read2::Intron3'::3’ΔHygR + 5’ΔHygR::Intron5'::Read1::AATCGTTATTATCCA::Read2::Intron3'::3’ΔHygR + 5’ΔHygR::Intron5'::Read1::AGGCTATATTAAATC::Read2::Intron3'::3’ΔHygR + 5’ΔHygR::Intron5'::Read1::AGTCAATCTGAAACT::Read2::Intron3'::3’ΔHygR + 5’ΔHygR::Intron5'::Read1::ATCCCTTCTTATTTC::Read2::Intron3'::3’ΔHygR + 5’ΔHygR::Intron5'::Read1::CCCCCATGTCATCTG::Read2::Intron3'::3’ΔHygR + 5’ΔHygR::Intron5'::Read1::CTCCACTTTAAGCCA::Read2::Intron3'::3’ΔHygR + 5’ΔHygR::Intron5'::Read1::TATCTCTGTAAAGCA::Read2::Intron3'::3’ΔHygR + 5’ΔHygR::Intron5'::Read1::TTTCCTTTTCATCAA::Read2::Intron3'::3’ΔHygR + 5’ΔHygR::Intron5'::Read1::CGACAGTTTGACTGC::Read2::Intron3'::3’ΔHygR + 5’ΔHygR::Intron5'::Read1::GCGCCATCTCATGAT::Read2::Intron3'::3’ΔHygR + 5’ΔHygR::Intron5'::Read1::GCTCGATTTTACACA::Read2::Intron3'::3’ΔHygR + 5’ΔHygR::Intron5'::Read1::GGACGTTGTGACATA::Read2::Intron3'::3’ΔHygR + 5’ΔHygR::Intron5'::Read1::TCACGTTATAAAGAG::Read2::Intron3'::3’ΔHygR + 5’ΔHygR::Intron5'::Read1::TCGCGCTCTTATCTT::Read2::Intron3'::3’ΔHygR + 5’ΔHygR::Intron5'::Read1::AAGCTCTATCAACCA::Read2::Intron3'::3’ΔHygR + 5’ΔHygR::Intron5'::Read1::CATCACTGTAAGGCT::Read2::Intron3'::3’ΔHygR + 5’ΔHygR::Intron5'::Read1::CCTCGCTATCATGTC::Read2::Intron3'::3’ΔHygR + 5’ΔHygR::Intron5'::Read1::CTCCCGTTTTATCTC::Read2::Intron3'::3’ΔHygR + 5’ΔHygR::Intron5'::Read1::TATCTTTCTCAGCCA::Read2::Intron3'::3’ΔHygR + 5’ΔHygR::Intron5'::Read1::TGCCAATGTAAGACC::Read2::Intron3'::3’ΔHygR + 5’ΔHygR::Intron5'::Read1::TTTCATTCTCATACC::Read2::Intron3'::3’ΔHygR + 5’ΔHygR::Intron5'::Read1::AACCATTGTAACAAA::Read2::Intron3'::3’ΔHygR + 5’ΔHygR::Intron5'::Read1::AATCCCTATCATTAT::Read2::Intron3'::3’ΔHygR + 5’ΔHygR::Intron5'::Read1::ATACCCTCTAATTCC::Read2::Intron3'::3’ΔHygR + 5’ΔHygR::Intron5'::Read1::ATGCTATTTAATTTC::Read2::Intron3'::3’ΔHygR + 5’ΔHygR::Intron5'::Read1::AGTCACTTTTACCAG::Read2::Intron3'::3’ΔHygR + 5’ΔHygR::Intron5'::Read1::CCGCAGTATCACCGC::Read2::Intron3'::3’ΔHygR + 5’ΔHygR::Intron5'::Read1::CCGCCGTCTAAAACC::Read2::Intron3'::3’ΔHygR + 5’ΔHygR::Intron5'::Read1::CTCCGATTTTAACGT::Read2::Intron3'::3’ΔHygR + 5’ΔHygR::Intron5'::Read1::GCGCTTTCTGATCTT::Read2::Intron3'::3’ΔHygR + 5’ΔHygR::Intron5'::Read1::TTACCATCTCAATAG::Read2::Intron3'::3’ΔHygR + 5’ΔHygR::Intron5'::Read1::AGGCGGTATCAGAGG::Read2::Intron3'::3’ΔHygR + 5’ΔHygR::Intron5'::Read1::CCACTGTCTTACGTA::Read2::Intron3'::3’ΔHygR + 5’ΔHygR::Intron5'::Read1::CCGCTATCTAACTAT::Read2::Intron3'::3’ΔHygR + 5’ΔHygR::Intron5'::Read1::CCTCCATCTGATAAC::Read2::Intron3'::3’ΔHygR + 5’ΔHygR::Intron5'::Read1::GCACGTTGTCATCTT::Read2::Intron3'::3’ΔHygR + 5’ΔHygR::Intron5'::Read1::GTGCTATCTTATTCC::Read2::Intron3'::3’ΔHygR +

5’ΔHygR::Intron5'::Read1::TAACAATTTTACCTT::Read2::Intron3'::3’ΔHygR + 5’ΔHygR::Intron5'::Read1::AGACAGTCTAACATC::Read2::Intron3'::3’ΔHygR + 5’ΔHygR::Intron5'::Read1::GCACCGTCTCACTCA::Read2::Intron3'::3’ΔHygR + 5’ΔHygR::Intron5'::Read1::GCCCCATATTAGAAG::Read2::Intron3'::3’ΔHygR + 5’ΔHygR::Intron5'::Read1::TAACCCTCTGATCCA::Read2::Intron3'::3’ΔHygR + 5’ΔHygR::Intron5'::Read1::TGACCCTCTTACGGC::Read2::Intron3'::3’ΔHygR + 5’ΔHygR::Intron5'::Read1::TGGCAATCTTATACC::Read2::Intron3'::3’ΔHygR + 5’ΔHygR::Intron5'::Read1::AGTCTGTTTAAGCAA::Read2::Intron3'::3’ΔHygR + 5’ΔHygR::Intron5'::Read1::ATCCACTATGAGACC::Read2::Intron3'::3’ΔHygR + 5’ΔHygR::Intron5'::Read1::CCCCAATCTCATTGT::Read2::Intron3'::3’ΔHygR + 5’ΔHygR::Intron5'::Read1::TTCCCTTATTACTAC::Read2::Intron3'::3’ΔHygR + 5’ΔHygR::Intron5'::Read1::ACACACTATCAGAAT::Read2::Intron3'::3’ΔHygR + 5’ΔHygR::Intron5'::Read1::CAACATTATCAGCAT::Read2::Intron3'::3’ΔHygR + 5’ΔHygR::Intron5'::Read1::TCCCCCTATTAGCCT::Read2::Intron3'::3’ΔHygR + 5’ΔHygR::Intron5'::Read1::TCTCACTATTATAAG::Read2::Intron3'::3’ΔHygR + 5’ΔHygR::Intron5'::Read1::TTACCCTTTGATCTC::Read2::Intron3'::3’ΔHygR + 5’ΔHygR::Intron5'::Read1::ATGCAATGTGATACG::Read2::Intron3'::3’ΔHygR + 5’ΔHygR::Intron5'::Read1::CGACGGTCTAAGGAG::Read2::Intron3'::3’ΔHygR + 5’ΔHygR::Intron5'::Read1::CGTCAGTGTCAATTC::Read2::Intron3'::3’ΔHygR + 5’ΔHygR::Intron5'::Read1::GCCCAATTTAATTTA::Read2::Intron3'::3’ΔHygR + 5’ΔHygR::Intron5'::Read1::GCGCCATATCACACA::Read2::Intron3'::3’ΔHygR + 5’ΔHygR::Intron5'::Read1::TAACGTTATAAAGCA::Read2::Intron3'::3’ΔHygR + 5’ΔHygR::Intron5'::Read1::TCACAGTCTAAGCCC::Read2::Intron3'::3’ΔHygR + 5’ΔHygR::Intron5'::Read1::TGTCTCTCTAAGATC::Read2::Intron3'::3’ΔHygR + 5’ΔHygR::Intron5'::Read1::AGACGCTCTCACAGC::Read2::Intron3'::3’ΔHygR + 5’ΔHygR::Intron5'::Read1::GTACATTTTTAAGTA::Read2::Intron3'::3’ΔHygR + 5’ΔHygR::Intron5'::Read1::GTTCCTTGTCACTAC::Read2::Intron3'::3’ΔHygR + 5’ΔHygR::Intron5'::Read1::GTTCGGTATTACCCA::Read2::Intron3'::3’ΔHygR + 5’ΔHygR::Intron5'::Read1::TGACCATTTGAAGTA::Read2::Intron3'::3’ΔHygR + 5’ΔHygR::Intron5'::Read1::CACCTTTATAATGTC::Read2::Intron3'::3’ΔHygR + 5’ΔHygR::Intron5'::Read1::GACCTTTTTGAACTT::Read2::Intron3'::3’ΔHygR + 5’ΔHygR::Intron5'::Read1::TACCATTGTAACGAC::Read2::Intron3'::3’ΔHygR + 5’ΔHygR::Intron5'::Read1::TTTCATTATTACTCT::Read2::Intron3'::3’ΔHygR + 5’ΔHygR::Intron5'::Read1::TTTCCATATAAGGTA::Read2::Intron3'::3’ΔHygR + 5’ΔHygR::Intron5'::Read1::ACACCATTTAACCCA::Read2::Intron3'::3’ΔHygR + 5’ΔHygR::Intron5'::Read1::ACACTTTGTTAAGGC::Read2::Intron3'::3’ΔHygR + 5’ΔHygR::Intron5'::Read1::ACGCGATCTCATTTA::Read2::Intron3'::3’ΔHygR + 5’ΔHygR::Intron5'::Read1::ATACTGTTTCACCAT::Read2::Intron3'::3’ΔHygR + 5’ΔHygR::Intron5'::Read1::CCTCGTTGTCAATAC::Read2::Intron3'::3’ΔHygR + 5’ΔHygR::Intron5'::Read1::CGACTTTCTCAGGTA::Read2::Intron3'::3’ΔHygR + 5’ΔHygR::Intron5'::Read1::CTACACTATGATATA::Read2::Intron3'::3’ΔHygR + 5’ΔHygR::Intron5'::Read1::CTGCCTTTTTAATCC::Read2::Intron3'::3’ΔHygR + 5’ΔHygR::Intron5'::Read1::GCCCAATCTAAGGTA::Read2::Intron3'::3’ΔHygR + 5’ΔHygR::Intron5'::Read1::GGTCGCTATTAAAAA::Read2::Intron3'::3’ΔHygR + 5’ΔHygR::Intron5'::Read1::TCACGATTTGAGGCC::Read2::Intron3'::3’ΔHygR + 5’ΔHygR::Intron5'::Read1::TCCCTATTTAAAACC::Read2::Intron3'::3’ΔHygR + 5’ΔHygR::Intron5'::Read1::TTGCAATGTGAATTT::Read2::Intron3'::3’ΔHygR + 5’ΔHygR::Intron5'::Read1::CCGCCTTATTAAACA::Read2::Intron3'::3’ΔHygR + 5’ΔHygR::Intron5'::Read1::CTACGCTGTAATCGT::Read2::Intron3'::3’ΔHygR + 5’ΔHygR::Intron5'::Read1::CTCCTTTCTCACCTG::Read2::Intron3'::3’ΔHygR + 5’ΔHygR::Intron5'::Read1::TAGCTCTGTCAAGTC::Read2::Intron3'::3’ΔHygR + 5’ΔHygR::Intron5'::Read1::TCACCTTCTTAAGCA::Read2::Intron3'::3’ΔHygR + 5’ΔHygR::Intron5'::Read1::TGGCGCTCTAATCGT::Read2::Intron3'::3’ΔHygR + 5’ΔHygR::Intron5'::Read1::ACCCAATTTTACTTC::Read2::Intron3'::3’ΔHygR + 5’ΔHygR::Intron5'::Read1::AGACCGTATAATCCC::Read2::Intron3'::3’ΔHygR + 5’ΔHygR::Intron5'::Read1::GTTCCTTGTTAGCCA::Read2::Intron3'::3’ΔHygR + 5’ΔHygR::Intron5'::Read1::TACCTGTTTCATTTT::Read2::Intron3'::3’ΔHygR + 5’ΔHygR::Intron5'::Read1::TATCCCTGTGACAGC::Read2::Intron3'::3’ΔHygR + 5’ΔHygR::Intron5'::Read1::TGACATTTTCAGTTA::Read2::Intron3'::3’ΔHygR + 5’ΔHygR::Intron5'::Read1::TTTCTATTTAAGAAT::Read2::Intron3'::3’ΔHygR + 5’ΔHygR::Intron5'::Read1::ACGCGGTCTCAGCTC::Read2::Intron3'::3’ΔHygR + 5’ΔHygR::Intron5'::Read1::CAACGGTCTTAACCT::Read2::Intron3'::3’ΔHygR + 5’ΔHygR::Intron5'::Read1::CAGCGATCTGACATA::Read2::Intron3'::3’ΔHygR + 5’ΔHygR::Intron5'::Read1::CTTCAGTTTGATTGC::Read2::Intron3'::3’ΔHygR + 5’ΔHygR::Intron5'::Read1::TACCTGTTTTACCAA::Read2::Intron3'::3’ΔHygR + 5’ΔHygR::Intron5'::Read1::TATCTGTCTAATAAA::Read2::Intron3'::3’ΔHygR + 5’ΔHygR::Intron5'::Read1::TGACCTTGTCAGCCG::Read2::Intron3'::3’ΔHygR + 5’ΔHygR::Intron5'::Read1::TGCCGATTTAATTTA::Read2::Intron3'::3’ΔHygR + 5’ΔHygR::Intron5'::Read1::TTACTATTTTAGTTC::Read2::Intron3'::3’ΔHygR + 5’ΔHygR::Intron5'::Read1::AATCTATATCACTTG::Read2::Intron3'::3’ΔHygR +

5’ΔHygR::Intron5'::Read1::ACTCTGTTTTAACAT::Read2::Intron3'::3’ΔHygR + 5’ΔHygR::Intron5'::Read1::AGTCCGTATTACATG::Read2::Intron3'::3’ΔHygR + 5’ΔHygR::Intron5'::Read1::ATACTATATAATTTA::Read2::Intron3'::3’ΔHygR + 5’ΔHygR::Intron5'::Read1::CGTCCCTTTGAGATA::Read2::Intron3'::3’ΔHygR + 5’ΔHygR::Intron5'::Read1::GCCCCTTTTCACAAA::Read2::Intron3'::3’ΔHygR + 5’ΔHygR::Intron5'::Read1::GCGCAATTTCACTTC::Read2::Intron3'::3’ΔHygR + 5’ΔHygR::Intron5'::Read1::TGACGATCTCAGCCA::Read2::Intron3'::3’ΔHygR + 5’ΔHygR::Intron5'::Read1::TTCCCGTATTACGTC::Read2::Intron3'::3’ΔHygR + 5’ΔHygR::Intron5'::Read1::AGACTATATTACAAC::Read2::Intron3'::3’ΔHygR + 5’ΔHygR::Intron5'::Read1::AGTCCGTATCAGGAA::Read2::Intron3'::3’ΔHygR + 5’ΔHygR::Intron5'::Read1::CATCTTTGTAACGTT::Read2::Intron3'::3’ΔHygR + 5’ΔHygR::Intron5'::Read1::CCTCTATATCATCTA::Read2::Intron3'::3’ΔHygR + 5’ΔHygR::Intron5'::Read1::CTGCGGTATTATTTC::Read2::Intron3'::3’ΔHygR + 5’ΔHygR::Intron5'::Read1::GCGCTCTGTTACCTG::Read2::Intron3'::3’ΔHygR + 5’ΔHygR::Intron5'::Read1::GTACTCTCTCAACAA::Read2::Intron3'::3’ΔHygR + 5’ΔHygR::Intron5'::Read1::TAACCATTTCACATA::Read2::Intron3'::3’ΔHygR + 5’ΔHygR::Intron5'::Read1::TAGCCCTGTCACTAC::Read2::Intron3'::3’ΔHygR + 5’ΔHygR::Intron5'::Read1::TCACCCTTTAAATAT::Read2::Intron3'::3’ΔHygR + 5’ΔHygR::Intron5'::Read1::AAGCATTGTTACCTT::Read2::Intron3'::3’ΔHygR + 5’ΔHygR::Intron5'::Read1::CGACTTTGTCAAACC::Read2::Intron3'::3’ΔHygR + 5’ΔHygR::Intron5'::Read1::CTCCGTTCTCATTCT::Read2::Intron3'::3’ΔHygR + 5’ΔHygR::Intron5'::Read1::GTTCTTTTTTAAATG::Read2::Intron3'::3’ΔHygR + 5’ΔHygR::Intron5'::Read1::TAACGCTTTTAATCC::Read2::Intron3'::3’ΔHygR + 5’ΔHygR::Intron5'::Read1::TACCAATCTTATCCT::Read2::Intron3'::3’ΔHygR + 5’ΔHygR::Intron5'::Read1::TCACAGTATAAAACC::Read2::Intron3'::3’ΔHygR + 5’ΔHygR::Intron5'::Read1::TCGCATTATTATTCG::Read2::Intron3'::3’ΔHygR + 5’ΔHygR::Intron5'::Read1::TTGCCGTTTCACTAC::Read2::Intron3'::3’ΔHygR + 5’ΔHygR::Intron5'::Read1::ACCCTATGTAAAAAC::Read2::Intron3'::3’ΔHygR + 5’ΔHygR::Intron5'::Read1::ACGCCATATGAATAG::Read2::Intron3'::3’ΔHygR + 5’ΔHygR::Intron5'::Read1::ACTCCTTTTCACTAA::Read2::Intron3'::3’ΔHygR + 5’ΔHygR::Intron5'::Read1::CGTCACTTTAATCTC::Read2::Intron3'::3’ΔHygR + 5’ΔHygR::Intron5'::Read1::TACCATTGTAATGTC::Read2::Intron3'::3’ΔHygR + 5’ΔHygR::Intron5'::Read1::TACCGTTCTAACCTA::Read2::Intron3'::3’ΔHygR + 5’ΔHygR::Intron5'::Read1::TAGCTATATCAAACC::Read2::Intron3'::3’ΔHygR + 5’ΔHygR::Intron5'::Read1::TCACCTTATTACACA::Read2::Intron3'::3’ΔHygR + 5’ΔHygR::Intron5'::Read1::TGACTTTGTCACTAT::Read2::Intron3'::3’ΔHygR + 5’ΔHygR::Intron5'::Read1::CCCCGCTCTAAGTAA::Read2::Intron3'::3’ΔHygR + 5’ΔHygR::Intron5'::Read1::CGCCTTTATCAACCA::Read2::Intron3'::3’ΔHygR + 5’ΔHygR::Intron5'::Read1::TACCCATGTAACCTA::Read2::Intron3'::3’ΔHygR + 5’ΔHygR::Intron5'::Read1::TCTCCTTTTAACAAC::Read2::Intron3'::3’ΔHygR + 5’ΔHygR::Intron5'::Read1::TTACGTTCTAACCAT::Read2::Intron3'::3’ΔHygR + 5’ΔHygR::Intron5'::Read1::TTTCTTTCTGACTTG::Read2::Intron3'::3’ΔHygR + 5’ΔHygR::Intron5'::Read1::AAACCGTTTCATGAT::Read2::Intron3'::3’ΔHygR + 5’ΔHygR::Intron5'::Read1::ATACCCTGTCATCCC::Read2::Intron3'::3’ΔHygR + 5’ΔHygR::Intron5'::Read1::ATCCCTTTTGACCTA::Read2::Intron3'::3’ΔHygR + 5’ΔHygR::Intron5'::Read1::ATGCCTTCTAACTAA::Read2::Intron3'::3’ΔHygR + 5’ΔHygR::Intron5'::Read1::CATCCGTCTCATCAA::Read2::Intron3'::3’ΔHygR + 5’ΔHygR::Intron5'::Read1::CGACAGTTTAATATC::Read2::Intron3'::3’ΔHygR + 5’ΔHygR::Intron5'::Read1::CTTCTATATCACCAT::Read2::Intron3'::3’ΔHygR + 5’ΔHygR::Intron5'::Read1::GTCCAGTGTAAGCCA::Read2::Intron3'::3’ΔHygR + 5’ΔHygR::Intron5'::Read1::GTGCTCTTTGATCCT::Read2::Intron3'::3’ΔHygR + 5’ΔHygR::Intron5'::Read1::TTACCTTATAAGCAT::Read2::Intron3'::3’ΔHygR + 5’ΔHygR::Intron5'::Read1::TTACGCTCTAACTCA::Read2::Intron3'::3’ΔHygR + 5’ΔHygR::Intron5'::Read1::TTTCTGTTTCACGTC::Read2::Intron3'::3’ΔHygR + 5’ΔHygR::Intron5'::Read1::AGTCGCTTTTAATGT::Read2::Intron3'::3’ΔHygR + 5’ΔHygR::Intron5'::Read1::ATCCGATCTAATTTA::Read2::Intron3'::3’ΔHygR + 5’ΔHygR::Intron5'::Read1::ATCCTGTTTTACTAA::Read2::Intron3'::3’ΔHygR + 5’ΔHygR::Intron5'::Read1::CGCCTCTGTGAAGTA::Read2::Intron3'::3’ΔHygR + 5’ΔHygR::Intron5'::Read1::GATCTGTCTTATTTG::Read2::Intron3'::3’ΔHygR + 5’ΔHygR::Intron5'::Read1::GGGCCATCTGAAGGT::Read2::Intron3'::3’ΔHygR + 5’ΔHygR::Intron5'::Read1::TGCCAATCTAAAAGC::Read2::Intron3'::3’ΔHygR + 5’ΔHygR::Intron5'::Read1::TGCCCCTTTTACGCT::Read2::Intron3'::3’ΔHygR + 5’ΔHygR::Intron5'::Read1::TTCCCATATGACAGT::Read2::Intron3'::3’ΔHygR + 5’ΔHygR::Intron5'::Read1::TTCCCTTCTTATATC::Read2::Intron3'::3’ΔHygR + 5’ΔHygR::Intron5'::Read1::ACTCACTGTAAGATC::Read2::Intron3'::3’ΔHygR + 5’ΔHygR::Intron5'::Read1::CAACACTCTGACACC::Read2::Intron3'::3’ΔHygR + 5’ΔHygR::Intron5'::Read1::CGACTAAAACAGATA::Read2::Intron3'::3’ΔHygR + 5’ΔHygR::Intron5'::Read1::CTTCGATTTCACGCG::Read2::Intron3'::3’ΔHygR + 5’ΔHygR::Intron5'::Read1::GGACAATTTAAAACA::Read2::Intron3'::3’ΔHygR + 5’ΔHygR::Intron5'::Read1::GGTCTTTATTACCTC::Read2::Intron3'::3’ΔHygR +

5’ΔHygR::Intron5'::Read1::GTTCAATGTCACCCA::Read2::Intron3'::3’ΔHygR + 5’ΔHygR::Intron5'::Read1::AAGCCTTATTAATCA::Read2::Intron3'::3’ΔHygR + 5’ΔHygR::Intron5'::Read1::ATACAGTATCATGCA::Read2::Intron3'::3’ΔHygR + 5’ΔHygR::Intron5'::Read1::ATACGCTCTAAACAA::Read2::Intron3'::3’ΔHygR + 5’ΔHygR::Intron5'::Read1::ATTCAATCTCAAGAC::Read2::Intron3'::3’ΔHygR + 5’ΔHygR::Intron5'::Read1::ATTCTATTTAATCCC::Read2::Intron3'::3’ΔHygR + 5’ΔHygR::Intron5'::Read1::CACCCTTCTCACGTC::Read2::Intron3'::3’ΔHygR + 5’ΔHygR::Intron5'::Read1::CATCGTTTTCAGTTG::Read2::Intron3'::3’ΔHygR + 5’ΔHygR::Intron5'::Read1::CTACGTTATCATTAT::Read2::Intron3'::3’ΔHygR + 5’ΔHygR::Intron5'::Read1::GTCCAGTTTTATACT::Read2::Intron3'::3’ΔHygR + 5’ΔHygR::Intron5'::Read1::TATCGATCTAACCTC::Read2::Intron3'::3’ΔHygR + 5’ΔHygR::Intron5'::Read1::TCACTTTTTTAATTG::Read2::Intron3'::3’ΔHygR + 5’ΔHygR::Intron5'::Read1::TCGCCTTATAACCTT::Read2::Intron3'::3’ΔHygR + 5’ΔHygR::Intron5'::Read1::TTTCTATGTAATGAC::Read2::Intron3'::3’ΔHygR + 5’ΔHygR::Intron5'::Read1::ACGCATTATAAGAGG::Read2::Intron3'::3’ΔHygR + 5’ΔHygR::Intron5'::Read1::CCTCACTTTCACAAA::Read2::Intron3'::3’ΔHygR + 5’ΔHygR::Intron5'::Read1::CTCCTCTTTTACACG::Read2::Intron3'::3’ΔHygR + 5’ΔHygR::Intron5'::Read1::GAACGGTTTTACACT::Read2::Intron3'::3’ΔHygR + 5’ΔHygR::Intron5'::Read1::GTACTTTTTTAGGGA::Read2::Intron3'::3’ΔHygR + 5’ΔHygR::Intron5'::Read1::TCGCGCTGTGATATT::Read2::Intron3'::3’ΔHygR + 5’ΔHygR::Intron5'::Read1::TCGCTCTGTCACCCG::Read2::Intron3'::3’ΔHygR + 5’ΔHygR::Intron5'::Read1::TGACTTTTTCACTCA::Read2::Intron3'::3’ΔHygR + 5’ΔHygR::Intron5'::Read1::TTACCATCTTAGCAT::Read2::Intron3'::3’ΔHygR + 5’ΔHygR::Intron5'::Read1::TTTCGCTATAACCTC::Read2::Intron3'::3’ΔHygR + 5’ΔHygR::Intron5'::Read1::ACCCCTTGTGATCCT::Read2::Intron3'::3’ΔHygR + 5’ΔHygR::Intron5'::Read1::CAACTCTCTCATTCC::Read2::Intron3'::3’ΔHygR + 5’ΔHygR::Intron5'::Read1::CCCCGGTCTAACGCC::Read2::Intron3'::3’ΔHygR + 5’ΔHygR::Intron5'::Read1::GACCGGTCTGATGGC::Read2::Intron3'::3’ΔHygR + 5’ΔHygR::Intron5'::Read1::TCACGATGTAAGCAC::Read2::Intron3'::3’ΔHygR + 5’ΔHygR::Intron5'::Read1::TTCCTTTATAACACA::Read2::Intron3'::3’ΔHygR + 5’ΔHygR::Intron5'::Read1::ACACCTTCTCACAAA::Read2::Intron3'::3’ΔHygR + 5’ΔHygR::Intron5'::Read1::AGGCTCTGTCATTAA::Read2::Intron3'::3’ΔHygR + 5’ΔHygR::Intron5'::Read1::CACCGCTCTTAAGAT::Read2::Intron3'::3’ΔHygR + 5’ΔHygR::Intron5'::Read1::CGTCATTTTTATTCC::Read2::Intron3'::3’ΔHygR + 5’ΔHygR::Intron5'::Read1::CTCCCCTCTCAGTAA::Read2::Intron3'::3’ΔHygR + 5’ΔHygR::Intron5'::Read1::GACCTTTTTCATTAC::Read2::Intron3'::3’ΔHygR + 5’ΔHygR::Intron5'::Read1::GCACAGTATGATGCC::Read2::Intron3'::3’ΔHygR + 5’ΔHygR::Intron5'::Read1::GCACGCTCTGAAACG::Read2::Intron3'::3’ΔHygR + 5’ΔHygR::Intron5'::Read1::GTGCGCTCTTATAAC::Read2::Intron3'::3’ΔHygR + 5’ΔHygR::Intron5'::Read1::TTACTTTGTCAGTTC::Read2::Intron3'::3’ΔHygR + 5’ΔHygR::Intron5'::Read1::ACACTTTTTCAAATC::Read2::Intron3'::3’ΔHygR + 5’ΔHygR::Intron5'::Read1::AGACGTTTTAATGTC::Read2::Intron3'::3’ΔHygR + 5’ΔHygR::Intron5'::Read1::CAGCGGTATTACTCG::Read2::Intron3'::3’ΔHygR + 5’ΔHygR::Intron5'::Read1::CATCGGTGTGACCGC::Read2::Intron3'::3’ΔHygR + 5’ΔHygR::Intron5'::Read1::CGACTGTCTGACCAC::Read2::Intron3'::3’ΔHygR + 5’ΔHygR::Intron5'::Read1::CTACGGTCTCATCAC::Read2::Intron3'::3’ΔHygR + 5’ΔHygR::Intron5'::Read1::CTCCCTTTTAATGGA::Read2::Intron3'::3’ΔHygR + 5’ΔHygR::Intron5'::Read1::CTGCAATCTAACAAA::Read2::Intron3'::3’ΔHygR + 5’ΔHygR::Intron5'::Read1::CTTCATTTTAAGCCA::Read2::Intron3'::3’ΔHygR + 5’ΔHygR::Intron5'::Read1::GTGCGTTTTAAGCAA::Read2::Intron3'::3’ΔHygR + 5’ΔHygR::Intron5'::Read1::GTTCTTTATCAGCCT::Read2::Intron3'::3’ΔHygR + 5’ΔHygR::Intron5'::Read1::TTCCATTGTCACGCC::Read2::Intron3'::3’ΔHygR + 5’ΔHygR::Intron5'::Read1::AAGCAATGTCAATGG::Read2::Intron3'::3’ΔHygR + 5’ΔHygR::Intron5'::Read1::ACACGGTGTAAGCTA::Read2::Intron3'::3’ΔHygR + 5’ΔHygR::Intron5'::Read1::ACACTGTTTCATTAT::Read2::Intron3'::3’ΔHygR + 5’ΔHygR::Intron5'::Read1::CGGCTTTGTTAATTG::Read2::Intron3'::3’ΔHygR + 5’ΔHygR::Intron5'::Read1::GAACTTTGTCAAACA::Read2::Intron3'::3’ΔHygR + 5’ΔHygR::Intron5'::Read1::GCACTCTCTTAGACA::Read2::Intron3'::3’ΔHygR + 5’ΔHygR::Intron5'::Read1::GCGCTCTTTAAAAAC::Read2::Intron3'::3’ΔHygR + 5’ΔHygR::Intron5'::Read1::GGTCACTGTAAACAC::Read2::Intron3'::3’ΔHygR + 5’ΔHygR::Intron5'::Read1::GGTCTGTGTTACTAG::Read2::Intron3'::3’ΔHygR + 5’ΔHygR::Intron5'::Read1::GTACTTTTTTACTCT::Read2::Intron3'::3’ΔHygR + 5’ΔHygR::Intron5'::Read1::GTCCATTGTCAGGTC::Read2::Intron3'::3’ΔHygR + 5’ΔHygR::Intron5'::Read1::GTGCCCTGTCAATAT::Read2::Intron3'::3’ΔHygR + 5’ΔHygR::Intron5'::Read1::TCCCCATTTAAAGCC::Read2::Intron3'::3’ΔHygR + 5’ΔHygR::Intron5'::Read1::TTCCATTTTTATCTC::Read2::Intron3'::3’ΔHygR + 5’ΔHygR::Intron5'::Read1::AGGCCATATGAGAGT::Read2::Intron3'::3’ΔHygR + 5’ΔHygR::Intron5'::Read1::ATGCGCTTTCAGACA::Read2::Intron3'::3’ΔHygR + 5’ΔHygR::Intron5'::Read1::CACCCGTTTCAAATT::Read2::Intron3'::3’ΔHygR + 5’ΔHygR::Intron5'::Read1::CATCGTTCTAATATC::Read2::Intron3'::3’ΔHygR +

5’ΔHygR::Intron5'::Read1::CGCCATTATAAGCAA::Read2::Intron3'::3’ΔHygR + 5’ΔHygR::Intron5'::Read1::CGTCTATTTCAATAG::Read2::Intron3'::3’ΔHygR + 5’ΔHygR::Intron5'::Read1::CTACCATCTCAATCC::Read2::Intron3'::3’ΔHygR + 5’ΔHygR::Intron5'::Read1::CTACCTTATAAGTAC::Read2::Intron3'::3’ΔHygR + 5’ΔHygR::Intron5'::Read1::CTACGCTCTTACCGC::Read2::Intron3'::3’ΔHygR + 5’ΔHygR::Intron5'::Read1::GAACCTTCTCAGTGC::Read2::Intron3'::3’ΔHygR + 5’ΔHygR::Intron5'::Read1::GCCCATTCTCAAACG::Read2::Intron3'::3’ΔHygR + 5’ΔHygR::Intron5'::Read1::TACCGATTTCATGCT::Read2::Intron3'::3’ΔHygR + 5’ΔHygR::Intron5'::Read1::TGCCTTTCTCAATGG::Read2::Intron3'::3’ΔHygR + 5’ΔHygR::Intron5'::Read1::TTTCTATCTAACGCC::Read2::Intron3'::3’ΔHygR + 5’ΔHygR::Intron5'::Read1::AACCGCTATAACCTC::Read2::Intron3'::3’ΔHygR + 5’ΔHygR::Intron5'::Read1::ACCCATTTTTAATCC::Read2::Intron3'::3’ΔHygR + 5’ΔHygR::Intron5'::Read1::GCCCAATGTCAATGA::Read2::Intron3'::3’ΔHygR + 5’ΔHygR::Intron5'::Read1::GCGCGCTTTTATTTG::Read2::Intron3'::3’ΔHygR + 5’ΔHygR::Intron5'::Read1::TGCCACTCTCAACCA::Read2::Intron3'::3’ΔHygR + 5’ΔHygR::Intron5'::Read1::TGGCCTTTTCAACCC::Read2::Intron3'::3’ΔHygR + 5’ΔHygR::Intron5'::Read1::TTGCATTTTAATTTA::Read2::Intron3'::3’ΔHygR + 5’ΔHygR::Intron5'::Read1::TTGCTCTCTAACCTT::Read2::Intron3'::3’ΔHygR + 5’ΔHygR::Intron5'::Read1::AACCTATTTGAGTCG::Read2::Intron3'::3’ΔHygR + 5’ΔHygR::Intron5'::Read1::CCACCGTATCAAATA::Read2::Intron3'::3’ΔHygR + 5’ΔHygR::Intron5'::Read1::CGGCTCTCTAAAGGA::Read2::Intron3'::3’ΔHygR + 5’ΔHygR::Intron5'::Read1::GACCGCTGTCATGTA::Read2::Intron3'::3’ΔHygR + 5’ΔHygR::Intron5'::Read1::GCACCATTTCACCCC::Read2::Intron3'::3’ΔHygR + 5’ΔHygR::Intron5'::Read1::GGCCATTTTTACATG::Read2::Intron3'::3’ΔHygR + 5’ΔHygR::Intron5'::Read1::TACCGCTGTAAGGTT::Read2::Intron3'::3’ΔHygR + 5’ΔHygR::Intron5'::Read1::TATCTTTGTTACCGT::Read2::Intron3'::3’ΔHygR + 5’ΔHygR::Intron5'::Read1::TCCCACTTTAAACAA::Read2::Intron3'::3’ΔHygR + 5’ΔHygR::Intron5'::Read1::TGCCCGTTTAATAAT::Read2::Intron3'::3’ΔHygR + 5’ΔHygR::Intron5'::Read1::AACCCCTGTCAAAAA::Read2::Intron3'::3’ΔHygR + 5’ΔHygR::Intron5'::Read1::ATCCCTTGTCACGCA::Read2::Intron3'::3’ΔHygR + 5’ΔHygR::Intron5'::Read1::CATCAGTATCATTTA::Read2::Intron3'::3’ΔHygR + 5’ΔHygR::Intron5'::Read1::CCGCGTTCTTACTGA::Read2::Intron3'::3’ΔHygR + 5’ΔHygR::Intron5'::Read1::GTGCATTTTAATGCA::Read2::Intron3'::3’ΔHygR + 5’ΔHygR::Intron5'::Read1::TCCCTTTTTTACAAA::Read2::Intron3'::3’ΔHygR + 5’ΔHygR::Intron5'::Read1::TCGCTTTATTACTAG::Read2::Intron3'::3’ΔHygR + 5’ΔHygR::Intron5'::Read1::TCTCTATATGATCCC::Read2::Intron3'::3’ΔHygR + 5’ΔHygR::Intron5'::Read1::TTGCTTTATAATCTT::Read2::Intron3'::3’ΔHygR + 5’ΔHygR::Intron5'::Read1::AGGCGATATAACTTC::Read2::Intron3'::3’ΔHygR + 5’ΔHygR::Intron5'::Read1::ATCCACTCTTACCCA::Read2::Intron3'::3’ΔHygR + 5’ΔHygR::Intron5'::Read1::ATTCGCTCTCAAATA::Read2::Intron3'::3’ΔHygR + 5’ΔHygR::Intron5'::Read1::CTTCCATTTCAAGGA::Read2::Intron3'::3’ΔHygR + 5’ΔHygR::Intron5'::Read1::CTTCGCTCTTAAGCT::Read2::Intron3'::3’ΔHygR + 5’ΔHygR::Intron5'::Read1::GCTCACTGTAATTCA::Read2::Intron3'::3’ΔHygR + 5’ΔHygR::Intron5'::Read1::TACCACTCTGAACGA::Read2::Intron3'::3’ΔHygR + 5’ΔHygR::Intron5'::Read1::TTCCGGTCTCATGTA::Read2::Intron3'::3’ΔHygR + 5’ΔHygR::Intron5'::Read1::TTGCTTTGTAACCTC::Read2::Intron3'::3’ΔHygR + 5’ΔHygR::Intron5'::Read1::ACGCTTTCTTACACA::Read2::Intron3'::3’ΔHygR + 5’ΔHygR::Intron5'::Read1::AGCCACTTTTACGTC::Read2::Intron3'::3’ΔHygR + 5’ΔHygR::Intron5'::Read1::ATCCAATCTAATTTC::Read2::Intron3'::3’ΔHygR + 5’ΔHygR::Intron5'::Read1::CAACGCTGTAATCTT::Read2::Intron3'::3’ΔHygR + 5’ΔHygR::Intron5'::Read1::CCACGGTGTTATTGA::Read2::Intron3'::3’ΔHygR + 5’ΔHygR::Intron5'::Read1::CCACTATTTTAACTA::Read2::Intron3'::3’ΔHygR + 5’ΔHygR::Intron5'::Read1::GATCCGTGTGACTGC::Read2::Intron3'::3’ΔHygR + 5’ΔHygR::Intron5'::Read1::GGACAATGTCATAAG::Read2::Intron3'::3’ΔHygR + 5’ΔHygR::Intron5'::Read1::GGACGATATTATTGA::Read2::Intron3'::3’ΔHygR + 5’ΔHygR::Intron5'::Read1::TCCCTCTTTCACTAT::Read2::Intron3'::3’ΔHygR + 5’ΔHygR::Intron5'::Read1::TGGCTTTTTCATCTC::Read2::Intron3'::3’ΔHygR + 5’ΔHygR::Intron5'::Read1::TTCCGATTTTAATAC::Read2::Intron3'::3’ΔHygR + 5’ΔHygR::Intron5'::Read1::TTGCAGTATGAGTGC::Read2::Intron3'::3’ΔHygR + 5’ΔHygR::Intron5'::Read1::ACTCACTTTTAATCC::Read2::Intron3'::3’ΔHygR + 5’ΔHygR::Intron5'::Read1::ATGCTTTCTCACGTC::Read2::Intron3'::3’ΔHygR + 5’ΔHygR::Intron5'::Read1::GCACTTTTTTATTAC::Read2::Intron3'::3’ΔHygR + 5’ΔHygR::Intron5'::Read1::TAACACTATGATCAC::Read2::Intron3'::3’ΔHygR + 5’ΔHygR::Intron5'::Read1::TACCGATCTTAATCC::Read2::Intron3'::3’ΔHygR + 5’ΔHygR::Intron5'::Read1::TCCCGATCTTAACAG::Read2::Intron3'::3’ΔHygR + 5’ΔHygR::Intron5'::Read1::TCTCCACATGATCAC::Read2::Intron3'::3’ΔHygR + 5’ΔHygR::Intron5'::Read1::TTTCTGTTTTATTAC::Read2::Intron3'::3’ΔHygR + 5’ΔHygR::Intron5'::Read1::CCTCTCTCTAATCAG::Read2::Intron3'::3’ΔHygR + 5’ΔHygR::Intron5'::Read1::CTGCCTTATGATTTA::Read2::Intron3'::3’ΔHygR + 5’ΔHygR::Intron5'::Read1::CTTCGTTCTTAACAG::Read2::Intron3'::3’ΔHygR +

5’ΔHygR::Intron5'::Read1::GAACTCTCTCATGCC::Read2::Intron3'::3’ΔHygR + 5’ΔHygR::Intron5'::Read1::GGTCATTCTTAAACT::Read2::Intron3'::3’ΔHygR + 5’ΔHygR::Intron5'::Read1::AGTCAGTATTACTTC::Read2::Intron3'::3’ΔHygR + 5’ΔHygR::Intron5'::Read1::CAGCATTATTAGTCT::Read2::Intron3'::3’ΔHygR + 5’ΔHygR::Intron5'::Read1::CGACAATTTGACTCT::Read2::Intron3'::3’ΔHygR + 5’ΔHygR::Intron5'::Read1::GTGCATTTTGACCGT::Read2::Intron3'::3’ΔHygR + 5’ΔHygR::Intron5'::Read1::GTTCTCTTTAATTAT::Read2::Intron3'::3’ΔHygR + 5’ΔHygR::Intron5'::Read1::TGGCCTTATGACGCT::Read2::Intron3'::3’ΔHygR + 5’ΔHygR::Intron5'::Read1::AGACCTTGTAATCAC::Read2::Intron3'::3’ΔHygR + 5’ΔHygR::Intron5'::Read1::CCGCCCTTTAATCCC::Read2::Intron3'::3’ΔHygR + 5’ΔHygR::Intron5'::Read1::CGACTTTTTTACTAG::Read2::Intron3'::3’ΔHygR + 5’ΔHygR::Intron5'::Read1::CGTCGTTCTGATCGC::Read2::Intron3'::3’ΔHygR + 5’ΔHygR::Intron5'::Read1::GCTCGTTCTCAGTCG::Read2::Intron3'::3’ΔHygR + 5’ΔHygR::Intron5'::Read1::GCTCGTTTTTATTCC::Read2::Intron3'::3’ΔHygR + 5’ΔHygR::Intron5'::Read1::GGCCATTCTAACACT::Read2::Intron3'::3’ΔHygR + 5’ΔHygR::Intron5'::Read1::GGTCCATATCACTAA::Read2::Intron3'::3’ΔHygR + 5’ΔHygR::Intron5'::Read1::GTGCGATTTAATCAT::Read2::Intron3'::3’ΔHygR + 5’ΔHygR::Intron5'::Read1::TACCGCTGTCACCTT::Read2::Intron3'::3’ΔHygR + 5’ΔHygR::Intron5'::Read1::AATCCCTGTCAAGCA::Read2::Intron3'::3’ΔHygR + 5’ΔHygR::Intron5'::Read1::AATCGATGTAAAACC::Read2::Intron3'::3’ΔHygR + 5’ΔHygR::Intron5'::Read1::AGACGATGTCACCCC::Read2::Intron3'::3’ΔHygR + 5’ΔHygR::Intron5'::Read1::ATCCGTTCTCAATTA::Read2::Intron3'::3’ΔHygR + 5’ΔHygR::Intron5'::Read1::CATCCATTTAATACC::Read2::Intron3'::3’ΔHygR + 5’ΔHygR::Intron5'::Read1::CCGCATTTTAACGCG::Read2::Intron3'::3’ΔHygR + 5’ΔHygR::Intron5'::Read1::GCACGCTCTCACATA::Read2::Intron3'::3’ΔHygR + 5’ΔHygR::Intron5'::Read1::TAGCGATTTTATCCC::Read2::Intron3'::3’ΔHygR + 5’ΔHygR::Intron5'::Read1::TGACCCTCTTATAGT::Read2::Intron3'::3’ΔHygR + 5’ΔHygR::Intron5'::Read1::TGACGATTTCACTCC::Read2::Intron3'::3’ΔHygR + 5’ΔHygR::Intron5'::Read1::TGTCGATTTTATCTA::Read2::Intron3'::3’ΔHygR + 5’ΔHygR::Intron5'::Read1::ACACACTTTGAATTT::Read2::Intron3'::3’ΔHygR + 5’ΔHygR::Intron5'::Read1::ACCCCGTTTAATTGT::Read2::Intron3'::3’ΔHygR + 5’ΔHygR::Intron5'::Read1::AGCCAATCTGAATGC::Read2::Intron3'::3’ΔHygR + 5’ΔHygR::Intron5'::Read1::GCGCGGTTTCACCAA::Read2::Intron3'::3’ΔHygR + 5’ΔHygR::Intron5'::Read1::AGTCTTTTTTAAACC::Read2::Intron3'::3’ΔHygR + 5’ΔHygR::Intron5'::Read1::CAACCTTATCATATT::Read2::Intron3'::3’ΔHygR + 5’ΔHygR::Intron5'::Read1::CCGCGTTATTACTTT::Read2::Intron3'::3’ΔHygR + 5’ΔHygR::Intron5'::Read1::CTACACTATTAAAAC::Read2::Intron3'::3’ΔHygR + 5’ΔHygR::Intron5'::Read1::GATCATTGTCAATTC::Read2::Intron3'::3’ΔHygR + 5’ΔHygR::Intron5'::Read1::GCCCTATCTTATCAT::Read2::Intron3'::3’ΔHygR + 5’ΔHygR::Intron5'::Read1::GCGCGCTATCAACTC::Read2::Intron3'::3’ΔHygR + 5’ΔHygR::Intron5'::Read1::GTCCGCTTTAATTGA::Read2::Intron3'::3’ΔHygR + 5’ΔHygR::Intron5'::Read1::TTTCCCTTTGATGGC::Read2::Intron3'::3’ΔHygR + 5’ΔHygR::Intron5'::Read1::AAGCCTTGTCAGCGA::Read2::Intron3'::3’ΔHygR + 5’ΔHygR::Intron5'::Read1::ACCCGTTATTACTAA::Read2::Intron3'::3’ΔHygR + 5’ΔHygR::Intron5'::Read1::CATCGTTATTACTCA::Read2::Intron3'::3’ΔHygR + 5’ΔHygR::Intron5'::Read1::GACCCATATCACTCA::Read2::Intron3'::3’ΔHygR + 5’ΔHygR::Intron5'::Read1::GACCTCTCTGAAGAT::Read2::Intron3'::3’ΔHygR + 5’ΔHygR::Intron5'::Read1::GTCCGATATCAGCAA::Read2::Intron3'::3’ΔHygR + 5’ΔHygR::Intron5'::Read1::TAGCCTTTTAATCGT::Read2::Intron3'::3’ΔHygR + 5’ΔHygR::Intron5'::Read1::AACCCTTGTCACTAA::Read2::Intron3'::3’ΔHygR + 5’ΔHygR::Intron5'::Read1::ACACCCTTTTAAAGT::Read2::Intron3'::3’ΔHygR + 5’ΔHygR::Intron5'::Read1::ATACACTCTTAAATG::Read2::Intron3'::3’ΔHygR + 5’ΔHygR::Intron5'::Read1::ATACCGTATGACAAA::Read2::Intron3'::3’ΔHygR + 5’ΔHygR::Intron5'::Read1::CACCAATATAATCAA::Read2::Intron3'::3’ΔHygR + 5’ΔHygR::Intron5'::Read1::CTGCTTTCTGACCAA::Read2::Intron3'::3’ΔHygR + 5’ΔHygR::Intron5'::Read1::GTCCATTATAATCCT::Read2::Intron3'::3’ΔHygR + 5’ΔHygR::Intron5'::Read1::TCACATTATCATACT::Read2::Intron3'::3’ΔHygR + 5’ΔHygR::Intron5'::Read1::TCCCACTTTAAGAAT::Read2::Intron3'::3’ΔHygR + 5’ΔHygR::Intron5'::Read1::TGTCCCTTTTAACTA::Read2::Intron3'::3’ΔHygR + 5’ΔHygR::Intron5'::Read1::ACACCCTATCAAATA::Read2::Intron3'::3’ΔHygR + 5’ΔHygR::Intron5'::Read1::AGACTTTTTAATGCT::Read2::Intron3'::3’ΔHygR + 5’ΔHygR::Intron5'::Read1::CAACCTTATGATCGG::Read2::Intron3'::3’ΔHygR + 5’ΔHygR::Intron5'::Read1::CCTCACTATCAGCGC::Read2::Intron3'::3’ΔHygR + 5’ΔHygR::Intron5'::Read1::TAACGATTTTACGTT::Read2::Intron3'::3’ΔHygR + 5’ΔHygR::Intron5'::Read1::ATACGTTCTAATGGT::Read2::Intron3'::3’ΔHygR + 5’ΔHygR::Intron5'::Read1::ATACTATCTCAAACA::Read2::Intron3'::3’ΔHygR + 5’ΔHygR::Intron5'::Read1::CATCAATATGAAACC::Read2::Intron3'::3’ΔHygR + 5’ΔHygR::Intron5'::Read1::CTACACTTTGAGTCC::Read2::Intron3'::3’ΔHygR + 5’ΔHygR::Intron5'::Read1::GACCCATATCAGGCA::Read2::Intron3'::3’ΔHygR + 5’ΔHygR::Intron5'::Read1::GCCCACTCTAATATA::Read2::Intron3'::3’ΔHygR +

5’ΔHygR::Intron5'::Read1::GGTCACTCTTAAGTT::Read2::Intron3'::3’ΔHygR + 5’ΔHygR::Intron5'::Read1::TACCGTTCTGAAAAT::Read2::Intron3'::3’ΔHygR + 5’ΔHygR::Intron5'::Read1::TACCTATATAATGCA::Read2::Intron3'::3’ΔHygR + 5’ΔHygR::Intron5'::Read1::TAGCGTTTTAATTCA::Read2::Intron3'::3’ΔHygR + 5’ΔHygR::Intron5'::Read1::TTACGTTATCACGCT::Read2::Intron3'::3’ΔHygR + 5’ΔHygR::Intron5'::Read1::TTTCCTTTTAACAAT::Read2::Intron3'::3’ΔHygR + 5’ΔHygR::Intron5'::Read1::GATCTTTGTTATCGA::Read2::Intron3'::3’ΔHygR + 5’ΔHygR::Intron5'::Read1::GTGCCTTCTTATTTT::Read2::Intron3'::3’ΔHygR + 5’ΔHygR::Intron5'::Read1::TACCTCTCTTAGTAA::Read2::Intron3'::3’ΔHygR + 5’ΔHygR::Intron5'::Read1::AAGCGCTTTAATCTA::Read2::Intron3'::3’ΔHygR + 5’ΔHygR::Intron5'::Read1::CACCTTTCTGAAACA::Read2::Intron3'::3’ΔHygR + 5’ΔHygR::Intron5'::Read1::CTACTTTATGAATGT::Read2::Intron3'::3’ΔHygR + 5’ΔHygR::Intron5'::Read1::CTTCTTTCTTAAAAT::Read2::Intron3'::3’ΔHygR + 5’ΔHygR::Intron5'::Read1::GCACCCTATTATCCT::Read2::Intron3'::3’ΔHygR + 5’ΔHygR::Intron5'::Read1::GTGCTGTTTTAGATT::Read2::Intron3'::3’ΔHygR + 5’ΔHygR::Intron5'::Read1::TAGCTGTGTCACTCC::Read2::Intron3'::3’ΔHygR + 5’ΔHygR::Intron5'::Read1::TAGCTTTGTCAATTT::Read2::Intron3'::3’ΔHygR + 5’ΔHygR::Intron5'::Read1::AATCCATCTTACGGC::Read2::Intron3'::3’ΔHygR + 5’ΔHygR::Intron5'::Read1::AATCCTTTTTATACC::Read2::Intron3'::3’ΔHygR + 5’ΔHygR::Intron5'::Read1::AGACTGTGTCACACG::Read2::Intron3'::3’ΔHygR + 5’ΔHygR::Intron5'::Read1::ATACTCTTTTATAGT::Read2::Intron3'::3’ΔHygR + 5’ΔHygR::Intron5'::Read1::CACCCCTATGACGCA::Read2::Intron3'::3’ΔHygR + 5’ΔHygR::Intron5'::Read1::CCACTCTGTCACATA::Read2::Intron3'::3’ΔHygR + 5’ΔHygR::Intron5'::Read1::CCACTTTATTACTAC::Read2::Intron3'::3’ΔHygR + 5’ΔHygR::Intron5'::Read1::CCCCTCTCTAAATAG::Read2::Intron3'::3’ΔHygR + 5’ΔHygR::Intron5'::Read1::TCCCGATCTCAATTT::Read2::Intron3'::3’ΔHygR + 5’ΔHygR::Intron5'::Read1::AAGCCTTTTAAATCG::Read2::Intron3'::3’ΔHygR + 5’ΔHygR::Intron5'::Read1::ATACTTTGTAAAAAA::Read2::Intron3'::3’ΔHygR + 5’ΔHygR::Intron5'::Read1::CCCCTATATGAAAAA::Read2::Intron3'::3’ΔHygR + 5’ΔHygR::Intron5'::Read1::TGACATTGTAAACCC::Read2::Intron3'::3’ΔHygR + 5’ΔHygR::Intron5'::Read1::ACCCAATCTCATCTA::Read2::Intron3'::3’ΔHygR + 5’ΔHygR::Intron5'::Read1::CACCTATTTCAGCAG::Read2::Intron3'::3’ΔHygR + 5’ΔHygR::Intron5'::Read1::CGTCTGTTTAAAATG::Read2::Intron3'::3’ΔHygR + 5’ΔHygR::Intron5'::Read1::GGTCTCTTTGATCTT::Read2::Intron3'::3’ΔHygR + 5’ΔHygR::Intron5'::Read1::GTTCATTATTAAACG::Read2::Intron3'::3’ΔHygR + 5’ΔHygR::Intron5'::Read1::TAACTTTCTCATAGG::Read2::Intron3'::3’ΔHygR + 5’ΔHygR::Intron5'::Read1::AACCTTTATAACCCA::Read2::Intron3'::3’ΔHygR + 5’ΔHygR::Intron5'::Read1::CACCCCTTTCATAGA::Read2::Intron3'::3’ΔHygR + 5’ΔHygR::Intron5'::Read1::CTCCCGTGTTATTAT::Read2::Intron3'::3’ΔHygR + 5’ΔHygR::Intron5'::Read1::GAACTCTTTAATAGA::Read2::Intron3'::3’ΔHygR + 5’ΔHygR::Intron5'::Read1::GATCGTTCTTATTTA::Read2::Intron3'::3’ΔHygR + 5’ΔHygR::Intron5'::Read1::TGCCCATGTGAATAC::Read2::Intron3'::3’ΔHygR + 5’ΔHygR::Intron5'::Read1::AACCCATTTAAAGTC::Read2::Intron3'::3’ΔHygR + 5’ΔHygR::Intron5'::Read1::AGACGCTATGACGGA::Read2::Intron3'::3’ΔHygR + 5’ΔHygR::Intron5'::Read1::ATCCACTCTGAACGG::Read2::Intron3'::3’ΔHygR + 5’ΔHygR::Intron5'::Read1::CACCTCTTTTACTAT::Read2::Intron3'::3’ΔHygR + 5’ΔHygR::Intron5'::Read1::CTCCCATGTTACCAC::Read2::Intron3'::3’ΔHygR + 5’ΔHygR::Intron5'::Read1::GACCGATGTTAACAG::Read2::Intron3'::3’ΔHygR + 5’ΔHygR::Intron5'::Read1::GGTCTTTCTAAAACT::Read2::Intron3'::3’ΔHygR + 5’ΔHygR::Intron5'::Read1::TAGCCCTTTAAACTC::Read2::Intron3'::3’ΔHygR + 5’ΔHygR::Intron5'::Read1::ACTCAGTTTGACATC::Read2::Intron3'::3’ΔHygR + 5’ΔHygR::Intron5'::Read1::AGACCATATTATGAT::Read2::Intron3'::3’ΔHygR + 5’ΔHygR::Intron5'::Read1::ATCCCATATGACTTA::Read2::Intron3'::3’ΔHygR + 5’ΔHygR::Intron5'::Read1::CATCCTTATTATATT::Read2::Intron3'::3’ΔHygR + 5’ΔHygR::Intron5'::Read1::CGCCCCTGTCAATAC::Read2::Intron3'::3’ΔHygR + 5’ΔHygR::Intron5'::Read1::GATCTCTTTAAGAAA::Read2::Intron3'::3’ΔHygR + 5’ΔHygR::Intron5'::Read1::AAACATTCTTAGGGA::Read2::Intron3'::3’ΔHygR + 5’ΔHygR::Intron5'::Read1::AAGCCTTGTGAGTTT::Read2::Intron3'::3’ΔHygR + 5’ΔHygR::Intron5'::Read1::ACCCGGTGTTATTTT::Read2::Intron3'::3’ΔHygR + 5’ΔHygR::Intron5'::Read1::AGGCTGTCTAATTTA::Read2::Intron3'::3’ΔHygR + 5’ΔHygR::Intron5'::Read1::CCCCTTTATAATGTA::Read2::Intron3'::3’ΔHygR + 5’ΔHygR::Intron5'::Read1::CTTCCGTATCACTAA::Read2::Intron3'::3’ΔHygR + 5’ΔHygR::Intron5'::Read1::GCTCAGTGTAATGCC::Read2::Intron3'::3’ΔHygR + 5’ΔHygR::Intron5'::Read1::GTCCATTATTAGCGT::Read2::Intron3'::3’ΔHygR + 5’ΔHygR::Intron5'::Read1::GTTCCTTATTATCAG::Read2::Intron3'::3’ΔHygR + 5’ΔHygR::Intron5'::Read1::TATCGTTATAACCTT::Read2::Intron3'::3’ΔHygR + 5’ΔHygR::Intron5'::Read1::CGTCTCTGTCAGCGC::Read2::Intron3'::3’ΔHygR + 5’ΔHygR::Intron5'::Read1::TCTCAGTCTAATTTG::Read2::Intron3'::3’ΔHygR + 5’ΔHygR::Intron5'::Read1::AGACCTTATCACGAA::Read2::Intron3'::3’ΔHygR + 5’ΔHygR::Intron5'::Read1::CTTCTCTGTCACGCA::Read2::Intron3'::3’ΔHygR +

5’ΔHygR::Intron5'::Read1::TTACGATATTACATC::Read2::Intron3'::3’ΔHygR + 5’ΔHygR::Intron5'::Read1::TATCTATCTTACACA::Read2::Intron3'::3’ΔHygR + 5’ΔHygR::Intron5'::Read1::TCCCCGTTTCAAATT::Read2::Intron3'::3’ΔHygR + 5’ΔHygR::Intron5'::Read1::TGGCCTTTTAAGGTA::Read2::Intron3'::3’ΔHygR + 5’ΔHygR::Intron5'::Read1::TGTCATTATGATTAT::Read2::Intron3'::3’ΔHygR + 5’ΔHygR::Intron5'::Read1::ATCCCTTATCACTCT::Read2::Intron3'::3’ΔHygR + 5’ΔHygR::Intron5'::Read1::CATCCGTGTCAATAT::Read2::Intron3'::3’ΔHygR + 5’ΔHygR::Intron5'::Read1::GCCCGATATCATCAT::Read2::Intron3'::3’ΔHygR + 5’ΔHygR::Intron5'::Read1::TCTCTCTCTTACATT::Read2::Intron3'::3’ΔHygR + 5’ΔHygR::Intron5'::Read1::TTCCCGTTTCAGCAT::Read2::Intron3'::3’ΔHygR + 5’ΔHygR::Intron5'::Read1::AACCTTTCTTAAAAA::Read2::Intron3'::3’ΔHygR + 5’ΔHygR::Intron5'::Read1::AGGCGCTTTTAACCA::Read2::Intron3'::3’ΔHygR + 5’ΔHygR::Intron5'::Read1::CCACTATATGACAAC::Read2::Intron3'::3’ΔHygR + 5’ΔHygR::Intron5'::Read1::CCCCCATATGAGCTC::Read2::Intron3'::3’ΔHygR + 5’ΔHygR::Intron5'::Read1::CTACTTTCTCAAATG::Read2::Intron3'::3’ΔHygR + 5’ΔHygR::Intron5'::Read1::CTGCATTCTCACTAT::Read2::Intron3'::3’ΔHygR + 5’ΔHygR::Intron5'::Read1::TAACACTTTTAAGTC::Read2::Intron3'::3’ΔHygR + 5’ΔHygR::Intron5'::Read1::TGGCGGTCTTACGAA::Read2::Intron3'::3’ΔHygR + 5’ΔHygR::Intron5'::Read1::TTCCCATATTATGAT::Read2::Intron3'::3’ΔHygR + 5’ΔHygR::Intron5'::Read1::AGACCCTTTTACTTT::Read2::Intron3'::3’ΔHygR + 5’ΔHygR::Intron5'::Read1::ATTCCATTTGACTTG::Read2::Intron3'::3’ΔHygR + 5’ΔHygR::Intron5'::Read1::CACCGATATGAATAA::Read2::Intron3'::3’ΔHygR + 5’ΔHygR::Intron5'::Read1::CGACACTGTTAGCGC::Read2::Intron3'::3’ΔHygR + 5’ΔHygR::Intron5'::Read1::GCGCAATATTAATAC::Read2::Intron3'::3’ΔHygR + 5’ΔHygR::Intron5'::Read1::TCTCAATCTAAGTAC::Read2::Intron3'::3’ΔHygR + 5’ΔHygR::Intron5'::Read1::CTGCGGTATCAGTAC::Read2::Intron3'::3’ΔHygR + 5’ΔHygR::Intron5'::Read1::GGACGCTTTAATGAC::Read2::Intron3'::3’ΔHygR + 5’ΔHygR::Intron5'::Read1::TCACTTTGTGATATC::Read2::Intron3'::3’ΔHygR + 5’ΔHygR::Intron5'::Read1::TCGCGCTCTAACACA::Read2::Intron3'::3’ΔHygR + 5’ΔHygR::Intron5'::Read1::TGACTTTCTAAGCTG::Read2::Intron3'::3’ΔHygR + 5’ΔHygR::Intron5'::Read1::CGTCAATGTAAGGTT::Read2::Intron3'::3’ΔHygR + 5’ΔHygR::Intron5'::Read1::GGACGTTGTTAGTTT::Read2::Intron3'::3’ΔHygR + 5’ΔHygR::Intron5'::Read1::TCCCGCTATCAGACT::Read2::Intron3'::3’ΔHygR + 5’ΔHygR::Intron5'::Read1::TTGCTCTATTATGAT::Read2::Intron3'::3’ΔHygR + 5’ΔHygR::Intron5'::Read1::AATCATTCTAACCAA::Read2::Intron3'::3’ΔHygR + 5’ΔHygR::Intron5'::Read1::ATACACTCTTATCCC::Read2::Intron3'::3’ΔHygR + 5’ΔHygR::Intron5'::Read1::ATGCGTTCTCATTCG::Read2::Intron3'::3’ΔHygR + 5’ΔHygR::Intron5'::Read1::ATTCAGTATAAATAT::Read2::Intron3'::3’ΔHygR + 5’ΔHygR::Intron5'::Read1::CACCACTATAACGCG::Read2::Intron3'::3’ΔHygR + 5’ΔHygR::Intron5'::Read1::CGGCACTCTCACCTG::Read2::Intron3'::3’ΔHygR + 5’ΔHygR::Intron5'::Read1::GAACCCTTTAAAAGA::Read2::Intron3'::3’ΔHygR + 5’ΔHygR::Intron5'::Read1::GCCCTATTTAATGTA::Read2::Intron3'::3’ΔHygR + 5’ΔHygR::Intron5'::Read1::GCCCTCTGTCAACAC::Read2::Intron3'::3’ΔHygR + 5’ΔHygR::Intron5'::Read1::AGCCACTTTGACAAT::Read2::Intron3'::3’ΔHygR + 5’ΔHygR::Intron5'::Read1::TACCATTTTCATGTG::Read2::Intron3'::3’ΔHygR + 5’ΔHygR::Intron5'::Read1::ACTCAGTGTGACTGG::Read2::Intron3'::3’ΔHygR + 5’ΔHygR::Intron5'::Read1::ATGCATTATCACACA::Read2::Intron3'::3’ΔHygR + 5’ΔHygR::Intron5'::Read1::CAATTGTCTTACCAT::Read2::Intron3'::3’ΔHygR + 5’ΔHygR::Intron5'::Read1::CAACACTCTCATTTA::Read2::Intron3'::3’ΔHygR + 5’ΔHygR::Intron5'::Read1::GAGCCATATTATTTG::Read2::Intron3'::3’ΔHygR + 5’ΔHygR::Intron5'::Read1::TCACTATTTAACCGA::Read2::Intron3'::3’ΔHygR + 5’ΔHygR::Intron5'::Read1::TGTCTATATTACCAT::Read2::Intron3'::3’ΔHygR + 5’ΔHygR::Intron5'::Read1::GAGCGTTTTTATTAT::Read2::Intron3'::3’ΔHygR + 5’ΔHygR::Intron5'::Read1::GTTCGCTGTGAATCA::Read2::Intron3'::3’ΔHygR + 5’ΔHygR::Intron5'::Read1::TCTCACTATTACCCC::Read2::Intron3'::3’ΔHygR + 5’ΔHygR::Intron5'::Read1::TTACATTTTGATTGA::Read2::Intron3'::3’ΔHygR + 5’ΔHygR::Intron5'::Read1::TTACGATGTAAACGT::Read2::Intron3'::3’ΔHygR + 5’ΔHygR::Intron5'::Read1::TTACTTTTTAAACCC::Read2::Intron3'::3’ΔHygR + 5’ΔHygR::Intron5'::Read1::ACACTGTGTTAAACC::Read2::Intron3'::3’ΔHygR + 5’ΔHygR::Intron5'::Read1::GCTCCTTATCAGGAG::Read2::Intron3'::3’ΔHygR + 5’ΔHygR::Intron5'::Read1::GTGCCTTTTAACGAC::Read2::Intron3'::3’ΔHygR + 5’ΔHygR::Intron5'::Read1::ATCCCGTGTAAAAAG::Read2::Intron3'::3’ΔHygR + 5’ΔHygR::Intron5'::Read1::ATCCTGTCTTACCAC::Read2::Intron3'::3’ΔHygR + 5’ΔHygR::Intron5'::Read1::ATGCGCTTTAATTAT::Read2::Intron3'::3’ΔHygR + 5’ΔHygR::Intron5'::Read1::CGCCAATCTAAGTAT::Read2::Intron3'::3’ΔHygR + 5’ΔHygR::Intron5'::Read1::GCTCAATATCAATTA::Read2::Intron3'::3’ΔHygR + 5’ΔHygR::Intron5'::Read1::TACCTCTCTGAATAC::Read2::Intron3'::3’ΔHygR + 5’ΔHygR::Intron5'::Read1::TCGCAATATAACACG::Read2::Intron3'::3’ΔHygR + 5’ΔHygR::Intron5'::Read1::TGTCCCTCTAAGCCA::Read2::Intron3'::3’ΔHygR + 5’ΔHygR::Intron5'::Read1::CCCCTCTCTCATCCG::Read2::Intron3'::3’ΔHygR +

5’ΔHygR::Intron5'::Read1::CGCCATTATTATCTG::Read2::Intron3'::3’ΔHygR + 5’ΔHygR::Intron5'::Read1::CTCCCATCTAATTCC::Read2::Intron3'::3’ΔHygR + 5’ΔHygR::Intron5'::Read1::GATCTTTTTTATTGT::Read2::Intron3'::3’ΔHygR + 5’ΔHygR::Intron5'::Read1::TCTCTTTATCACACA::Read2::Intron3'::3’ΔHygR + 5’ΔHygR::Intron5'::Read1::TTCCGCTATAATCGC::Read2::Intron3'::3’ΔHygR + 5’ΔHygR::Intron5'::Read1::ACACTCTATGAGATT::Read2::Intron3'::3’ΔHygR + 5’ΔHygR::Intron5'::Read1::CACCACTTTGAAGAA::Read2::Intron3'::3’ΔHygR + 5’ΔHygR::Intron5'::Read1::CACCCTTTTTATGAA::Read2::Intron3'::3’ΔHygR + 5’ΔHygR::Intron5'::Read1::CTGCCGTCTGATAGA::Read2::Intron3'::3’ΔHygR + 5’ΔHygR::Intron5'::Read1::CTGCGTTCTAACAGT::Read2::Intron3'::3’ΔHygR + 5’ΔHygR::Intron5'::Read1::GAACATTGTCACTCA::Read2::Intron3'::3’ΔHygR + 5’ΔHygR::Intron5'::Read1::TTTCCTTGTGATTTA::Read2::Intron3'::3’ΔHygR + 5’ΔHygR::Intron5'::Read1::TTTCTTTATGACCGC::Read2::Intron3'::3’ΔHygR + 5’ΔHygR::Intron5'::Read1::CAACATTTTTATACA::Read2::Intron3'::3’ΔHygR + 5’ΔHygR::Intron5'::Read1::CCGCAGTATAATACT::Read2::Intron3'::3’ΔHygR + 5’ΔHygR::Intron5'::Read1::GGTCCATGTTAGTCA::Read2::Intron3'::3’ΔHygR + 5’ΔHygR::Intron5'::Read1::TAGCCATTTTATTAT::Read2::Intron3'::3’ΔHygR + 5’ΔHygR::Intron5'::Read1::TGGCTTTATTATTCG::Read2::Intron3'::3’ΔHygR + 5’ΔHygR::Intron5'::Read1::CCACACTATGAATGC::Read2::Intron3'::3’ΔHygR + 5’ΔHygR::Intron5'::Read1::GCGCCTTTTAATTTT::Read2::Intron3'::3’ΔHygR + 5’ΔHygR::Intron5'::Read1::CTACTTTCTTACAAT::Read2::Intron3'::3’ΔHygR + 5’ΔHygR::Intron5'::Read1::CTTCTCTATGATTTA::Read2::Intron3'::3’ΔHygR + 5’ΔHygR::Intron5'::Read1::GCGCCATCTGATAAA::Read2::Intron3'::3’ΔHygR + 5’ΔHygR::Intron5'::Read1::AACCGCTTTGACGCC::Read2::Intron3'::3’ΔHygR + 5’ΔHygR::Intron5'::Read1::AGTCATTTTAAGTAC::Read2::Intron3'::3’ΔHygR + 5’ΔHygR::Intron5'::Read1::TGCCATTCTTATTTA::Read2::Intron3'::3’ΔHygR + 5’ΔHygR::Intron5'::Read1::AACCTTTATCACCTT::Read2::Intron3'::3’ΔHygR + 5’ΔHygR::Intron5'::Read1::CAACAGTCTTATAAA::Read2::Intron3'::3’ΔHygR + 5’ΔHygR::Intron5'::Read1::CAACCATATTACTCC::Read2::Intron3'::3’ΔHygR + 5’ΔHygR::Intron5'::Read1::GGACTATTTCAGTTG::Read2::Intron3'::3’ΔHygR + 5’ΔHygR::Intron5'::Read1::CCCCCATGTTAGTGC::Read2::Intron3'::3’ΔHygR + 5’ΔHygR::Intron5'::Read1::CGACAGTATGACGCC::Read2::Intron3'::3’ΔHygR + 5’ΔHygR::Intron5'::Read1::GTTCTATATTACGCT::Read2::Intron3'::3’ΔHygR + 5’ΔHygR::Intron5'::Read1::ATACATTTTCAGCGC::Read2::Intron3'::3’ΔHygR + 5’ΔHygR::Intron5'::Read1::AGACTGTATGAGGAT::Read2::Intron3'::3’ΔHygR + 5’ΔHygR::Intron5'::Read1::CAACCCTGTTATCGT::Read2::Intron3'::3’ΔHygR + 5’ΔHygR::Intron5'::Read1::CTGCAGTGTCAACCC::Read2::Intron3'::3’ΔHygR + 5’ΔHygR::Intron5'::Read1::GCACGCTCTGATATT::Read2::Intron3'::3’ΔHygR + 5’ΔHygR::Intron5'::Read1::TCCCAATGTAAAAGT::Read2::Intron3'::3’ΔHygR + 5’ΔHygR::Intron5'::Read1::ATCCTATGTAAATTC::Read2::Intron3'::3’ΔHygR + 5’ΔHygR::Intron5'::Read1::CTGCAGTGTTAGGCC::Read2::Intron3'::3’ΔHygR + 5’ΔHygR::Intron5'::Read1::CTGCGTTTTCATCAA::Read2::Intron3'::3’ΔHygR + 5’ΔHygR::Intron5'::Read1::GCGCTGTGTGATCAA::Read2::Intron3'::3’ΔHygR + 5’ΔHygR::Intron5'::Read1::CACCGCTTTTAACTT::Read2::Intron3'::3’ΔHygR + 5’ΔHygR::Intron5'::Read1::ATGCCGTTTTAATTT::Read2::Intron3'::3’ΔHygR + 5’ΔHygR::Intron5'::Read1::GGCCTCTCTGCACTC::Read2::Intron3'::3’ΔHygR + 5’ΔHygR::Intron5'::Read1::TTCCCTATGTCAGCG::Read2::Intron3'::3’ΔHygR + 5’ΔHygR::Intron5'::Read1::GTGCTGTTGTGAGCG::Read2::Intron3'::3’ΔHygR + 5’ΔHygR::Intron5'::Read1::GTTCCCTCTAACGTC::Read2::Intron3'::3’ΔHygR + 5’ΔHygR::Intron5'::Read1::ACGCGGTCTCACTAA::Read2::Intron3'::3’ΔHygR + 5’ΔHygR::Intron5'::Read1::ATATTAACATGGTGC::Read2::Intron3'::3’ΔHygR + 5’ΔHygR::Intron5'::Read1::ATCTTTATACCGTAA::Read2::Intron3'::3’ΔHygR + 5’ΔHygR::Intron5'::Read1::GCCCGCTCTGAGTAT::Read2::Intron3'::3’ΔHygR + 5’ΔHygR::Intron5'::Read1::TACCACTATGAACCT::Read2::Intron3'::3’ΔHygR + 5’ΔHygR::Intron5'::Read1::TGTTTCAGAGAGAGT::Read2::Intron3'::3’ΔHygR + 5’ΔHygR::Intron5'::Read1::CAACTCTTTTACCGT::Read2::Intron3'::3’ΔHygR + 5’ΔHygR::Intron5'::Read1::GTATTAAGATAGGCT::Read2::Intron3'::3’ΔHygR + 5’ΔHygR::Intron5'::Read1::TTCCGGTGTTAGGCC::Read2::Intron3'::3’ΔHygR + 5’ΔHygR::Intron5'::Read1::CTCCCCTATTAAATC::Read2::Intron3'::3’ΔHygR + 5’ΔHygR::Intron5'::Read1::TAACATTTTTAACGA::Read2::Intron3'::3’ΔHygR + 5’ΔHygR::Intron5'::Read1::AAGCGGTTTTAGCTA::Read2::Intron3'::3’ΔHygR + 5’ΔHygR::Intron5'::Read1::TTGCCTTCTCACTCT::Read2::Intron3'::3’ΔHygR + 5’ΔHygR::Intron5'::Read1::AGTCTGTTTTAACGC::Read2::Intron3'::3’ΔHygR + 5’ΔHygR::Intron5'::Read1::TCGCTGTTTAACACC::Read2::Intron3'::3’ΔHygR + 5’ΔHygR::Intron5'::Read1::ACTCTTTATAATCAC::Read2::Intron3'::3’ΔHygR + 5’ΔHygR::Intron5'::Read1::TCACAGTATCATGAA::Read2::Intron3'::3’ΔHygR + 5’ΔHygR::Intron5'::Read1::ACCCTCTTTGAATCC::Read2::Intron3'::3’ΔHygR + 5’ΔHygR::Intron5'::Read1::TTCCCTTTTCACCAG::Read2::Intron3'::3’ΔHygR + 5’ΔHygR::Intron5'::Read1::CACCATTATGAGCTA::Read2::Intron3'::3’ΔHygR + 5’ΔHygR::Intron5'::Read1::CAACCTTTTAATGTC::Read2::Intron3'::3’ΔHygR +

5’ΔHygR::Intron5'::Read1::ATGCCATATAACTTA::Read2::Intron3'::3’ΔHygR + 5’ΔHygR::Intron5'::Read1::GAGCACTGTTATTAA::Read2::Intron3'::3’ΔHygR + 5’ΔHygR::Intron5'::Read1::TCACATTTTCATATC::Read2::Intron3'::3’ΔHygR + 5’ΔHygR::Intron5'::Read1::ACCCCATGTAACCCC::Read2::Intron3'::3’ΔHygR + 5’ΔHygR::Intron5'::Read1::ACTCAGTGTTAACAT::Read2::Intron3'::3’ΔHygR + 5’ΔHygR::Intron5'::Read1::ATTCGATTTAATCCT::Read2::Intron3'::3’ΔHygR + 5’ΔHygR::Intron5'::Read1::TGACAAATCTAAGAT::Read2::Intron3'::3’ΔHygR + 5’ΔHygR::Intron5'::Read1::GGAATCATAGAGCCA::Read2::Intron3'::3’ΔHygR + 5’ΔHygR::Intron5'::Read1::TCAGTTAGAAGGTGA::Read2::Intron3'::3’ΔHygR + 5’ΔHygR::Intron5'::Read1::TTACCATGCTTAGTA::Read2::Intron3'::3’ΔHygR + 5’ΔHygR::Intron5'::Read1::ATCCCCTGTAACATA::Read2::Intron3'::3’ΔHygR + 5’ΔHygR::Intron5'::Read1::ATGATAAGATGGGCA::Read2::Intron3'::3’ΔHygR + 5’ΔHygR::Intron5'::Read1::GATATCAAAGGGATT::Read2::Intron3'::3’ΔHygR + 5’ΔHygR::Intron5'::Read1::AAGCTTATATGGGTG::Read2::Intron3'::3’ΔHygR + 5’ΔHygR::Intron5'::Read1::AATTTAACAGGGGGT::Read2::Intron3'::3’ΔHygR + 5’ΔHygR::Intron5'::Read1::ACACGACTTAGATAA::Read2::Intron3'::3’ΔHygR + 5’ΔHygR::Intron5'::Read1::AGTATCATAGAGCTT::Read2::Intron3'::3’ΔHygR + 5’ΔHygR::Intron5'::Read1::ATCATGAAACGGTTT::Read2::Intron3'::3’ΔHygR + 5’ΔHygR::Intron5'::Read1::CTATTAAGAAAGAAT::Read2::Intron3'::3’ΔHygR + 5’ΔHygR::Intron5'::Read1::GACCTCTCTAATATC::Read2::Intron3'::3’ΔHygR + 5’ΔHygR::Intron5'::Read1::GGTGTCATAGCGAGA::Read2::Intron3'::3’ΔHygR + 5’ΔHygR::Intron5'::Read1::ACACCCTCTAAGATC::Read2::Intron3'::3’ΔHygR + 5’ΔHygR::Intron5'::Read1::CTACAATCTCACGTT::Read2::Intron3'::3’ΔHygR + 5’ΔHygR::Intron5'::Read1::GAACCATCTCAGCAC::Read2::Intron3'::3’ΔHygR + 5’ΔHygR::Intron5'::Read1::TTTCGTTATGAGATC::Read2::Intron3'::3’ΔHygR + 5’ΔHygR::Intron5'::Read1::TTCCTCTGTAACGCT::Read2::Intron3'::3’ΔHygR + 5’ΔHygR::Intron5'::Read1::CCTCTTTTTCATACC::Read2::Intron3'::3’ΔHygR + 5’ΔHygR::Intron5'::Read1::TCGCCATTTTAAACA::Read2::Intron3'::3’ΔHygR + 5’ΔHygR::Intron5'::Read1::TGGCCGTATCATACT::Read2::Intron3'::3’ΔHygR + 5’ΔHygR::Intron5'::Read1::CTGCCTTTTCAAATC::Read2::Intron3'::3’ΔHygR + 5’ΔHygR::Intron5'::Read1::TTTCTCTATTATATC::Read2::Intron3'::3’ΔHygR + 5’ΔHygR::Intron5'::Read1::GGGCTCTCTGAAGCA::Read2::Intron3'::3’ΔHygR + 5’ΔHygR::Intron5'::Read1::GTACTTTCTTATACT::Read2::Intron3'::3’ΔHygR + 5’ΔHygR::Intron5'::Read1::TGGCTCTATGATTCC::Read2::Intron3'::3’ΔHygR + 5’ΔHygR::Intron5'::Read1::TACCCTTATCAAGTC::Read2::Intron3'::3’ΔHygR + 5’ΔHygR::Intron5'::Read1::CAACGTTCTAACCCC::Read2::Intron3'::3’ΔHygR + 5’ΔHygR::Intron5'::Read1::TTTCATTCTAATAAT::Read2::Intron3'::3’ΔHygR + 5’ΔHygR::Intron5'::Read1::ATACATTATTAGAAC::Read2::Intron3'::3’ΔHygR + 5’ΔHygR::Intron5'::Read1::AAACACTGTCATGTC::Read2::Intron3'::3’ΔHygR + 5’ΔHygR::Intron5'::Read1::CACCGCTCTTAATTA::Read2::Intron3'::3’ΔHygR + 5’ΔHygR::Intron5'::Read1::TCGCACTGTAACACT::Read2::Intron3'::3’ΔHygR + 5’ΔHygR::Intron5'::Read1::TACCATTTTCACACC::Read2::Intron3'::3’ΔHygR + 5’ΔHygR::Intron5'::Read1::GATCACTTTAACTCG::Read2::Intron3'::3’ΔHygR + 5’ΔHygR::Intron5'::Read1::TAACTATTTAAGCGG::Read2::Intron3'::3’ΔHygR + 5’ΔHygR::Intron5'::Read1::TATCAATCTCATCGC::Read2::Intron3'::3’ΔHygR + 5’ΔHygR::Intron5'::Read1::CCACGGTATAATATC::Read2::Intron3'::3’ΔHygR + 5’ΔHygR::Intron5'::Read1::ACATTAAAAGCGACT::Read2::Intron3'::3’ΔHygR + 5’ΔHygR::Intron5'::Read1::ATTGTAATAGGGCGG::Read2::Intron3'::3’ΔHygR + 5’ΔHygR::Intron5'::Read1::GTACTTATAAGGTGA::Read2::Intron3'::3’ΔHygR + 5’ΔHygR::Intron5'::Read1::TGCATGACAGAGGGT::Read2::Intron3'::3’ΔHygR + 5’ΔHygR::Intron5'::Read1::AGACTTATACTGCGC::Read2::Intron3'::3’ΔHygR + 5’ΔHygR::Intron5'::Read1::ATTCTCAGAGGGGGA::Read2::Intron3'::3’ΔHygR + 5’ΔHygR::Intron5'::Read1::TGGCCCTCTCAGATC::Read2::Intron3'::3’ΔHygR + 5’ΔHygR::Intron5'::Read1::TGGTTAATACCGGCC::Read2::Intron3'::3’ΔHygR + 5’ΔHygR::Intron5'::Read1::TTACACTATTAACGG::Read2::Intron3'::3’ΔHygR + 5’ΔHygR::Intron5'::Read1::ATCCTTTGTAAACGA::Read2::Intron3'::3’ΔHygR + 5’ΔHygR::Intron5'::Read1::CACCATTTTCATCGA::Read2::Intron3'::3’ΔHygR + 5’ΔHygR::Intron5'::Read1::ATGCAGTTTTAATCG::Read2::Intron3'::3’ΔHygR + 5’ΔHygR::Intron5'::Read1::CACCCCTATAATCGA::Read2::Intron3'::3’ΔHygR + 5’ΔHygR::Intron5'::Read1::TCAGTTACATAGTTG::Read2::Intron3'::3’ΔHygR + 5’ΔHygR::Intron5'::Read1::AGATTAATAACGGGG::Read2::Intron3'::3’ΔHygR + 5’ΔHygR::Intron5'::Read1::CATCCATATTAGTTA::Read2::Intron3'::3’ΔHygR + 5’ΔHygR::Intron5'::Read1::TGGCTCTCTTATTCC::Read2::Intron3'::3’ΔHygR + 5’ΔHygR::Intron5'::Read1::CCTCTCTGTTAACCA::Read2::Intron3'::3’ΔHygR + 5’ΔHygR::Intron5'::Read1::TGTCGATGTAATTAG::Read2::Intron3'::3’ΔHygR + 5’ΔHygR::Intron5'::Read1::GTGCTATCTCACTAT::Read2::Intron3'::3’ΔHygR + 5’ΔHygR::Intron5'::Read1::TATCGTTCTGACAGA::Read2::Intron3'::3’ΔHygR + 5’ΔHygR::Intron5'::Read1::TACCTCTTTTAATCC::Read2::Intron3'::3’ΔHygR + 5’ΔHygR::Intron5'::Read1::AGTCCATATGATTCA::Read2::Intron3'::3’ΔHygR + 5’ΔHygR::Intron5'::Read1::CTACCATCTTATCTC::Read2::Intron3'::3’ΔHygR +

5’ΔHygR::Intron5'::Read1::TCACATTCTCACTAC::Read2::Intron3'::3’ΔHygR + 5’ΔHygR::Intron5'::Read1::TAGCCCTATTACCTT::Read2::Intron3'::3’ΔHygR + 5’ΔHygR::Intron5'::Read1::CATCCCTCTAACTCA::Read2::Intron3'::3’ΔHygR + 5’ΔHygR::Intron5'::Read1::TTTCCTTTTTAGAAC::Read2::Intron3'::3’ΔHygR + 5’ΔHygR::Intron5'::Read1::AAACTATTTTACCAT::Read2::Intron3'::3’ΔHygR + 5’ΔHygR::Intron5'::Read1::TCTCGTTTTGAGATC::Read2::Intron3'::3’ΔHygR + 5’ΔHygR::Intron5'::Read1::ACCCCATCTCATCAC::Read2::Intron3'::3’ΔHygR + 5’ΔHygR::Intron5'::Read1::TACCTATATTAGGCG::Read2::Intron3'::3’ΔHygR + 5’ΔHygR::Intron5'::Read1::ACGCGTGTAATCTTA::Read2::Intron3'::3’ΔHygR + 5’ΔHygR::Intron5'::Read1::AGAGTTACAAGGATG::Read2::Intron3'::3’ΔHygR + 5’ΔHygR::Intron5'::Read1::ATACGTTATGATCTC::Read2::Intron3'::3’ΔHygR + 5’ΔHygR::Intron5'::Read1::CGCCTCTCTCAGTTT::Read2::Intron3'::3’ΔHygR + 5’ΔHygR::Intron5'::Read1::TGAATGAAATAGAGC::Read2::Intron3'::3’ΔHygR + 5’ΔHygR::Intron5'::Read1::AAGCTTTTTGATAGT::Read2::Intron3'::3’ΔHygR + 5’ΔHygR::Intron5'::Read1::CAACGCTCTTACTCA::Read2::Intron3'::3’ΔHygR + 5’ΔHygR::Intron5'::Read1::TACCAGTATCACCCA::Read2::Intron3'::3’ΔHygR + 5’ΔHygR::Intron5'::Read1::TATCCCTTTTACGCT::Read2::Intron3'::3’ΔHygR + 5’ΔHygR::Intron5'::Read1::TTTCATTGTAAGCTC::Read2::Intron3'::3’ΔHygR + 5’ΔHygR::Intron5'::Read1::GAACAGTTTCAAATT::Read2::Intron3'::3’ΔHygR + 5’ΔHygR::Intron5'::Read1::ATGCTATCTCAGGGT::Read2::Intron3'::3’ΔHygR + 5’ΔHygR::Intron5'::Read1::TCCCTGTTTTACTTA::Read2::Intron3'::3’ΔHygR + 5’ΔHygR::Intron5'::Read1::TTTCCGTCTGATAGT::Read2::Intron3'::3’ΔHygR + 5’ΔHygR::Intron5'::Read1::TACCTTTTTAAGTTT::Read2::Intron3'::3’ΔHygR + 5’ΔHygR::Intron5'::Read1::GACCTTTGTTAGTCC::Read2::Intron3'::3’ΔHygR + 5’ΔHygR::Intron5'::Read1::GAGCCATATGATTCA::Read2::Intron3'::3’ΔHygR + 5’ΔHygR::Intron5'::Read1::TCCCCATGTCACTCA::Read2::Intron3'::3’ΔHygR + 5’ΔHygR::Intron5'::Read1::CCGCTGTCTTAAATG::Read2::Intron3'::3’ΔHygR + 5’ΔHygR::Intron5'::Read1::TATCGGTTTCAGTCA::Read2::Intron3'::3’ΔHygR + 5’ΔHygR::Intron5'::Read1::GCTCATTCTGACGTA::Read2::Intron3'::3’ΔHygR + 5’ΔHygR::Intron5'::Read1::ACACTATTTCATCTC::Read2::Intron3'::3’ΔHygR + 5’ΔHygR::Intron5'::Read1::GCACCATCTGATGCC::Read2::Intron3'::3’ΔHygR + 5’ΔHygR::Intron5'::Read1::TAGACATTTTCCTGT::Read2::Intron3'::3’ΔHygR + 5’ΔHygR::Intron5'::Read1::ACCCGGTATAACTTC::Read2::Intron3'::3’ΔHygR + 5’ΔHygR::Intron5'::Read1::TTGCTCTCTTATCTG::Read2::Intron3'::3’ΔHygR + 5’ΔHygR::Intron5'::Read1::ATACATTCTTACACT::Read2::Intron3'::3’ΔHygR +hsp-16.41p::Cas9(dpiRNA)::tbb-2 3’ UTR + prsp-27::NeoR::unc- 54 3' UTR + U6p:: GCGAAGTGACGGTAGACCGT]; fxSi47[ prsp-0p::5’ΔHygR:: GCGAAGTGACGGTAGACCGT ::3’ΔHygR::unc-54 3’::LoxP]

PX818 profile 1 fxEx27[5’ΔHygR::Intron5'::Read1::CGACAGTATGACGCC::Read2::Intron3'::3’ΔHygR + 5’ΔHygR::Intron5'::Read1::AACCGCTTTGACGCC::Read2::Intron3'::3’ΔHygR + 5’ΔHygR::Intron5'::Read1::TACCGATGTTACATC::Read2::Intron3'::3’ΔHygR + 5’ΔHygR::Intron5'::Read1::CTGCAGTGTTAGGCC::Read2::Intron3'::3’ΔHygR + 5’ΔHygR::Intron5'::Read1::GCACGCTCTGATATT::Read2::Intron3'::3’ΔHygR + 5’ΔHygR::Intron5'::Read1::ATACATTTTCAGCGC::Read2::Intron3'::3’ΔHygR + 5’ΔHygR::Intron5'::Read1::CTGCAGTGTCAACCC::Read2::Intron3'::3’ΔHygR + 5’ΔHygR::Intron5'::Read1::ATCCTATGTAAATTC::Read2::Intron3'::3’ΔHygR + 5’ΔHygR::Intron5'::Read1::GTTCTATATTACGCT::Read2::Intron3'::3’ΔHygR + 5’ΔHygR::Intron5'::Read1::GCGCTGTGTGATCAA::Read2::Intron3'::3’ΔHygR + 5’ΔHygR::Intron5'::Read1::CTGCGTTTTCATCAA::Read2::Intron3'::3’ΔHygR + 5’ΔHygR::Intron5'::Read1::CACCGCTTTTAACTT::Read2::Intron3'::3’ΔHygR + 5’ΔHygR::Intron5'::Read1::GTTCACTATAAGTCT::Read2::Intron3'::3’ΔHygR + 5’ΔHygR::Intron5'::Read1::GACCTCTCTAATATC::Read2::Intron3'::3’ΔHygR + 5’ΔHygR::Intron5'::Read1::CCCCTGTTTAAAGAT::Read2::Intron3'::3’ΔHygR + 5’ΔHygR::Intron5'::Read1::AGACGGTTTTAACTC::Read2::Intron3'::3’ΔHygR + 5’ΔHygR::Intron5'::Read1::TACCTATATTAGGCG::Read2::Intron3'::3’ΔHygR + 5’ΔHygR::Intron5'::Read1::TAACTTTGTAATTCT::Read2::Intron3'::3’ΔHygR + 5’ΔHygR::Intron5'::Read1::CTCCGTTGTCAATTA::Read2::Intron3'::3’ΔHygR + 5’ΔHygR::Intron5'::Read1::TATCCCTATGACTAT::Read2::Intron3'::3’ΔHygR + 5’ΔHygR::Intron5'::Read1::CAGCTTTTTAAAGAT::Read2::Intron3'::3’ΔHygR + 5’ΔHygR::Intron5'::Read1::GCGCTGAAAATGTAT::Read2::Intron3'::3’ΔHygR + 5’ΔHygR::Intron5'::Read1::CCTCCATCTCATGCT::Read2::Intron3'::3’ΔHygR + 5’ΔHygR::Intron5'::Read1::GCACGTTGTCATCTT::Read2::Intron3'::3’ΔHygR + 5’ΔHygR::Intron5'::Read1::ATTCCATCTTACCAC::Read2::Intron3'::3’ΔHygR + 5’ΔHygR::Intron5'::Read1::GGCCTAACACTGCAG::Read2::Intron3'::3’ΔHygR + 5’ΔHygR::Intron5'::Read1::CGCCTAATATAGGTA::Read2::Intron3'::3’ΔHygR + 5’ΔHygR::Intron5'::Read1::CATCTTGTTCCCATC::Read2::Intron3'::3’ΔHygR + 5’ΔHygR::Intron5'::Read1::AATATCAGAGCGTGC::Read2::Intron3'::3’ΔHygR + 5’ΔHygR::Intron5'::Read1::GAATTTACATAGGAT::Read2::Intron3'::3’ΔHygR + 5’ΔHygR::Intron5'::Read1::AGACTTATAGTGAAC::Read2::Intron3'::3’ΔHygR +

5’ΔHygR::Intron5'::Read1::CACCTATTTCACAAC::Read2::Intron3'::3’ΔHygR + 5’ΔHygR::Intron5'::Read1::GGCGTCATACTGTCG::Read2::Intron3'::3’ΔHygR + 5’ΔHygR::Intron5'::Read1::TTTCATTCTCATATC::Read2::Intron3'::3’ΔHygR + 5’ΔHygR::Intron5'::Read1::AGCGTAATATAGAAC::Read2::Intron3'::3’ΔHygR + 5’ΔHygR::Intron5'::Read1::GATATTAGAGAGGTC::Read2::Intron3'::3’ΔHygR + 5’ΔHygR::Intron5'::Read1::GTCGTGCAAGTTTGA::Read2::Intron3'::3’ΔHygR + 5’ΔHygR::Intron5'::Read1::TAGACATTTTCCTGT::Read2::Intron3'::3’ΔHygR + 5’ΔHygR::Intron5'::Read1::CGCCAATTTAACTAC::Read2::Intron3'::3’ΔHygR + 5’ΔHygR::Intron5'::Read1::ATCTTTAAAAAGCTG::Read2::Intron3'::3’ΔHygR + 5’ΔHygR::Intron5'::Read1::GTATGTGTTATTTTT::Read2::Intron3'::3’ΔHygR + 5’ΔHygR::Intron5'::Read1::TAATTGACAACGGAG::Read2::Intron3'::3’ΔHygR + 5’ΔHygR::Intron5'::Read1::GATGTAACATCGGTA::Read2::Intron3'::3’ΔHygR + 5’ΔHygR::Intron5'::Read1::TAAGTTATATCGTAC::Read2::Intron3'::3’ΔHygR + 5’ΔHygR::Intron5'::Read1::AATAGTGTCACATGA::Read2::Intron3'::3’ΔHygR + 5’ΔHygR::Intron5'::Read1::TACCTTCTGTCTGTT::Read2::Intron3'::3’ΔHygR + 5’ΔHygR::Intron5'::Read1::CAGCAGTGTCAACAA::Read2::Intron3'::3’ΔHygR + 5’ΔHygR::Intron5'::Read1::CATAGTTTTCCTTAT::Read2::Intron3'::3’ΔHygR + 5’ΔHygR::Intron5'::Read1::CTCCCTTATCATTGC::Read2::Intron3'::3’ΔHygR + 5’ΔHygR::Intron5'::Read1::ACCCGCTTTTAGTTA::Read2::Intron3'::3’ΔHygR + 5’ΔHygR::Intron5'::Read1::ATAGTCATAGGGATA::Read2::Intron3'::3’ΔHygR + 5’ΔHygR::Intron5'::Read1::CCGCTGTCTTAAATG::Read2::Intron3'::3’ΔHygR + 5’ΔHygR::Intron5'::Read1::GGCGTCAAAGCGGTT::Read2::Intron3'::3’ΔHygR + 5’ΔHygR::Intron5'::Read1::AACTCTATGGCACAG::Read2::Intron3'::3’ΔHygR + 5’ΔHygR::Intron5'::Read1::AACCTTTATCATGGA::Read2::Intron3'::3’ΔHygR + 5’ΔHygR::Intron5'::Read1::ACACTATTTCATCTC::Read2::Intron3'::3’ΔHygR + 5’ΔHygR::Intron5'::Read1::ACTCGGTTTCAACCA::Read2::Intron3'::3’ΔHygR + 5’ΔHygR::Intron5'::Read1::AGAATTACAAAGTTA::Read2::Intron3'::3’ΔHygR + 5’ΔHygR::Intron5'::Read1::TTGATGAAAACGCAG::Read2::Intron3'::3’ΔHygR + 5’ΔHygR::Intron5'::Read1::GGGTTGACACTGCAG::Read2::Intron3'::3’ΔHygR + 5’ΔHygR::Intron5'::Read1::TACCCCTTTTAAACT::Read2::Intron3'::3’ΔHygR + 5’ΔHygR::Intron5'::Read1::ACACTCTATCACCCC::Read2::Intron3'::3’ΔHygR + 5’ΔHygR::Intron5'::Read1::ACCCGGTATAACTTC::Read2::Intron3'::3’ΔHygR + 5’ΔHygR::Intron5'::Read1::GAGCAATTTAATCAT::Read2::Intron3'::3’ΔHygR + 5’ΔHygR::Intron5'::Read1::ATACATTCTTACACT::Read2::Intron3'::3’ΔHygR + 5’ΔHygR::Intron5'::Read1::CACCCCTGTGACTAC::Read2::Intron3'::3’ΔHygR + 5’ΔHygR::Intron5'::Read1::CCTCTGTATTATCCC::Read2::Intron3'::3’ΔHygR + 5’ΔHygR::Intron5'::Read1::TATCTTTGTAACATA::Read2::Intron3'::3’ΔHygR + 5’ΔHygR::Intron5'::Read1::CACTTTCTTTCTTAC::Read2::Intron3'::3’ΔHygR + 5’ΔHygR::Intron5'::Read1::GATAATATACCTAGT::Read2::Intron3'::3’ΔHygR + 5’ΔHygR::Intron5'::Read1::GCTCATTCTGACGTA::Read2::Intron3'::3’ΔHygR + 5’ΔHygR::Intron5'::Read1::TTGATCACACAGCGC::Read2::Intron3'::3’ΔHygR + 5’ΔHygR::Intron5'::Read1::CTTCAGTGTCAATGT::Read2::Intron3'::3’ΔHygR + 5’ΔHygR::Intron5'::Read1::TAACGGTATGAATAA::Read2::Intron3'::3’ΔHygR + 5’ΔHygR::Intron5'::Read1::TATCGGTTTCAGTCA::Read2::Intron3'::3’ΔHygR + 5’ΔHygR::Intron5'::Read1::TCGCGTTCTCATTCA::Read2::Intron3'::3’ΔHygR + 5’ΔHygR::Intron5'::Read1::AACCGATTTAAAAAA::Read2::Intron3'::3’ΔHygR + 5’ΔHygR::Intron5'::Read1::CGCCATTCTTAATTT::Read2::Intron3'::3’ΔHygR + 5’ΔHygR::Intron5'::Read1::GAGAGTTGTAATTCA::Read2::Intron3'::3’ΔHygR + 5’ΔHygR::Intron5'::Read1::ATCTTTAAACAGGGG::Read2::Intron3'::3’ΔHygR + 5’ΔHygR::Intron5'::Read1::AAACCATCTAAAAAA::Read2::Intron3'::3’ΔHygR + 5’ΔHygR::Intron5'::Read1::CTTCAATATTACGCA::Read2::Intron3'::3’ΔHygR + 5’ΔHygR::Intron5'::Read1::AACGCCTTCCAAAGG::Read2::Intron3'::3’ΔHygR + 5’ΔHygR::Intron5'::Read1::CACCACTCTAATAAC::Read2::Intron3'::3’ΔHygR + 5’ΔHygR::Intron5'::Read1::AAGTTAAAAGCGGTG::Read2::Intron3'::3’ΔHygR + 5’ΔHygR::Intron5'::Read1::CCTCGTTCTGATCCA::Read2::Intron3'::3’ΔHygR + 5’ΔHygR::Intron5'::Read1::CGCCCATATTACCTA::Read2::Intron3'::3’ΔHygR + 5’ΔHygR::Intron5'::Read1::GCACCATCTGATGCC::Read2::Intron3'::3’ΔHygR + 5’ΔHygR::Intron5'::Read1::TCCCCATGTCACTCA::Read2::Intron3'::3’ΔHygR + 5’ΔHygR::Intron5'::Read1::TGCCTGTGTAATCTC::Read2::Intron3'::3’ΔHygR + 5’ΔHygR::Intron5'::Read1::GAACATTTGAATTTG::Read2::Intron3'::3’ΔHygR +hsp-16.41p::Cas9(dpiRNA)::tbb-2 3’ UTR + prsp-27::NeoR::unc- 54 3' UTR + U6p:: GCGAAGTGACGGTAGACCGT]; fxSi47[ prsp-0p::5’ΔHygR:: GCGAAGTGACGGTAGACCGT ::3’ΔHygR::unc-54 3’::LoxP]

PX818 profile 2 fxEx28[5’ΔHygR::Intron5'::Read1::GCACGTTGTCATCTT::Read2::Intron3'::3’ΔHygR + 5’ΔHygR::Intron5'::Read1::CCTCCATCTCATGCT::Read2::Intron3'::3’ΔHygR + 5’ΔHygR::Intron5'::Read1::ATTCCATCTTACCAC::Read2::Intron3'::3’ΔHygR + 5’ΔHygR::Intron5'::Read1::TTTCATTCTCATATC::Read2::Intron3'::3’ΔHygR + 5’ΔHygR::Intron5'::Read1::AAGATGACAACGTGC::Read2::Intron3'::3’ΔHygR + 5’ΔHygR::Intron5'::Read1::CCACCATCTCATCAT::Read2::Intron3'::3’ΔHygR +

5’ΔHygR::Intron5'::Read1::AAACCATCTAAACAA::Read2::Intron3'::3’ΔHygR + 5’ΔHygR::Intron5'::Read1::GATATGAGAATGAAA::Read2::Intron3'::3’ΔHygR + 5’ΔHygR::Intron5'::Read1::GTGGTAAGATGGAAT::Read2::Intron3'::3’ΔHygR + 5’ΔHygR::Intron5'::Read1::AGCATGAGATGGAGG::Read2::Intron3'::3’ΔHygR + 5’ΔHygR::Intron5'::Read1::TAACATTCTCATATA::Read2::Intron3'::3’ΔHygR + 5’ΔHygR::Intron5'::Read1::TACCGATGTTACATC::Read2::Intron3'::3’ΔHygR + 5’ΔHygR::Intron5'::Read1::GTATGTGTTATTTTT::Read2::Intron3'::3’ΔHygR + 5’ΔHygR::Intron5'::Read1::TAGACATTTTCCTGT::Read2::Intron3'::3’ΔHygR + 5’ΔHygR::Intron5'::Read1::GCACGCTCTGATATT::Read2::Intron3'::3’ΔHygR + 5’ΔHygR::Intron5'::Read1::CTGCAGTGTTAGGCC::Read2::Intron3'::3’ΔHygR + 5’ΔHygR::Intron5'::Read1::AACCGCTTTGACGCC::Read2::Intron3'::3’ΔHygR + 5’ΔHygR::Intron5'::Read1::CTGCAGTGTCAACCC::Read2::Intron3'::3’ΔHygR + 5’ΔHygR::Intron5'::Read1::ATACATTTTCAGCGC::Read2::Intron3'::3’ΔHygR + 5’ΔHygR::Intron5'::Read1::CCCCTGTTTAAAGAT::Read2::Intron3'::3’ΔHygR + 5’ΔHygR::Intron5'::Read1::GCGCTGTGTGATCAA::Read2::Intron3'::3’ΔHygR + 5’ΔHygR::Intron5'::Read1::GTTCTATATTACGCT::Read2::Intron3'::3’ΔHygR + 5’ΔHygR::Intron5'::Read1::GTCGTGCAAGTTTGA::Read2::Intron3'::3’ΔHygR + 5’ΔHygR::Intron5'::Read1::CGACAGTATGACGCC::Read2::Intron3'::3’ΔHygR + 5’ΔHygR::Intron5'::Read1::CACCGCTTTTAACTT::Read2::Intron3'::3’ΔHygR + 5’ΔHygR::Intron5'::Read1::CTGCGTTTTCATCAA::Read2::Intron3'::3’ΔHygR + 5’ΔHygR::Intron5'::Read1::GTTCACTATAAGTCT::Read2::Intron3'::3’ΔHygR + 5’ΔHygR::Intron5'::Read1::AGACGGTTTTAACTC::Read2::Intron3'::3’ΔHygR + 5’ΔHygR::Intron5'::Read1::ATCCTATGTAAATTC::Read2::Intron3'::3’ΔHygR + 5’ΔHygR::Intron5'::Read1::CTCCCTTATCATTGC::Read2::Intron3'::3’ΔHygR + 5’ΔHygR::Intron5'::Read1::TAACTTTGTAATTCT::Read2::Intron3'::3’ΔHygR + 5’ΔHygR::Intron5'::Read1::TACCTATATTAGGCG::Read2::Intron3'::3’ΔHygR + 5’ΔHygR::Intron5'::Read1::TTTCCTTGTCAGATT::Read2::Intron3'::3’ΔHygR + 5’ΔHygR::Intron5'::Read1::GCACCATCTGATGCC::Read2::Intron3'::3’ΔHygR + 5’ΔHygR::Intron5'::Read1::TGCCCTTGTCATCCT::Read2::Intron3'::3’ΔHygR + 5’ΔHygR::Intron5'::Read1::GACCTCTCTAATATC::Read2::Intron3'::3’ΔHygR + 5’ΔHygR::Intron5'::Read1::TATCGGTTTCAGTCA::Read2::Intron3'::3’ΔHygR + 5’ΔHygR::Intron5'::Read1::AACCTTTATCATGGA::Read2::Intron3'::3’ΔHygR + 5’ΔHygR::Intron5'::Read1::ACTCGGTTTCAACCA::Read2::Intron3'::3’ΔHygR + 5’ΔHygR::Intron5'::Read1::TAACGGTATGAATAA::Read2::Intron3'::3’ΔHygR + 5’ΔHygR::Intron5'::Read1::AGTCTATCTCAGCCT::Read2::Intron3'::3’ΔHygR + 5’ΔHygR::Intron5'::Read1::CTTCTGTGTGATCAC::Read2::Intron3'::3’ΔHygR + 5’ΔHygR::Intron5'::Read1::GAGCAATTTAATCAT::Read2::Intron3'::3’ΔHygR + 5’ΔHygR::Intron5'::Read1::GCACACTATAACATT::Read2::Intron3'::3’ΔHygR + 5’ΔHygR::Intron5'::Read1::TGCCCATCTGAAAAG::Read2::Intron3'::3’ΔHygR + 5’ΔHygR::Intron5'::Read1::CAGCTTTTTAAAGAT::Read2::Intron3'::3’ΔHygR + 5’ΔHygR::Intron5'::Read1::CCACGGTCTCACAGT::Read2::Intron3'::3’ΔHygR + 5’ΔHygR::Intron5'::Read1::GAGAGTTGTAATTCA::Read2::Intron3'::3’ΔHygR + 5’ΔHygR::Intron5'::Read1::CCTCTGTATTATCCC::Read2::Intron3'::3’ΔHygR + 5’ΔHygR::Intron5'::Read1::AAACCATCTAACCAA::Read2::Intron3'::3’ΔHygR + 5’ΔHygR::Intron5'::Read1::CGCCAATTTAACTAC::Read2::Intron3'::3’ΔHygR + 5’ΔHygR::Intron5'::Read1::AATCCATCTAAACAA::Read2::Intron3'::3’ΔHygR + 5’ΔHygR::Intron5'::Read1::CGCCATTCTTAATTT::Read2::Intron3'::3’ΔHygR + 5’ΔHygR::Intron5'::Read1::CACCTATTTCACAAC::Read2::Intron3'::3’ΔHygR + 5’ΔHygR::Intron5'::Read1::TACCAATTTGACCAT::Read2::Intron3'::3’ΔHygR + 5’ΔHygR::Intron5'::Read1::TTGCTCTCTTATCTG::Read2::Intron3'::3’ΔHygR + 5’ΔHygR::Intron5'::Read1::ATACATTCTTACACT::Read2::Intron3'::3’ΔHygR + 5’ΔHygR::Intron5'::Read1::TACCCCTTTTAAACT::Read2::Intron3'::3’ΔHygR + 5’ΔHygR::Intron5'::Read1::TTTCACTATCAATCC::Read2::Intron3'::3’ΔHygR + 5’ΔHygR::Intron5'::Read1::TCGCGTTCTCATTCA::Read2::Intron3'::3’ΔHygR + 5’ΔHygR::Intron5'::Read1::AGCAGGAGATGGAGG::Read2::Intron3'::3’ΔHygR + 5’ΔHygR::Intron5'::Read1::TTCCTTTTTAACCCT::Read2::Intron3'::3’ΔHygR + 5’ΔHygR::Intron5'::Read1::AACCATTTTCACTAT::Read2::Intron3'::3’ΔHygR + 5’ΔHygR::Intron5'::Read1::ACCCGGTATAACTTC::Read2::Intron3'::3’ΔHygR + 5’ΔHygR::Intron5'::Read1::TAACAATTTCATCGT::Read2::Intron3'::3’ΔHygR + 5’ΔHygR::Intron5'::Read1::CTCCGTTGTCAATTA::Read2::Intron3'::3’ΔHygR + 5’ΔHygR::Intron5'::Read1::GTCCTGTTTCATCCC::Read2::Intron3'::3’ΔHygR + 5’ΔHygR::Intron5'::Read1::TATCTTTGTAACATA::Read2::Intron3'::3’ΔHygR + 5’ΔHygR::Intron5'::Read1::CCGCATTTTTATCCT::Read2::Intron3'::3’ΔHygR + 5’ΔHygR::Intron5'::Read1::GGTCAATTTTAACTT::Read2::Intron3'::3’ΔHygR + 5’ΔHygR::Intron5'::Read1::TCTCCGTATTACCTT::Read2::Intron3'::3’ΔHygR + 5’ΔHygR::Intron5'::Read1::AACGCCTTCCAAAGG::Read2::Intron3'::3’ΔHygR + 5’ΔHygR::Intron5'::Read1::ACACTATTTCATCTC::Read2::Intron3'::3’ΔHygR + 5’ΔHygR::Intron5'::Read1::ACCCGTTTTAACTCT::Read2::Intron3'::3’ΔHygR + 5’ΔHygR::Intron5'::Read1::CACCCCTGTGACTAC::Read2::Intron3'::3’ΔHygR + 5’ΔHygR::Intron5'::Read1::CATCTATTTTAACTC::Read2::Intron3'::3’ΔHygR +

5’ΔHygR::Intron5'::Read1::CCACCATTTTATTTG::Read2::Intron3'::3’ΔHygR + 5’ΔHygR::Intron5'::Read1::TATCCCTATGACTAT::Read2::Intron3'::3’ΔHygR + 5’ΔHygR::Intron5'::Read1::TGACCTTTTCAAAGA::Read2::Intron3'::3’ΔHygR + 5’ΔHygR::Intron5'::Read1::TGCCAGTCTCAATAA::Read2::Intron3'::3’ΔHygR + 5’ΔHygR::Intron5'::Read1::TTACCATGTTAGCCA::Read2::Intron3'::3’ΔHygR + 5’ΔHygR::Intron5'::Read1::CGCCCATATTACCTA::Read2::Intron3'::3’ΔHygR + 5’ΔHygR::Intron5'::Read1::CGTCACTATGATTAA::Read2::Intron3'::3’ΔHygR + 5’ΔHygR::Intron5'::Read1::GAACATTTGAATTTG::Read2::Intron3'::3’ΔHygR + 5’ΔHygR::Intron5'::Read1::GCTCATTCTGACGTA::Read2::Intron3'::3’ΔHygR + 5’ΔHygR::Intron5'::Read1::TACGTTTTTTCGAAG::Read2::Intron3'::3’ΔHygR + 5’ΔHygR::Intron5'::Read1::TCACATTCTGACACA::Read2::Intron3'::3’ΔHygR + 5’ΔHygR::Intron5'::Read1::TCCCCATGTCACTCA::Read2::Intron3'::3’ΔHygR + 5’ΔHygR::Intron5'::Read1::TTTCAATGTTATCAA::Read2::Intron3'::3’ΔHygR + 5’ΔHygR::Intron5'::Read1::TTTCCCTTTGACTTC::Read2::Intron3'::3’ΔHygR + 5’ΔHygR::Intron5'::Read1::ACACTCTATCACCCC::Read2::Intron3'::3’ΔHygR + 5’ΔHygR::Intron5'::Read1::AGGCTTTGTCAAACG::Read2::Intron3'::3’ΔHygR + 5’ΔHygR::Intron5'::Read1::CAACACTATGACTTT::Read2::Intron3'::3’ΔHygR + 5’ΔHygR::Intron5'::Read1::CGTCTGTTTTAGTAC::Read2::Intron3'::3’ΔHygR + 5’ΔHygR::Intron5'::Read1::CTGCCGTGTTACATC::Read2::Intron3'::3’ΔHygR + 5’ΔHygR::Intron5'::Read1::GTTCCATATCAGTTT::Read2::Intron3'::3’ΔHygR + 5’ΔHygR::Intron5'::Read1::GTTCTTTTTGAACCA::Read2::Intron3'::3’ΔHygR + 5’ΔHygR::Intron5'::Read1::TCCCTCTCTAATCAT::Read2::Intron3'::3’ΔHygR + 5’ΔHygR::Intron5'::Read1::TTACACTATCATTCG::Read2::Intron3'::3’ΔHygR + 5’ΔHygR::Intron5'::Read1::CACCATTTTAAACGA::Read2::Intron3'::3’ΔHygR + 5’ΔHygR::Intron5'::Read1::CATGGTATGACCACT::Read2::Intron3'::3’ΔHygR + 5’ΔHygR::Intron5'::Read1::CCACCTTTTCACTCG::Read2::Intron3'::3’ΔHygR + 5’ΔHygR::Intron5'::Read1::CGTCCATCTTATCAA::Read2::Intron3'::3’ΔHygR + 5’ΔHygR::Intron5'::Read1::GAACGGTTGTAAGTT::Read2::Intron3'::3’ΔHygR + 5’ΔHygR::Intron5'::Read1::GGACTATTTGACCTC::Read2::Intron3'::3’ΔHygR + 5’ΔHygR::Intron5'::Read1::TAGCAATGTTACGAA::Read2::Intron3'::3’ΔHygR + 5’ΔHygR::Intron5'::Read1::TCCCGTTTTCATCCA::Read2::Intron3'::3’ΔHygR + 5’ΔHygR::Intron5'::Read1::TCCCTCTATAAGTCT::Read2::Intron3'::3’ΔHygR + 5’ΔHygR::Intron5'::Read1::TCGCTATGTAACGTC::Read2::Intron3'::3’ΔHygR + 5’ΔHygR::Intron5'::Read1::TTGCCTTCTCAGGTG::Read2::Intron3'::3’ΔHygR + 5’ΔHygR::Intron5'::Read1::ACCCGCTTTTAGTTA::Read2::Intron3'::3’ΔHygR + 5’ΔHygR::Intron5'::Read1::AGGCAATGTAACGAC::Read2::Intron3'::3’ΔHygR + 5’ΔHygR::Intron5'::Read1::AGTCGCTTTTATAGC::Read2::Intron3'::3’ΔHygR + 5’ΔHygR::Intron5'::Read1::CAAGTTATGACCCTT::Read2::Intron3'::3’ΔHygR + 5’ΔHygR::Intron5'::Read1::CAAGTTCTTACATCT::Read2::Intron3'::3’ΔHygR + 5’ΔHygR::Intron5'::Read1::CACCGGTCTCAACCA::Read2::Intron3'::3’ΔHygR + 5’ΔHygR::Intron5'::Read1::CATCCGTATGAAGTT::Read2::Intron3'::3’ΔHygR + 5’ΔHygR::Intron5'::Read1::CTACAATCTAACCGT::Read2::Intron3'::3’ΔHygR + 5’ΔHygR::Intron5'::Read1::CTACCCTTTTAACCT::Read2::Intron3'::3’ΔHygR + 5’ΔHygR::Intron5'::Read1::GAATCTGTCGCTGGT::Read2::Intron3'::3’ΔHygR + 5’ΔHygR::Intron5'::Read1::GTCCACTCTAATTCA::Read2::Intron3'::3’ΔHygR + 5’ΔHygR::Intron5'::Read1::GTCCGCTCTAACATC::Read2::Intron3'::3’ΔHygR + 5’ΔHygR::Intron5'::Read1::TAATGTGTTTCTCCC::Read2::Intron3'::3’ΔHygR + 5’ΔHygR::Intron5'::Read1::TTACATTCTTATCCG::Read2::Intron3'::3’ΔHygR + 5’ΔHygR::Intron5'::Read1::TTGCAATATCAGTTG::Read2::Intron3'::3’ΔHygR + 5’ΔHygR::Intron5'::Read1::CAACGGTTTCAGCAT::Read2::Intron3'::3’ΔHygR + 5’ΔHygR::Intron5'::Read1::CATCCTCTAACCTTG::Read2::Intron3'::3’ΔHygR + 5’ΔHygR::Intron5'::Read1::CATTATTTCTCGGCC::Read2::Intron3'::3’ΔHygR + 5’ΔHygR::Intron5'::Read1::CCACATTCTAATACG::Read2::Intron3'::3’ΔHygR + 5’ΔHygR::Intron5'::Read1::GGTCGGTCTCAAGAT::Read2::Intron3'::3’ΔHygR + 5’ΔHygR::Intron5'::Read1::TCCCTATTTTATACA::Read2::Intron3'::3’ΔHygR + 5’ΔHygR::Intron5'::Read1::TCGCGCTTTAATTCT::Read2::Intron3'::3’ΔHygR + 5’ΔHygR::Intron5'::Read1::TGCCCTTTTCAGTAC::Read2::Intron3'::3’ΔHygR + 5’ΔHygR::Intron5'::Read1::AACGCTATACCGATC::Read2::Intron3'::3’ΔHygR + 5’ΔHygR::Intron5'::Read1::AAGGCTGTTGCGTCG::Read2::Intron3'::3’ΔHygR + 5’ΔHygR::Intron5'::Read1::ATTCGTTCTGATTTG::Read2::Intron3'::3’ΔHygR + 5’ΔHygR::Intron5'::Read1::CCGCTGTCTTAAATG::Read2::Intron3'::3’ΔHygR + 5’ΔHygR::Intron5'::Read1::GACCAGTCTCACTTG::Read2::Intron3'::3’ΔHygR + 5’ΔHygR::Intron5'::Read1::GCACATTCTAATACA::Read2::Intron3'::3’ΔHygR + 5’ΔHygR::Intron5'::Read1::GCCCGGTATCACTCC::Read2::Intron3'::3’ΔHygR + 5’ΔHygR::Intron5'::Read1::ACTAGACCTCGCTGC::Read2::Intron3'::3’ΔHygR + 5’ΔHygR::Intron5'::Read1::CTGGTACCGTAGGGT::Read2::Intron3'::3’ΔHygR + 5’ΔHygR::Intron5'::Read1::TAGCGCTGTGATATT::Read2::Intron3'::3’ΔHygR + 5’ΔHygR::Intron5'::Read1::TCGCTGTATGAGGAT::Read2::Intron3'::3’ΔHygR + 5’ΔHygR::Intron5'::Read1::AATCAATCTCATAAA::Read2::Intron3'::3’ΔHygR + 5’ΔHygR::Intron5'::Read1::AGCCTGTCTCAACAG::Read2::Intron3'::3’ΔHygR +

5’ΔHygR::Intron5'::Read1::CAGCCATCTCAATTC::Read2::Intron3'::3’ΔHygR + 5’ΔHygR::Intron5'::Read1::CATCTTGTTCCCATC::Read2::Intron3'::3’ΔHygR + 5’ΔHygR::Intron5'::Read1::GATCTTTCTCAACAT::Read2::Intron3'::3’ΔHygR + 5’ΔHygR::Intron5'::Read1::CCACCCTATCATGCC::Read2::Intron3'::3’ΔHygR + 5’ΔHygR::Intron5'::Read1::GTGCCGTATCAACTC::Read2::Intron3'::3’ΔHygR + 5’ΔHygR::Intron5'::Read1::TTTCGCTTTTACCCT::Read2::Intron3'::3’ΔHygR + 5’ΔHygR::Intron5'::Read1::AAACGATGTCAATTA::Read2::Intron3'::3’ΔHygR + 5’ΔHygR::Intron5'::Read1::ACTCTATGTCATCTA::Read2::Intron3'::3’ΔHygR + 5’ΔHygR::Intron5'::Read1::ATGCCATATTACCCC::Read2::Intron3'::3’ΔHygR + 5’ΔHygR::Intron5'::Read1::CTTCCATCTCATCAC::Read2::Intron3'::3’ΔHygR + 5’ΔHygR::Intron5'::Read1::GACCCATTTTACTAC::Read2::Intron3'::3’ΔHygR + 5’ΔHygR::Intron5'::Read1::GCACGTTTTAACATA::Read2::Intron3'::3’ΔHygR + 5’ΔHygR::Intron5'::Read1::TAAGTTATATCGTAC::Read2::Intron3'::3’ΔHygR + 5’ΔHygR::Intron5'::Read1::AGTCCTTTTGACGTA::Read2::Intron3'::3’ΔHygR + 5’ΔHygR::Intron5'::Read1::CCGCAGTTTCAATCA::Read2::Intron3'::3’ΔHygR + 5’ΔHygR::Intron5'::Read1::TTTCCATGTCACATC::Read2::Intron3'::3’ΔHygR + 5’ΔHygR::Intron5'::Read1::CTCCGCTATTACAAA::Read2::Intron3'::3’ΔHygR + 5’ΔHygR::Intron5'::Read1::GGACATTTTGAATTT::Read2::Intron3'::3’ΔHygR + 5’ΔHygR::Intron5'::Read1::GGCCTATTTAACTTA::Read2::Intron3'::3’ΔHygR + 5’ΔHygR::Intron5'::Read1::TAGCTCTCTCATCCT::Read2::Intron3'::3’ΔHygR + 5’ΔHygR::Intron5'::Read1::TCTCCTTATAAAGGC::Read2::Intron3'::3’ΔHygR + 5’ΔHygR::Intron5'::Read1::TCTCTTTGTTATCAC::Read2::Intron3'::3’ΔHygR + 5’ΔHygR::Intron5'::Read1::AAGCCCTATAATACC::Read2::Intron3'::3’ΔHygR + 5’ΔHygR::Intron5'::Read1::CTTCGCTGTCATATT::Read2::Intron3'::3’ΔHygR + 5’ΔHygR::Intron5'::Read1::TAGGCTTTAACAAAT::Read2::Intron3'::3’ΔHygR + 5’ΔHygR::Intron5'::Read1::AAGCCTATTCCTCCT::Read2::Intron3'::3’ΔHygR + 5’ΔHygR::Intron5'::Read1::ACACCTTATGATAGA::Read2::Intron3'::3’ΔHygR + 5’ΔHygR::Intron5'::Read1::CGGGTTATTGACACA::Read2::Intron3'::3’ΔHygR + 5’ΔHygR::Intron5'::Read1::GCACACTCTGAGACC::Read2::Intron3'::3’ΔHygR + 5’ΔHygR::Intron5'::Read1::GCCCGTTCTCATTGA::Read2::Intron3'::3’ΔHygR + 5’ΔHygR::Intron5'::Read1::TAACTGTCTGAATAC::Read2::Intron3'::3’ΔHygR + 5’ΔHygR::Intron5'::Read1::TGCCAATCTTATCCA::Read2::Intron3'::3’ΔHygR + 5’ΔHygR::Intron5'::Read1::TTAAATTATCACATG::Read2::Intron3'::3’ΔHygR + 5’ΔHygR::Intron5'::Read1::TTTCCTGGTGGTTGT::Read2::Intron3'::3’ΔHygR + 5’ΔHygR::Intron5'::Read1::CTACACTCTCATGTT::Read2::Intron3'::3’ΔHygR + 5’ΔHygR::Intron5'::Read1::CTGCAATGTCACTAA::Read2::Intron3'::3’ΔHygR + 5’ΔHygR::Intron5'::Read1::GACGATGTGTCTCAT::Read2::Intron3'::3’ΔHygR + 5’ΔHygR::Intron5'::Read1::AATAGTGTCACATGA::Read2::Intron3'::3’ΔHygR + 5’ΔHygR::Intron5'::Read1::AACTCTATGGCACAG::Read2::Intron3'::3’ΔHygR + 5’ΔHygR::Intron5'::Read1::CATAGTTTTCCTTAT::Read2::Intron3'::3’ΔHygR + 5’ΔHygR::Intron5'::Read1::TACCTTCTGTCTGTT::Read2::Intron3'::3’ΔHygR + 5’ΔHygR::Intron5'::Read1::CTACCCTATCATCAC::Read2::Intron3'::3’ΔHygR + 5’ΔHygR::Intron5'::Read1::CTTAGCGTGCTATGT::Read2::Intron3'::3’ΔHygR + 5’ΔHygR::Intron5'::Read1::GATAATATACCTAGT::Read2::Intron3'::3’ΔHygR + 5’ΔHygR::Intron5'::Read1::GGGTGAACGGCGATG::Read2::Intron3'::3’ΔHygR + 5’ΔHygR::Intron5'::Read1::GTAGGCGCACTCATG::Read2::Intron3'::3’ΔHygR + 5’ΔHygR::Intron5'::Read1::TCCCTGATGGATTTC::Read2::Intron3'::3’ΔHygR + 5’ΔHygR::Intron5'::Read1::TCGCAATTTCAATTT::Read2::Intron3'::3’ΔHygR + 5’ΔHygR::Intron5'::Read1::TATCGTTCTAAACGA::Read2::Intron3'::3’ΔHygR + 5’ΔHygR::Intron5'::Read1::TCGCGATGTCATACC::Read2::Intron3'::3’ΔHygR + 5’ΔHygR::Intron5'::Read1::AATCAGTATAACAGC::Read2::Intron3'::3’ΔHygR + 5’ΔHygR::Intron5'::Read1::TGGTGTTTGATTCTG::Read2::Intron3'::3’ΔHygR + 5’ΔHygR::Intron5'::Read1::TTCTCGCTTTTAGGT::Read2::Intron3'::3’ΔHygR + 5’ΔHygR::Intron5'::Read1::AAGCGACTCGACAAG::Read2::Intron3'::3’ΔHygR + 5’ΔHygR::Intron5'::Read1::CGACTGTCTTAAACC::Read2::Intron3'::3’ΔHygR + 5’ΔHygR::Intron5'::Read1::GTACCGTTTAACGTC::Read2::Intron3'::3’ΔHygR + 5’ΔHygR::Intron5'::Read1::TGTACCAACCCATTG::Read2::Intron3'::3’ΔHygR + 5’ΔHygR::Intron5'::Read1::TTTTGCACAATTCAT::Read2::Intron3'::3’ΔHygR +hsp-16.41p::Cas9(dpiRNA)::tbb-2 3’ UTR + prsp-27::NeoR::unc- 54 3' UTR + U6p:: GCGAAGTGACGGTAGACCGT]; fxSi47[ prsp-0p::5’ΔHygR:: GCGAAGTGACGGTAGACCGT ::3’ΔHygR::unc-54 3’::LoxP]
