## Supplementary figures and images for "High-Throughput Library Transgenesis in *Caenorhabditis elegans* via Transgenic Arrays Resulting in Diversity of Integrated Sequences (TARDIS)"

### Figure5_S1A_unedited.tif

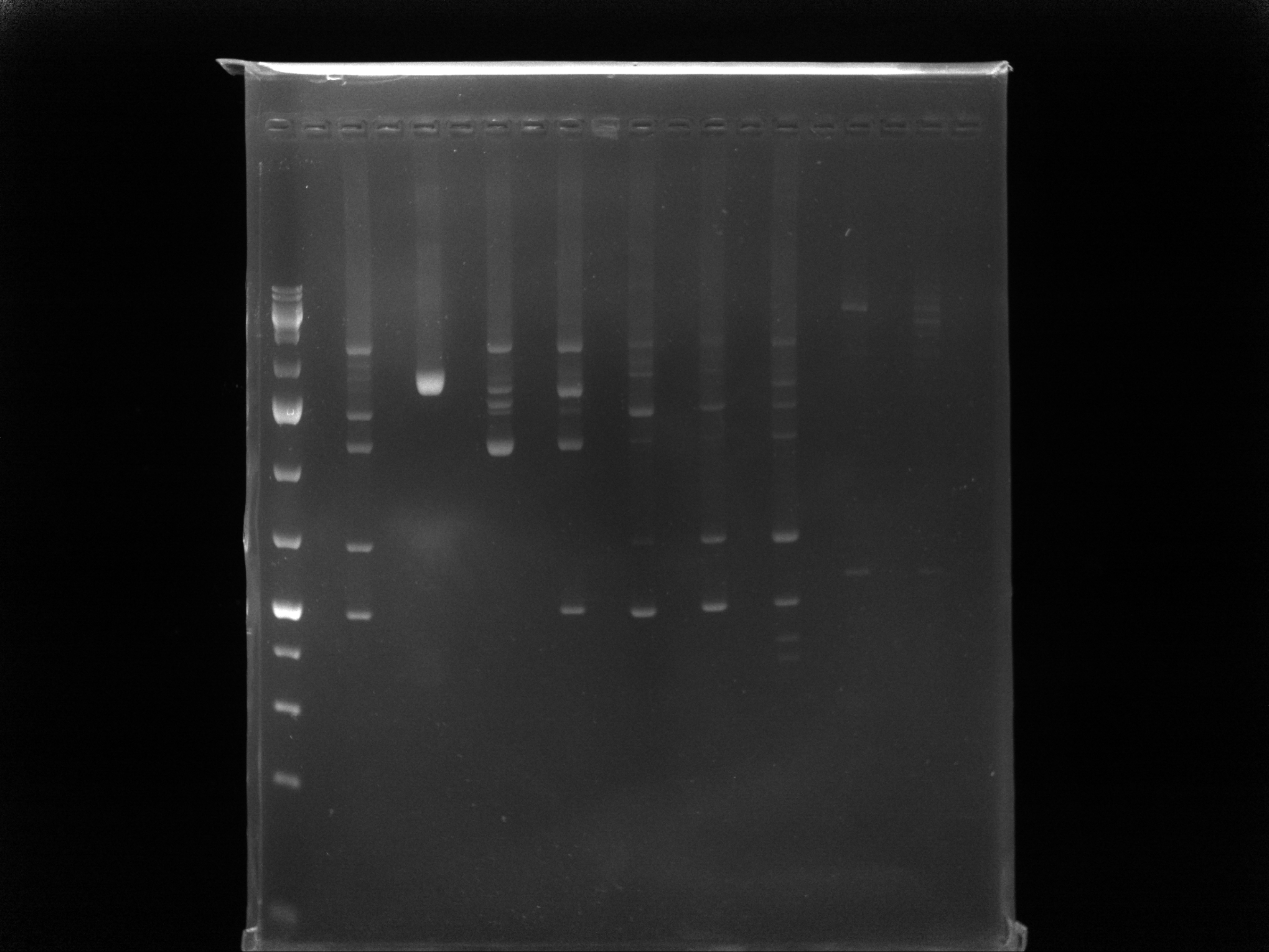

### Figure5_S1C_unedited.tif

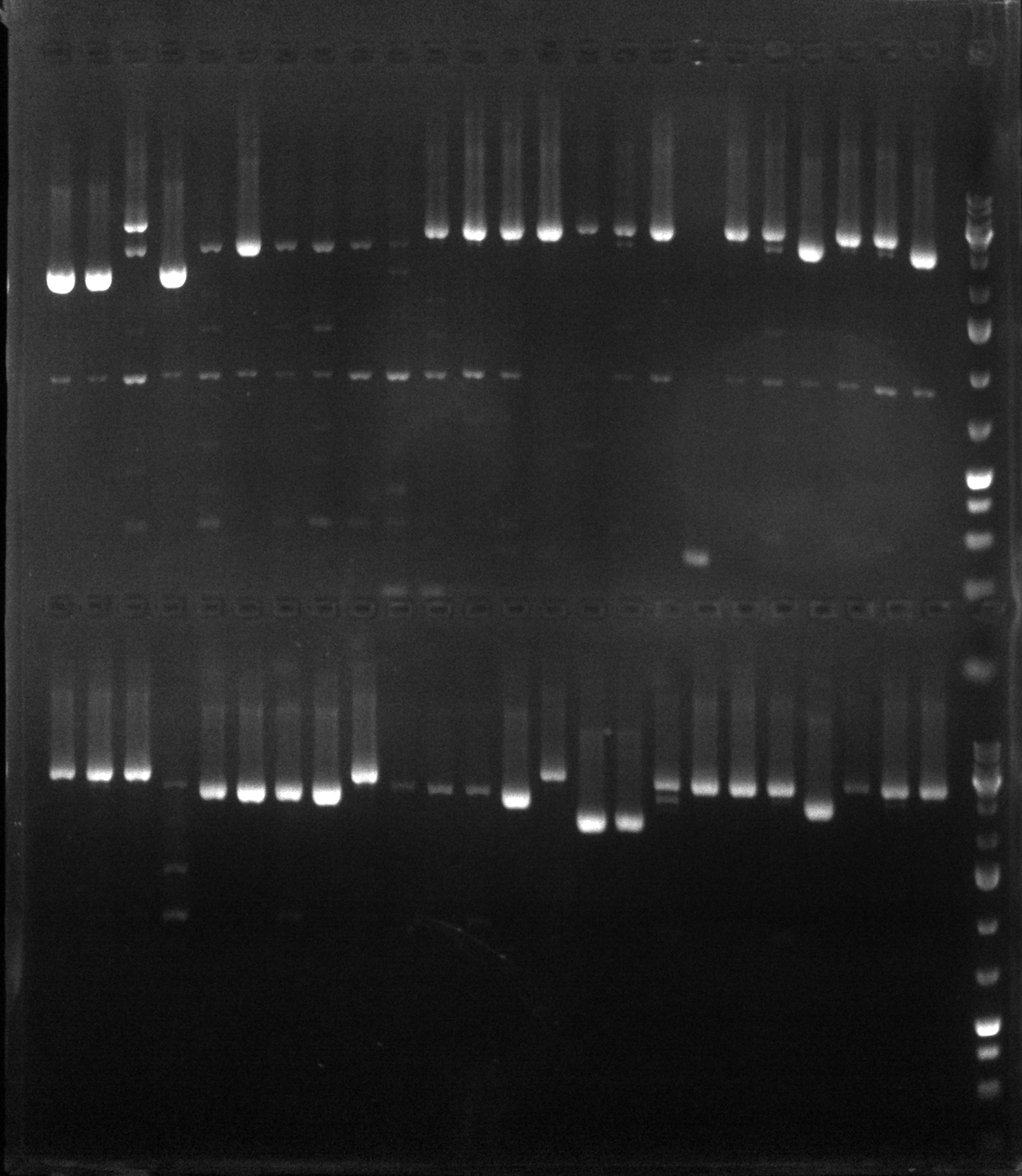
